# Seasonal succession of nano- and picoplankton communities in Lake Constance: conserved dynamics despite compositional shifts under contrasting mixing and oxygen regimes

**DOI:** 10.64898/2026.06.24.733152

**Authors:** Corentin Fournier, David Schleheck

## Abstract

Lake Constance is a pre-Alpine, monomictic, oligotrophic lake situated at the southern end of Germany composed of two main water bodies: deep, oligotrophic Upper Lake Constance (ULC) and the shallow, more mesotrophic Lower Lake Constance (LLC). To date, no sequencing-based study exists of the seasonal succession of the microbial plankton in Lake Constance. Over one-year, microbial plankton communities were sampled biweekly from the top 20 m of the water column in both sites and separated into nanoplankton (NP) and picoplankton (PP). Communities were analysed using rDNA amplicon sequencing: NP samples were analysed by 18S rDNA, and PP samples by 18S and 16S rDNA sequencing. Temporal community diversity was compared between sites and the effect of two major environmental perturbations, winter vertical mixing in ULC and oxygen depletion of the bottom-water layer in LLC, on the community was examined. Despite strong environmental contrasts, microbial plankton communities exhibited conserved seasonal temporal dynamics across basins. In contrast, pronounced compositional shifts occurred during mixing and oxygen depletion events. Approximately 20% of detected taxa were positively associated with these events, with log fold changes reaching 9.82, reflecting rare or undetectable taxa outside these periods. Taxa favoured by these perturbations commonly exhibited high metabolic flexibility, including mixotrophy, fermentation, or anaerobic respiration, or possessed functional traits conferring tolerance to altered redox and mixing regimes. Our results suggest that the temporal dynamics of freshwater microbial plankton communities are driven by deterministic processes and highlight the profound impact of large, and less known, environmental changes on these communities.

## Introduction

Lake Constance is a large pre-Alpine, warm-monomictic, oligotrophic lake located at the southern end of Germany and composed of two connected basins: the deep, oligotrophic Upper Lake Constance (ULC) and the shallower, more mesotrophic Lower Lake Constance (LLC). Thanks to decades of extensive monitoring (Straile and Geller 1998; Sabel et al. 2020), Lake Constance has become a model system in limnology. For example, Sommer and colleagues used data generated at Lake Constance to develop the Plankton Ecology Group (PEG) model, now used as a standard template to describe patterns of seasonal plankton succession in oligotrophic and eutrophic lakes. (Sommer et al. 2012). While this work has allowed for a detailed understanding of the phytoplankton and zooplankton dynamics, it relies mostly on microscope observation, with no use of more recent techniques like sequencing. Furthermore, very few studies have focused on Lake Constance bacterioplankton communities, and sequencing based studies are very recent (Zwisler et al. 2003; Fournier et al. 2021, 2024).

Traditional microscopy-based approaches to study plankton communities are limited in their ability to reveal the complete community diversity, (Rodriguez et al. 2002; Naselli-Flores 2014; Xiao et al. 2014) and cannot be applied to study heterotrophic pico- and bacterioplankton communities. Molecular fingerprinting techniques have been the initial means to study bacterioplankton community diversity, providing the first insight into Lake Constance bacterioplankton community composition. (Zwisler et al. 2003). The advent of next-generation sequencing, particularly amplicon sequencing targeting the 16S and 18S rDNA taxonomic markers, now enables more detailed characterisation of bacterial and micro-eukaryotic communities across plankton size classes. These approaches have become essential for understanding freshwater ecosystem structure and functioning. (Newton et al. 2007; Salmaso et al. 2018, 2020).

The microbial plankton community plays a central role in aquatic ecosystems by sustaining primary production, regulating oxygen availability, and mediating nutrient recycling within food webs, all of which maintain the stability of aquatic ecosystems (Cole et al. 1988; Lewis et al. 2020). In the context of accelerating environmental changes and anthropogenic disturbances establishing molecular baselines of these communities is increasingly important. It is therefore becoming urgent to create plankton DNA-sample repositories for Lake Constance to study its present microbial plankton community, which could then serve as a reference for comparison with future changes.

We started routine sampling of the total DNA of the nano- and picoplankton community of Lake Constance every two weeks and established a DNA bank for the year 2018 of these two plankton size classes. Integrated water sampling from 0 – 20 m was chosen as an approximate representation of the photic zone (epilimnion), that represents the highest primary production. Sampling was carried out in two sites of the lake, the Überlingersee basin of Upper Lake Constance (in the following abbreviated ULC), and Zellersee basin of Lower Lake Constance (abbreviated LLC), which offer marked differences in environmental conditions. The ULC site at Überlingersee is deep (up to 145 m), with a water column yearly oxic and oligotrophic, and undergo a vertical mixing during the cold period. The LLC is shallow, with a water depth of approx. 22 m at the sampling site, has a water column less oligotrophic, and presents a strong oxycline from mid-August to early November, resulting in depletion of oxygen at the deepest layer. Both the winter water vertical mixing and the depletion of oxygen lead to drastic and sudden changes in environmental conditions (Diehl et al. 2002; Carey 2023; Limnologischer Zustand des Bodensees)

This dataset allows us to study how the temporal dynamics and community composition of the microbial plankton differ between two sites of Lake Constance. The following questions were addressed. Despite being characterised by different environmental conditions, does the compositional diversity of the communities in these two sites follow the same pattern of temporal dynamics? The winter water vertical mixing and the oxygen depletion affect the whole water column and deeply and suddenly modify the environmental conditions. Are there specific microbial taxa positively selected, or is it randomly distributed in the communities, and in what proportion? This study establishes a high-resolution, molecular reference for Lake Constance microbial plankton and provides, for the first time, insights into the temporal and spatial dynamics of microbial plankton communities in a well-characterised freshwater ecosystem.

## Materials and Methods

### Study sites, sampling campaign and filtration procedure

No permits were required for accessing the sampling sites and for the water sampling. Samples were taken at the routine sampling site Wallhausen in the Überlinger See (47.7571°N, 9.1273°E), a north-western arm of ULC with a maximal depth of 140 m, and in Zeller See (47.716557 °N, 8.991767 °E), a shallow north-western part of LLC with a maximal depth of about 20 m. We sampled the approximate photic zone (epilimnion), which represents the 0 – 20 m depth surface layer. Integrated samples were collected on a boat using an integrating water sampler (Model IWS II, Hydro-Bios, Germany). The sampling campaign started in March 2018 and is still ongoing. This study presents results based on samples from March 2018 to March 2019. Samples were taken every two weeks and are stored in stainless steel barrels (19 L soda-kegs). Table S1 summarises the sampling dates and their metadata. Four biological replicates were collected each time (n = 4).

Filtrations were done directly on the boat in order to reduce alteration of the samples. A pressure of 2 bars was applied in the barrels, using a pressurized air tank. The barrels were connected to a series of filter holders (Swinnex 47 mm filter holder; Millipore) using PVC tubing. The pore-sizes of the filters were: 180 µm (hydrophilic nylon net, 47 mm diameter; Millipore) to remove zooplankton and larger particles, and 5.0 µm to collect the nanoplankton, and 0.2 µm to collect the picoplankton, each using polycarbonate membrane filters (Isopore, 47 mm diameter; Millipore). The Swinnex filter holders, with the filters assembled, were wrapped in aluminium foil, autoclaved and then transported onto the boat. Filtration was done following the protocol developed in our laboratory (Fournier et al. 2024). Each filter was transferred by using sterile forceps and stored when being rolled into 15 ml Falcon tubes, with the filter side harbouring the collected biomass facing to the inside of the tube. The tubes were stored on dry ice on the boat and in the laboratory at -20°C until the DNA was extracted. Only the 5.0 and 0.2 µm filters were used for the downstream analysis.

### Molecular work: DNA extraction, library preparation and sequencing

The DNA extraction protocol used was adapted from Rusch et al. 2007 (Rusch et al. 2007) and from the JGI protocol. Extraction started by thawing the filters and adding 3 ml of lysis buffer, composed of EDTA, EGTA and Tris-HCl [pH 8.0], each at a final concentration of 50 mM. 1 min of ultrasonication (Sonorex Super RK 510, Bandelin, Germany) was applied, followed by adding 0.1- and 1-mm diameter zirconium beads (approximately 200 µg of each) and 15 min of vortexing at highest speed; the tubes were held vertically during vortexing. Lysozyme (2.5 mg/ml) was added, and the tubes incubated for 1 h at 37°C in a shaking incubator. Sodium dodecyl sulphate (SDS) at a final concentration of 1% and proteinase K (500 µg/ml) were added, followed by 2 h of incubation at 55°C. After one hour of incubation, proteinase K was added again at the same concentration. A final step was done consisting of 10 min incubation at 65°C with 0.07 vol. of 5M NaCl and 0.07 vol. of CTAB/NaCl (preheated at 65°C). DNA was then purified by adding 1 vol. of phenol/chloroform/isoamyalcohol, followed by 20 min of centrifugation at 13,000 RCF and 4°C. After recovering the supernatant in a new 15 ml Falcon tube, 1 vol. of chloroform/isoamyalcohol was added, followed by 20 min of centrifugation at 13,000 rcf and 4°C. The supernatant was recovered in a new 15 ml Falcon tube. 0.5 µl of glycogen (Thermo Fisher Scientific, USA) and 0.7 vol. of isopropanol were added to the tube, followed by incubation for 15 min at 20°C. Another centrifugation was done for 25 min at 15,000 RCF and 4°C. The isopropanol was decanted, and the DNA pellet was washed using 500 µl ice-cold 70% ethanol. After a final centrifugation of 5 min, the ethanol was removed and the pellet was dried in air for approx. 5 min. Then, 50 µl of PCR-grade water was added to each tube. Finally, the DNA concentration was measured using a Nanodrop 2000c spectrophotometer (Thermo Fisher Scientific, USA), and the DNA quality examined after agarose gel electrophoresis.

The 180 – 5 µm size class nanoplankton (NP) was studied using 18S rRNA gene-fragment amplification, targeting the eukaryotic nanoplankton community (18S-NP). The 5 – 0.2 µm size class picoplankton (PP) was studied using both 16S and 18S rRNA gene-fragment amplification, targeting respectively the eukaryotic picoplankton (18S-PP) and the bacterioplankton (16S-PP). A two-step PCR approach was used for the DNA library preparation with Illumina dual indexing. The primer pairs for the first PCR amplified the DNA fragment of interest and included a universal 5’-sequence tail. Amplification of the V3-V5 hypervariable region of the 16S rRNA gene (569 bp) was performed using the 357F and 926R primers, respectively (Schuurman et al. 2004; Walters et al. 2016). Amplification of the V4 hypervariable region of the 18S rRNA gene (378 bp) was done using the 573F and 951R primers (Mangot et al. 2013). Table 1 includes the sequences of the universal 5’-tail and the rDNA primer sequences. PCR was performed with 0.02 U/µl of Phusion High Fidelity DNA polymerase, 1X Phusion HF buffer, and 200 µM of dNTPs (New England Biolabs, USA). Extracted DNA was added at a final concentration of 0.12 ng/µl. A final concentration of 0.5 µM of each primer was used. The number of PCR cycles was 10. It comprised the initial denaturation step of 3 min at 98°C, the denaturation of 45 s at 98°C, primer annealing step of 20 s at 62.4°C, extension step of 8 s at 72°C, and the final extension step of 5 min at 72°C. The reactions were done using a T100 Thermal Cycler (Bio-Rad, USA). PCR products of the first PCR were used as template for the second PCR with Illumina adapters. The sets of primer pairs used included an individual index (barcode sequence) specific to each sample and the Illumina adapters. The same conditions were used as with the first PCR, but the number of cycles was 20. PCR products of both steps were each purified following the Agencourt AMPure XP protocol, with a 0.9 and 0.8 vol. beads, and a final elution volume of 11 μl and 14 μl for the first and second purification, respectively. PCR products were pooled in equivalent quantities and sent to Eurofins GATC Biotech for sequencing using Illumina MiSeq 2*300 bp with the Microbiome Profiling Indexing Only-package. Each amplification, 16S and 18S DNA amplicon, was sequenced independently in duplicate.

**Table 1:**
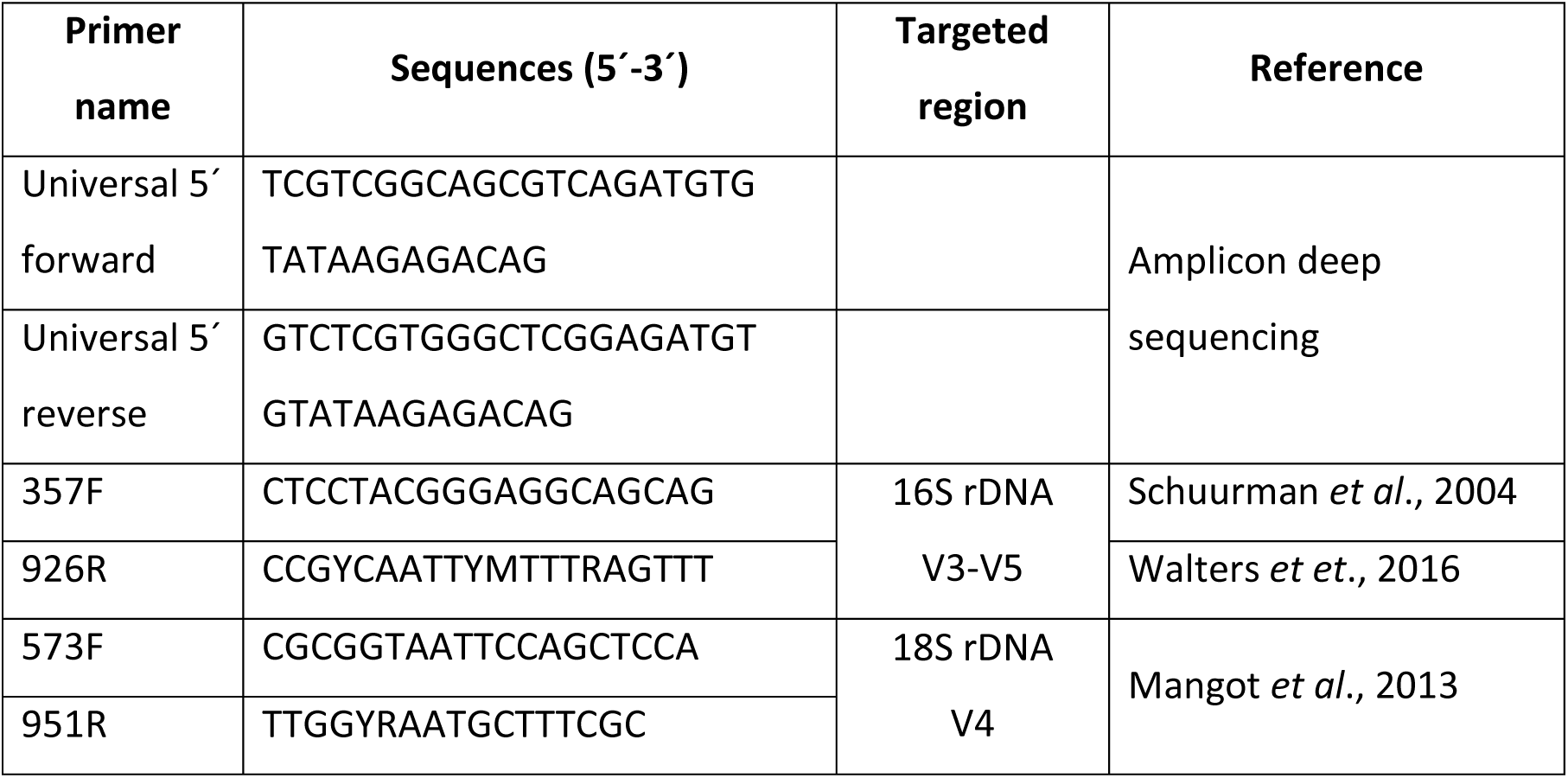
Sequence and targeted region of the primers used for the amplification of the 16S and 18S rRNA gene for barcoded amplicon sequencing library preparation.

### Bioinformatic pipeline

The sequencing data provided by Eurofins-GATC Biotech were paired-end. Quality of the sequences was checked using FastQC (Simon 2010). The poor quality of the 16S rDNA reverse reads made the merging of the paired end reads not reliable. Therefore, the analysis of the 16S rDNA dataset was done using only the forward reads. Reads were trimmed using Trimmomatic (Bolger et al. 2014), discarding reads with a Phred score below 3 at the start and the end of the reads, and an average quality of 11 over a window of 3 bases within the reads. The 18S rDNA reads were also cropped at a maximum size of 285 to remove bases with lowest quality in the merging area. The 18S rDNA paired end reads were merged using NGmerge with parameters of a minimum overlap of 30 bases and an acceptable error rate of maximum 10% (Gaspar 2018). Read quality was checked between each of the previously described steps. Denoising and taxonomic affiliation were then performed in QIIME2 2020.11. Filtration of chimeras using the consensus method, denoising, and dereplication of the quality reads were done using the denoise and dereplicate single-end sequences (DADA2, denoise-single) and a read learn of 2,000,000 reads for the training error model (Callahan et al. 2016). For the 16S rDNA database, taxonomic affiliation was done using the classify-consensus VSEARCH program and the TaxAss pipeline. (Rohwer et al. 2018), which used both a general database, SILVA_138, and a freshwater ecosystem-specific database, FreshTrain (Newton et al. 2011). Four thresholds of identity were used for the taxonomic assignment: 80%, 90%, 97%, and 100%. Outputs were then compared to check for consistency, and to keep the deepest taxonomic affiliation while having a minimum of unassigned sequences. The 18S rDNA taxonomic affiliation was performed using Blast+ with the top 5 hits, Megablast, on the SSU_eukaryote_rRNA reference database. After checking the consistency of the top-5 hits, the top one was kept and its Unique Subject Taxonomy ID (STAXIDs) was linked to the full lineage using a custom reference database: the reference database was created from the NCBI archive new_taxdump using NCBItax2lin. A parallel taxonomic affiliation was performed using the classify-consensus VSEARCH program with an 80% identity threshold and the reference database PR2 (Protist Ribosomal Database). Both taxonomies were compared to verify the consistency of the taxonomy. Raw datasets are available under the bioproject number PRJNA1478098.

### Biostatistics

The dataset was split by sampling site (ULC and LLC) and plankton type (18S-NP, 18S-PP, and 16S-PP), followed by the filtration of the lowest abundant ASVs. ASVs represented by less than 0.01% in relative abundance in less than 5% of the samples were discarded. ASV affiliated to Mitochondria and Chloroplast for the 16S rDNA datasets, and ASV affiliated to Metazoa for the 18S rDNA datasets, were also discarded. No rarefaction to a minimum number of reads for all samples was applied on the datasets. Samples with a number of reads below 8907, threshold chosen based on the rarefaction curve, were discarded.

All statistical analyses were performed on the R software (R core team 2021) using primarily the package Phyloseq (McMurdie and Holmes 2013) and vegan (Oksanen et al. 2015). The Shannon-Wiener and Simpson diversity indices were used to observe the alpha diversity, with Observed ASVs and the Pielou index as a complement for richness and evenness. The beta diversity was visualised by Principal Component Analysis (PCA) based on the PhilR distance (phylogenetic isometric log-ratio transform) (Silverman et al. 2021). Ellipses were calculated following a multivariate t-distribution, with a confidence level of 0.95. Correlations between distance matrices (ULC versus LLC) were performed using a Procrustes analysis and Mantel test. The Procrustes analysis is a statistical shape analysis which compares the geometry of two datasets, here the temporal dynamics of the microbial plankton community in ULC and LLC, while the Mantel test evaluates the correlation between two dissimilarity matrices. Permutational multivariate analysis of variance (PermANOVA) with 999 permutations and Analysis of Similarity (ANOSIM) were used to test the difference in the community composition. Homogeneity of variance between groups was analysed using betadisper, which calculates the average distance to the group median. PermANOVA, ANOSIM, and betadisper tests were all performed on the PhilR distance matrix. P-value threshold for significance was set to 0.05. The pseudo-F value and R2 of the PermANOVA can be interpreted as the effect size and the percentage of variance explained by the tested groups, respectively. The ANOSIM statistic, F-value, which ranges from 0 to 1, represents the similarity of the groups tested, with a high value representing a high degree of dissimilarity.

Analysis of the microbial plankton community presenting increased relative abundance during ULC winter vertical mixing and LLC oxygen depletion events was performed using the LinDA method (linear models for differential abundance analysis of microbiome compositional data) (Zhou et al. 2022). Count data were used to perform the analysis. The p-value was adjusted using the false discovery rate method of Benjamini & Hochberg. ASV differential abundances were tested comparing the winter water vertical mixing period versus water stratification for ULC and oxygen depleted bottom water layer versus full oxic water column. The adjusted p-value threshold was set to 0.01. ASVs were considered positively selected if its p-value was below the threshold, and the log fold change (logFC), representing the ratio of relative abundance between the conditions, was above 2. A logFC of 2 represents a minimum fourfold difference of abundance between the tested periods.

## Results

This study covered a full annual cycle from March 2018 to March 2019 and involved sampling at Upper Lake Constance (ULC) and Lower Lake Constance (LLC) every two weeks, for a total of 26 sampling dates. Microbial plankton biomass was collected and separated into nanoplankton (NP; 180–5 µm) and picoplankton (PP; 5–0.2 µm) size classes. Three biological replicates were used for the downstream analysis, resulting in 78 samples per dataset. For each site, three datasets were created: 18S rDNA amplicon sequencing of the NP and PP fractions (18S-NP and 18S-PP, respectively), and 16S rDNA amplicon sequencing of the PP fraction (16S-PP), totalling six datasets. The quality of the 16S rDNA reverse reads was too low for use in the analysis, so we only used the forward reads. Amplicons still comprise enough taxonomic information to cover the complete V3- and a large proportion of the V4-hypervariable region. Two independent Illumina paired-end sequencing runs were performed on each sample to examine variability introduced by sequencing and increase the number of reads by pooling them.

After filtering out low-quality samples, there were no time gaps in our datasets (Table S1). Sample numbers ranged from 70 to 78 per dataset. Two replicates from 10 April 2018 in the ULC 18S NP dataset and from 20 November 2018 in the LLC 16S-PP dataset were removed. Although no standard deviation could be calculated for the alpha diversity, this did not affect the downstream analysis, as the samples were clustered into larger groups (Table S1). Following bioinformatics processing and filtration of low-abundant ASVs, the final datasets comprised 2.15×10⁷ reads and a total of 5,416 ASVs across all datasets (Table S2).

### Plankton diversity dynamics

The first question addressed was whether there were differences in the temporal dynamics of microbial plankton community diversity between two sites of Lake Constance with different environmental conditions. To this extent, the alpha and beta diversity of the three communities examined, eukaryotic nano- (18S-NP) and picoplankton (18S-PP), and prokaryotic picoplankton (16S-PP), were analysed and compared.

The 18S-NP and 18S-PP communities displayed a similar alpha diversity pattern between ULC and LLC, peaking during the warmer period from spring to summer. The 18S LLC communities both showed an earlier increase and decrease in diversity than the ULC community (Fig. 1AB). This increase was due to a higher number of ASVs being detected during this time (Fig. S1AB). The fact that the Simpson index increased less intensely than the Shannon index indicated that the additional ASVs detected belonged to the less abundant community (Fig. 1AB and Fig. S2AB). High diversity was observed in the prokaryotic picoplankton communities (16S-PP) during winter for both ULC and LLC (Fig. 1C). These increases in diversity coincided with increased evenness (Fig. S3C), indicating a more even distribution of the relative abundances of the detected ASVs. The diversity of both ULC and LLC 16S-PP communities followed similar patterns, except during the mid-summer to late-autumn period when 16S-PP diversity was lowest in ULC and highest in LLC. This difference was due to the detection of 400–600 additional ASVs in the LLC community during this period (Fig. S1C).

**Figure 1:**
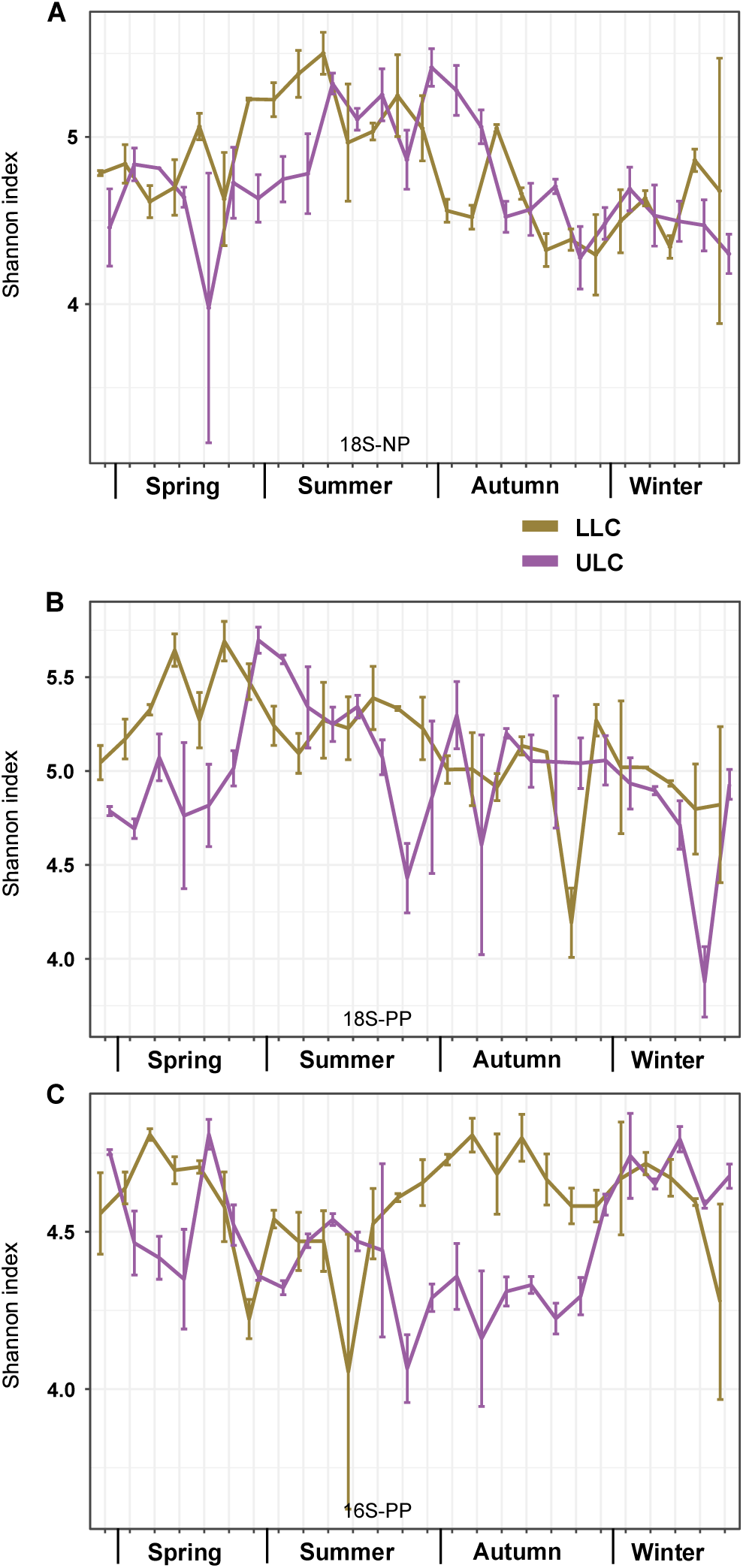
Shannon diversity calculated per sampling site and across the sampling dates. The purple lines correspond to the Upper Lake Constance (ULC) and the gold lines correspond to the Lower Lake Constance (LLC) sampling site. The upper panel (A) shows the indices for the eukaryotic nanoplankton (18S-NP dataset), the central panel (B) for the eukaryotic picoplankton (18S-PP dataset) and the lower panel (C) for the prokaryotic picoplankton (16S-PP dataset). The x-axis represents the different seasons (for exact sampling dates, please see Table S1). Note that the y-axes have different scales for each of the three graphs. Graphical representations of the Simpson indices are shown in the Supplemental information file (Figure S1) as well as for the richness (Figure S2) and evenness (Figure S3).

The biodiversity analysis (beta diversity) had two objectives. The first objective was to test whether the ULC and LLC communities exhibited similar seasonal diversity dynamics. A cyclic temporal dynamic was observed in all three plankton communities in both the ULC and the LLC (Fig. S4A–F). Procrustes analysis and Mantel tests confirmed the similarity of the temporal dynamics of the three plankton communities between ULC and LLC. The results of both tests consisted of correlation values, p-values and sums of squares for the Procrustes analysis. All p-values were below the chosen significance threshold of 0.05 and the correlation values ranged from medium to high, depending on the test. The sum of squares value for the Procrustes analysis was also low (Table 2 and Fig. S5). These results suggest that all three plankton communities follow similar temporal patterns of biodiversity in both the ULC and the LLC.

**Table 2:** Results of the Mantel tests and Procrutes analysis testing the correlation of the temporal patterns of plankton biodiversity change in ULC compared to LLC.

| Microbial plankton | Procrustes analysis |  |  | Mantel test |  |
| --- | --- | --- | --- | --- | --- |
|  | Correlation | Sum of squares | p-value | Correlation | p-value |
| **18S-NP** | 0.79 | 0.38 | <0.001 | 0.45 | <0.001 |
| **18S-PP** | 0.87 | 0.24 | <0.001 | 0.67 | <0.001 |
| **16S-PP** | 0.84 | 0.29 | <0.001 | 0.64 | <0.001 |

The second objective was to compare the biodiversity of the communities at the two sites to determine their level of similarity. To achieve this, distance matrices were calculated for each plankton type, combining the ULC and LLC samples, and visualised using PCA (Fig. 2). Excluding the samples representing the winter water vertical mixing and oxygen depletion events, the ULC and LLC 16S NP and -PP samples visually clustered well together (Fig. 2AB). The 16S-PP communities displayed clear visual differences in biodiversity between the two sites, with the water vertical mixing and oxygen depletion events clustering together and being more isolated (Fig. 2C). For 18S-NP, while less visible, the ULC 18S-PP and 16S-PP communities tend to align with the second PCA axis, while the LLC communities align with the first PCA axis. This indicates different drivers of biodiversity between the two sites. The PERMANOVA and ANOSIM results are summarised in Table 3. Based on the p-values, there were significant differences in all three plankton community compositions based on the sampling sites, which supports the visual observations of the 16S-PP communities. However, the 18S-NP and 18S-PP communities showed a low pseudo-F ratio and R² value for PERMANOVA, as well as a mild ANOSIM statistic F value (Table 3), indicating similarity in community biodiversity between the two sites. These conflicting results will be discussed in more detail. For all three communities, samples taken during ULC vertical mixing and LLC oxygen depletion in deep water were the most isolated and clustered (Fig. 2 and Fig. S6). These results suggest that these events strongly affect the microbial plankton communities, causing a sudden shift in composition. We therefore addressed the question of whether specific taxonomic groups were positively affected, and to what extent the microbial plankton community composition was modified.

**Figure 2:**
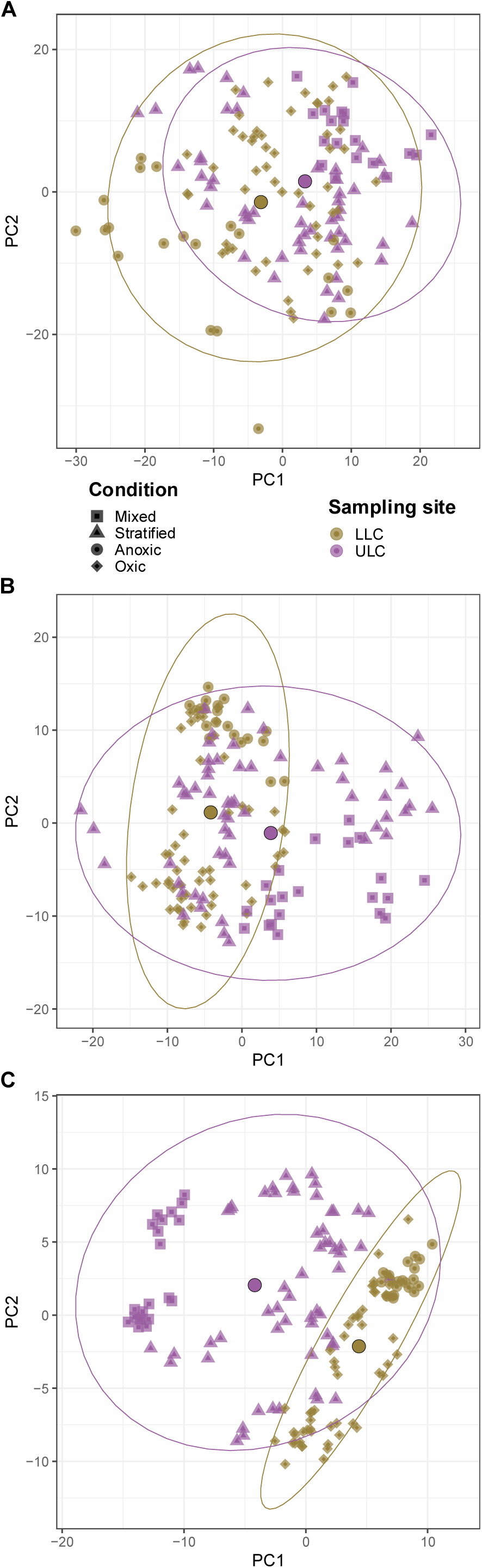
Principal Component Analysis of nano- and picoplankton biodiversity across the seasonal sampling of the different parts of the lake. Dissimilarity matrices calculated based on the phylogenetic isometric log-ratio (PhilR)-transformed Euclidean distance matrix. Colours indicate the sampling site; gold, LLC; purple, ULC. Shape indicates the different conditions observed in the water. Square represent ULC water vertical mixing samples and triangle the water stratified samples. Round represent LLC oxygen depleted bottom water samples and diamond full oxic column samples. The large circle with black outline corresponds to the centroid per sampling site. Ellipses were calculated following a multivariate t-distribution with a confidence level of 0.95 per sampling site. The top panel (A) shows the data for the eukaryotic nanoplankton (18S-NP), with the ellipses overlapping, indicating similar biodiversity dynamics in ULC and LLC and across the seasons. The central panel (B) shows the data for the eukaryotic picoplankton (18S-PP), indicating a certain separation of biodiversity change, even though the ellipses overlapped. The lower panel (C) shows the data for the prokaryotic picoplankton (16S-PP), indicating a clear separation of biodiversity change between ULC and LLC across the seasons. For all three communities, samples representing the ULC vertical mixing and LLC oxygen depletion event were the most isolated one.

**Table 3:** Results of PermANOVA and ANOSIM testing the biodiversity differences of the three plankton communities between ULC and LLC.

| Microbial plankton | PermANOVA |  |  | ANOSIM |  |
| --- | --- | --- | --- | --- | --- |
| | $R^2$ | Pseudo-F ratio | p-value | Statistic R | p-value |
| 18S-NP | 0.08 | 12.56 | <0.001 | 0.17 | <0.001 |
| 18S-PP | 0.09 | 14.83 | <0.001 | 0.25 | <0.001 |
| 16S-PP | 0.17 | 31.53 | <0.001 | 0.50 | <0.001 |

### Microbial plankton community affected by specific environmental conditions

The detection of taxa that were positively affected by vertical water mixing in the ULC and by oxygen depletion in the bottom water layer in the LLC was performed using linear models for differential abundance analysis (LinDA). These models analyse the differential relative abundance of ASVs in our datasets. Comparisons were made between the thermally stratified or mixed water column in the ULC and the full oxic water column or the oxygen-depleted bottom water layer in the LLC, with respect to the three plankton communities: 18S-NP, 18S-PP and 16S-PP. The outputs of the LinDA analysis are summarised in Table S3 for the ULC analysis and Table S4 for the LLC analysis. The number of ASVs affected in relation to the total number of ASVs detected is summarised in Table 4. The number of ASVs impacted by family are presented in Table S5 for the ULC analysis and Table S6 for LLC analysis.

**Table 4:**
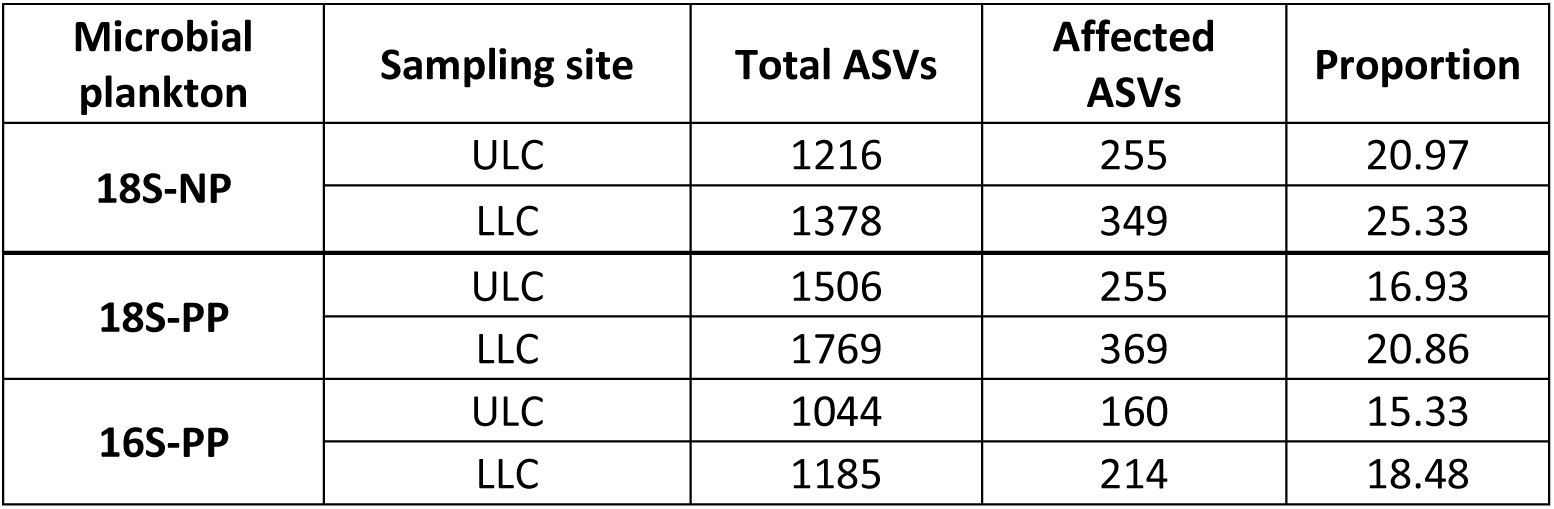
Number of affected ASVs per community and sampling site and their proportion in relation to the total ASVs number in the community.

### ULC vertical mixing microbial plankton community

Winter vertical mixing in the Upper Lake Community (ULC) had a significant effect on the three plankton communities (18S-NP, 18S-PP and 16S-PP). Between 15.33% and 20.97% of the ASVs in these communities were positively affected by this event (Table 4). LogFC, representing the ratio of the difference in abundance between the two conditions, could reach 9.62 (Table S3). The ASVs associated with these LogFC values were at extremely low abundances, below the detection level, during the period of water thermal stratification, but increased sharply during vertical mixing of the water. Aggregating the relative abundance of all ASVs affected positively and comparing the averages between mixed and stratified water revealed that the relative abundance increased during mixing events by a factor ranging from 3.33 (18S-NP) to 8.92 (16S-PP) (Fig. 4).

**Figure 4:**
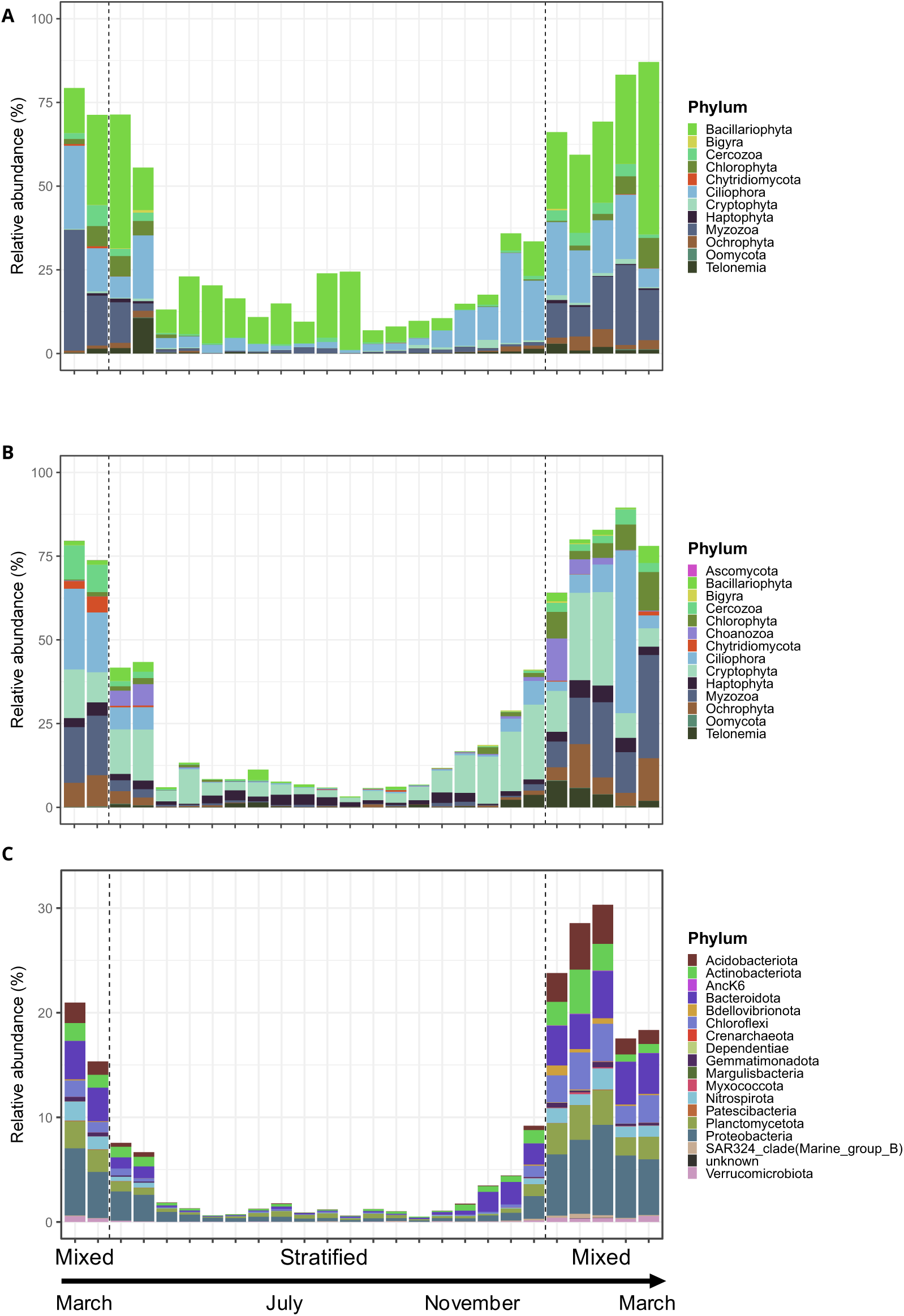
Relative abundances of ASVs, detected by LinDA as having a higher abundance during the vertical mixing in ULC. The stacked bar plots illustrate the relative abundance, in percentages, of ASVs, coloured by phylum, relative to relative abundance of the total community. X axis represents the different sampling date (Table S1) (**A**) eukaryotic nanoplankton (18S-NP), (**B**) eukaryotic picoplankton (18S-PP) and (**C**) prokaryotic picoplankton (16S-PP).

In the 18S-NP community, 100% of the ASVs in the Amphisiellidae (Ciliophora) and Noelaerhabdaceae (Haptophyta) families were affected. The Chytridiaceae (Chytridiomycota) family had more than 75% of its ASVs impacted, as did the Holophryidae (Ciliophora) family, the Peronosporaceae (Oomycota) family, and the Stephanodiscaceae (Bacillariophyta) family (Table S4). Apart from Stephanodiscaceae, which had 35 affected ASVs, all the other families with high levels of impact were represented by four or five ASVs each. Gymnodiniaceae (Myzozoa) had only 54.5% of their ASVs affected, and these all belonged to the Lepidodinium genus. Higher logFC was observed for ASVs affiliated with Strombidiidae (Ciliophora), Gymnodiniaceae, Chlamydomonadaceae (Chlorophyta) and Paraphysomonadaceae (Ochrophyta) (Table S3). When the relative abundance of positively affected ASVs was aggregated at family level, Stephanodiscaceae represented up to 44.92% of the total community on a single sampling date and Gymnodiniaceae had a relative abundance ranging from 0.03% during stratification to a maximum of 33.9% during vertical mixing (Fig. S7).

In the 18S-PP community, 100% of impacted ASVs were found in Amphisiellidae and Ancistridae (Ciliophora), Chlorellales (incertae sedis) and Mychonastaceae (both Chlorophyta), and Peronosporaceae. Stephanodiscaceae, Geminigeraceae (Cryptophyta) and Noelaerhabdaceae also had over 75% of their ASVs affected. Apart from Stephanodiscaceae and Geminigeraceae, which had 18 and 21 affected ASVs respectively, the other highly impacted families were represented by fewer than four ASVs (Table S4). Gymnodiniaceae was affected in only 51.9% of ASVs, but the majority of these were affiliated with Lepidodinium. The highest LogFC values were observed for ASVs affiliated with Gymnodiniaceae, Paraphysomonadaceae, Strombidiidae and Colepidae (Ciliophora) (Table S3). Aggregated at the family level, Colepidae represented up to 47.45% of the total community in early March 2019. The other dominant families were Geminigeraceae, Gymnodiniaceae, Paraphysomonadaceae, Dolichomastigaceae (Chlorophyta) and Protaspidae (Cercozoa) (Fig. S8).

The 16S RNA dataset contained 14 families with at least 75% of ASVs affected by vertical mixing, including 10 families with 100% of ASVs affected. These families were Anck6 (Anck6), Nitrosopumilaceae (Crenarchaeota), Bacteriap25 (Myxococcota), SAR11 clade I, EF100-94H03 and KI89A_clade (all Proteobacteria), KD4-96 and P2-11E (both Chloroflexi), Gaiellaceae (Actinobacteriota), and BD2-11 terrestrial group (Gemmatimonadota). All of these families were represented by a single ASV. The families with 75% of ASVs affected were Vicinamibacteraceae and Bryobacteraceae (both Acidobacteriota), Reyranellaceae (Proteobacteria), and JG30-KF-CM66 (Chloroflexi). These families were represented by 15, 3, 3, and 6 affected ASVs, respectively (Table S4). Although the ASVs with the highest logFC values were taxonomically diverse (Table S3), Anck6 and Crenarchaeota were almost exclusively detected during vertical mixing (Fig. S9). The vertical mixing had a positive effect on the following dominant families: Anaerolineaceae (Chloroflexi), Vicinamibacteraceae, Nitrospiraceae (Nitrospirota), BSV26 and Chitinophagaceae (both Bacteroidota), as well as Gemmataceae and Phycisphaeraceae (both Planctomycetota) (Fig. S9).

### LLC oxygen depleted bottom water layer microbial plankton community

Similarly, bottom water oxygen depletion in the LLC had a significant impact on the three plankton communities (18S-NP, 18S-PP and 16S-PP). Between 18.48% and 25.33% of ASVs in these communities were positively affected by this event (Table 4), with LogFC values similar to those observed in the analysis of water-vertical-mixing communities (Table S3). Aggregating the relative abundance of all ASVs positively affected, and comparing the average between oxic and oxygen-depleted periods revealed an increase in relative abundance during the oxygen depletion event by a factor ranging from 3.57 (18S-NP) to 6.00 (18S-PP) (Fig. 5).

**Figure 5:**
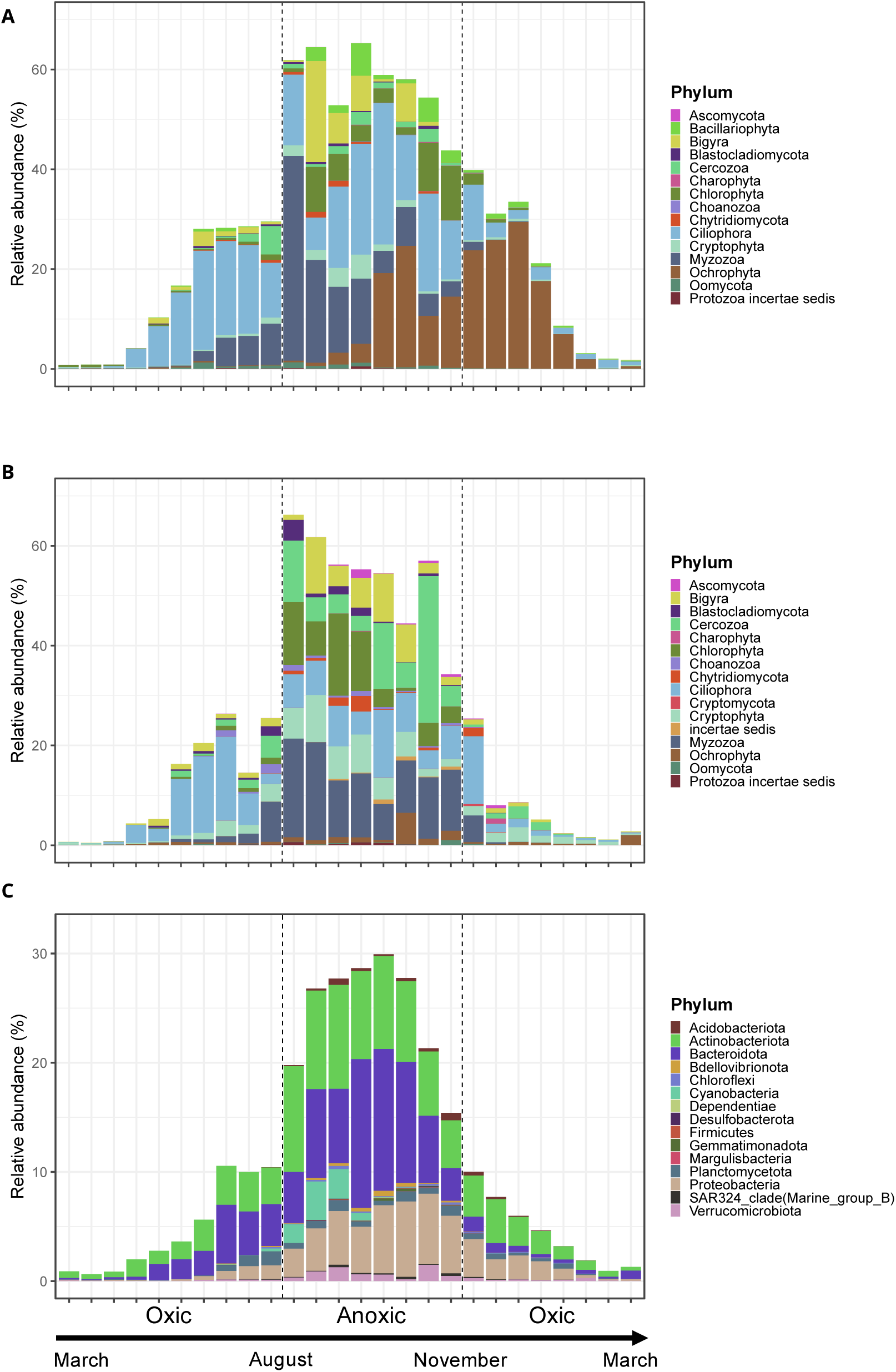
Relative abundances of ASVs, detected by LinDA as having a higher abundance during the oxygen depletion event in LLC. The stacked bar plots illustrate the relative abundance, in percentages, of ASV, coloured by phylum, relative to relative abundance of the total community. X axis represents the different sampling date (Table S1) (**A**) eukaryotic nanoplankton (18S-NP), (**B**) eukaryotic picoplankton (18S-PP) and (**C**) prokaryotic picoplankton (16S-PP).

A total of 19 families had 100% of their ASVs affected by the oxygen depletion event and a further three had 70% or more affected. Most of these families were represented by fewer than four ASVs and belonged to the phyla Ciliophora, Myzozoa, Chlorophyta, Charophyta, Choanozoa and Cercozoa (ordered by the number of families affected; see Table S4). Notable exceptions were Holostichidae (Ciliophora), with 12 affected ASVs, and Peridiniaceae and Suessiaceae (both Myzozoa), with 22 and 31 affected ASVs, respectively (Table S4). All of the ASVs with the highest logFC were taxonomically affiliated with families belonging to Myzozoa and Ciliophora (Table S3). When relative abundances were aggregated at family level, Suessiaceae showed the greatest increase, representing 31.7% of the total community during the initial phase of oxygen depletion. ASVs affiliated with Ciliophora, which were detected as being statistically more abundant in oxygen-depleted water, also peaked when the oxygen concentration started to decrease in the bottom layer (Fig. 5A and Fig. S10).

The 18S-PP dataset comprised 18 families, over 70% of which were affected by oxygen-depleted bottom water layer events. Of these families, 13 had 100% of their ASVs affected. Eleven families were represented by fewer than four ASVs and were affiliated with Myzozoa, Ciliophora, Chlorophyta, Cercozoa and Ochrophyta (Table S4). The families with the highest representation, where all ASVs were affected, were Radiococcaceae (Chlorophyta), Uebelmesseromycetaceae (Chytridiomycota), and Chaetomellaceae (Ascomycota), with 16, 9, and 5 ASVs respectively. The ASVs with the highest logFC were mainly affiliated with the Myzozoa, Cercozoa, Bigyra and Ciliophora phyla (Table S3). The dominating families in terms of relative abundance were Protozoa familia incertae sedis, Radiococcaceae (Chlorophyta), Cryptomonadaceae, Prorocentraceae and Colepidae (Fig. S11). Temporal dynamics can also be observed between families within a phylum. ASVs that were more abundant during the oxygen depletion event and that were affiliated with Myzozoa were affiliated with Suessiaceae and Podolampaceae during the early period of oxygen depletion, followed by Prorocentraceae during the middle period and Gymnodiniaceae during the final period of the oxygen depletion event (Figure S11).

The family Coxiellaceae (Proteobacteria) was the only one in which 100% of ASVs were affected by the oxygen depletion event, with 2 ASVs affected. The second most impacted family was BSV26, with 66.7% (2 out of 3) of its ASVs affected. Rhodocyclaceae (Proteobacteria) was represented by 30 ASVs, of which 53.3% were impacted by the oxygen depletion event (Table S4). Their genus affiliation showed that almost all Sulfitalea and Ferribacterium ASVs were affected. The same dynamic was observed in Lentimicrobiaceae (Bacteroidota), with ASVs only affiliated to an unidentified genus. The taxonomic affiliation of the ASVs with the highest logFC showed a dominance of ASVs affiliated with Proteobacteria (Rhodocyclaceae and an unclassified Gammaproteobacteria), as well as Bacteroidota (Chitinophagales and Flavobacteriales) (Table S3). In terms of relative abundance, the community was dominated by ASVs affiliated with the Actinobacteria cluster I family (acI), the Bacteroidota cluster I family (bacI), the Cyanobacteria family Cyanobiaceae, and the Rhodocyclaceae family. Only three out of the 36 acI ASVs were positively affected by this event, but they were highly dynamic in terms of relative abundance, ranging from 0.28% during the full oxic period to a maximum of 8.82% during the oxygen depletion event. Several ASVs were only detectable during the oxygen depletion period and were affiliated with Cyanobiaceae, Rhodocyclaceae, Methylomonadaceae (Proteobacteria), Anaerolineaceae and Gemmatimonadaceae (Gemmatimonadota) (Fig. S12).

## Discussion

Lake Constance has been extensively studied with respect to seasonal plankton succession and has served as a model system for the development of plankton ecology models, including the Plankton Ecology Group (PEG) framework (Sommer et al. 2012). However, to date, no DNA-sample repository and sequencing-based study exists of the seasonal succession of the nano- and picoplankton in Lake Constance. To address this gap, we initiated a routine biweekly sampling campaign from March 2018 and established a DNA repository of the nano- and picoplankton epilimnetic communities. This sampling campaign has been continued from spring 2021 onwards (i.e., after the Corona pandemic) until today, while for the present study, only one seasonal cycle, from March 2018 until March 2019, was analysed. Climate change is expected to intensify thermal stratification in Lake Constance, leading to increasingly incomplete winter mixing and, thus, to more persistent decreases of oxygen concentration in the deep water, as observed during the period 2013–2016 and 2022 to 2023 (Limnologischer Zustand des Bodensees), with profound consequences for lake ecosystem functioning. In this context, the DNA-repository will provide a critical reference for detecting future changes in microbial plankton community composition. It also represents a valuable resource for monitoring toxic bloom-forming cyanobacteria, which pose a growing threat in pre-Alpine lakes (Jacquet et al. 2005; Fournier et al. 2021).

In the present study, we focused on a first observation of the taxa detectable in ULC and LLC nano- and picoplankton using a general 16S- and 18S-rDNA based approach over one year. The use of generalist 16S and 18S rDNA amplicons delivers a good description of microbial community structure and composition. The amplicon size and choice of hypervariable region still limit robust taxonomic affiliation, especially at the deepest taxonomic levels. To ensure robust analysis, it was decided to work primarily with family as the deepest taxonomic rank. The use of primers pair targeting a specific group of organisms, or a sequence the whole community metagenome, would give a finer taxonomic resolution and access to the community function potential.

PermANOVA and ANOSIM, testing differences in community biodiversity between ULC and LLC, showed contradicting results for the 18S-NP and 18S-PP communities. Heterogeneity of variance can lead to confounding location and dispersion effects in PermANOVA and ANOSIM, which is enhanced by unbalanced design (Anderson 2017). Both heterogeneity of variance and unbalanced design, as a result of removal of replicates of low quality, were present in this study. While corrections are existing (Anderson et al. 2017), none were implemented in the R function adonis2() function at the time of the study. Thus, the results of the biodiversity comparison between the ULC and LLC 18S-NP and 18S-PP communities must be considered carefully due to this confounding effect, and no definitive conclusions could be drawn regarding differences in biodiversity between these communities.

### Microbial plankton diversity

Across all three microbial plankton communities studied, we observed highly similar cyclical seasonal dynamics between the Upper and Lower Lake Constance, despite marked differences in physico-chemical conditions. Eukaryotic nano- and picoplankton diversity peaked during the warmer, stratified period, which presents better light conditions and high nutrient availability, supporting a larger phytoplankton population, followed by an increased predatory and parasitic protist activity (Sommer 1985; Salmaso et al. 2020). The bacterioplankton diversity was highest during cold periods, especially during the water vertical mixing. The origin of this increase of diversity in the water vertical mixing is debated, with either a physical import of the hypolimnion bacterioplankton to the surface water or the change in environmental conditions promoting the growth of already present low abundant taxa. Our results were more supportive of the latter, as we observed an increase in evenness rather than richness during this period, meaning that already detected low abundant taxa have increased in abundance at this period. These patterns are consistent with observations from other Alpine and oligotrophic lakes, indicating a strong seasonal influence on the structure of the microbial plankton community (García et al. 2015; Salmaso et al. 2018, 2020).

The spatial and temporal structures of microbial communities can be driven by deterministic processes, such as selection by environmental factors and biotic interactions, as well as stochastic processes, such as species drift, speciation, and dispersal (Menéndez-Serra et al. 2023). The relative impact of these processes on community structure depends on the type of microbial community (Larsen et al. 2023), the studied ecosystem (Riddley et al. 2025), the seasonality and strength of the environmental pressures (Sommer et al. 2012; Menéndez-Serra et al. 2023), and the scale of the study (Aguilar and Sommaruga 2020). The large scale of this study made it more susceptible to deterministic processes than stochastic ones. Therefore, the similarity in the temporal dynamics of diversity of the three freshwater plankton communities studied, combined with similar patterns observed in other freshwater lakes, suggests that deterministic processes are the dominant drivers of community dynamics. This deterministic influence was not limited to diversity metrics. Winter water vertical mixing and oxygen depletion acted as environmental filters, selecting specific subsets of taxa that suddenly constituted the dominant group in the community. The potential non-random nature of these responses is explored in detail below and supports the observation that community reorganisation during mixing and oxygen depletion is driven by taxon-specific ecological traits rather than stochastic processes.

### Vertical mixing impact on the microbial plankton community

Winter water vertical mixing corresponds to several sudden changes in epilimnetic physico-chemical conditions, including water turbulence, low irradiance, nutrient regeneration, reoxygenation and low water temperature (Diehl et al. 2002; Limnologischer Zustand des Bodensees). These changes act as environmental filters on the microbial plankton community, and our results show that these filters favour specific groups of taxa with particular ecological traits.

During the warmer stratified period, nutrient concentration in the epilimnion decreases due to microbial plankton activity, while nutrient concentration in the hypolimnion increases (Limnologischer Zustand des Bodensees). Winter mixing is the main driver of nutrient regeneration in the epilimnion, enabling taxa that favour less oligotrophic conditions to grow and outcompete organisms specific to oligotrophic conditions. Previous studies have shown a positive relationship between the Ciliate families Colepidae and Holophryidae, the Haptophyte family Neolaerhabdaceae and the Cercozoan family Protaspidae, and vertical mixing and nutrient regeneration (Müller et al. 1991; Egge et al. 2015; Caracciolo et al. 2022). A previous study has observed a positive relationship between oligotrophic conditions and certain Actinobacteria bacteria in lake mesocosms (Haukka et al. 2006) while another have found that bacteria affiliated with Planctomycetota, Chloroflexi, and Bacteroidota increased in relative abundance during vertical mixing of the water column in the oligotrophic Flathead Lake (Evans et al. 2024). These findings support the observations made in this study.

Physical disturbance further reshapes community structure by reducing sinking rates and altering the sinking rates of cells. Some dinoflagellate species can sense turbulence and adjust their behaviour by forming chains, increasing their swimming speed, or adapting their swimming trajectories in order to protect themselves from turbulence-induced mortality. Lepidodinium, found in our dataset, is closely related to the genus Gymnodinium, known to modify its behaviour from single cells to chains in response to upwelling (Smayda 2010). The reduced sinking rate enables larger cells, which typically sink more quickly during thermal stratification, to remain in the surface water. This lower sinking rate could act synergistically with nutrient regeneration to allow larger species with a low volume-to-surface ratio to uptake nutrients more efficiently. Together, these factors could create an environment favourable for the growth and outcompeting of larger species, such as centric Stephanodiscaceae diatoms, over smaller species (Diehl et al. 2002; Winder and Hunter 2008).

During this period, protists that relied less on light acquisition and were capable of an alternative pathway also seemed to hold an ecological advantage. Several of the taxa detected in this study are known mixotrophs, such as the dinoflagellate Lepidodinium, and Mychonastalean and Chlamydomonadalean algae (Paranjape et al. 2016; Millette et al. 2021; Liu et al. 2021). The lower irradiance present during vertical mixing may trigger the mixotrophic behaviour of some dinoflagellates (Bielewicz et al. 2011; Millette et al. 2017).

Physical disturbance is also responsible for homogenising the microbial community throughout the water column (Mukherjee et al. 2024), as well as resuspending sinking organic matter and sediments. This study showed that vertical mixing events allowed microbial taxa that are usually very low in abundance to grow, rather than large-scale import of microbial taxa from the hypolimnion. Still, some taxa remained below the detection limit outside of these events and are representative of the oxygenated hypolimnion community, such as Nitrosopumilaceae, Anaerolineaceae, Phycisphaeraceae, Gemmataceae and Nitrospiraceae (Newton et al. 2011; Klotz et al. 2022; Evans et al. 2024), or are commonly detected in freshwater and marine sediments, such as the KI89A clade, BD2-11, and Acidobacteria taxa (Newton et al. 2011; Vavourakis et al. 2018; Scicchitano et al. 2022). The hypolimnion protist community, which is mainly composed of parasitic and saprophytic particle-associated organisms, likely thrives in the epilimnion water, presumably due to the homogenisation of environmental conditions in the water column, resulting in a homogenised community along the water column, or due to the resuspension of organic matter (Mukherjee et al. 2024).

Indeed, the oxygenated hypolimnion water contains recalcitrant organic matter that sinks to the bottom of the lake. This matter is produced by the microbial carbon pump or comes from land (Legendre et al. 2015). The resuspension of organic matter in the water column by vertical mixing provides nutrients for microbial taxa capable of degrading the matter, allowing these taxa to grow and increase in relative abundance. Physical disturbance was shown to inhibit Chytridiomycota infection of phytoplankton (McKindles et al. 2021).The relative abundance increase of Chytridiomycota in this study could be the results of a shift towards saprophytic behaviour, whereby feeding on the resuspended particulate organic matter (Gleason et al. 2008). Similarly, several bacterial taxa known to degrade recalcitrant organic matter were detected in our datasets, including Chitinophagaceae, Gemmataceae and Vicinamibacteraceae. (Silva et al. 2020; Rutere et al. 2020; Stuij et al. 2024).

Despite the strong and consistent effects of vertical mixing on microbial plankton communities, many of the bacterial lineages that are positively affected remain poorly characterised, with few cultivated representatives. Nevertheless, vertical mixing has a tremendous effect on microbial plankton communities. Surviving eukaryotic cells will serve as a seed bank for the next spring bloom, with experiments showing that weak vertical mixing favours cyanobacterial blooms over diatom blooms (Smayda 2010; Visser et al. 2016). This shift in community composition will likely propagate to phytoplankton grazers, parasites, and microorganisms that rely on cross-feeding, cascading to the rest of the trophic network. Improved characterisation of the microbial taxa that respond to vertical mixing is therefore essential to understanding how mixing-driven disturbances influence the stability of freshwater ecosystems.

### Impact of the water oxygen depletion on the microbial plankton community

Oxygen depletion is commonly observed when the consumption of oxygen by microorganisms exceeds its supply. Seasonally, it can be observed when an increase in nutrient concentration occurs through the sedimentation of detritus following increased phytoplankton productivity and thermal stratification, which prevents reoxygenation. As most biogeochemical cycles are strongly influenced by the presence of molecular oxygen, its depletion fundamentally alters redox conditions, leading to the accumulation of reduced compounds such as ammonia, phosphorus, hydrogen sulfide, methane and dissolved iron and manganese (Kosolapov et al. 2003; Biderre-Petit et al. 2011; Limnologischer Zustand des Bodensees). While there were no measurements of hydrogen sulfide or methane available for Lake Constance during the 2018–2019 season, nitrate, manganese, and ferric iron were monitored throughout the year, with accumulation observed in the oxygen-depleted water (Limnologischer Zustand des Bodensees). The absence of oxygen acts as an environmental filter on microbial communities, favouring those adapted to low or no oxygen.

Oxygen depletion tends to select for eukaryotic microorganisms that are capable of mixotrophy or heterotrophy, and that possess alternative metabolic pathways. This gives them an ecological advantage in oxygen-depleted conditions. Many of the taxa that were found to be more abundant during oxygen depletion in this study and in previous research exhibit such nutritional strategies (Atteia et al. 2013; Kamp et al. 2015; Gauns et al. 2020). Dinoflagellates appear particularly well adapted to low-oxygen conditions; mesocosm experiments have shown that they grow under oxygen depletion due to their more efficient utilisation of nutrients released from the degradation of organic matter and elution from sediments (Hinode et al. 2019). Fermentation and anaerobic respiration have also been documented in various eukaryotic taxa, including Chlorophyta, Ciliates, Fungi (mostly Chytridiomycota) and some nanoflagellates (Fenchel 2012; Atteia et al. 2013). A broad variety of eukaryotic microorganisms possess capacities affiliated with nitrogen cycles, such as nitrate reduction, denitrification, and dissimilatory nitrate reduction to ammonium (DNRA), which enable survival under oxygen depletion. Some Ciliates are capable of nitrate reduction, while others, isolated from anoxic freshwater basins, possess endosymbionts that can perform denitrification (Kamp et al. 2015; Zeller et al. 2026). Oxygen depletion has also been shown to trigger “fungal denitrification” and “ammonia fermentation” in certain fungal taxa, enabling survival in oxygen-depleted conditions (Kamp et al. 2015).

Ciliates may benefit further from oxygen depletion through morphological and trophic flexibility. Intracellular chloroplast sequestration has been observed in mixotrophic Ciliates, enabling them to continue to be supplied with oxygen by phototrophic endosymbionts (Esteban et al. 2010). Ciliates can also gain an advantage from their morphological and size diversity. Thanks to this diversity, Ciliates not only prey on bacteria, but also graze on phytoplankton (Lischke et al. 2016). This may explain why we observed their relative abundance increase when oxygen started to deplete, as this combines with a large phytoplankton biomass.

Oxygen depletion creates a redox transition in the water column, defining microbial ecological niches by controlling the availability of terminal electron acceptors for respiration. Consequently, specific bacterial and archaeal groups that rely on nitrate, nitrite, sulphate, manganese, ferric iron or carbon dioxide instead of oxygen are selected, and the presence of these groups is governed by the availability of these compounds. The increased abundance of the obligate anaerobic, Fe(III)-reducing bacterium Ferribacterium is consistent with the accumulation of ferrous iron that was measured during oxygen depletion (Cummings et al. 1999; Limnologischer Zustand des Bodensees). Lentimicrobium, a genus affiliated with oxygen depletion in our study and belonging to the Lentimicrobiaceae family, is strictly anaerobic and capable of DNRA, in line with the accumulation of nitrate in the absence of oxygen (Sun et al. 2016). Although information on the concentration of hydrogen sulfide and other reduced sulfur species was unavailable, the detection of the anoxygenic sulfur-oxidising genus Sulfuritalea, as well as ASVs affiliated with green and purple sulfur bacteria, indicates the euxinic nature of the oxygen-depleted water in Lake Constance (Sperfeld et al. 2019).

Several bacterial families that were detected during oxygen depletion are also known to degrade complex organic matter anaerobically, such as the Anaerolineaceae family, which can metabolise complex polysaccharides through fermentation and degradation (McIlroy et al. 2017). Chitinophagaceae and Flavobacteriales can anaerobically degrade chitin (Beier and Bertilsson 2011; Rosenberg 2014) and the detection of acI (Actinobacteria cluster I) bacteria could be attributed to their ability to utilise the products of chitin hydrolysis(Beier and Bertilsson 2011).

We also detected bacterial taxa that are typically described as microaerophilic and that live at the oxic–anoxic transition zone. The presence of the aerobic methane-oxidiser Methylomonadaceae indicates the production of methane in oxygen-depleted water, which is then transported upwards to the oxic water where it is used by Methylomonadaceae (Cabrol et al. 2020). Similarly, Sideroxydans is a facultative Fe(II) oxidiser that thrives in microaerophilic niches at the oxic–anoxic transition(Emerson et al. 2013).

The water column in Lower Lake Constance was not completely devoid of oxygen; only the lower part of the column was affected. This resulted in samples integrating oxic, microoxic and anoxic environments. This likely reduced the detectability of strictly anaerobic and microaerophilic taxa living at the oxic–anoxic transition. A more refined analysis would require discrete samples taken at different depths to elucidate the microbial communities of the oxygen-depleted water of Lower Lake Constance with greater resolution.

## Conclusions

Despite pronounced physico-chemical differences between the two basins, microbial plankton communities exhibited highly similar cyclical temporal dynamics across all three planktonic subsets. This indicates that, at the scale investigated, deterministic seasonal forcing imposes a dominant and recurrent structure on microbial plankton diversity, largely independent of local environmental conditions. In contrast, episodic disturbances—winter vertical mixing in Upper Lake Constance and deep-water oxygen depletion in Lower Lake Constance—induced rapid and pronounced compositional shifts. These events acted as environmental filters, favouring taxa with traits such as metabolic flexibility or tolerance to physical disturbance and low-oxygen conditions. Importantly, our results suggest that these shifts are driven mainly by changes in the relative abundance of low-abundance taxa already present in the community. The strong and taxon-specific responses to these disturbances highlight microbial plankton as sensitive indicators of environmental change and as active contributors to ecosystem functioning. Many responsive bacterial lineages remain poorly characterised, emphasizing the need for improved functional and physiological knowledge of freshwater microbial diversity. The establishment of a long-term DNA repository for Lake Constance provides a critical baseline for detecting future community shifts. Increasing water temperatures are expected to strengthen thermal stratification, weaken winter mixing, and promote more frequent and persistent oxygen depletion events. Such changes are likely to alter nutrient regeneration, oxygen availability, and redox structure in the water column, with cascading effects on microbial community organisation, trophic interactions, and lake ecosystem stability. By linking microbial dynamics to environmental drivers, this study advances our capacity to predict how freshwater ecosystems may respond to future climatic forcing. Future integration of this long-term framework with depth-resolved and functional approaches will be essential to resolve microbial metabolic responses and species interactions in greater detail.

## Supporting information

Supplementary data total

## Author Contributions

Conceptualization, D.S, C.F; methodology, D.S, C.F; Investigation, C.F; formal analysis, C.F; data curation, C.F; writing—original draft preparation, C.F; writing—review and editing, D.S, C.F; project administration, D.S; funding acquisition, D.S. All authors have read and agreed to the published version of the manuscript.

## Data Availability Statement

The Illumina sequence data presented in this study is accessible at NCBI under the bioproject number PRJNA1478098.

## Acknowledgments

We would like to highlight the excellent support received from Alfred Sulger, Angelika Seifried, Pia Mahler and Josef Halder at the Limnological Institute of University of Konstanz for managing the ship cruises. Further, we would like thank all members of the AG Schleheck, the RTG-R3, and especially Tina Romer and Frank Peeters, for their support.

