## Supplementary data total for "Seasonal succession of nano- and picoplankton communities in Lake Constance: conserved dynamics despite compositional shifts under contrasting mixing and oxygen regimes"

[1] Microbial Ecology and Limnic Microbiology. Limnological Institute. Department of Biology. University of Konstanz. D-78457 Konstanz. Germany

*Shared corresponding authorship:

.

**Table S1: Date of sampling campaign by sampling sites.**

| Sampling site | Date | Season | Condition | Sampling site | Date | Season | Condition |
|---|---|---|---|---|---|---|---|
| ULC | 13.03.2018 | Winter | Mixed | LLC | 15.03.18 | Winter | Oxic |
| ULC | 27.03.2018 | Spring | Mixed | LLC | 29.03.18 | Spring | Oxic |
| ULC | 10.04.2018 | Spring | Stratified | LLC | 14.04.18 | Spring | Oxic |
| ULC | 27.04.2018 | Spring | Stratified | LLC | 26.04.18 | Spring | Oxic |
| ULC | 08.05.2018 | Spring | Stratified | LLC | 09.05.18 | Spring | Oxic |
| ULC | 22.05.2018 | Spring | Stratified | LLC | 24.05.18 | Spring | Oxic |
| ULC | 05.06.2018 | Spring | Stratified | LLC | 07.06.18 | Spring | Oxic |
| ULC | 19.06.2018 | Summer | Stratified | LLC | 21.06.18 | Summer | Oxic |
| ULC | 03.07.2018 | Summer | Stratified | LLC | 05.07.18 | Summer | Oxic |
| ULC | 17.07.2018 | Summer | Stratified | LLC | 19.07.18 | Summer | Oxic |
| ULC | 31.07.2018 | Summer | Stratified | LLC | 02.08.18 | Summer | Oxic |
| ULC | 14.08.2018 | Summer | Stratified | LLC | 16.08.18 | Summer | Anoxic |
| ULC | 28.08.2018 | Summer | Stratified | LLC | 30.08.18 | Summer | Anoxic |
| ULC | 11.09.2018 | Summer | Stratified | LLC | 13.09.18 | Summer | Anoxic |
| ULC | 21.09.2018 | Autumn | Stratified | LLC | 01.10.18 | Autumn | Anoxic |
| ULC | 09.10.2018 | Autumn | Stratified | LLC | 11.10.18 | Autumn | Anoxic |
| ULC | 23.10.2018 | Autumn | Stratified | LLC | 25.10.18 | Autumn | Anoxic |
| ULC | 06.11.2018 | Autumn | Stratified | LLC | 08.11.2018 | Autumn | Anoxic |
| ULC | 22.11.2018 | Autumn | Stratified | LLC | 20.11.2018 | Autumn | Oxic |
| ULC | 04.12.2018 | Autumn | Stratified | LLC | 06.12.2018 | Autumn | Oxic |
| ULC | 18.12.2018 | Autumn | Stratified | LLC | 20.12.2018 | Autumn | Oxic |
| ULC | 11.01.2019 | Winter | Mixed | LLC | 10.01.2019 | Winter | Oxic |
| ULC | 29.01.2019 | Winter | Mixed | LLC | 31.01.2019 | Winter | Oxic |
| ULC | 12.02.2019 | Winter | Mixed | LLC | 14.02.2019 | Winter | Oxic |
| ULC | 26.02.2019 | Winter | Mixed | LLC | 27.02.2019 | Winter | Oxic |
| ULC | 12.03.2019 | Winter | Mixed | LLC | 19.03.2019 | Winter | Oxic |

**Table S2: Overview of sample number per microbial plankton and sampling sites, at individual steps of the bioinformatics processing with the final number of reads in each dataset and the number of detected ASVs per microplankton type and sampling site.** Raw files paired end correspond to the sequencing files send by the sequencing company. Files merge reads are after merging forward and reverse reads together. Files concatenate correspond to the two individual Illumina sequencing run merge together. Files after filtration correspond to the number of files having a minimum of 10000 reads after filtering low abundant ASV and represent the number of samples used for the analysis. The total number of reads column indicate as well the number of reads in each dataset after the bioinformatic process and filtration of low abundant ASVs. The number of ASVs represent the one that passed the filtration of low abundant ASV (see Material and Methods section for filtration parameters). Abbreviations used: 18S-NP, eukaryotic nanoplankton dataset; 18S-PP, eukaryotic picoplankton dataset; 16S-PP, prokaryotic picoplankton dataset; ULC, Upper Lake Constance (*Wallhausen* sampling site); LLC, Lower Lake Constance (*Zeller See* sampling site); ASV, amplicon sequencing variant.

| Microbial plankton | Sampling site | Raw files paired end | Files merge reads | Files per site | Files Concatenate | Files After filtration | Total number of reads | ASVs number by microbial plankton | ASVs number by site |
|---|---|---|---|---|---|---|---|---|---|
| **18S-NP** | ULC | 624 | 312 | 156 | 78 | 73 | 2718202 | 1708 | 1216 |
| | LLC | | | 156 | 78 | 70 | 2382616 | | 1378 |
| **18S-PP** | ULC | 624 | 312 | 156 | 78 | 76 | 3553391 | 2195 | 1506 |
| | LLC | | | 156 | 78 | 72 | 3868407 | | 1769 |
| **16S-PP** | ULC | 624 | 312 | 156 | 78 | 78 | 4080533 | 1513 | 1044 |
| | LLC | | | 156 | 78 | 75 | 4944881 | | 1185 |

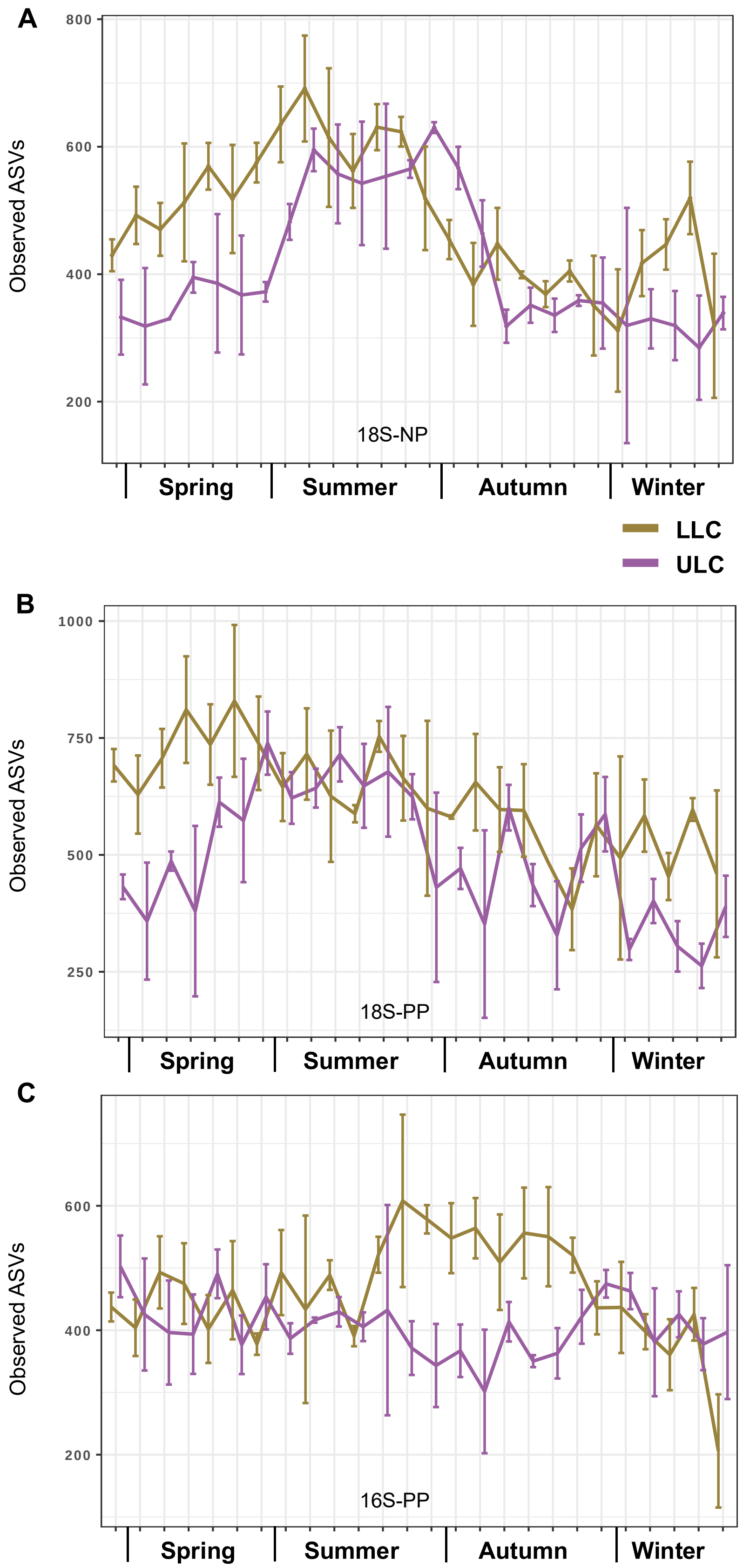

**Figure S1: Observed ASVs richness calculated per sampling site and across the sampling dates.** The purple lines correspond to the Upper Lake Constance (ULC) and the gold lines correspond to the Lower Lake Constance (LLC) sampling site. The upper panel (A) shows the indices for the eukaryotic nanoplankton (18S-NP dataset), the central panel (B) for the eukaryotic picoplankton (18S-PP dataset) and the lower panel (C) for the prokaryotic picoplankton (16S-PP dataset). The x-axis represents the different sampling date (Table S1) labelled by seasons. Note that the y-axes have different scale for each of the three graphs.

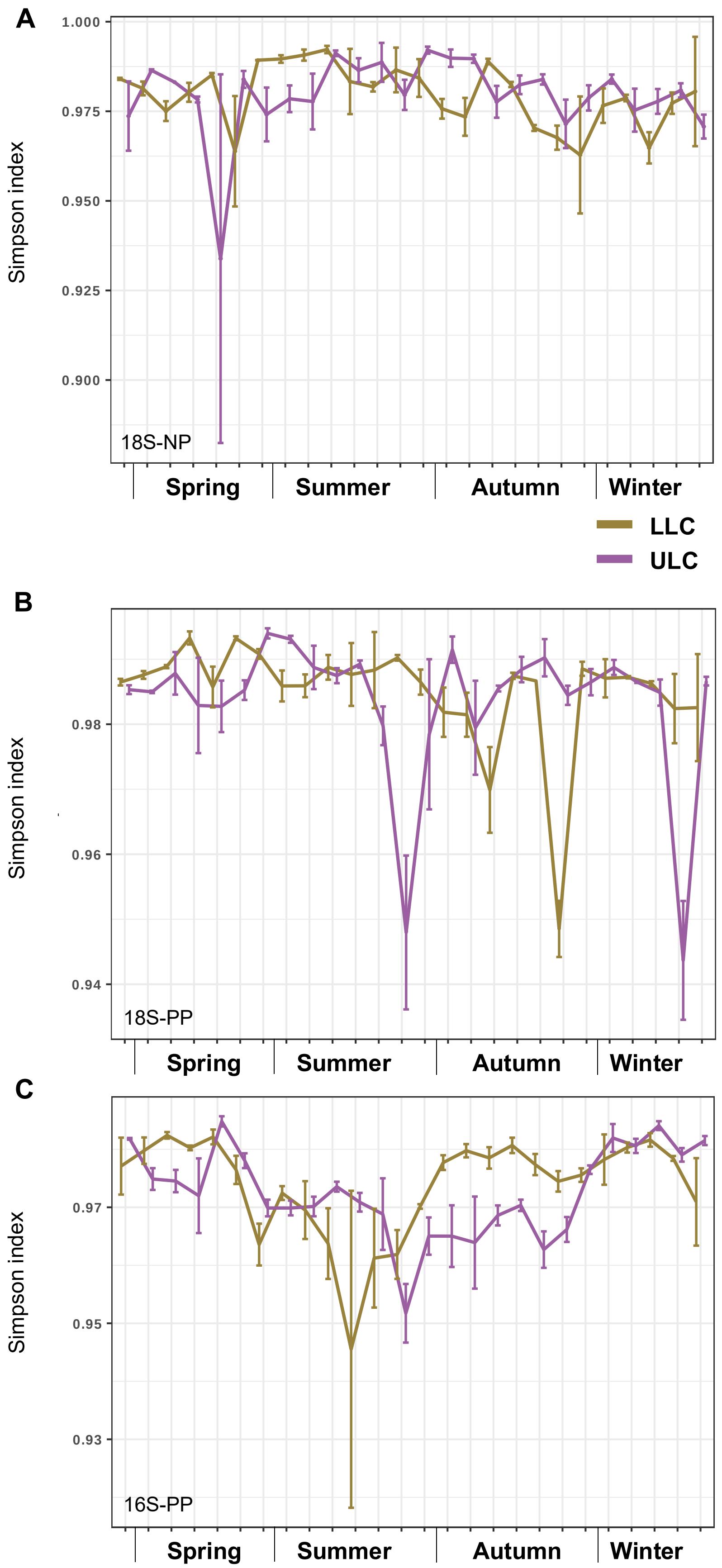

**Figure S2: Simpson diversity index diversity calculated per sampling site and across the sampling dates.** The purple lines correspond to the Upper Lake Constance (ULC) and the gold lines correspond to the Lower Lake Constance (LLC) sampling site. The upper panel (A) shows the indices for the eukaryotic nanoplankton (18S-NP dataset), the central panel (B) for the eukaryotic picoplankton (18S-PP dataset) and the lower panel (C) for the prokaryotic picoplankton (16S-PP dataset). The x-axis represents the different sampling date (Table S1) labelled by seasons. Note that the y-axes have different scale for each of the three graphs.

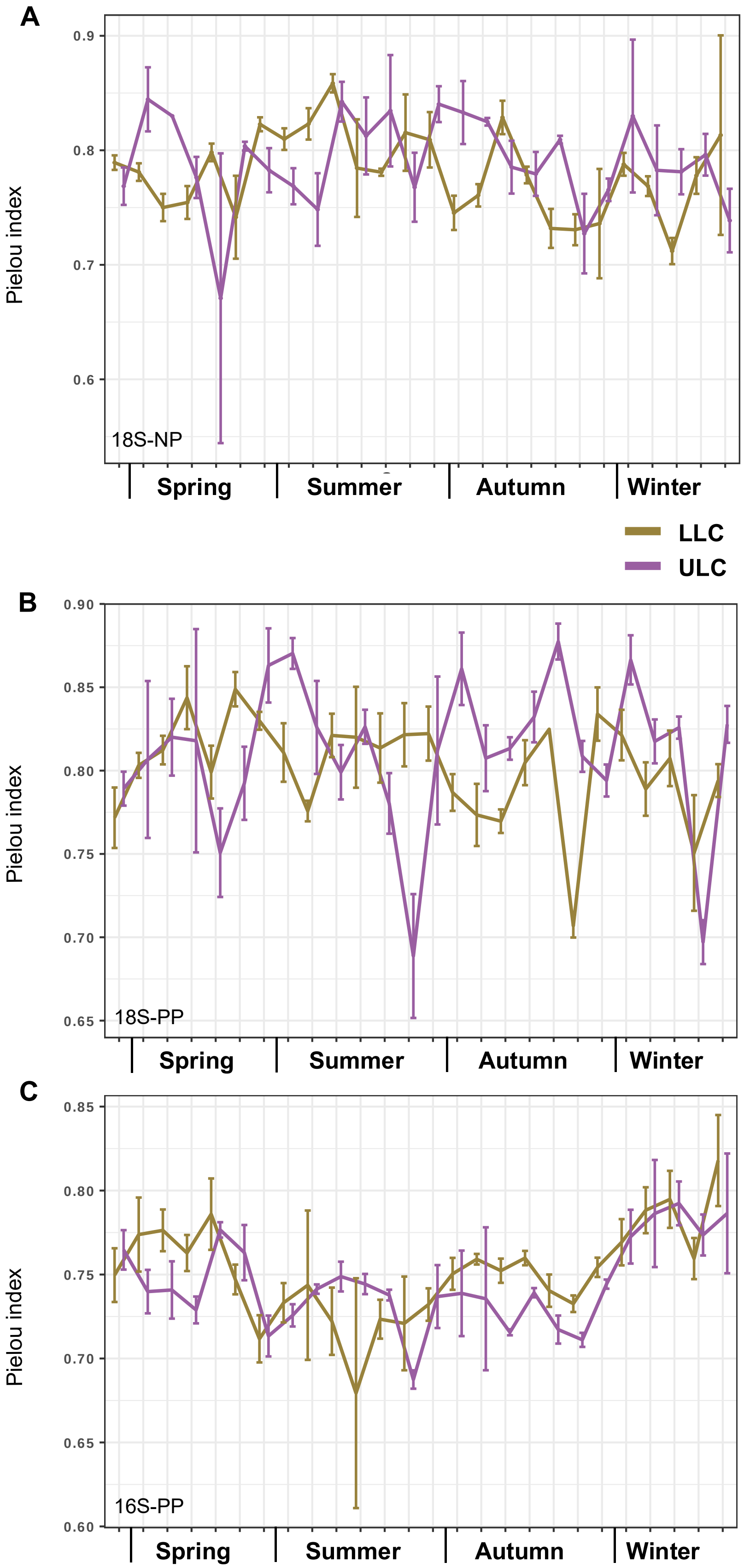

**Figure S3: Pielou evenness index diversity calculated per sampling site and across the sampling dates.** The purple lines correspond to the Upper Lake Constance (ULC) and the gold lines correspond to the Lower Lake Constance (LLC) sampling site. The upper panel (A) shows the indices for the eukaryotic nanoplankton (18S-NP dataset), the central panel (B) for the eukaryotic picoplankton (18S-PP dataset) and the lower panel (C) for the prokaryotic picoplankton (16S-PP dataset). The x-axis represents the different sampling date (Table S1) labelled by seasons. Note that the y-axes have different scale for each of the three graphs.

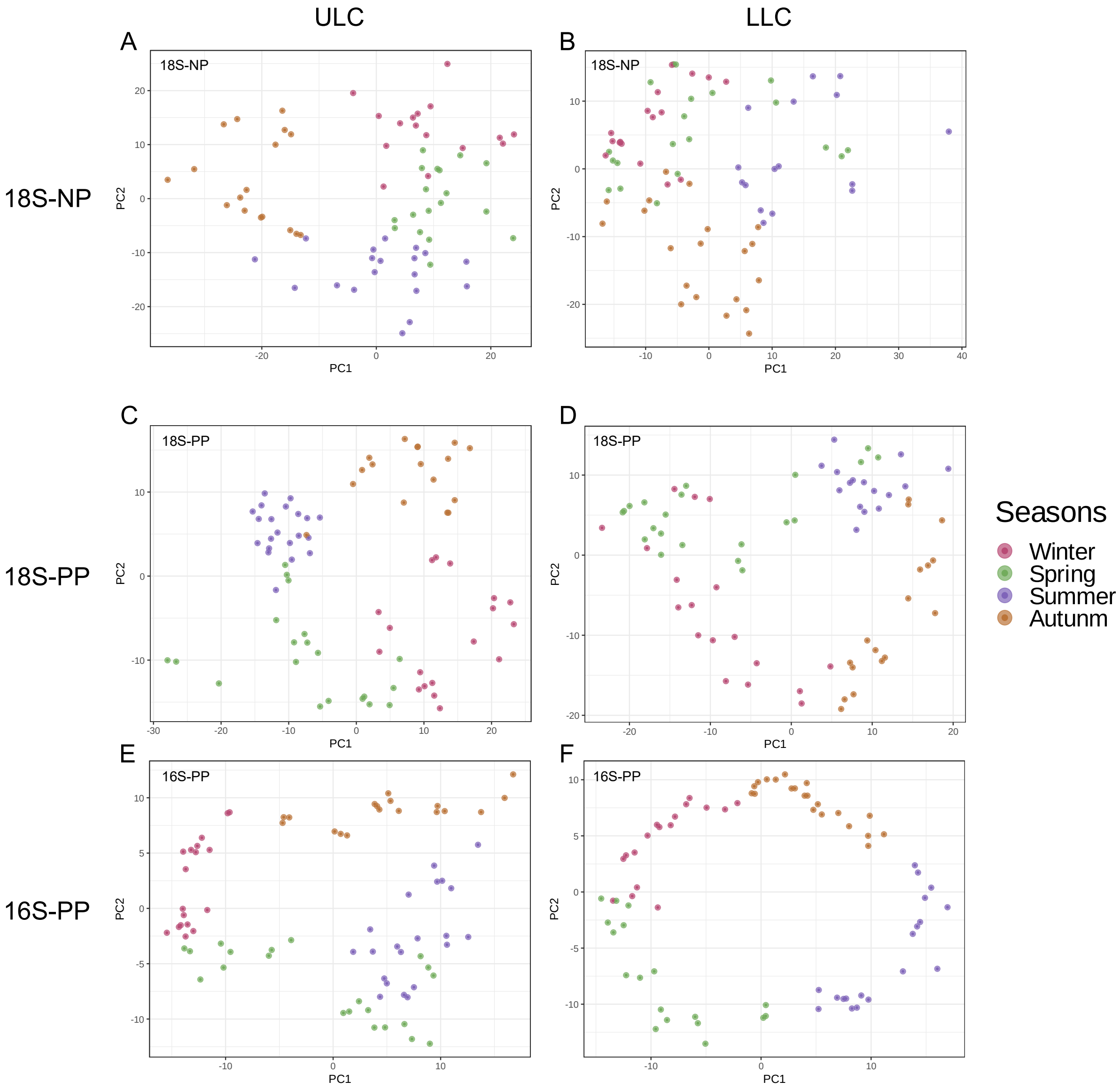

**Figure S4: Temporal dynamic of the microplankton community observed in ULC and LLC independently.** Dissimilarity matrix calculated based on the phylogenetic isometric log-ratio (PhilR) transformed Euclidean distance matrix and visualize by Principal Component Analysis (PCA). Colors indicate the seasons. The top panels (AB) show the data for the eukaryotic nanoplankton (18S-NP). The centrals panel (CD) show the data for the eukaryotic picoplankton (18S-PP). The lower panels (EF) show the data for the prokaryotic picoplankton (16S-PP). The right panels (A - C - E) show the ULC temporal biodiversity and the left panels (B - D - F) show the LLC temporal biodiversity.

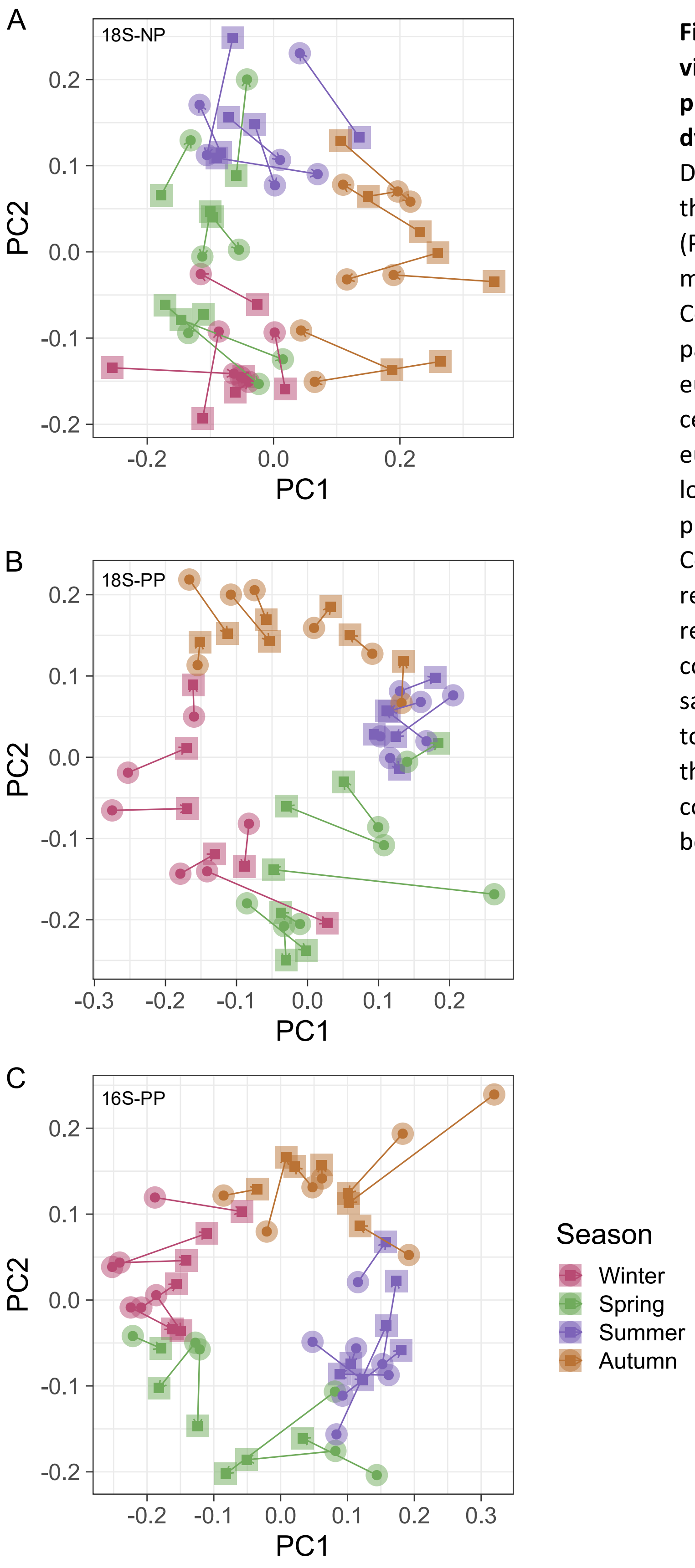

**Figure S5: Procrustes analysis visualization of the nano- and picoplankton temporal biodiversity dynamic between ULC and LLC.** Dissimilarity matrix calculated based on the phylogenetic isometric log-ratio (PhilR) transformed Euclidean distance matrix and visualize by Principal Component Analysis (PCA). The top panel (A) shows the data for the eukaryotic nanoplankton (18S-NP). The central panel (B) shows the data for the eukaryotic picoplankton (18S-PP). The lower panel (C) shows the data for the prokaryotic picoplankton (16S-PP). Colors indicate the seasons. Square represent ULC samples and circle represent LLC samples. Segment connecting points indicates which samples from ULC dataset correspond to the samples in the LLC dataset. Each three plankton showed high concordance of temporal dynamic between ULC and LLC

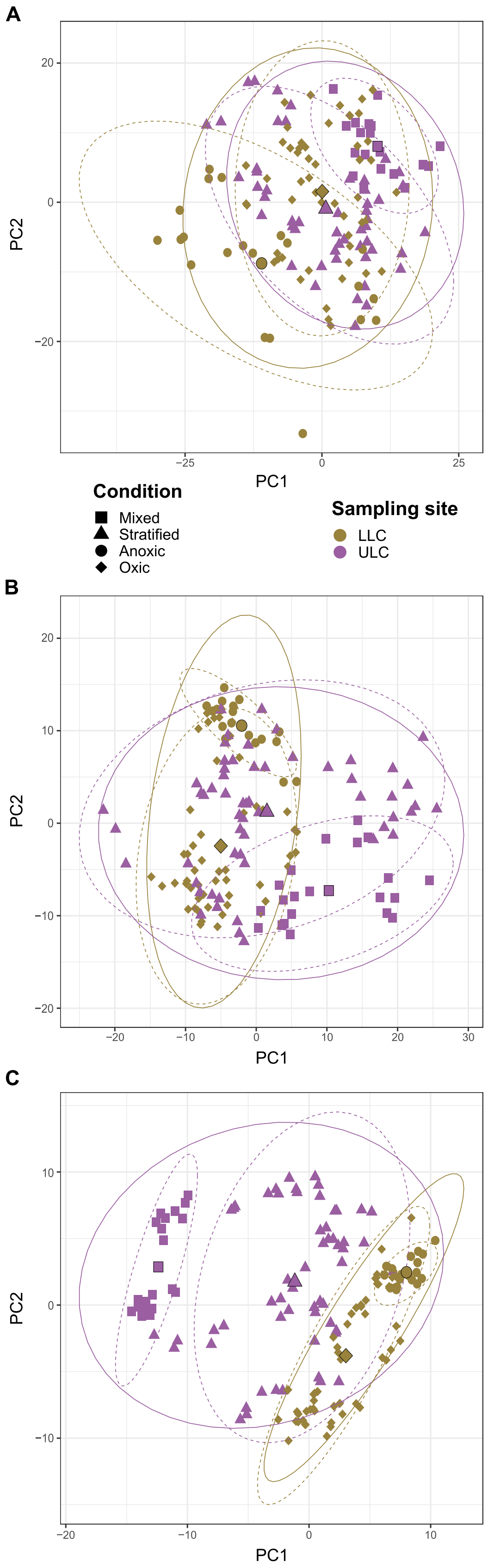

**Figure S6: Principal Component Analysis of nano- and picoplankton biodiversity across the seasonal sampling of the different parts of the lake.** Dissimilarity matrices calculated based on the phylogenetic isometric log-ratio (PhilR)-transformed Euclidean distance matrix. Colors indicate the sampling site; gold, LLC; purple, ULC. Shape indicates the different conditions observed in the water. Square represent ULC water vertical mixing samples and triangle the water stratified samples. Round represent LLC bottom water anoxic event samples and diamond full oxic column samples. Centroids were calculated per condition and are represented by larger points coloured by sampling site and shaped by conditions. Ellipses were calculated following a multivariate t-distribution with a confidence level of 0.95. Two sets of ellipses were drawn. Dotted ellipses represented the conditions and full ellipses represented the sampling sites The top panel (A) shows the data for the eukaryotic nanoplankton (18S-NP), the central panel (B) shows the data for the eukaryotic picoplankton (18S-PP) and the lower panel (C) shows the data for the prokaryotic picoplankton (16S-PP) For all three communities, samples representing the ULC vertical mixing (square points) and LLC anoxic event (round points) were the most isolated one, mostly responsible for the statistical significance difference of biodiversity between ULC and LLC calculated by PermANOVA and ANOSIM (table 3 and 4).

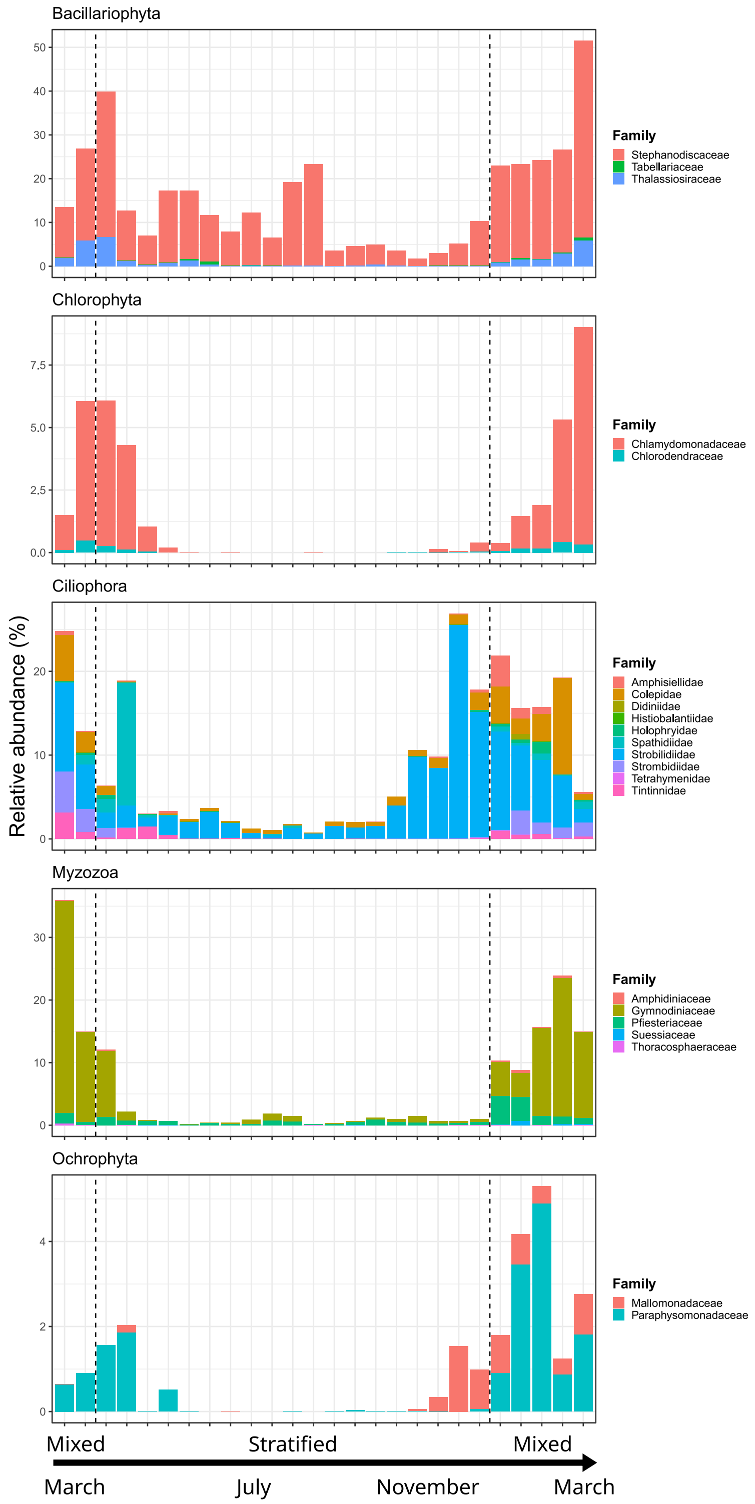

**Figure S7: Relative abundances of the main taxa having a higher abundance during the vertical mixing in ULC in the 18S-NP dataset found by LinDA.** The stacked bar plots illustrate the relative abundance, in percentages, of ASVs. ASVs are grouped and coloured by family and separated by phyla. X axis represents the different sampling date (table S1) and y axis the relative abundance. Beware the difference in x axis.

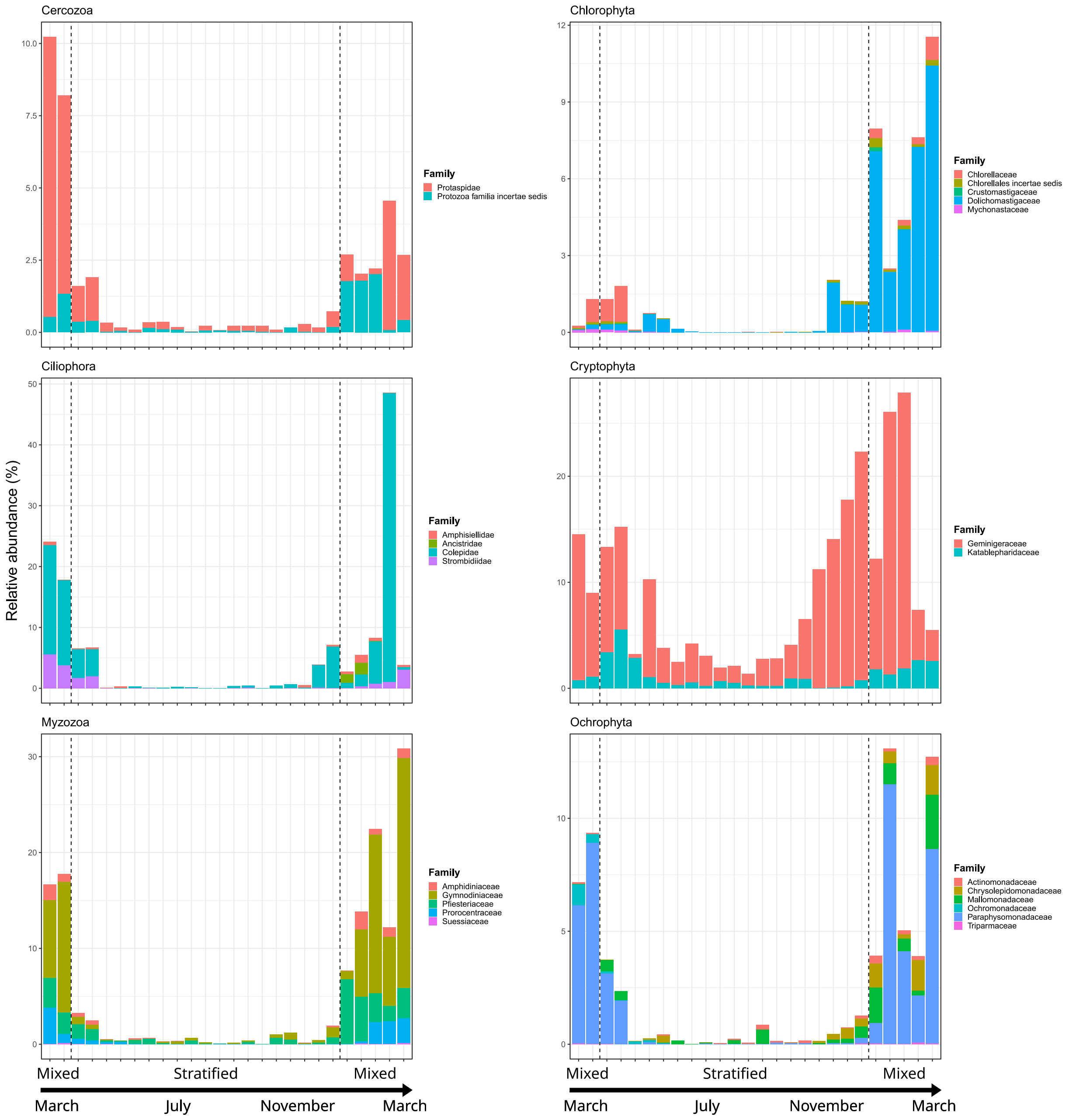

**Figure S8: Relative abundances of the main taxa having a higher abundance during the vertical mixing in ULC in the 18S-PP dataset found by LinDA.** The stacked bar plots illustrate the relative abundance, in percentages, of ASVs. ASVs are grouped and coloured by family and separated by phyla. X axis represents the different sampling date (table S1) and y axis the relative abundance. Beware the difference in x axis.

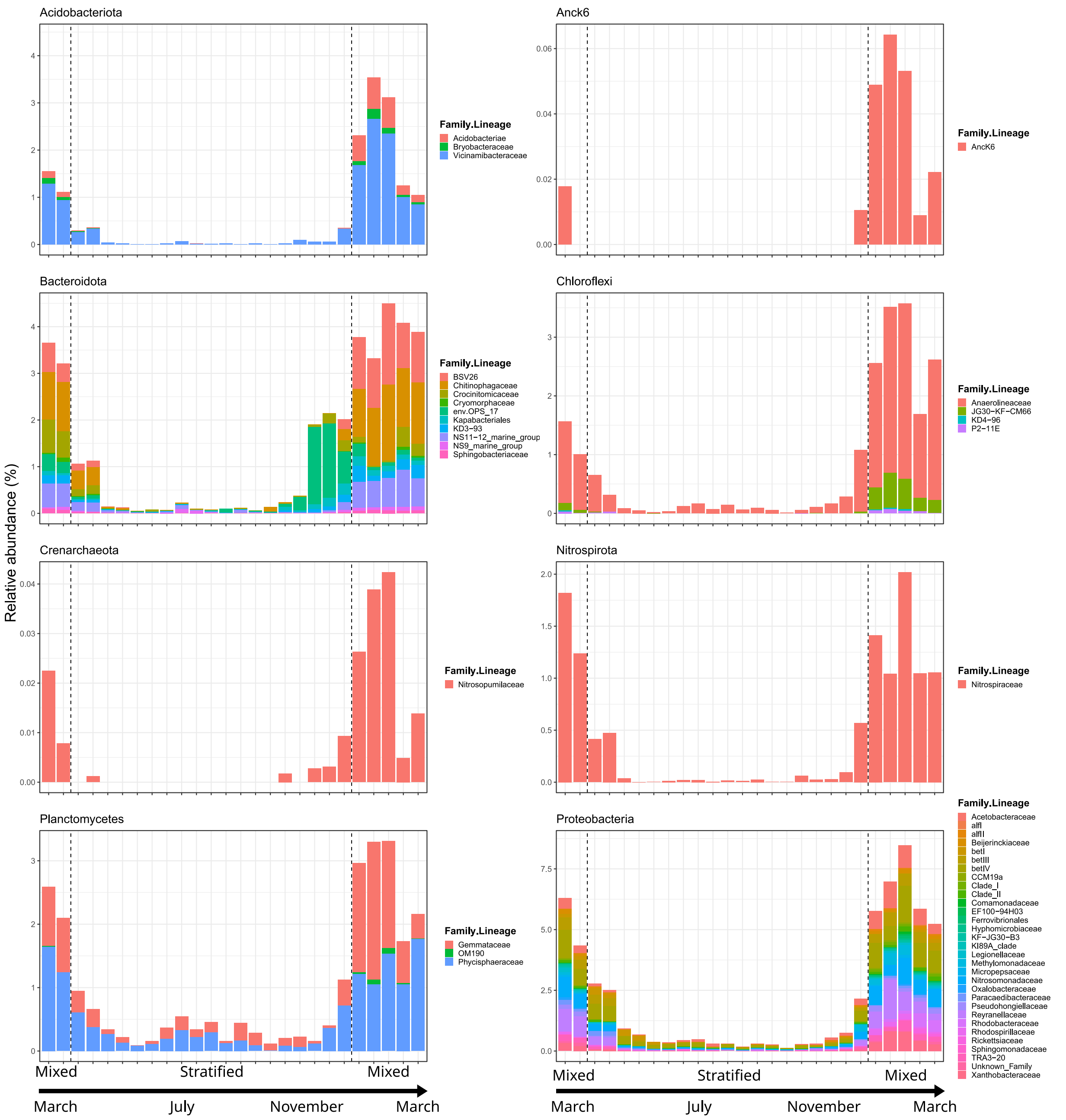

**Figure S9: Relative abundances of the main taxa having a higher abundance during the vertical mixing in ULC in the 16S-PP dataset found by LinDA.** The stacked bar plots illustrate the relative abundance, in percentages, of ASVs. ASVs are grouped and coloured by family and separated by phyla. X axis represents the different sampling date (table S1) and y axis the relative abundance. Beware the difference in x axis.

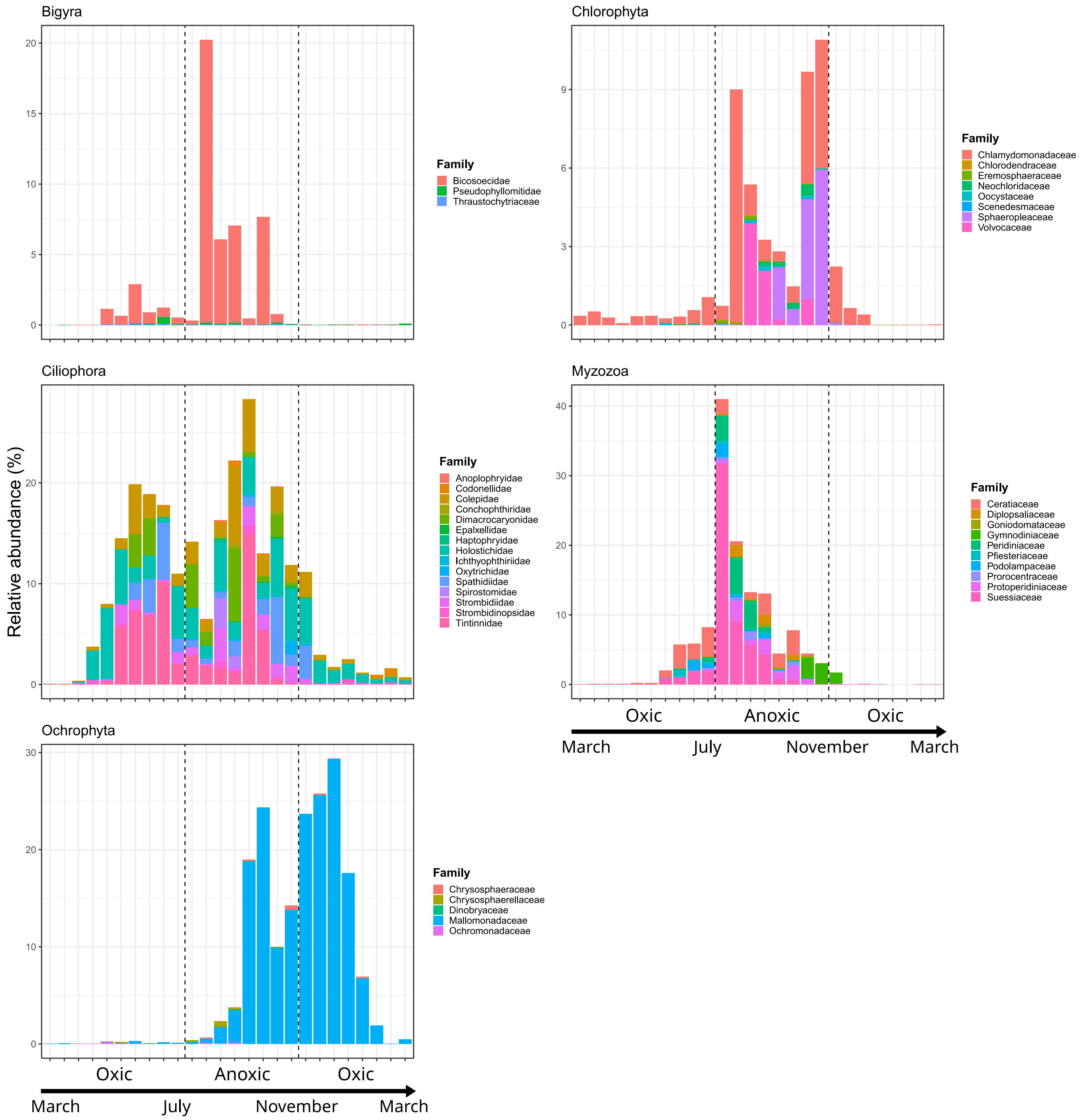

**Figure S10: Relative abundances of the main taxa having a higher abundance during the anoxic deep-water in LLC in the 18S-NP dataset found by LinDA.** The stacked bar plots illustrate the relative abundance, in percentages, of ASV. ASVs are grouped and coloured by family and separated by phyla. X axis represents the different sampling date (table S1) and y axis the relative abundance. Beware the difference in x axis.

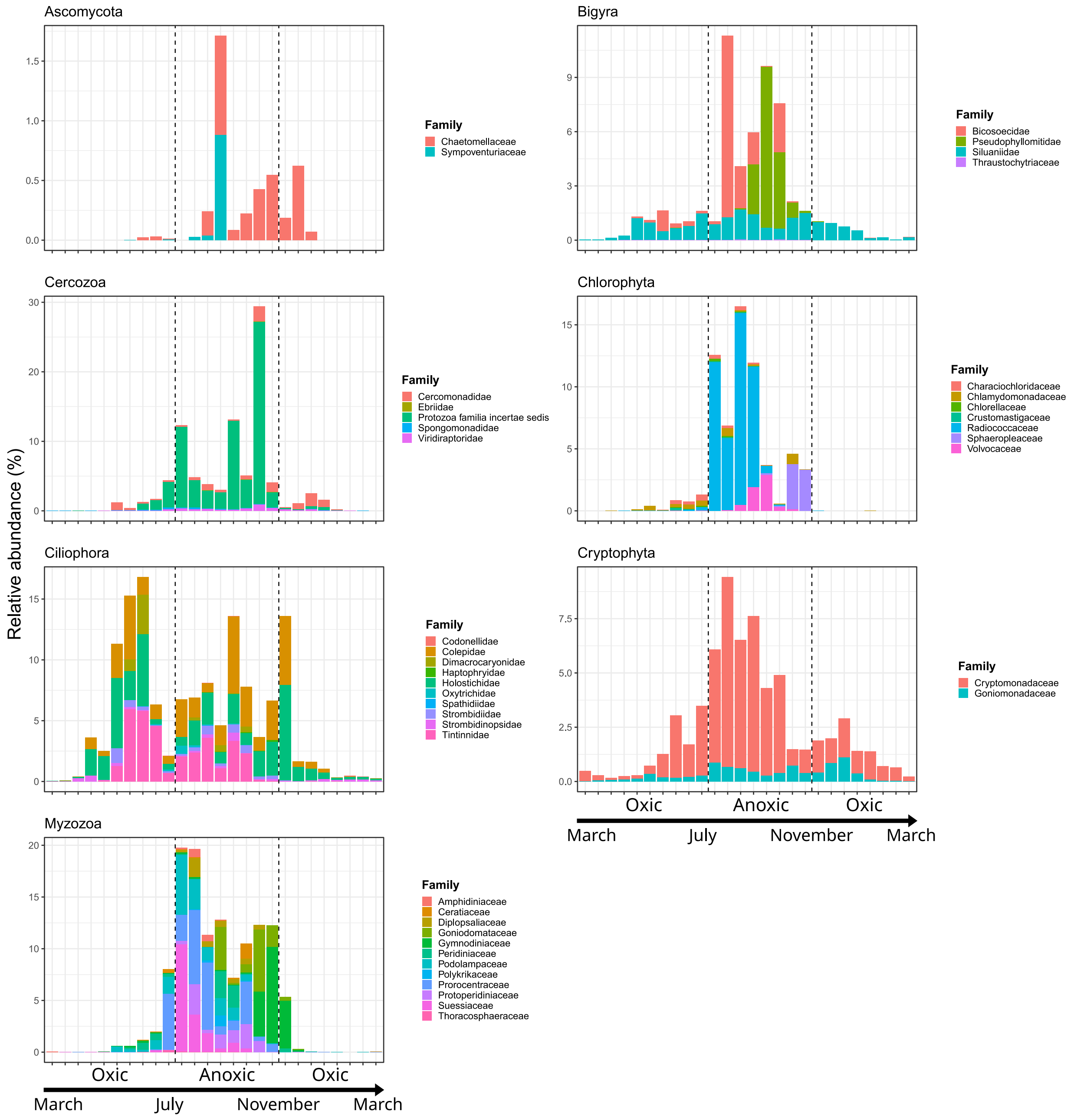

**Figure S11: Relative abundances of the main taxa having a higher abundance during the anoxic deep-LLC in the 18S-PP dataset found by LinDA.** The stacked bar plots illustrate the relative abundance, in percentages, of ASV. ASVs are grouped and coloured by family and separated by phyla. X axis represents the different sampling date (table S1) and y axis the relative abundance. Beware the difference in x axis.

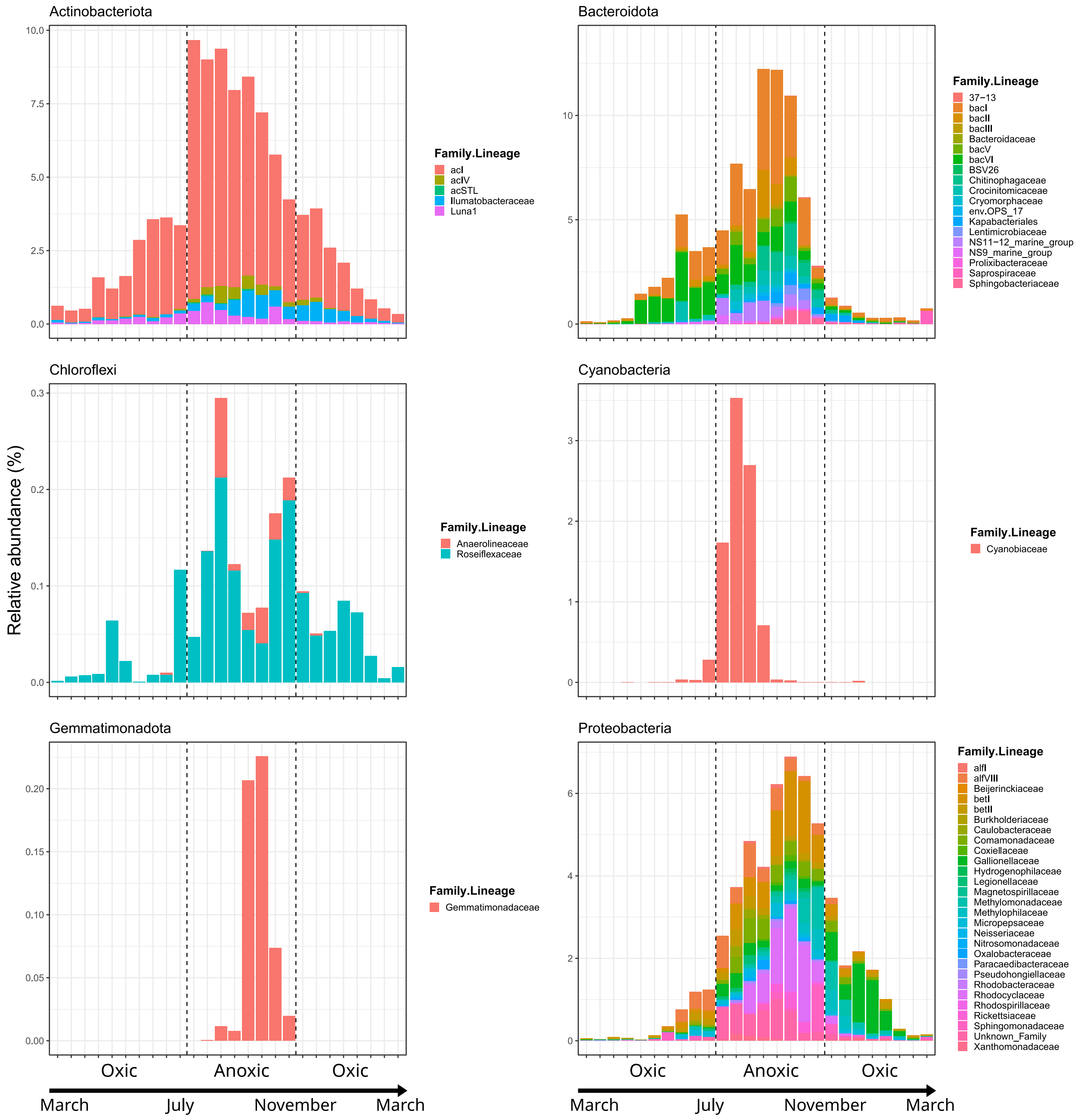

**Figure S12: Relative abundances of the main taxa having a higher abundance during the anoxic deep-LLC in the 16S-PP dataset found by LinDA.** The stacked bar plots illustrate the relative abundance, in percentages, of ASV. ASVs are grouped and coloured by family and separated by phyla. X axis represents the different sampling date (table S1) and y axis the relative abundance. Beware the difference in x axis.

**Table S3: Linear Models for Differential Abundance (LinDA) outputs for the composition of microbial plankton composition differences between the stratified and mixed water column in ULC.** The outputs consist of ASVs that pass the threshold of Log2FC above 2 and an adjusted p-value below 0.01, along with their taxonomic affiliation up to family level. The ASVs are categorised as follows: eukaryotic nanoplankton (18S-NP), eukaryotic picoplankton (18S-PP) and prokaryotic picoplankton (16S-PP).

| Microbial plankton | ASV | log2FC | adjusted p-value | Kingdoms | Phylum | Class | Order | Family |
|---|---|---|---|---|---|---|---|---|
| 18S-NP | ASV1962 | 8.25 | 1.90E-21 | Eukaryota | Ciliophora | Spirotrichea | Oligotrichia | Strombidiidae |
| | ASV1915 | 8.09 | 3.33E-21 | Eukaryota | Ciliophora | Spirotrichea | Oligotrichia | Strombidiidae |
| | ASV499 | 7.80 | 3.52E-13 | Eukaryota | Myzozoa | Dinophyceae | Gymnodiniales | Gymnodiniaceae |
| | ASV1674 | 7.67 | 3.87E-18 | Eukaryota | Ciliophora | Spirotrichea | Oligotrichia | Strombidiidae |
| | ASV797 | 7.66 | 7.45E-21 | Eukaryota | Ciliophora | Spirotrichea | Oligotrichia | Strombidiidae |
| | ASV1897 | 7.60 | 2.36E-11 | Eukaryota | Chlorophyta | Chlorophyceae | Chlamydomonadales | Chlamydomonadaceae |
| | ASV1827 | 7.60 | 1.34E-13 | Eukaryota | Myzozoa | Dinophyceae | Gymnodiniales | Gymnodiniaceae |
| | ASV2480 | 7.49 | 4.46E-12 | Eukaryota | Chlorophyta | Chlorophyceae | Chlamydomonadales | Chlamydomonadaceae |
| | ASV1680 | 7.48 | 1.50E-14 | Eukaryota | Myzozoa | Dinophyceae | Gymnodiniales | Gymnodiniaceae |
| | ASV91 | 7.46 | 1.62E-11 | Eukaryota | Chlorophyta | Chlorophyceae | Chlamydomonadales | Chlamydomonadaceae |
| | ASV1860 | 7.44 | 1.72E-13 | Eukaryota | Myzozoa | Dinophyceae | Gymnodiniales | Gymnodiniaceae |
| | ASV2738 | 7.44 | 1.12E-10 | Eukaryota | Chlorophyta | Chlorophyceae | Chlamydomonadales | Chlamydomonadaceae |
| | ASV229 | 7.20 | 5.32E-15 | Eukaryota | Myzozoa | Dinophyceae | Gymnodiniales | Gymnodiniaceae |
| | ASV313 | 7.16 | 2.16E-15 | Eukaryota | Myzozoa | Dinophyceae | Gymnodiniales | Gymnodiniaceae |
| | ASV2169 | 7.11 | 2.05E-16 | Eukaryota | Myzozoa | Dinophyceae | Gymnodiniales | Gymnodiniaceae |
| | ASV1102 | 7.01 | 3.05E-15 | Eukaryota | Myzozoa | Dinophyceae | Gymnodiniales | Gymnodiniaceae |
| | ASV2298 | 7.00 | 2.11E-17 | Eukaryota | Cercozoa | Thecofilosea | Cryomonadida | Protaspidae |
| | ASV1819 | 6.94 | 5.05E-13 | Eukaryota | Ochrophyta | Chrysophyceae | Chromulinales | Paraphysomonadaceae |
| | ASV1320 | 6.90 | 4.63E-13 | Eukaryota | Ochrophyta | Chrysophyceae | Chromulinales | Paraphysomonadaceae |
| | ASV2759 | 6.87 | 5.51E-17 | Eukaryota | Myzozoa | Dinophyceae | Gymnodiniales | Gymnodiniaceae |
| | ASV2100 | 6.85 | 1.23E-15 | Eukaryota | Myzozoa | Dinophyceae | Gymnodiniales | Gymnodiniaceae |
| | ASV2182 | 6.72 | 5.99E-15 | Eukaryota | Myzozoa | Dinophyceae | Gymnodiniales | Gymnodiniaceae |
| | ASV207 | 6.34 | 2.68E-15 | Eukaryota | Ochrophyta | Chrysophyceae | Chromulinales | Paraphysomonadaceae |

| | | | | | | | | |
|---|---|---|---|---|---|---|---|---|
| | ASV268 | 6.27 | 7.65E-12 | Eukaryota | Bacillariophyta | Mediophyceae | Thalassiosirales | Thalassiosiraceae |
| | ASV2464 | 6.23 | 2.52E-12 | Eukaryota | Cercozoa | Thecofilosea | Cryomonadida | Protaspidae |
| | ASV1543 | 6.10 | 1.46E-13 | Eukaryota | Myzozoa | Dinophyceae | Gymnodiniales | Gymnodiniaceae |
| | ASV2740 | 6.08 | 2.53E-10 | Eukaryota | Ciliophora | Prostomatea | Prorodontida | Colepidae |
| | ASV2643 | 6.02 | 4.19E-15 | Eukaryota | Ochrophyta | Chrysophyceae | Chromulinales | Paraphysomonadaceae |
| | ASV378 | 5.95 | 1.29E-13 | Eukaryota | Ochrophyta | Chrysophyceae | Chromulinales | Paraphysomonadaceae |
| | ASV2207 | 5.94 | 1.69E-12 | Eukaryota | Ochrophyta | Chrysophyceae | Chromulinales | Paraphysomonadaceae |
| | ASV2792 | 5.84 | 4.75E-10 | Eukaryota | Ciliophora | Prostomatea | Prorodontida | Colepidae |
| | ASV687 | 5.60 | 1.22E-09 | Eukaryota | Ciliophora | Prostomatea | Prorodontida | Colepidae |
| | ASV2009 | 5.57 | 2.74E-11 | Eukaryota | Bacillariophyta | Mediophyceae | Thalassiosirales | Thalassiosiraceae |
| | ASV1131 | 5.52 | 3.23E-10 | Eukaryota | Ciliophora | Prostomatea | Prorodontida | Colepidae |
| | ASV2423 | 5.50 | 2.64E-14 | Eukaryota | Chlorophyta | Chlorodendrophyceae | Chlorodendrales | Chlorodendraceae |
| | ASV1257 | 5.50 | 3.33E-11 | Eukaryota | Bacillariophyta | Mediophyceae | Stephanodiscales | Stephanodiscaceae |
| | ASV989 | 5.40 | 9.78E-11 | Eukaryota | Bacillariophyta | Mediophyceae | Thalassiosirales | Thalassiosiraceae |
| | ASV242 | 5.34 | 5.99E-15 | Eukaryota | Chlorophyta | Chlorodendrophyceae | Chlorodendrales | Chlorodendraceae |
| | ASV483 | 5.33 | 2.02E-09 | Eukaryota | Ciliophora | Spirotrichea | Stichotrichida | Amphisiellidae |
| | ASV1928 | 5.32 | 2.98E-12 | Eukaryota | Bacillariophyta | Mediophyceae | Thalassiosirales | Thalassiosiraceae |
| | ASV2864 | 5.28 | 2.10E-08 | Eukaryota | Ciliophora | Spirotrichea | Tintinnida | Tintinnidae |
| | ASV1729 | 5.24 | 2.02E-15 | Eukaryota | Ochrophyta | Chrysophyceae | Chromulinales | Paraphysomonadaceae |
| | ASV2264 | 5.12 | 6.29E-15 | Eukaryota | Ochrophyta | Chrysophyceae | Chromulinales | Paraphysomonadaceae |
| | ASV1871 | 5.08 | 2.29E-16 | Eukaryota | Myzozoa | Dinophyceae | Gymnodiniales | Gymnodiniaceae |
| | ASV815 | 5.08 | 1.17E-15 | Eukaryota | Myzozoa | Dinophyceae | Gymnodiniales | Gymnodiniaceae |
| | ASV833 | 5.07 | 7.48E-15 | Eukaryota | Chlorophyta | Chlorodendrophyceae | Chlorodendrales | Chlorodendraceae |
| | ASV2062 | 5.06 | 6.02E-16 | Eukaryota | Myzozoa | Dinophyceae | Gymnodiniales | Gymnodiniaceae |
| | ASV1523 | 5.06 | 1.19E-10 | Eukaryota | Bacillariophyta | Mediophyceae | Thalassiosirales | Thalassiosiraceae |
| | ASV2437 | 5.01 | 2.68E-15 | Eukaryota | Myzozoa | Dinophyceae | Amphidiniales | Amphidiniaceae |
| | ASV1796 | 4.97 | 2.39E-13 | Eukaryota | Cercozoa | Thecofilosea | Cryomonadida | Protaspidae |
| | ASV1820 | 4.94 | 4.14E-07 | Eukaryota | Ciliophora | Spirotrichea | Tintinnida | Tintinnidae |
| | ASV620 | 4.92 | 5.27E-09 | Eukaryota | Ciliophora | Prostomatea | Prorodontida | Colepidae |
| | ASV1544 | 4.91 | 6.24E-08 | Eukaryota | Ciliophora | Spirotrichea | Stichotrichida | Amphisiellidae |

| | | | | | | | | |
|---|---|---|---|---|---|---|---|---|
| | ASV853 | 4.90 | 3.35E-09 | Eukaryota | Ciliophora | Spirotrichea | Stichotrichida | Amphisiellidae |
| | ASV724 | 4.85 | 3.39E-08 | Eukaryota | Ciliophora | Prostomatea | Prorodontida | Colepidae |
| | ASV2382 | 4.85 | 2.75E-08 | Eukaryota | Bacillariophyta | Mediophyceae | Stephanodiscales | Stephanodiscaceae |
| | ASV2060 | 4.84 | 2.54E-08 | Eukaryota | Ciliophora | Spirotrichea | Stichotrichida | Amphisiellidae |
| | ASV2345 | 4.80 | 1.15E-06 | Eukaryota | Bacillariophyta | Mediophyceae | Stephanodiscales | Stephanodiscaceae |
| | ASV1017 | 4.79 | 6.55E-08 | Eukaryota | Ciliophora | Litostomatea | Haptorida | Spathidiidae |
| | ASV1306 | 4.76 | 2.16E-15 | Eukaryota | Myzozoa | Dinophyceae | Gymnodiniales | Gymnodiniaceae |
| | ASV2275 | 4.75 | 5.51E-17 | Eukaryota | Myzozoa | Dinophyceae | Gymnodiniales | Gymnodiniaceae |
| | ASV2299 | 4.73 | 1.33E-10 | Eukaryota | Bacillariophyta | Mediophyceae | Thalassiosirales | Thalassiosiraceae |
| | ASV1436 | 4.72 | 6.29E-15 | Eukaryota | Cercozoa | Thecofilosea | Cryomonadida | Protaspidae |
| | ASV1026 | 4.71 | 1.26E-09 | Eukaryota | Cercozoa | Thecofilosea | Cryomonadida | Protaspidae |
| | ASV1336 | 4.67 | 2.11E-07 | Eukaryota | Bacillariophyta | Mediophyceae | Stephanodiscales | Stephanodiscaceae |
| | ASV1609 | 4.67 | 3.92E-08 | Eukaryota | Ciliophora | Prostomatea | Prorodontida | Colepidae |
| | ASV33 | 4.64 | 6.84E-07 | Eukaryota | Ciliophora | Spirotrichea | Tintinnida | Tintinnidae |
| | ASV1347 | 4.64 | 2.29E-10 | Eukaryota | Ciliophora | Prostomatea | Prorodontida | Holophryidae |
| | ASV1717 | 4.63 | 4.93E-11 | Eukaryota | Ciliophora | Prostomatea | Prorodontida | Holophryidae |
| | ASV63 | 4.62 | 3.68E-06 | Eukaryota | Bacillariophyta | Mediophyceae | Stephanodiscales | Stephanodiscaceae |
| | ASV819 | 4.62 | 1.90E-09 | Eukaryota | Cercozoa | Protozoa classis incertae sedis | Protozoa ordo incertae sedis | Protozoa familia incertae sedis |
| | ASV1340 | 4.62 | 1.22E-14 | Eukaryota | Myzozoa | Dinophyceae | Amphidiniales | Amphidiniaceae |
| | ASV2320 | 4.54 | 4.97E-14 | Eukaryota | Chlorophyta | Chlorodendrophyceae | Chlorodendrales | Chlorodendraceae |
| | ASV2741 | 4.54 | 1.48E-08 | Eukaryota | Haptophyta | Coccolithophyceae | Isochrysidales | Noelaerhabdaceae |
| | ASV2362 | 4.54 | 9.65E-14 | Eukaryota | Myzozoa | Dinophyceae | Gymnodiniales | Gymnodiniaceae |
| | ASV2633 | 4.47 | 5.32E-07 | Eukaryota | Bacillariophyta | Mediophyceae | Stephanodiscales | Stephanodiscaceae |
| | ASV1431 | 4.45 | 1.30E-09 | Eukaryota | Cercozoa | Thecofilosea | Cryomonadida | Protaspidae |
| | ASV1670 | 4.45 | 3.98E-10 | Eukaryota | Ciliophora | Prostomatea | Prorodontida | Holophryidae |
| | ASV416 | 4.45 | 9.52E-07 | Eukaryota | Ciliophora | Spirotrichea | Tintinnida | Tintinnidae |
| | ASV2425 | 4.43 | 8.21E-11 | Eukaryota | Myzozoa | Dinophyceae | Peridiniales | Pfiesteriaceae |
| | ASV684 | 4.37 | 7.38E-13 | Eukaryota | Ochrophyta | Chrysophyceae | Chromulinales | Paraphysomonadaceae |
| | ASV942 | 4.35 | 4.71E-12 | Eukaryota | Myzozoa | Dinophyceae | Peridiniales | Pfiesteriaceae |
| | ASV2035 | 4.34 | 3.93E-08 | Eukaryota | Haptophyta | Coccolithophyceae | Isochrysidales | Noelaerhabdaceae |

| | | | | | | | | |
|---|---|---|---|---|---|---|---|---|
| | ASV2321 | 4.33 | 1.14E-10 | Eukaryota | Ciliophora | Prostomatea | Prorodontida | Holophryidae |
| | ASV2501 | 4.31 | 5.16E-09 | Eukaryota | Haptophyta | Coccolithophyceae | Isochrysidales | Noelaerhabdaceae |
| | ASV310 | 4.30 | 3.03E-14 | Eukaryota | Myzozoa | Dinophyceae | Gymnodiniales | Gymnodiniaceae |
| | ASV353 | 4.29 | 3.49E-07 | Eukaryota | Ciliophora | Litostomatea | Haptorida | Spathidiidae |
| | ASV153 | 4.29 | 2.14E-07 | Eukaryota | Bacillariophyta | Mediophyceae | Stephanodiscales | Stephanodiscaceae |
| | ASV1163 | 4.29 | 6.62E-08 | Eukaryota | Bacillariophyta | Mediophyceae | Stephanodiscales | Stephanodiscaceae |
| | ASV303 | 4.28 | 5.26E-07 | Eukaryota | Bacillariophyta | Mediophyceae | Stephanodiscales | Stephanodiscaceae |
| | ASV2789 | 4.24 | 5.52E-12 | Eukaryota | Myzozoa | Dinophyceae | Amphidiniales | Amphidiniaceae |
| | ASV1328 | 4.21 | 4.46E-12 | Eukaryota | Myzozoa | Dinophyceae | Peridiniales | Pfiesteriaceae |
| | ASV1994 | 4.17 | 7.86E-16 | Eukaryota | Myzozoa | Dinophyceae | Gymnodiniales | Gymnodiniaceae |
| | ASV2199 | 4.14 | 7.05E-07 | Eukaryota | Ciliophora | Litostomatea | Haptorida | Spathidiidae |
| | ASV2466 | 4.13 | 2.86E-12 | Eukaryota | Myzozoa | Dinophyceae | Peridiniales | Pfiesteriaceae |
| | ASV874 | 4.12 | 1.84E-08 | Eukaryota | Bacillariophyta | Mediophyceae | Stephanodiscales | Stephanodiscaceae |
| | ASV849 | 4.11 | 3.79E-07 | Eukaryota | Bacillariophyta | Mediophyceae | Stephanodiscales | Stephanodiscaceae |
| | ASV2259 | 4.06 | 1.48E-07 | Eukaryota | Bacillariophyta | Mediophyceae | Stephanodiscales | Stephanodiscaceae |
| | ASV2284 | 4.04 | 2.17E-10 | Eukaryota | Cercozoa | Thecofilosea | Cryomonadida | Protaspidae |
| | ASV743 | 4.04 | 1.09E-06 | Eukaryota | Ciliophora | Litostomatea | Haptorida | Spathidiidae |
| | ASV547 | 4.04 | 2.17E-10 | Eukaryota | Ochrophyta | Chrysophyceae | Chromulinales | Paraphysomonadaceae |
| | ASV2648 | 4.01 | 1.19E-12 | Eukaryota | Myzozoa | Dinophyceae | Amphidiniales | Amphidiniaceae |
| | ASV65 | 3.99 | 2.27E-05 | Eukaryota | Ochrophyta | Chrysophyceae | Synurales | Mallomonadaceae |
| | ASV605 | 3.90 | 2.05E-14 | Eukaryota | Myzozoa | Dinophyceae | Suessiales | Suessiaceae |
| | ASV2078 | 3.88 | 9.11E-09 | Eukaryota | Bacillariophyta | Mediophyceae | Stephanodiscales | Stephanodiscaceae |
| | ASV2604 | 3.87 | 6.33E-07 | Eukaryota | Cryptophyta | Cryptophyceae | Pyrenomonadales | Geminigeraceae |
| | ASV1300 | 3.86 | 1.21E-05 | Eukaryota | Ciliophora | Prostomatea | Prorodontida | Colepidae |
| | ASV2852 | 3.83 | 3.10E-07 | Eukaryota | Haptophyta | Coccolithophyceae | Isochrysidales | Noelaerhabdaceae |
| | ASV321 | 3.83 | 8.93E-07 | Eukaryota | Bacillariophyta | Mediophyceae | Stephanodiscales | Stephanodiscaceae |
| | ASV2465 | 3.79 | 2.07E-13 | Eukaryota | Myzozoa | Dinophyceae | Suessiales | Suessiaceae |
| | ASV2762 | 3.77 | 0.000134 | Eukaryota | Ochrophyta | Chrysophyceae | Synurales | Mallomonadaceae |
| | ASV1575 | 3.76 | 2.92E-06 | Eukaryota | Bacillariophyta | Mediophyceae | Stephanodiscales | Stephanodiscaceae |
| | ASV2109 | 3.62 | 2.53E-10 | Eukaryota | Bacillariophyta | Mediophyceae | Thalassiosirales | Thalassiosiraceae |

| | | | | | | | | |
|---|---|---|---|---|---|---|---|---|
| | ASV2051 | 3.57 | 3.13E-07 | Eukaryota | Ciliophora | Prostomatea | Prorodontida | Colepidae |
| | ASV2784 | 3.56 | 6.17E-08 | Eukaryota | Cercozoa | Protozoa classis incertae sedis | Protozoa ordo incertae sedis | Protozoa familia incertae sedis |
| | ASV465 | 3.55 | 0.000187 | Eukaryota | Telonemia | Telonemia classis ineditae | Telonemida | Telonemia familia ineditae |
| | ASV2564 | 3.45 | 2.45E-05 | Eukaryota | Cryptophyta | Cryptophyceae | Pyrenomonadales | Geminigeraceae |
| | ASV2649 | 3.45 | 3.77E-07 | Eukaryota | Myzozoa | Dinophyceae | Gymnodiniales | Gymnodiniaceae |
| | ASV1639 | 3.44 | 1.16E-05 | Eukaryota | Ochrophyta | Chrysophyceae | Synurales | Mallomonadaceae |
| | ASV606 | 3.43 | 1.12E-06 | Eukaryota | Ciliophora | Prostomatea | Prorodontida | Colepidae |
| | ASV956 | 3.42 | 0.000388 | Eukaryota | Telonemia | Telonemia classis ineditae | Telonemida | Telonemia familia ineditae |
| | ASV1867 | 3.40 | 6.43E-12 | Eukaryota | Ciliophora | Litostomatea | Haptorida | Didiniidae |
| | ASV495 | 3.40 | 8.36E-06 | Eukaryota | Cercozoa | Thecofilosea | Cryomonadida | Protaspidae |
| | ASV2075 | 3.38 | 2.92E-07 | Eukaryota | Ochrophyta | Chrysophyceae | Synurales | Mallomonadaceae |
| | ASV2644 | 3.36 | 3.47E-07 | Eukaryota | Oomycota | Peronosporea | Peronosporales | Peronosporaceae |
| | ASV2351 | 3.35 | 1.82E-07 | Fungi | Chytridiomycota | Chytridiomycetes | Chytridiales | Chytridiaceae |
| | ASV1354 | 3.34 | 2.34E-11 | Eukaryota | Myzozoa | Dinophyceae | Suessiales | Suessiaceae |
| | ASV814 | 3.33 | 3.13E-05 | Eukaryota | Cryptophyta | Cryptophyceae | Pyrenomonadales | Geminigeraceae |
| | ASV1821 | 3.32 | 1.38E-05 | Eukaryota | Ochrophyta | Chrysophyceae | Synurales | Mallomonadaceae |
| | ASV1335 | 3.30 | 1.09E-08 | Eukaryota | Ochrophyta | Chrysophyceae | Synurales | Mallomonadaceae |
| | ASV542 | 3.27 | 7.98E-07 | Eukaryota | Myzozoa | Dinophyceae | Gymnodiniales | Gymnodiniaceae |
| | ASV813 | 3.25 | 1.83E-09 | Eukaryota | Ochrophyta | Chrysophyceae | Synurales | Mallomonadaceae |
| | ASV2164 | 3.18 | 1.61E-07 | Eukaryota | Ciliophora | Litostomatea | Haptorida | Spathidiidae |
| | ASV683 | 3.17 | 1.12E-09 | Eukaryota | Cercozoa | Thecofilosea | Cryomonadida | Protaspidae |
| | ASV2211 | 3.17 | 0.000105 | Eukaryota | Ochrophyta | Chrysophyceae | Synurales | Mallomonadaceae |
| | ASV2703 | 3.17 | 0.002314 | Eukaryota | Telonemia | Telonemia classis ineditae | Telonemida | Telonemia familia ineditae |
| | ASV933 | 3.15 | 0.000154 | Eukaryota | Ciliophora | Prostomatea | Prorodontida | Colepidae |
| | ASV1205 | 3.13 | 9.69E-11 | Eukaryota | Ochrophyta | Chrysophyceae | Chromulinales | Paraphysomonadaceae |
| | ASV297 | 3.11 | 1.71E-07 | Eukaryota | Oomycota | Peronosporea | Peronosporales | Peronosporaceae |
| | ASV1784 | 3.10 | 3.71E-05 | Eukaryota | Cryptophyta | Cryptophyceae | Pyrenomonadales | Geminigeraceae |
| | ASV1432 | 3.07 | 5.00E-07 | Eukaryota | Ciliophora | Prostomatea | Prorodontida | Colepidae |
| | ASV1591 | 3.07 | 6.48E-07 | Fungi | Chytridiomycota | Chytridiomycetes | Chytridiales | Chytridiaceae |
| | ASV2879 | 3.05 | 5.38E-06 | Fungi | Chytridiomycota | Chytridiomycetes | Chytridiales | Chytridiaceae |

| | | | | | | | | |
|---|---|---|---|---|---|---|---|---|
| | ASV2494 | 3.05 | 2.88E-06 | Eukaryota | Myzozoa | Dinophyceae | Gymnodiniales | Gymnodiniaceae |
| | ASV987 | 3.03 | 6.70E-10 | Eukaryota | Myzozoa | Dinophyceae | Suessiales | Suessiaceae |
| | ASV2709 | 3.03 | 7.73E-08 | Fungi | Chytridiomycota | Chytridiomycetes | Chytridiales | Chytridiaceae |
| | ASV1536 | 3.01 | 3.19E-11 | Eukaryota | Ciliophora | Litostomatea | Haptorida | Didiniidae |
| | ASV865 | 3.01 | 2.12E-07 | Eukaryota | Bacillariophyta | Mediophyceae | Stephanodiscales | Stephanodiscaceae |
| | ASV2805 | 3.01 | 0.001127 | Eukaryota | Cercozoa | Thecofilosea | Cryomonadida | Protaspidae |
| | ASV254 | 3.01 | 0.003915 | Eukaryota | Telonemia | Telonemia classis ineditae | Telonemida | Telonemia familia ineditae |
| | ASV1133 | 3.00 | 4.38E-10 | Eukaryota | Ciliophora | Litostomatea | Haptorida | Didiniidae |
| | ASV1473 | 3.00 | 6.75E-06 | Eukaryota | Cercozoa | Protozoa classis incertae sedis | Protozoa ordo incertae sedis | Protozoa familia incertae sedis |
| | ASV277 | 3.00 | 8.30E-07 | Eukaryota | Cercozoa | Protozoa classis incertae sedis | Protozoa ordo incertae sedis | Protozoa familia incertae sedis |
| | ASV527 | 2.99 | 4.55E-12 | Eukaryota | Ciliophora | Prostomatea | Prorodontida | Colepidae |
| | ASV2766 | 2.99 | 1.05E-05 | Eukaryota | Ciliophora | Prostomatea | Prorodontida | Colepidae |
| | ASV2408 | 2.96 | 1.33E-06 | Eukaryota | Ciliophora | Litostomatea | Haptorida | Spathidiidae |
| | ASV2343 | 2.95 | 0.000193 | Eukaryota | Ochrophyta | Chrysophyceae | Synurales | Mallomonadaceae |
| | ASV2483 | 2.94 | 6.02E-05 | Eukaryota | Cercozoa | Protozoa classis incertae sedis | Protozoa ordo incertae sedis | Protozoa familia incertae sedis |
| | ASV1760 | 2.94 | 4.70E-05 | Eukaryota | Bacillariophyta | Mediophyceae | Stephanodiscales | Stephanodiscaceae |
| | ASV723 | 2.92 | 0.000117 | Eukaryota | Cercozoa | Thecofilosea | Cryomonadida | Protaspidae |
| | ASV518 | 2.91 | 0.000181 | Eukaryota | Bacillariophyta | Mediophyceae | Stephanodiscales | Stephanodiscaceae |
| | ASV69 | 2.91 | 4.70E-05 | Eukaryota | Ciliophora | Prostomatea | Prorodontida | Colepidae |
| | ASV2288 | 2.89 | 0.000282 | Eukaryota | Bacillariophyta | Mediophyceae | Stephanodiscales | Stephanodiscaceae |
| | ASV110 | 2.87 | 7.36E-11 | Eukaryota | Ciliophora | Litostomatea | Haptorida | Didiniidae |
| | ASV2828 | 2.86 | 0.001558 | Eukaryota | Telonemia | Telonemia classis ineditae | Telonemida | Telonemia familia ineditae |
| | ASV1596 | 2.81 | 0.000207 | Eukaryota | Cercozoa | Thecofilosea | Cryomonadida | Protaspidae |
| | ASV2392 | 2.78 | 0.005765 | Eukaryota | Ciliophora | Spirotrichea | Choreotrichida | Strobilidiidae |
| | ASV53 | 2.77 | 3.61E-06 | Eukaryota | Myzozoa | Dinophyceae | Peridiniales | Pfiesteriaceae |
| | ASV1268 | 2.77 | 2.01E-13 | Eukaryota | Ciliophora | Prostomatea | Prorodontida | Colepidae |
| | ASV119 | 2.76 | 1.58E-05 | Eukaryota | Bacillariophyta | Mediophyceae | Stephanodiscales | Stephanodiscaceae |
| | ASV28 | 2.76 | 0.000213 | Eukaryota | Cercozoa | Protozoa classis incertae sedis | Protozoa ordo incertae sedis | Protozoa familia incertae sedis |
| | ASV1528 | 2.75 | 1.69E-09 | Eukaryota | Ciliophora | Litostomatea | Haptorida | Spathidiidae |
| | ASV319 | 2.75 | 1.33E-05 | Eukaryota | Ochrophyta | Chrysophyceae | Synurales | Mallomonadaceae |

| | | | | | | | | |
|---|---|---|---|---|---|---|---|---|
| | ASV1445 | 2.72 | 7.44E-08 | Eukaryota | Cercozoa | Thecofilosea | Cryomonadida | Protaspidae |
| | ASV1707 | 2.72 | 0.005481 | Eukaryota | Telonemia | Telonemia classis ineditae | Telonemida | Telonemia familia ineditae |
| | ASV2693 | 2.72 | 0.000271 | Eukaryota | Bacillariophyta | Mediophyceae | Stephanodiscales | Stephanodiscaceae |
| | ASV100 | 2.71 | 0.001193 | Eukaryota | Cryptophyta | Cryptophyceae | Kathablepharidacea | Katablepharidaceae |
| | ASV1254 | 2.70 | 0.001305 | Eukaryota | Cryptophyta | Cryptophyceae | Pyrenomonadales | Geminigeraceae |
| | ASV350 | 2.68 | 0.001122 | Eukaryota | Ciliophora | Prostomatea | Prorodontida | Colepidae |
| | ASV1477 | 2.66 | 0.000574 | Eukaryota | Cryptophyta | Cryptophyceae | Kathablepharidacea | Katablepharidaceae |
| | ASV1678 | 2.65 | 0.002314 | Eukaryota | Bacillariophyta | Bacillariophyceae | Rhabdonematales | Tabellariaceae |
| | ASV1085 | 2.64 | 2.04E-05 | Eukaryota | Myzozoa | Dinophyceae | Gymnodiniales | Gymnodiniaceae |
| | ASV1629 | 2.64 | 8.24E-06 | Eukaryota | Cercozoa | Thecofilosea | Cryomonadida | Protaspidae |
| | ASV2481 | 2.64 | 0.00169 | Eukaryota | Cercozoa | Thecofilosea | Cryomonadida | Protaspidae |
| | ASV6 | 2.64 | 1.36E-05 | Eukaryota | Myzozoa | Dinophyceae | Peridiniales | Pfiesteriaceae |
| | ASV263 | 2.63 | 0.000104 | Eukaryota | Bigyra | Bikosea | Bicosoecida | Bicosoecidae |
| | ASV1882 | 2.62 | 1.40E-06 | Eukaryota | Cercozoa | Protozoa classis incertae sedis | Protozoa ordo incertae sedis | Protozoa familia incertae sedis |
| | ASV1418 | 2.62 | 5.09E-06 | Eukaryota | Myzozoa | Dinophyceae | Peridiniales | Pfiesteriaceae |
| | ASV841 | 2.61 | 0.000294 | Eukaryota | Cercozoa | Protozoa classis incertae sedis | Protozoa ordo incertae sedis | Protozoa familia incertae sedis |
| | ASV2151 | 2.61 | 0.000705 | Eukaryota | Bacillariophyta | Mediophyceae | Stephanodiscales | Stephanodiscaceae |
| | ASV1054 | 2.59 | 1.92E-07 | Eukaryota | Oomycota | Peronosporea | Peronosporales | Peronosporaceae |
| | ASV1402 | 2.58 | 0.000268 | Eukaryota | Bigyra | Bikosea | Bicosoecida | Bicosoecidae |
| | ASV366 | 2.57 | 5.15E-09 | Eukaryota | Cercozoa | Granofilosea | Leucodictyida | Massisteriidae |
| | ASV1518 | 2.57 | 8.20E-10 | Eukaryota | Myzozoa | Dinophyceae | Thoracosphaerales | Thoracosphaeraceae |
| | ASV1060 | 2.55 | 2.29E-10 | Eukaryota | Bacillariophyta | Mediophyceae | Stephanodiscales | Stephanodiscaceae |
| | ASV253 | 2.54 | 7.35E-05 | Eukaryota | Cercozoa | Sarcomonadea | Cercomonadida | Cercomonadidae |
| | ASV2027 | 2.54 | 0.003299 | Eukaryota | Bacillariophyta | Bacillariophyceae | Rhabdonematales | Tabellariaceae |
| | ASV225 | 2.54 | 1.13E-10 | Eukaryota | Bacillariophyta | Mediophyceae | Stephanodiscales | Stephanodiscaceae |
| | ASV2094 | 2.54 | 0.001013 | Eukaryota | Cryptophyta | Cryptophyceae | Pyrenomonadales | Geminigeraceae |
| | ASV2192 | 2.53 | 1.50E-10 | Eukaryota | Bacillariophyta | Mediophyceae | Stephanodiscales | Stephanodiscaceae |
| | ASV1218 | 2.52 | 0.000286 | Eukaryota | Bacillariophyta | Mediophyceae | Stephanodiscales | Stephanodiscaceae |
| | ASV2253 | 2.52 | 1.49E-10 | Eukaryota | Bacillariophyta | Mediophyceae | Stephanodiscales | Stephanodiscaceae |
| | ASV1276 | 2.51 | 4.48E-06 | Eukaryota | Ciliophora | Litostomatea | Haptorida | Spathidiidae |

| | | | | | | | | |
|---|---|---|---|---|---|---|---|---|
| | ASV2271 | 2.50 | 3.43E-05 | Eukaryota | Myzozoa | Dinophyceae | Peridiniales | Pfiesteriaceae |
| | ASV2170 | 2.50 | 0.002905 | Eukaryota | Ciliophora | Prostomatea | Prorodontida | Colepidae |
| | ASV1437 | 2.50 | 6.40E-09 | Eukaryota | Ciliophora | Litostomatea | Haptorida | Spathidiidae |
| | ASV2416 | 2.49 | 3.17E-08 | Eukaryota | Myzozoa | Dinophyceae | Thoracosphaerales | Thoracosphaeraceae |
| | ASV2715 | 2.49 | 0.00104 | Eukaryota | Cryptophyta | Cryptophyceae | Pyrenomonadales | Geminigeraceae |
| | ASV1269 | 2.48 | 0.00169 | Eukaryota | Cercozoa | Thecofilosea | Cryomonadida | Protaspidae |
| | ASV2725 | 2.48 | 3.45E-11 | Eukaryota | Bacillariophyta | Mediophyceae | Stephanodiscales | Stephanodiscaceae |
| | ASV893 | 2.48 | 1.16E-10 | Eukaryota | Bacillariophyta | Mediophyceae | Stephanodiscales | Stephanodiscaceae |
| | ASV1219 | 2.48 | 0.001102 | Eukaryota | Ciliophora | Prostomatea | Prorodontida | Colepidae |
| | ASV1135 | 2.48 | 4.93E-11 | Eukaryota | Bacillariophyta | Mediophyceae | Stephanodiscales | Stephanodiscaceae |
| | ASV2711 | 2.46 | 0.000712 | Eukaryota | Bacillariophyta | Bacillariophyceae | Rhabdonematales | Tabellariaceae |
| | ASV2670 | 2.44 | 1.85E-05 | Eukaryota | Cercozoa | Protozoa classis incertae sedis | Protozoa ordo incertae sedis | Protozoa familia incertae sedis |
| | ASV116 | 2.43 | 5.52E-07 | Eukaryota | Ochrophyta | Chrysophyceae | Synurales | Mallomonadaceae |
| | ASV2653 | 2.41 | 0.000574 | Eukaryota | Cryptophyta | Cryptophyceae | Kathablepharidacea | Katablepharidaceae |
| | ASV1705 | 2.40 | 7.07E-06 | Eukaryota | Ciliophora | Prostomatea | Prorodontida | Colepidae |
| | ASV1511 | 2.39 | 0.000976 | Eukaryota | Bacillariophyta | Mediophyceae | Stephanodiscales | Stephanodiscaceae |
| | ASV675 | 2.39 | 4.80E-05 | Eukaryota | Bacillariophyta | Mediophyceae | Stephanodiscales | Stephanodiscaceae |
| | ASV2791 | 2.37 | 7.36E-11 | Eukaryota | Bacillariophyta | Mediophyceae | Stephanodiscales | Stephanodiscaceae |
| | ASV719 | 2.37 | 8.82E-06 | Eukaryota | Cercozoa | Thecofilosea | Cryomonadida | Protaspidae |
| | ASV2132 | 2.36 | 0.006794 | Eukaryota | Ciliophora | Prostomatea | Prorodontida | Colepidae |
| | ASV1294 | 2.34 | 0.001122 | Eukaryota | Cryptophyta | Cryptophyceae | Kathablepharidacea | Katablepharidaceae |
| | ASV2412 | 2.33 | 0.000615 | Eukaryota | Cryptophyta | Cryptophyceae | Pyrenomonadales | Geminigeraceae |
| | ASV972 | 2.33 | 1.06E-05 | Eukaryota | Ciliophora | Litostomatea | Haptorida | Spathidiidae |
| | ASV2432 | 2.31 | 2.88E-06 | Eukaryota | Haptophyta | Coccolithophyceae | Prymnesiales | Chrysochromulinaceae |
| | ASV211 | 2.29 | 2.02E-06 | Eukaryota | Oomycota | Peronosporea | Peronosporales | Peronosporaceae |
| | ASV2641 | 2.26 | 4.18E-05 | Eukaryota | Cercozoa | Protozoa classis incertae sedis | Protozoa ordo incertae sedis | Protozoa familia incertae sedis |
| | ASV470 | 2.25 | 0.002368 | Eukaryota | Haptophyta | Coccolithophyceae | Prymnesiales | Chrysochromulinaceae |
| | ASV1369 | 2.24 | 5.05E-05 | Eukaryota | Ciliophora | Oligohymenophorea | Hymenostomatida | Tetrahymenidae |
| | ASV1547 | 2.23 | 0.003861 | Eukaryota | Ciliophora | Oligohymenophorea | Pleuronematida | Histiobalantiidae |
| | ASV1545 | 2.22 | 0.000498 | Eukaryota | Cryptophyta | Cryptophyceae | Kathablepharidacea | Katablepharidaceae |

| | | | | | | | | |
|---|---|---|---|---|---|---|---|---|
| | ASV1721 | 2.17 | 6.86E-07 | Eukaryota | Cercozoa | Protozoa classis incertae sedis | Protozoa ordo incertae sedis | Protozoa familia incertae sedis |
| | ASV1650 | 2.17 | 0.008898 | Eukaryota | Haptophyta | Coccolithophyceae | Prymnesiales | Chrysochromulinaceae |
| | ASV258 | 2.15 | 2.79E-06 | Eukaryota | Bigyra | incertae sedis | incertae sedis | Pseudophyllomitidae |
| | ASV2687 | 2.14 | 0.005822 | Eukaryota | Ciliophora | Prostomatea | Prorodontida | Colepidae |
| | ASV2504 | 2.14 | 0.000947 | Eukaryota | Cercozoa | Thecofilosea | Cryomonadida | Protaspidae |
| | ASV1953 | 2.13 | 6.67E-07 | Eukaryota | Ochrophyta | Chrysophyceae | Synurales | Mallomonadaceae |
| | ASV265 | 2.13 | 0.000308 | Eukaryota | Bacillariophyta | Mediophyceae | Stephanodiscales | Stephanodiscaceae |
| | ASV589 | 2.12 | 0.00052 | Eukaryota | Ciliophora | Spirotrichea | Choreotrichida | Strobilidiidae |
| | ASV2503 | 2.10 | 9.94E-07 | Eukaryota | Ciliophora | Litostomatea | Haptorida | Spathidiidae |
| | ASV921 | 2.10 | 0.004844 | Eukaryota | Cercozoa | Thecofilosea | Cryomonadida | Protaspidae |
| | ASV1548 | 2.09 | 0.000673 | Eukaryota | Ciliophora | Spirotrichea | Choreotrichida | Strobilidiidae |
| | ASV2793 | 2.08 | 0.009101 | Eukaryota | Ciliophora | Spirotrichea | Choreotrichida | Strobilidiidae |
| | ASV2768 | 2.08 | 0.000308 | Eukaryota | Bacillariophyta | Mediophyceae | Stephanodiscales | Stephanodiscaceae |
| | ASV337 | 2.08 | 0.005909 | Eukaryota | Ciliophora | Litostomatea | Haptorida | Spathidiidae |
| | ASV1625 | 2.08 | 0.003438 | Eukaryota | Ciliophora | Prostomatea | Prorodontida | Colepidae |
| | ASV2396 | 2.07 | 0.000724 | Eukaryota | Ciliophora | Spirotrichea | Choreotrichida | Strobilidiidae |
| | ASV2673 | 2.07 | 0.002623 | Eukaryota | Ciliophora | Litostomatea | Haptorida | Spathidiidae |
| | ASV2463 | 2.06 | 0.00064 | Eukaryota | Ciliophora | Spirotrichea | Choreotrichida | Strobilidiidae |
| | ASV738 | 2.06 | 5.08E-06 | Eukaryota | Ochrophyta | Chrysophyceae | Synurales | Mallomonadaceae |
| | ASV2679 | 2.05 | 5.05E-05 | Eukaryota | Ciliophora | Oligohymenophorea | Hymenostomatida | Tetrahymenidae |
| | ASV587 | 2.01 | 6.27E-05 | Eukaryota | Cercozoa | Protozoa classis incertae sedis | Protozoa ordo incertae sedis | Protozoa familia incertae sedis |
| | ASV134 | 2.00 | 0.001532 | Eukaryota | Cercozoa | Thecofilosea | Cryomonadida | Protaspidae |
| 18S-PP | ASV207 | 9.82 | 1.10E-20 | Eukaryota | Ochrophyta | Chrysophyceae | Chromulinales | Paraphysomonadaceae |
| | ASV2643 | 9.21 | 1.08E-17 | Eukaryota | Ochrophyta | Chrysophyceae | Chromulinales | Paraphysomonadaceae |
| | ASV1729 | 8.45 | 5.11E-20 | Eukaryota | Ochrophyta | Chrysophyceae | Chromulinales | Paraphysomonadaceae |
| | ASV2264 | 8.31 | 1.72E-18 | Eukaryota | Ochrophyta | Chrysophyceae | Chromulinales | Paraphysomonadaceae |
| | ASV499 | 7.81 | 3.33E-14 | Eukaryota | Myzozoa | Dinophyceae | Gymnodiniales | Gymnodiniaceae |
| | ASV1860 | 7.78 | 4.04E-14 | Eukaryota | Myzozoa | Dinophyceae | Gymnodiniales | Gymnodiniaceae |
| | ASV1827 | 7.38 | 1.84E-13 | Eukaryota | Myzozoa | Dinophyceae | Gymnodiniales | Gymnodiniaceae |

| | | | | | | | | |
|---|---|---|---|---|---|---|---|---|
| | ASV2100 | 7.30 | 2.89E-19 | Eukaryota | Myzozoa | Dinophyceae | Gymnodiniales | Gymnodiniaceae |
| | ASV1680 | 7.21 | 1.95E-12 | Eukaryota | Myzozoa | Dinophyceae | Gymnodiniales | Gymnodiniaceae |
| | ASV229 | 7.15 | 2.40E-14 | Eukaryota | Myzozoa | Dinophyceae | Gymnodiniales | Gymnodiniaceae |
| | ASV1297 | 7.14 | 1.29E-13 | Eukaryota | Myzozoa | Dinophyceae | Prorocentrales | Prorocentraceae |
| | ASV2759 | 7.11 | 3.53E-19 | Eukaryota | Myzozoa | Dinophyceae | Gymnodiniales | Gymnodiniaceae |
| | ASV2169 | 7.09 | 4.17E-17 | Eukaryota | Myzozoa | Dinophyceae | Gymnodiniales | Gymnodiniaceae |
| | ASV2182 | 7.03 | 5.06E-14 | Eukaryota | Myzozoa | Dinophyceae | Gymnodiniales | Gymnodiniaceae |
| | ASV1543 | 7.01 | 3.66E-21 | Eukaryota | Myzozoa | Dinophyceae | Gymnodiniales | Gymnodiniaceae |
| | ASV1396 | 6.97 | 5.23E-13 | Eukaryota | Myzozoa | Dinophyceae | Prorocentrales | Prorocentraceae |
| | ASV313 | 6.91 | 9.47E-14 | Eukaryota | Myzozoa | Dinophyceae | Gymnodiniales | Gymnodiniaceae |
| | ASV1962 | 6.90 | 1.43E-12 | Eukaryota | Ciliophora | Oligotrichea | Oligotrichida | Strombidiidae |
| | ASV1674 | 6.88 | 5.96E-13 | Eukaryota | Ciliophora | Oligotrichea | Oligotrichida | Strombidiidae |
| | ASV1915 | 6.87 | 5.24E-12 | Eukaryota | Ciliophora | Oligotrichea | Oligotrichida | Strombidiidae |
| | ASV2792 | 6.80 | 4.28E-09 | Eukaryota | Ciliophora | Prostomatea | Prorodontida | Colepidae |
| | ASV1102 | 6.70 | 3.45E-14 | Eukaryota | Myzozoa | Dinophyceae | Gymnodiniales | Gymnodiniaceae |
| | ASV582 | 6.66 | 5.23E-13 | Eukaryota | Myzozoa | Dinophyceae | Prorocentrales | Prorocentraceae |
| | ASV1131 | 6.58 | 7.20E-10 | Eukaryota | Ciliophora | Prostomatea | Prorodontida | Colepidae |
| | ASV2740 | 6.56 | 4.42E-09 | Eukaryota | Ciliophora | Prostomatea | Prorodontida | Colepidae |
| | ASV862 | 6.52 | 5.23E-13 | Eukaryota | Myzozoa | Dinophyceae | Prorocentrales | Prorocentraceae |
| | ASV687 | 6.50 | 1.20E-08 | Eukaryota | Ciliophora | Prostomatea | Prorodontida | Colepidae |
| | ASV797 | 6.33 | 4.11E-11 | Eukaryota | Ciliophora | Oligotrichea | Oligotrichida | Strombidiidae |
| | ASV2120 | 5.96 | 5.48E-08 | Eukaryota | Chlorophyta | Mamiellophyceae | Dolichomastigales | Dolichomastigaceae |
| | ASV2843 | 5.95 | 8.35E-09 | Eukaryota | Chlorophyta | Mamiellophyceae | Dolichomastigales | Dolichomastigaceae |
| | ASV1308 | 5.90 | 1.09E-08 | Eukaryota | Chlorophyta | Mamiellophyceae | Dolichomastigales | Dolichomastigaceae |
| | ASV2789 | 5.88 | 1.15E-11 | Eukaryota | Myzozoa | Dinophyceae | Amphidiniales | Amphidiniaceae |
| | ASV2437 | 5.64 | 2.51E-10 | Eukaryota | Myzozoa | Dinophyceae | Amphidiniales | Amphidiniaceae |
| | ASV547 | 5.61 | 2.40E-14 | Eukaryota | Ochrophyta | Chrysophyceae | Chromulinales | Paraphysomonadaceae |
| | ASV2648 | 5.61 | 2.04E-12 | Eukaryota | Myzozoa | Dinophyceae | Amphidiniales | Amphidiniaceae |

| | | | | | | | | |
|---|---|---|---|---|---|---|---|---|
| | ASV1340 | 5.54 | 2.68E-11 | Eukaryota | Myzozoa | Dinophyceae | Amphidiniales | Amphidiniaceae |
| | ASV1757 | 5.51 | 5.58E-07 | Eukaryota | Chlorophyta | Mamiellophyceae | Dolichomastigales | Dolichomastigaceae |
| | ASV684 | 5.51 | 9.47E-14 | Eukaryota | Ochrophyta | Chrysophyceae | Chromulinales | Paraphysomonadaceae |
| | ASV1854 | 5.24 | 1.29E-10 | Eukaryota | Chlorophyta | Trebouxiophyceae | Chlorellales | Chlorellaceae |
| | ASV2355 | 5.22 | 2.64E-11 | Eukaryota | Chlorophyta | Trebouxiophyceae | Chlorellales | Chlorellaceae |
| | ASV483 | 5.21 | 1.31E-09 | Eukaryota | Ciliophora | Spirotrichea | Stichotrichida | Amphisiellidae |
| | ASV2062 | 5.12 | 9.57E-17 | Eukaryota | Myzozoa | Dinophyceae | Gymnodiniales | Gymnodiniaceae |
| | ASV1647 | 5.11 | 5.55E-11 | Eukaryota | Chlorophyta | Trebouxiophyceae | Chlorellales | Chlorellaceae |
| | ASV1328 | 5.09 | 7.56E-12 | Eukaryota | Myzozoa | Dinophyceae | Peridiniales | Pfiesteriaceae |
| | ASV2425 | 5.07 | 4.04E-12 | Eukaryota | Myzozoa | Dinophyceae | Peridiniales | Pfiesteriaceae |
| | ASV2277 | 5.06 | 1.64E-09 | Eukaryota | Cercozoa | Protozoa classis incertae sedis | Protozoa ordo incertae sedis | Protozoa familia incertae sedis |
| | ASV1544 | 5.05 | 1.65E-09 | Eukaryota | Ciliophora | Spirotrichea | Stichotrichida | Amphisiellidae |
| | ASV942 | 4.99 | 1.99E-12 | Eukaryota | Myzozoa | Dinophyceae | Peridiniales | Pfiesteriaceae |
| | ASV1168 | 4.97 | 5.24E-12 | Eukaryota | Cercozoa | Protozoa classis incertae sedis | Protozoa ordo incertae sedis | Protozoa familia incertae sedis |
| | ASV723 | 4.96 | 3.25E-09 | Eukaryota | Cercozoa | Thecofilosea | Cryomonadida | Protaspidae |
| | ASV2401 | 4.83 | 1.45E-10 | Eukaryota | Chlorophyta | Trebouxiophyceae | Chlorellales | Chlorellaceae |
| | ASV1819 | 4.82 | 3.40E-08 | Eukaryota | Ochrophyta | Chrysophyceae | Chromulinales | Paraphysomonadaceae |
| | ASV2060 | 4.81 | 3.14E-09 | Eukaryota | Ciliophora | Spirotrichea | Stichotrichida | Amphisiellidae |
| | ASV921 | 4.77 | 1.05E-07 | Eukaryota | Cercozoa | Thecofilosea | Cryomonadida | Protaspidae |
| | ASV2582 | 4.76 | 8.78E-11 | Eukaryota | Choanozoa | Filasterea | Filasterea ordo incertae sedis | Filasterea familia incertae sedis |
| | ASV1306 | 4.75 | 1.49E-14 | Eukaryota | Myzozoa | Dinophyceae | Gymnodiniales | Gymnodiniaceae |
| | ASV2481 | 4.75 | 3.29E-07 | Eukaryota | Cercozoa | Thecofilosea | Cryomonadida | Protaspidae |
| | ASV853 | 4.74 | 5.44E-10 | Eukaryota | Ciliophora | Spirotrichea | Stichotrichida | Amphisiellidae |
| | ASV815 | 4.70 | 9.47E-14 | Eukaryota | Myzozoa | Dinophyceae | Gymnodiniales | Gymnodiniaceae |
| | ASV2471 | 4.64 | 5.93E-07 | Eukaryota | Ochrophyta | Chrysophyceae | Chromulinales | Chrysolepidomonadaceae |
| | ASV2206 | 4.60 | 4.27E-10 | Eukaryota | Choanozoa | Filasterea | Filasterea ordo incertae sedis | Filasterea familia incertae sedis |
| | ASV6 | 4.59 | 3.28E-07 | Eukaryota | Myzozoa | Dinophyceae | Peridiniales | Pfiesteriaceae |
| | ASV1999 | 4.57 | 8.98E-08 | Eukaryota | Choanozoa | Choanoflagellata | Craspedida | Codonosigaceae |

| | | | | | | | | |
|---|---|---|---|---|---|---|---|---|
| | ASV1320 | 4.52 | 9.85E-08 | Eukaryota | Ochrophyta | Chrysophyceae | Chromulinales | Paraphysomonadaceae |
| | ASV897 | 4.48 | 6.12E-08 | Eukaryota | Choanozoa | Choanoflagellata | Craspedida | Codonosigaceae |
| | ASV1184 | 4.44 | 7.20E-10 | Eukaryota | Choanozoa | Filasterea | Filasterea ordo incertae sedis | Filasterea familia incertae sedis |
| | ASV620 | 4.44 | 3.45E-06 | Eukaryota | Ciliophora | Prostomatea | Prorodontida | Colepidae |
| | ASV1418 | 4.44 | 9.78E-08 | Eukaryota | Myzozoa | Dinophyceae | Peridiniales | Pfiesteriaceae |
| | ASV1609 | 4.42 | 7.93E-06 | Eukaryota | Ciliophora | Prostomatea | Prorodontida | Colepidae |
| | ASV2271 | 4.40 | 9.01E-07 | Eukaryota | Myzozoa | Dinophyceae | Peridiniales | Pfiesteriaceae |
| | ASV53 | 4.39 | 1.53E-07 | Eukaryota | Myzozoa | Dinophyceae | Peridiniales | Pfiesteriaceae |
| | ASV2466 | 4.37 | 1.45E-10 | Eukaryota | Myzozoa | Dinophyceae | Peridiniales | Pfiesteriaceae |
| | ASV2791 | 4.37 | 4.06E-06 | Eukaryota | Bacillariophyta | Mediophyceae | Stephanodiscales | Stephanodiscaceae |
| | ASV1300 | 4.37 | 2.32E-06 | Eukaryota | Ciliophora | Prostomatea | Prorodontida | Colepidae |
| | ASV232 | 4.33 | 2.62E-10 | Eukaryota | Bacillariophyta | Mediophyceae | Thalassiosirales | Thalassiosiraceae |
| | ASV1205 | 4.28 | 9.47E-14 | Eukaryota | Ochrophyta | Chrysophyceae | Chromulinales | Paraphysomonadaceae |
| | ASV1294 | 4.26 | 7.71E-07 | Eukaryota | Cryptophyta | Cryptophyceae | Kathablepharidacea | Katablepharidaceae |
| | ASV2075 | 4.23 | 1.71E-05 | Eukaryota | Ochrophyta | Chrysophyceae | Synurales | Mallomonadaceae |
| | ASV724 | 4.20 | 4.34E-05 | Eukaryota | Ciliophora | Prostomatea | Prorodontida | Colepidae |
| | ASV241 | 4.19 | 7.20E-10 | Eukaryota | Choanozoa | Filasterea | Filasterea ordo incertae sedis | Filasterea familia incertae sedis |
| | ASV225 | 4.18 | 1.06E-05 | Eukaryota | Bacillariophyta | Mediophyceae | Stephanodiscales | Stephanodiscaceae |
| | ASV1994 | 4.11 | 8.15E-16 | Eukaryota | Myzozoa | Dinophyceae | Gymnodiniales | Gymnodiniaceae |
| | ASV319 | 4.09 | 1.66E-05 | Eukaryota | Ochrophyta | Chrysophyceae | Synurales | Mallomonadaceae |
| | ASV733 | 4.05 | 1.26E-11 | Eukaryota | Cercozoa | Protozoa classis incertae sedis | Protozoa ordo incertae sedis | Protozoa familia incertae sedis |
| | ASV996 | 4.04 | 1.59E-09 | Eukaryota | Bacillariophyta | Mediophyceae | Thalassiosirales | Thalassiosiraceae |
| | ASV615 | 4.04 | 2.15E-07 | Eukaryota | Choanozoa | Choanoflagellata | Craspedida | Codonosigaceae |
| | ASV841 | 4.03 | 8.87E-09 | Eukaryota | Cercozoa | Protozoa classis incertae sedis | Protozoa ordo incertae sedis | Protozoa familia incertae sedis |
| | ASV100 | 4.03 | 5.35E-06 | Eukaryota | Cryptophyta | Cryptophyceae | Kathablepharidacea | Katablepharidaceae |
| | ASV721 | 4.02 | 1.94E-05 | Eukaryota | Ochrophyta | Chrysophyceae | Chromulinales | Chrysolepidomonadaceae |
| | ASV2725 | 4.01 | 1.40E-06 | Eukaryota | Bacillariophyta | Mediophyceae | Stephanodiscales | Stephanodiscaceae |
| | ASV1473 | 4.00 | 3.83E-09 | Eukaryota | Cercozoa | Protozoa classis incertae sedis | Protozoa ordo incertae sedis | Protozoa familia incertae sedis |

| | | | | | | | | |
|---|---|---|---|---|---|---|---|---|
| | ASV205 | 3.99 | 2.96E-07 | Eukaryota | Ochrophyta | Chrysophyceae | Chromulinales | Chrysolepidomonadaceae |
| | ASV1825 | 3.95 | 1.57E-06 | Eukaryota | Choanozoa | Choanoflagellata | Craspedida | Codonosigaceae |
| | ASV2198 | 3.94 | 3.31E-07 | Eukaryota | Ochrophyta | Chrysophyceae | Chromulinales | Chrysolepidomonadaceae |
| | ASV819 | 3.92 | 1.58E-06 | Eukaryota | Cercozoa | Protozoa classis incertae sedis | Protozoa ordo incertae sedis | Protozoa familia incertae sedis |
| | ASV713 | 3.80 | 5.51E-08 | Eukaryota | Myzozoa | Dinophyceae | Gymnodiniales | Gymnodiniaceae |
| | ASV956 | 3.78 | 0.000426 | Eukaryota | Telonemia | Telonemia classis ineditae | Telonemida | Telonemia familia ineditae |
| | ASV1497 | 3.78 | 9.94E-08 | Eukaryota | Ciliophora | Oligohymenophorea | Thigmotrichida | Ancistridae |
| | ASV1477 | 3.77 | 1.46E-05 | Eukaryota | Cryptophyta | Cryptophyceae | Kathablepharidacea | Katablepharidaceae |
| | ASV2818 | 3.77 | 8.72E-10 | Eukaryota | Ochrophyta | Dictyochophyceae | Pedinellales | Actinomonadaceae |
| | ASV1901 | 3.75 | 2.89E-09 | Eukaryota | Bigyra | incertae sedis | incertae sedis | Pseudophyllomitidae |
| | ASV456 | 3.74 | 1.40E-09 | Eukaryota | Ochrophyta | Dictyochophyceae | Pedinellales | Actinomonadaceae |
| | ASV1053 | 3.73 | 8.24E-10 | Eukaryota | Myzozoa | Dinophyceae | Amphidiniales | Amphidiniaceae |
| | ASV1290 | 3.73 | 4.11E-11 | Eukaryota | Myzozoa | Dinophyceae | Amphidiniales | Amphidiniaceae |
| | ASV2013 | 3.72 | 9.81E-07 | Eukaryota | Haptophyta | Coccolithophyceae | Prymnesiales | Chrysochromulinaceae |
| | ASV2051 | 3.71 | 0.00016 | Eukaryota | Ciliophora | Prostomatea | Prorodontida | Colepidae |
| | ASV465 | 3.70 | 0.000378 | Eukaryota | Telonemia | Telonemia classis ineditae | Telonemida | Telonemia familia ineditae |
| | ASV2766 | 3.70 | 0.000106 | Eukaryota | Ciliophora | Prostomatea | Prorodontida | Colepidae |
| | ASV2644 | 3.66 | 9.34E-12 | Eukaryota | Oomycota | Peronosporea | Peronosporales | Peronosporaceae |
| | ASV2192 | 3.66 | 0.000127 | Eukaryota | Bacillariophyta | Mediophyceae | Stephanodiscales | Stephanodiscaceae |
| | ASV2467 | 3.66 | 5.32E-08 | Eukaryota | Ciliophora | Oligohymenophorea | Thigmotrichida | Ancistridae |
| | ASV401 | 3.66 | 2.84E-07 | Eukaryota | Chlorophyta | Trebouxiophyceae | Chlorellales | Chlorellales incertae sedis |
| | ASV378 | 3.65 | 5.38E-07 | Eukaryota | Ochrophyta | Chrysophyceae | Chromulinales | Paraphysomonadaceae |
| | ASV2483 | 3.65 | 3.68E-06 | Eukaryota | Cercozoa | Protozoa classis incertae sedis | Protozoa ordo incertae sedis | Protozoa familia incertae sedis |
| | ASV1957 | 3.65 | 2.19E-11 | Eukaryota | Ochrophyta | Chrysophyceae | Chromulinales | Paraphysomonadaceae |
| | ASV599 | 3.65 | 5.55E-07 | Eukaryota | Haptophyta | Coccolithophyceae | Prymnesiales | Chrysochromulinaceae |
| | ASV1269 | 3.64 | 1.57E-05 | Eukaryota | Cercozoa | Thecofilosea | Cryomonadida | Protaspidae |
| | ASV738 | 3.61 | 1.72E-05 | Eukaryota | Ochrophyta | Chrysophyceae | Synurales | Mallomonadaceae |
| | ASV1542 | 3.60 | 1.12E-05 | Eukaryota | Cryptophyta | Cryptophyceae | Kathablepharidacea | Katablepharidaceae |

| | | | | | | | | |
|---|---|---|---|---|---|---|---|---|
| | ASV1907 | 3.57 | 5.51E-08 | Eukaryota | Ciliophora | Oligohymenophorea | Thigmotrichida | Ancistridae |
| | ASV31 | 3.56 | 5.39E-07 | Eukaryota | Haptophyta | Coccolithophyceae | Prymnesiales | Chrysochromulinaceae |
| | ASV2432 | 3.51 | 2.54E-06 | Eukaryota | Haptophyta | Coccolithophyceae | Prymnesiales | Chrysochromulinaceae |
| | ASV606 | 3.48 | 0.000554 | Eukaryota | Ciliophora | Prostomatea | Prorodontida | Colepidae |
| | ASV2207 | 3.47 | 9.87E-07 | Eukaryota | Ochrophyta | Chrysophyceae | Chromulinales | Paraphysomonadaceae |
| | ASV812 | 3.46 | 5.02E-09 | Eukaryota | Ochrophyta | Chrysophyceae | Chromulinales | Paraphysomonadaceae |
| | ASV1934 | 3.45 | 3.27E-07 | Eukaryota | Chlorophyta | Trebouxiophyceae | Chlorellales | Chlorellales incertae sedis |
| | ASV981 | 3.45 | 8.02E-07 | Eukaryota | Cercozoa | Thecofilosea | Cryomonadida | Protaspidae |
| | ASV637 | 3.45 | 1.83E-10 | Eukaryota | Myzozoa | Dinophyceae | Amphidiniales | Amphidiniaceae |
| | ASV1432 | 3.45 | 0.000169 | Eukaryota | Ciliophora | Prostomatea | Prorodontida | Colepidae |
| | ASV1953 | 3.44 | 2.76E-05 | Eukaryota | Ochrophyta | Chrysophyceae | Synurales | Mallomonadaceae |
| | ASV1591 | 3.43 | 0.000224 | Fungi | Chytridiomycota | Chytridiomycetes | Chytridiales | Chytridiaceae |
| | ASV1135 | 3.43 | 3.76E-05 | Eukaryota | Bacillariophyta | Mediophyceae | Stephanodiscales | Stephanodiscaceae |
| | ASV1721 | 3.43 | 1.29E-11 | Eukaryota | Cercozoa | Protozoa classis incertae sedis | Protozoa ordo incertae sedis | Protozoa familia incertae sedis |
| | ASV1354 | 3.43 | 5.77E-12 | Eukaryota | Myzozoa | Dinophyceae | Suessiales | Suessiaceae |
| | ASV2351 | 3.42 | 0.000613 | Fungi | Chytridiomycota | Chytridiomycetes | Chytridiales | Chytridiaceae |
| | ASV2879 | 3.41 | 0.00051 | Fungi | Chytridiomycota | Chytridiomycetes | Chytridiales | Chytridiaceae |
| | ASV2703 | 3.38 | 0.001537 | Eukaryota | Telonemia | Telonemia classis ineditae | Telonemida | Telonemia familia ineditae |
| | ASV170 | 3.38 | 3.64E-07 | Eukaryota | Cryptophyta | Cryptophyceae | Pyrenomonadales | Geminigeraceae |
| | ASV893 | 3.37 | 0.000196 | Eukaryota | Bacillariophyta | Mediophyceae | Stephanodiscales | Stephanodiscaceae |
| | ASV1189 | 3.37 | 4.83E-06 | Eukaryota | Cryptophyta | Cryptophyceae | Pyrenomonadales | Geminigeraceae |
| | ASV2465 | 3.35 | 1.01E-13 | Eukaryota | Myzozoa | Dinophyceae | Suessiales | Suessiaceae |
| | ASV2275 | 3.33 | 7.92E-14 | Eukaryota | Myzozoa | Dinophyceae | Gymnodiniales | Gymnodiniaceae |
| | ASV2397 | 3.31 | 2.42E-06 | Eukaryota | Myzozoa | Dinophyceae | Gymnodiniales | Gymnodiniaceae |
| | ASV2345 | 3.29 | 7.58E-06 | Eukaryota | Bacillariophyta | Mediophyceae | Stephanodiscales | Stephanodiscaceae |
| | ASV1871 | 3.28 | 2.70E-12 | Eukaryota | Myzozoa | Dinophyceae | Gymnodiniales | Gymnodiniaceae |
| | ASV2551 | 3.27 | 8.85E-08 | Fungi | Chytridiomycota | Chytridiomycetes | Rhizophydiales | Uebelmesseromycetaceae |
| | ASV2709 | 3.25 | 0.00026 | Fungi | Chytridiomycota | Chytridiomycetes | Chytridiales | Chytridiaceae |

| | | | | | | | | |
|---|---|---|---|---|---|---|---|---|
| | ASV254 | 3.20 | 0.004609 | Eukaryota | Telonemia | Telonemia classis ineditae | Telonemida | Telonemia familia ineditae |
| | ASV2504 | 3.19 | 9.01E-07 | Eukaryota | Cercozoa | Thecofilosea | Cryomonadida | Protaspidae |
| | ASV122 | 3.17 | 0.001178 | Eukaryota | Choanozoa | Choanoflagellata | Craspedida | Codonosigaceae |
| | ASV2475 | 3.13 | 7.10E-11 | Eukaryota | Ochrophyta | Bolidophyceae | Parmales | Triparmaceae |
| | ASV1724 | 3.12 | 4.50E-07 | Fungi | Chytridiomycota | Chytridiomycetes | Rhizophydiales | Uebelmesseromycetaceae |
| | ASV605 | 3.11 | 1.95E-12 | Eukaryota | Myzozoa | Dinophyceae | Suessiales | Suessiaceae |
| | ASV977 | 3.09 | 6.10E-07 | Eukaryota | Chlorophyta | Trebouxiophyceae | Chlorellales | Chlorellales incertae sedis |
| | ASV258 | 3.09 | 8.82E-06 | Eukaryota | Bigyra | incertae sedis | incertae sedis | Pseudophyllomitidae |
| | ASV307 | 3.07 | 0.000963 | Eukaryota | Choanozoa | Choanoflagellata | Craspedida | Codonosigaceae |
| | ASV933 | 3.06 | 0.00045 | Eukaryota | Ciliophora | Prostomatea | Prorodontida | Colepidae |
| | ASV2641 | 3.05 | 2.74E-07 | Eukaryota | Cercozoa | Protozoa classis incertae sedis | Protozoa ordo incertae sedis | Protozoa familia incertae sedis |
| | ASV2637 | 3.02 | 3.28E-06 | Eukaryota | Myzozoa | Dinophyceae | Gymnodiniales | Gymnodiniaceae |
| | ASV907 | 3.02 | 0.000813 | Eukaryota | Cryptophyta | Cryptophyceae | Pyrenomonadales | Geminigeraceae |
| | ASV1663 | 2.99 | 5.46E-12 | Eukaryota | Myzozoa | Dinophyceae | Amphidiniales | Amphidiniaceae |
| | ASV1936 | 2.96 | 3.76E-05 | Eukaryota | Myzozoa | Dinophyceae | Gymnodiniales | Gymnodiniaceae |
| | ASV290 | 2.96 | 1.48E-05 | Eukaryota | Cryptophyta | Cryptophyceae | Pyrenomonadales | Geminigeraceae |
| | ASV297 | 2.95 | 1.00E-08 | Eukaryota | Oomycota | Peronosporea | Peronosporales | Peronosporaceae |
| | ASV2784 | 2.95 | 7.58E-06 | Eukaryota | Cercozoa | Protozoa classis incertae sedis | Protozoa ordo incertae sedis | Protozoa familia incertae sedis |
| | ASV234 | 2.91 | 1.17E-06 | Fungi | Chytridiomycota | Chytridiomycetes | Rhizophydiales | Uebelmesseromycetaceae |
| | ASV1060 | 2.89 | 0.00072 | Eukaryota | Bacillariophyta | Mediophyceae | Stephanodiscales | Stephanodiscaceae |
| | ASV1471 | 2.89 | 2.52E-05 | Eukaryota | Myzozoa | Dinophyceae | Gymnodiniales | Gymnodiniaceae |
| | ASV2094 | 2.87 | 2.73E-06 | Eukaryota | Cryptophyta | Cryptophyceae | Pyrenomonadales | Geminigeraceae |
| | ASV2649 | 2.87 | 3.47E-08 | Eukaryota | Myzozoa | Dinophyceae | Gymnodiniales | Gymnodiniaceae |
| | ASV2035 | 2.85 | 0.000274 | Eukaryota | Haptophyta | Coccolithophyceae | Isochrysidales | Noelaerhabdaceae |
| | ASV277 | 2.85 | 1.94E-07 | Eukaryota | Cercozoa | Protozoa classis incertae sedis | Protozoa ordo incertae sedis | Protozoa familia incertae sedis |
| | ASV2185 | 2.85 | 0.000644 | Eukaryota | Cryptophyta | Cryptophyceae | Kathablepharidacea | Katablepharidaceae |
| | ASV2660 | 2.84 | 8.12E-07 | Eukaryota | Telonemia | Telonemia classis ineditae | Telonemida | Telonemia familia ineditae |
| | ASV1234 | 2.83 | 7.20E-10 | Eukaryota | Ochrophyta | Bolidophyceae | Parmales | Triparmaceae |

| | | | | | | | | |
|---|---|---|---|---|---|---|---|---|
| | ASV1314 | 2.83 | 1.23E-06 | Eukaryota | Ochrophyta | Chrysophyceae | Synurales | Mallomonadaceae |
| | ASV543 | 2.81 | 2.02E-07 | Eukaryota | Ochrophyta | Chrysophyceae | Chromulinales | Paraphysomonadaceae |
| | ASV83 | 2.80 | 1.15E-07 | Eukaryota | Telonemia | Telonemia classis ineditae | Telonemida | Telonemia familia ineditae |
| | ASV24 | 2.80 | 0.000124 | Eukaryota | Myzozoa | Dinophyceae | Gymnodiniales | Gymnodiniaceae |
| | ASV1476 | 2.79 | 8.57E-08 | Eukaryota | Ochrophyta | Chrysophyceae | Chromulinales | Paraphysomonadaceae |
| | ASV63 | 2.79 | 0.000116 | Eukaryota | Bacillariophyta | Mediophyceae | Stephanodiscales | Stephanodiscaceae |
| | ASV2412 | 2.75 | 0.001807 | Eukaryota | Cryptophyta | Cryptophyceae | Pyrenomonadales | Geminigeraceae |
| | ASV591 | 2.74 | 0.001447 | Eukaryota | Cryptophyta | Cryptophyceae | Pyrenomonadales | Geminigeraceae |
| | ASV1644 | 2.73 | 0.001489 | Eukaryota | Cryptophyta | Cryptophyceae | Pyrenomonadales | Geminigeraceae |
| | ASV1677 | 2.73 | 1.80E-06 | Fungi | Chytridiomycota | Chytridiomycetes | Rhizophydiales | Uebelmesseromycetaceae |
| | ASV2253 | 2.69 | 0.001171 | Eukaryota | Bacillariophyta | Mediophyceae | Stephanodiscales | Stephanodiscaceae |
| | ASV1254 | 2.67 | 3.27E-06 | Eukaryota | Cryptophyta | Cryptophyceae | Pyrenomonadales | Geminigeraceae |
| | ASV2715 | 2.65 | 5.45E-06 | Eukaryota | Cryptophyta | Cryptophyceae | Pyrenomonadales | Geminigeraceae |
| | ASV2831 | 2.64 | 2.33E-06 | Eukaryota | Telonemia | Telonemia classis ineditae | Telonemida | Telonemia familia ineditae |
| | ASV181 | 2.63 | 8.66E-05 | Eukaryota | Myzozoa | Dinophyceae | Gymnodiniales | Gymnodiniaceae |
| | ASV1174 | 2.60 | 0.000113 | Eukaryota | Chlorophyta | Trebouxiophyceae | Chlorellales | Chlorellales incertae sedis |
| | ASV1545 | 2.59 | 0.00088 | Eukaryota | Cryptophyta | Cryptophyceae | Kathablepharidacea | Katablepharidaceae |
| | ASV354 | 2.58 | 0.000708 | Eukaryota | Ochrophyta | Chrysophyceae | Chromulinales | Paraphysomonadaceae |
| | ASV1642 | 2.57 | 0.00989 | Eukaryota | Choanozoa | Choanoflagellata | Craspedida | Codonosigaceae |
| | ASV517 | 2.56 | 4.10E-07 | Eukaryota | Cryptophyta | Cryptophyceae | Pyrenomonadales | Geminigeraceae |
| | ASV2741 | 2.56 | 0.001066 | Eukaryota | Haptophyta | Coccolithophyceae | Isochrysidales | Noelaerhabdaceae |
| | ASV2697 | 2.55 | 0.000105 | Eukaryota | Myzozoa | Dinophyceae | Gymnodiniales | Gymnodiniaceae |
| | ASV1219 | 2.55 | 0.003167 | Eukaryota | Ciliophora | Prostomatea | Prorodontida | Colepidae |
| | ASV1784 | 2.55 | 3.87E-05 | Eukaryota | Cryptophyta | Cryptophyceae | Pyrenomonadales | Geminigeraceae |
| | ASV814 | 2.55 | 2.76E-05 | Eukaryota | Cryptophyta | Cryptophyceae | Pyrenomonadales | Geminigeraceae |
| | ASV422 | 2.54 | 8.40E-06 | Eukaryota | Cryptophyta | Cryptophyceae | Pyrenomonadales | Geminigeraceae |
| | ASV129 | 2.53 | 0.00083 | Eukaryota | Cryptophyta | Cryptophyceae | Pyrenomonadales | Geminigeraceae |
| | ASV2653 | 2.52 | 0.001066 | Eukaryota | Cryptophyta | Cryptophyceae | Kathablepharidacea | Katablepharidaceae |

| | | | | | | | | |
|---|---|---|---|---|---|---|---|---|
| | ASV1287 | 2.52 | 9.26E-07 | Eukaryota | Haptophyta | Coccolithophyceae | Prymnesiales | Chrysochromulinaceae |
| | ASV177 | 2.50 | 0.009365 | Eukaryota | Choanozoa | Choanoflagellata | Craspedida | Codonosigaceae |
| | ASV69 | 2.48 | 0.003326 | Eukaryota | Ciliophora | Prostomatea | Prorodontida | Colepidae |
| | ASV273 | 2.46 | 2.32E-05 | Eukaryota | Chlorophyta | Chlorophyceae | Sphaeropleales | Mychonastaceae |
| | ASV2410 | 2.45 | 1.84E-05 | Eukaryota | Ochrophyta | Chrysophyceae | Synurales | Mallomonadaceae |
| | ASV1385 | 2.45 | 0.000602 | Eukaryota | Ochrophyta | Dictyochophyceae | Pedinellales | Actinomonadaceae |
| | ASV2604 | 2.45 | 5.10E-05 | Eukaryota | Cryptophyta | Cryptophyceae | Pyrenomonadales | Geminigeraceae |
| | ASV2476 | 2.43 | 0.001667 | Eukaryota | Ochrophyta | Chrysophyceae | Chromulinales | Paraphysomonadaceae |
| | ASV2768 | 2.42 | 0.000116 | Eukaryota | Bacillariophyta | Mediophyceae | Stephanodiscales | Stephanodiscaceae |
| | ASV1707 | 2.41 | 0.001794 | Eukaryota | Telonemia | Telonemia classis ineditae | Telonemida | Telonemia familia ineditae |
| | ASV2520 | 2.40 | 0.000186 | Eukaryota | Oomycota | Hyphochytrea | Pirsoniales | Pirsoniaceae |
| | ASV625 | 2.38 | 6.19E-06 | Eukaryota | Haptophyta | Coccolithophyceae | Prymnesiales | Chrysochromulinaceae |
| | ASV1378 | 2.38 | 0.001644 | Eukaryota | Ochrophyta | Dictyochophyceae | Pedinellales | Actinomonadaceae |
| | ASV2670 | 2.38 | 2.19E-05 | Eukaryota | Cercozoa | Protozoa classis incertae sedis | Protozoa ordo incertae sedis | Protozoa familia incertae sedis |
| | ASV789 | 2.37 | 7.95E-06 | Eukaryota | Haptophyta | Coccolithophyceae | Prymnesiales | Chrysochromulinaceae |
| | ASV1145 | 2.36 | 0.001462 | Eukaryota | Cryptophyta | Cryptophyceae | Pyrenomonadales | Geminigeraceae |
| | ASV1465 | 2.35 | 0.002368 | Eukaryota | Bigyra | incertae sedis | incertae sedis | Pseudophyllomitidae |
| | ASV1951 | 2.35 | 1.07E-05 | Eukaryota | Cercozoa | Protozoa classis incertae sedis | Protozoa ordo incertae sedis | Protozoa familia incertae sedis |
| | ASV364 | 2.34 | 0.000162 | Eukaryota | Oomycota | Hyphochytrea | Pirsoniales | Pirsoniaceae |
| | ASV2382 | 2.33 | 9.48E-05 | Eukaryota | Bacillariophyta | Mediophyceae | Stephanodiscales | Stephanodiscaceae |
| | ASV868 | 2.32 | 2.90E-06 | Eukaryota | Haptophyta | Coccolithophyceae | Prymnesiales | Chrysochromulinaceae |
| | ASV874 | 2.32 | 0.003559 | Eukaryota | Bacillariophyta | Mediophyceae | Stephanodiscales | Stephanodiscaceae |
| | ASV1113 | 2.29 | 0.000769 | Eukaryota | Cryptophyta | Cryptophyceae | Pyrenomonadales | Geminigeraceae |
| | ASV1327 | 2.27 | 1.17E-05 | Eukaryota | Cercozoa | Thecofilosea | Cryomonadida | Protaspidae |
| | ASV1786 | 2.27 | 4.67E-06 | Eukaryota | Choanozoa | Choanoflagellata | Acanthoecida | Acanthoecidae |
| | ASV400 | 2.26 | 4.10E-05 | Eukaryota | Cryptophyta | Cryptophyceae | Pyrenomonadales | Geminigeraceae |
| | ASV2564 | 2.24 | 1.21E-05 | Eukaryota | Cryptophyta | Cryptophyceae | Pyrenomonadales | Geminigeraceae |
| | ASV1991 | 2.21 | 5.95E-05 | Eukaryota | Ochrophyta | Chrysophyceae | Ochromonadales | Ochromonadaceae |

| | | | | | | | | |
|---|---|---|---|---|---|---|---|---|
| | ASV1530 | 2.20 | 0.007392 | Eukaryota | Bigyra | incertae sedis | incertae sedis | Pseudophyllomitidae |
| | ASV1595 | 2.19 | 1.17E-05 | Eukaryota | Myzozoa | Dinophyceae | Gymnodiniales | Gymnodiniaceae |
| | ASV380 | 2.18 | 3.91E-05 | Eukaryota | Bigyra | incertae sedis | incertae sedis | Pseudophyllomitidae |
| | ASV367 | 2.17 | 3.19E-05 | Eukaryota | Ochrophyta | Dictyochophyceae | Pedinellales | Actinomonadaceae |
| | ASV2685 | 2.16 | 9.51E-05 | Eukaryota | Chlorophyta | Chlorophyceae | Sphaeropleales | Mychonastaceae |
| | ASV1096 | 2.16 | 7.07E-07 | Eukaryota | Telonemia | Telonemia classis ineditae | Telonemida | Telonemia familia ineditae |
| | ASV2852 | 2.12 | 0.002709 | Eukaryota | Haptophyta | Coccolithophyceae | Isochrysidales | Noelaerhabdaceae |
| | ASV65 | 2.12 | 1.90E-05 | Eukaryota | Ochrophyta | Chrysophyceae | Synurales | Mallomonadaceae |
| | ASV265 | 2.11 | 0.000889 | Eukaryota | Bacillariophyta | Mediophyceae | Stephanodiscales | Stephanodiscaceae |
| | ASV119 | 2.10 | 0.000138 | Eukaryota | Bacillariophyta | Mediophyceae | Stephanodiscales | Stephanodiscaceae |
| | ASV518 | 2.09 | 0.006294 | Eukaryota | Bacillariophyta | Mediophyceae | Stephanodiscales | Stephanodiscaceae |
| | ASV2825 | 2.08 | 2.40E-05 | Eukaryota | Ochrophyta | Dictyochophyceae | Pedinellales | Actinomonadaceae |
| | ASV1275 | 2.07 | 0.000575 | Eukaryota | Chlorophyta | Mamiellophyceae | Dolichomastigales | Crustomastigaceae |
| | ASV321 | 2.07 | 9.06E-05 | Eukaryota | Bacillariophyta | Mediophyceae | Stephanodiscales | Stephanodiscaceae |
| | ASV536 | 2.07 | 0.003777 | Eukaryota | Ochrophyta | Chrysophyceae | Ochromonadales | Ochromonadaceae |
| | ASV666 | 2.06 | 0.005107 | Eukaryota | Cryptophyta | Cryptophyceae | Kathablepharidacea | Katablepharidaceae |
| | ASV2237 | 2.06 | 0.005293 | Eukaryota | Ochrophyta | Chrysophyceae | Ochromonadales | Ochromonadaceae |
| | ASV865 | 2.05 | 0.000511 | Eukaryota | Bacillariophyta | Mediophyceae | Stephanodiscales | Stephanodiscaceae |
| | ASV2761 | 2.04 | 0.0006 | Eukaryota | Chlorophyta | Chlorophyceae | Sphaeropleales | Mychonastaceae |
| | ASV993 | 2.04 | 1.44E-05 | Eukaryota | Ochrophyta | Dictyochophyceae | Pedinellales | Actinomonadaceae |
| | ASV2616 | 2.04 | 4.94E-05 | Fungi | Ascomycota | Leotiomycetes | Erysiphales | Erysiphaceae |
| | ASV844 | 2.02 | 0.00035 | Eukaryota | Cryptophyta | Cryptophyceae | Pyrenomonadales | Geminigeraceae |
| | ASV2762 | 2.02 | 0.000173 | Eukaryota | Ochrophyta | Chrysophyceae | Synurales | Mallomonadaceae |
| | ASV1079 | 2.01 | 0.00072 | Eukaryota | Ochrophyta | Chrysophyceae | Chromulinales | Paraphysomonadaceae |
| 16S-PP | ASV267 | 7.54 | 1.20E-18 | Bacteria | Bacteroidota | Kryptonia | Kryptoniales | BSV26 |
| | ASV203 | 7.15 | 4.55E-29 | Bacteria | Acidobacteriota | Acidobacteriae | Acidobacteriae | Acidobacteriae |
| | ASV95 | 6.86 | 1.06E-15 | Bacteria | Bacteroidota | Bacteroidia | Chitinophagales | Chitinophagaceae |
| | ASV171 | 6.65 | 2.59E-19 | Bacteria | Acidobacteriota | Vicinamibacteria | Vicinamibacterales | uncultured |

| | | | | | | | | |
|---|---|---|---|---|---|---|---|---|
| | ASV889 | 6.52 | 1.41E-35 | Bacteria | Planctomycetota | Phycisphaerae | Phycisphaerales | Phycisphaeraceae |
| | ASV431 | 6.39 | 4.47E-27 | Bacteria | Proteobacteria | Alphaproteobacteria | Rhizobiales | Xanthobacteraceae |
| | ASV16 | 6.37 | 7.51E-24 | Bacteria | Proteobacteria | Alphaproteobacteria | Rhizobiales | Xanthobacteraceae |
| | ASV468 | 6.26 | 1.96E-15 | Bacteria | Proteobacteria | Gammaproteobacteria | Burkholderiales | Nitrosomonadaceae |
| | ASV382 | 6.20 | 2.07E-26 | Bacteria | Acidobacteriota | Vicinamibacteria | Vicinamibacterales | Vicinamibacteraceae |
| | ASV813 | 6.07 | 6.89E-13 | Bacteria | Nitrospirota | Nitrospiria | Nitrospirales | Nitrospiraceae |
| | ASV509 | 5.94 | 6.10E-23 | Bacteria | Bdellovibrionota | Bdellovibrionia | Bdellovibrionales | Bdellovibrionaceae |
| | ASV695 | 5.93 | 3.56E-27 | Bacteria | Bacteroidota | Bacteroidia | Sphingobacteriales | NS11-12_marine_group |
| | ASV104 | 5.84 | 1.07E-26 | Bacteria | Proteobacteria | Alphaproteobacteria | Acetobacterales | Acetobacteraceae |
| | ASV276 | 5.78 | 1.17E-20 | Bacteria | Planctomycetota | Phycisphaerae | Phycisphaerales | Phycisphaeraceae |
| | ASV55 | 5.73 | 5.75E-12 | Bacteria | Nitrospirota | Nitrospiria | Nitrospirales | Nitrospiraceae |
| | ASV280 | 5.47 | 8.75E-21 | Bacteria | Acidobacteriota | Vicinamibacteria | Vicinamibacterales | Vicinamibacteraceae |
| | ASV780 | 5.46 | 7.42E-28 | Bacteria | Chloroflexi | JG30-KF-CM66 | JG30-KF-CM66 | JG30-KF-CM66 |
| | ASV971 | 5.43 | 9.42E-12 | Bacteria | Proteobacteria | Alphaproteobacteria | Reyranellales | Reyranellaceae |
| | ASV927 | 5.32 | 1.04E-13 | Bacteria | Gemmatimonadota | Gemmatimonadetes | Gemmatimonadales | Gemmatimonadaceae |
| | ASV90 | 5.19 | 7.78E-36 | Bacteria | Proteobacteria | Alphaproteobacteria | Rhizobiales | KF-JG30-B3 |
| | ASV507 | 5.13 | 4.22E-12 | Bacteria | Bacteroidota | Bacteroidia | Chitinophagales | Chitinophagaceae |
| | ASV389 | 5.11 | 5.91E-25 | Bacteria | Chloroflexi | JG30-KF-CM66 | JG30-KF-CM66 | JG30-KF-CM66 |
| | ASV928 | 5.00 | 5.36E-22 | Bacteria | Verrucomicrobiota | Verrucomicrobiae | Opitutales | Opitutaceae |
| | ASV916 | 4.98 | 5.46E-16 | Bacteria | Proteobacteria | Gammaproteobacteria | Oceanospirillales | Pseudohongiellaceae |
| | ASV451 | 4.87 | 6.12E-11 | Bacteria | Acidobacteriota | Vicinamibacteria | Vicinamibacterales | Vicinamibacteraceae |
| | ASV924 | 4.81 | 3.62E-11 | Bacteria | Chloroflexi | Anaerolineae | Anaerolineales | Anaerolineaceae |
| | ASV933 | 4.73 | 1.82E-18 | Bacteria | Bacteroidota | Bacteroidia | Sphingobacteriales | NS11-12_marine_group |
| | ASV275 | 4.72 | 1.17E-20 | Bacteria | Acidobacteriota | Vicinamibacteria | Vicinamibacterales | Vicinamibacteraceae |
| | ASV388 | 4.63 | 3.02E-20 | Bacteria | Proteobacteria | Alphaproteobacteria | Reyranellales | Reyranellaceae |
| | ASV640 | 4.62 | 6.03E-26 | Bacteria | Bacteroidota | Bacteroidia | Sphingobacteriales | NS11-12_marine_group |
| | ASV92 | 4.54 | 5.72E-13 | Bacteria | Proteobacteria | Alphaproteobacteria | Acetobacterales | Acetobacteraceae |
| | ASV198 | 4.53 | 4.32E-21 | Bacteria | Acidobacteriota | Acidobacteriae | Bryobacterales | Bryobacteraceae |

| | | | | | | | | |
|---|---|---|---|---|---|---|---|---|
| | ASV498 | 4.51 | 1.33E-09 | Bacteria | Bacteroidota | Bacteroidia | Sphingobacteriales | KD3-93 |
| | ASV972 | 4.47 | 1.17E-26 | Bacteria | Acidobacteriota | Vicinamibacteria | Vicinamibacterales | Vicinamibacteraceae |
| | ASV420 | 4.45 | 4.24E-21 | Bacteria | Acidobacteriota | Vicinamibacteria | Vicinamibacterales | uncultured |
| | ASV535 | 4.44 | 1.49E-22 | Bacteria | Actinobacteriota | Thermoleophilia | Gaiellales | uncultured |
| | ASV642 | 4.33 | 2.20E-10 | Bacteria | Proteobacteria | Gammaproteobacteria | Burkholderiales | Nitrosomonadaceae |
| | ASV965 | 4.29 | 1.11E-15 | Bacteria | Actinobacteriota | Acidimicrobiia | Microtrichales | Iamiaceae |
| | ASV1004 | 4.29 | 2.51E-22 | Bacteria | Bacteroidota | Bacteroidia | Chitinophagales | Chitinophagaceae |
| | ASV490 | 4.28 | 4.18E-26 | Bacteria | Chloroflexi | JG30-KF-CM66 | JG30-KF-CM66 | JG30-KF-CM66 |
| | ASV597 | 4.26 | 7.93E-11 | Bacteria | Verrucomicrobiota | Verrucomicrobiae | Pedosphaerales | Pedosphaeraceae |
| | ASV227 | 4.22 | 2.80E-08 | Bacteria | Proteobacteria | Gammaproteobacteria | Burkholderiales | betIII |
| | ASV638 | 4.17 | 3.94E-21 | Bacteria | Verrucomicrobiota | Verrucomicrobiae | Pedosphaerales | Pedosphaeraceae |
| | ASV832 | 4.15 | 1.58E-13 | Bacteria | Proteobacteria | Alphaproteobacteria | SAR11_clade | Clade_I |
| | ASV199 | 4.04 | 1.76E-10 | Bacteria | Bacteroidota | Bacteroidia | Sphingobacteriales | NS11-12_marine_group |
| | ASV804 | 4.03 | 6.10E-23 | Bacteria | Proteobacteria | Alphaproteobacteria | Rhodospirillales | Rhodospirillaceae |
| | ASV605 | 4.00 | 6.25E-12 | Bacteria | Bacteroidota | Bacteroidia | Sphingobacteriales | Sphingobacteriaceae |
| | ASV840 | 3.98 | 1.62E-23 | Bacteria | Gemmatimonadota | BD2-11_terrestrial_group | BD2-11_terrestrial_group | BD2-11_terrestrial_group |
| | ASV394 | 3.95 | 2.14E-12 | Bacteria | Gemmatimonadota | Gemmatimonadetes | Gemmatimonadales | Gemmatimonadaceae |
| | ASV671 | 3.91 | 2.24E-18 | Bacteria | Proteobacteria | Alphaproteobacteria | Rhodospirillales | Rhodospirillaceae |
| | ASV511 | 3.85 | 6.54E-17 | Bacteria | Chloroflexi | P2-11E | P2-11E | P2-11E |
| | ASV401 | 3.81 | 3.80E-09 | Bacteria | Proteobacteria | Alphaproteobacteria | Rhizobiales | Beijerinckiaceae |
| | ASV85 | 3.80 | 6.45E-08 | Bacteria | Actinobacteriota | Actinobacteria | PeM15 | PeM15 |
| | ASV417 | 3.77 | 7.87E-16 | Bacteria | Acidobacteriota | Vicinamibacteria | Vicinamibacterales | Vicinamibacteraceae |
| | ASV542 | 3.73 | 5.68E-14 | Bacteria | Proteobacteria | Alphaproteobacteria | Ferrovibrionales | Ferrovibrionales |
| | ASV213 | 3.73 | 2.40E-12 | Bacteria | Chloroflexi | Anaerolineae | Anaerolineales | Anaerolineaceae |
| | ASV876 | 3.70 | 5.18E-13 | Bacteria | Proteobacteria | Gammaproteobacteria | CCM19a | CCM19a |
| | ASV849 | 3.67 | 9.81E-15 | Bacteria | Proteobacteria | Gammaproteobacteria | Burkholderiales | TRA3-20 |
| | ASV205 | 3.62 | 2.08E-05 | Bacteria | Bacteroidota | Bacteroidia | Flavobacteriales | Crocinitomicaceae |
| | ASV1031 | 3.62 | 3.71E-08 | Bacteria | Proteobacteria | Gammaproteobacteria | Burkholderiales | Nitrosomonadaceae |

| | | | | | | | | |
|---|---|---|---|---|---|---|---|---|
| | ASV449 | 3.58 | 1.40E-09 | Bacteria | Acidobacteriota | Vicinamibacteria | Vicinamibacterales | Vicinamibacteraceae |
| | ASV581 | 3.58 | 4.63E-08 | Bacteria | Proteobacteria | Alphaproteobacteria | Rhodobacterales | Rhodobacteraceae |
| | ASV589 | 3.57 | 1.27E-06 | Bacteria | SAR324_clade(Marine_group_B) | SAR324_clade(Marine_group_B) | SAR324_clade(Marine_group_B) | SAR324_clade(Marine_group_B) |
| | ASV152 | 3.56 | 2.38E-20 | Bacteria | Proteobacteria | Alphaproteobacteria | Rhodospirillales | Rhodospirillaceae |
| | ASV907 | 3.55 | 5.89E-10 | Bacteria | Proteobacteria | Gammaproteobacteria | Burkholderiales | TRA3-20 |
| | ASV230 | 3.45 | 2.38E-07 | Bacteria | Proteobacteria | Gammaproteobacteria | Methylococcales | Methylomonadaceae |
| | ASV687 | 3.39 | 3.20E-06 | Bacteria | Actinobacteriota | Thermoleophilia | Gaiellales | uncultured |
| | ASV925 | 3.39 | 1.47E-09 | Bacteria | Proteobacteria | Alphaproteobacteria | Acetobacterales | Acetobacteraceae |
| | ASV437 | 3.36 | 3.92E-07 | Bacteria | Proteobacteria | Gammaproteobacteria | Burkholderiales | TRA3-20 |
| | ASV60 | 3.35 | 1.03E-15 | Bacteria | AncK6 | AncK6 | AncK6 | AncK6 |
| | ASV83 | 3.34 | 6.94E-22 | Bacteria | Proteobacteria | Alphaproteobacteria | uncultured | uncultured |
| | ASV184 | 3.34 | 2.02E-17 | Bacteria | Acidobacteriota | Vicinamibacteria | Vicinamibacterales | uncultured |
| | ASV293 | 3.34 | 5.46E-16 | Bacteria | Verrucomicrobiota | Verrucomicrobiae | Pedosphaerales | Pedosphaeraceae |
| | ASV464 | 3.32 | 4.23E-17 | Bacteria | Planctomycetota | OM190 | OM190 | OM190 |
| | ASV419 | 3.32 | 2.57E-10 | Bacteria | Proteobacteria | Alphaproteobacteria | Rhodobacterales | Rhodobacteraceae |
| | ASV854 | 3.31 | 5.05E-16 | Bacteria | Actinobacteriota | Acidimicrobiia | Microtrichales | Ilumatobacteraceae |
| | ASV327 | 3.28 | 6.10E-12 | Bacteria | Planctomycetota | Phycisphaerae | Phycisphaerales | Phycisphaeraceae |
| | ASV471 | 3.28 | 1.97E-16 | Bacteria | Acidobacteriota | Vicinamibacteria | Vicinamibacterales | Vicinamibacteraceae |
| | ASV815 | 3.24 | 9.14E-07 | Bacteria | Bacteroidota | Bacteroidia | Sphingobacteriales | NS11-12_marine_group |
| | ASV1036 | 3.24 | 4.23E-17 | Bacteria | Acidobacteriota | Vicinamibacteria | Vicinamibacterales | Vicinamibacteraceae |
| | ASV483 | 3.23 | 2.53E-10 | Bacteria | Proteobacteria | Gammaproteobacteria | Legionellales | Legionellaceae |
| | ASV462 | 3.22 | 1.17E-12 | Bacteria | Acidobacteriota | Acidobacteriae | Bryobacterales | Bryobacteraceae |
| | ASV760 | 3.21 | 5.34E-09 | Bacteria | Proteobacteria | Gammaproteobacteria | Burkholderiales | Oxalobacteraceae |
| | ASV711 | 3.18 | 3.32E-18 | Bacteria | Proteobacteria | Gammaproteobacteria | Burkholderiales | Comamonadaceae |
| | ASV87 | 3.17 | 1.01E-06 | Bacteria | Bacteroidota | Bacteroidia | Flavobacteriales | NS9_marine_group |
| | ASV76 | 3.14 | 9.04E-08 | Bacteria | Proteobacteria | Gammaproteobacteria | Oceanospirillales | Pseudohongiellaceae |
| | ASV297 | 3.11 | 1.90E-14 | Bacteria | Bdellovibrionota | Oligoflexia | 0319-6G20 | 0319-6G20 |
| | ASV473 | 3.11 | 1.26E-14 | Bacteria | Verrucomicrobiota | Verrucomicrobiae | Pedosphaerales | Pedosphaeraceae |

| | | | | | | | | |
|---|---|---|---|---|---|---|---|---|
| | ASV966 | 3.10 | 6.49E-10 | Bacteria | Bdellovibrionota | Oligoflexia | Oligoflexales | uncultured |
| | ASV225 | 3.08 | 1.07E-06 | Bacteria | Proteobacteria | Gammaproteobacteria | Gammaproteobacteria_Incertae_Sedis | Unknown_Family |
| | ASV514 | 3.06 | 3.38E-14 | Bacteria | Proteobacteria | Alphaproteobacteria | Zavarziniales | uncultured |
| | ASV302 | 3.03 | 1.21E-07 | Bacteria | Proteobacteria | Gammaproteobacteria | Methylococcales | Methylomonadaceae |
| | ASV208 | 3.02 | 1.50E-08 | Bacteria | Verrucomicrobiota | Verrucomicrobiae | Chthoniobacterales | Chthoniobacteraceae |
| | ASV365 | 3.02 | 1.13E-12 | Bacteria | Proteobacteria | Alphaproteobacteria | Acetobacterales | Acetobacteraceae |
| | ASV317 | 3.01 | 4.42E-06 | Bacteria | Planctomycetota | Phycisphaerae | Phycisphaerales | Phycisphaeraceae |
| | ASV864 | 3.01 | 1.06E-15 | Bacteria | Proteobacteria | Alphaproteobacteria | uncultured | uncultured |
| | ASV143 | 3.00 | 2.20E-13 | Bacteria | Bdellovibrionota | Bdellovibrionia | Bdellovibrionales | Bdellovibrionaceae |
| | ASV718 | 3.00 | 4.14E-16 | Bacteria | Chloroflexi | JG30-KF-CM66 | JG30-KF-CM66 | JG30-KF-CM66 |
| | ASV107 | 3.00 | 0.001317 | Bacteria | Proteobacteria | Gammaproteobacteria | Burkholderiales | betI |
| | ASV1037 | 2.99 | 5.85E-07 | Bacteria | Proteobacteria | Alphaproteobacteria | Micropepsales | Micropepsaceae |
| | ASV1010 | 2.98 | 4.83E-12 | Bacteria | Proteobacteria | Alphaproteobacteria | Rhodospirillales | uncultured |
| | ASV970 | 2.97 | 7.36E-10 | Bacteria | Planctomycetota | Planctomycetes | Gemmatales | Gemmataceae |
| | ASV736 | 2.94 | 9.64E-15 | Bacteria | Chloroflexi | JG30-KF-CM66 | JG30-KF-CM66 | JG30-KF-CM66 |
| | ASV826 | 2.91 | 1.18E-06 | Bacteria | Proteobacteria | Gammaproteobacteria | Burkholderiales | TRA3-20 |
| | ASV369 | 2.91 | 0.006793 | Bacteria | Bacteroidota | Bacteroidia | Sphingobacteriales | env.OPS_17 |
| | ASV962 | 2.87 | 2.87E-14 | Bacteria | Acidobacteriota | Vicinamibacteria | Vicinamibacterales | Vicinamibacteraceae |
| | ASV773 | 2.87 | 9.34E-14 | Bacteria | Proteobacteria | Alphaproteobacteria | Rhizobiales | Xanthobacteraceae |
| | ASV758 | 2.87 | 9.43E-08 | Bacteria | Proteobacteria | Gammaproteobacteria | Burkholderiales | betIV |
| | ASV785 | 2.84 | 1.92E-09 | Bacteria | Proteobacteria | Gammaproteobacteria | Burkholderiales | TRA3-20 |
| | ASV14 | 2.82 | 7.58E-07 | Bacteria | Patescibacteria | Saccharimonadia | Saccharimonadales | Saccharimonadales |
| | ASV512 | 2.79 | 7.08E-16 | Bacteria | Acidobacteriota | Vicinamibacteria | Vicinamibacterales | Vicinamibacteraceae |
| | ASV98 | 2.79 | 4.76E-06 | Bacteria | Proteobacteria | Gammaproteobacteria | Burkholderiales | betIV |
| | ASV395 | 2.78 | 9.64E-15 | Bacteria | Proteobacteria | Alphaproteobacteria | Sphingomonadales | Sphingomonadaceae |
| | ASV258 | 2.76 | 3.94E-14 | Bacteria | Verrucomicrobiota | Verrucomicrobiae | Pedosphaerales | Pedosphaeraceae |
| | ASV502 | 2.75 | 2.52E-06 | Bacteria | Proteobacteria | Gammaproteobacteria | Legionellales | Legionellaceae |
| | ASV151 | 2.73 | 1.83E-10 | Archaea | Crenarchaeota | Nitrososphaeria | Nitrosopumilales | Nitrosopumilaceae |

| | | | | | | | | |
|---|---|---|---|---|---|---|---|---|
| | ASV572 | 2.70 | 2.74E-07 | Bacteria | Proteobacteria | Alphaproteobacteria | SAR11_clade | Clade_II |
| | ASV286 | 2.70 | 3.54E-13 | Bacteria | Desulfobacterota | uncultured | uncultured | uncultured |
| | ASV459 | 2.69 | 5.89E-10 | Bacteria | Myxococcota | Polyangia | Blfdi19 | Blfdi19 |
| | ASV567 | 2.67 | 0.000236 | Bacteria | Proteobacteria | Alphaproteobacteria | Paracaedibacterales | Paracaedibacteraceae |
| | ASV477 | 2.62 | 2.99E-12 | Bacteria | Acidobacteriota | Vicinamibacteria | Vicinamibacterales | Vicinamibacteraceae |
| | ASV882 | 2.60 | 9.49E-15 | Bacteria | Proteobacteria | Alphaproteobacteria | uncultured | uncultured |
| | ASV109 | 2.57 | 1.22E-11 | Bacteria | Acidobacteriota | Vicinamibacteria | Vicinamibacterales | uncultured |
| | ASV602 | 2.54 | 7.69E-08 | Bacteria | Planctomycetota | Phycisphaerae | Phycisphaerales | Phycisphaeraceae |
| | ASV1034 | 2.54 | 4.95E-11 | Bacteria | Chloroflexi | JG30-KF-CM66 | JG30-KF-CM66 | JG30-KF-CM66 |
| | ASV1028 | 2.52 | 2.11E-07 | Bacteria | Proteobacteria | Alphaproteobacteria | Acetobacterales | Acetobacteraceae |
| | ASV870 | 2.51 | 1.35E-07 | Bacteria | Bacteroidota | Bacteroidia | Sphingobacteriales | KD3-93 |
| | ASV166 | 2.51 | 7.36E-10 | Bacteria | Actinobacteriota | Thermoleophilia | Solirubrobacterales | Solirubrobacteraceae |
| | ASV339 | 2.51 | 1.53E-14 | Bacteria | Acidobacteriota | Vicinamibacteria | Vicinamibacterales | Vicinamibacteraceae |
| | ASV646 | 2.50 | 2.41E-11 | Bacteria | Proteobacteria | Alphaproteobacteria | Rhodospirillales | Rhodospirillaceae |
| | ASV1007 | 2.49 | 0.000285 | Bacteria | Proteobacteria | Alphaproteobacteria | Reyranellales | Reyranellaceae |
| | ASV1005 | 2.48 | 3.71E-09 | Bacteria | Actinobacteriota | Acidimicrobiia | Microtrichales | Ilumatobacteraceae |
| | ASV396 | 2.47 | 8.12E-09 | Bacteria | Proteobacteria | Alphaproteobacteria | Rhizobiales | Hyphomicrobiaceae |
| | ASV442 | 2.46 | 1.24E-11 | Bacteria | Acidobacteriota | Vicinamibacteria | Vicinamibacterales | Vicinamibacteraceae |
| | ASV392 | 2.44 | 4.10E-11 | Bacteria | Proteobacteria | Alphaproteobacteria | Puniceispirillales | EF100-94H03 |
| | ASV592 | 2.44 | 0.000206 | Bacteria | Proteobacteria | Alphaproteobacteria | Caulobacterales | alfII |
| | ASV99 | 2.42 | 8.30E-13 | Bacteria | Actinobacteriota | Thermoleophilia | Gaiellales | Gaiellaceae |
| | ASV906 | 2.40 | 1.83E-10 | Bacteria | Acidobacteriota | Vicinamibacteria | Vicinamibacterales | Vicinamibacteraceae |
| | ASV1041 | 2.38 | 8.99E-11 | Bacteria | Gemmatimonadota | Gemmatimonadetes | Gemmatimonadales | Gemmatimonadaceae |
| | ASV775 | 2.36 | 0.000398 | Bacteria | Verrucomicrobiota | Verrucomicrobiae | Opitutales | Opitutaceae |
| | ASV142 | 2.35 | 5.08E-05 | Bacteria | Planctomycetota | Phycisphaerae | Phycisphaerales | Phycisphaeraceae |
| | ASV333 | 2.34 | 3.64E-11 | Bacteria | Proteobacteria | Alphaproteobacteria | Rhizobiales | Xanthobacteraceae |
| | ASV606 | 2.32 | 1.08E-12 | Bacteria | Proteobacteria | Alphaproteobacteria | Puniceispirillales | uncultured |
| | ASV655 | 2.31 | 8.24E-08 | Bacteria | Proteobacteria | Alphaproteobacteria | Rhizobiales | Hyphomicrobiaceae |

| | | | | | | | | |
|---|---|---|---|---|---|---|---|---|
| | ASV693 | 2.30 | 4.86E-05 | Bacteria | Actinobacteriota | Acidimicrobiia | IMCC26256 | IMCC26256 |
| | ASV220 | 2.26 | 3.01E-08 | Bacteria | Actinobacteriota | Actinobacteria | Frankiales | Sporichthyaceae |
| | ASV686 | 2.24 | 1.91E-06 | Bacteria | Proteobacteria | Gammaproteobacteria | KI89A_clade | KI89A_clade |
| | ASV545 | 2.19 | 2.33E-08 | Bacteria | Myxococcota | Polyangia | Blfdi19 | Blfdi19 |
| | ASV958 | 2.17 | 0.000793 | Bacteria | Bacteroidota | Kapabacteria | Kapabacteriales | Kapabacteriales |
| | ASV10 | 2.16 | 4.25E-11 | Bacteria | Chloroflexi | KD4-96 | KD4-96 | KD4-96 |
| | ASV134 | 2.10 | 7.72E-08 | Bacteria | Proteobacteria | Alphaproteobacteria | Rhizobiales | alfI |
| | ASV197 | 2.10 | 0.000607 | Bacteria | Bacteroidota | Bacteroidia | Flavobacteriales | Cryomorphaceae |
| | ASV841 | 2.10 | 1.50E-05 | Bacteria | Dependentiae | Babeliae | Babeliales | UBA12409 |
| | ASV636 | 2.07 | 1.68E-07 | Bacteria | Verrucomicrobiota | Verrucomicrobiae | Pedosphaerales | Pedosphaeraceae |
| | ASV771 | 2.06 | 2.46E-10 | Bacteria | Acidobacteriota | Acidobacteriae | Bryobacterales | Bryobacteraceae |
| | ASV728 | 2.05 | 1.14E-09 | Bacteria | Margulisbacteria | Margulisbacteria | Margulisbacteria | Margulisbacteria |
| | ASV515 | 2.05 | 0.001537 | Bacteria | Proteobacteria | Alphaproteobacteria | Rickettsiales | Rickettsiaceae |
| | ASV283 | 2.05 | 3.72E-09 | Bacteria | Myxococcota | bacteriap25 | bacteriap25 | bacteriap25 |
| | ASV618 | 2.04 | 5.73E-09 | Bacteria | Proteobacteria | Gammaproteobacteria | Burkholderiales | Nitrosomonadaceae |
| | ASV491 | 2.01 | 0.000578 | Bacteria | Planctomycetota | Phycisphaerae | Phycisphaerales | Phycisphaeraceae |

**Table S4: Linear Models for Differential Abundance (LinDA) outputs for the composition of microbial plankton composition differences between the oxic and anoxic water column in LLC.** The outputs consist of ASVs that pass the threshold of Log2FC above 2 and an adjusted p-value below 0.01, along with their taxonomic affiliation up to family level. The ASVs are categorised as follows: eukaryotic nanoplankton (18S-NP), eukaryotic picoplankton (18S-PP) and prokaryotic picoplankton (16S-PP).

| Microbial plankton | ASV | log2FC | adjusted p-value | Kingdoms | Phylum | Class | Order | Family |
|---|---|---|---|---|---|---|---|---|
| 18S-NP | ASV711 | 7.06 | 9.68E-23 | Eukaryota | Myzozoa | Dinophyceae | Peridiniales | Protoperidiniaceae |
| | ASV1217 | 6.87 | 3.36E-21 | Eukaryota | Myzozoa | Dinophyceae | Peridiniales | Protoperidiniaceae |
| | ASV2302 | 6.80 | 4.33E-23 | Eukaryota | Myzozoa | Dinophyceae | Peridiniales | Protoperidiniaceae |
| | ASV184 | 6.53 | 2.26E-22 | Eukaryota | Myzozoa | Dinophyceae | Peridiniales | Protoperidiniaceae |
| | ASV1010 | 6.18 | 5.93E-14 | Eukaryota | Ciliophora | Spirotrichea | Urostylida | Holostichidae |
| | ASV1615 | 6.15 | 1.57E-13 | Eukaryota | Ciliophora | Spirotrichea | Urostylida | Holostichidae |
| | ASV257 | 5.96 | 2.86E-10 | Eukaryota | Ciliophora | Litostomatea | Haptorida | Dimacrocaryonidae |
| | ASV1157 | 5.91 | 3.32E-14 | Eukaryota | Ciliophora | Spirotrichea | Urostylida | Holostichidae |
| | ASV2577 | 5.85 | 5.87E-11 | Eukaryota | Ciliophora | Litostomatea | Haptorida | Dimacrocaryonidae |
| | ASV2220 | 5.83 | 5.17E-10 | Eukaryota | Ciliophora | Litostomatea | Haptorida | Dimacrocaryonidae |
| | ASV1025 | 5.72 | 5.58E-11 | Eukaryota | Ciliophora | Litostomatea | Haptorida | Dimacrocaryonidae |
| | ASV389 | 5.68 | 2.32E-19 | Eukaryota | Myzozoa | Dinophyceae | Peridiniales | Diplopsaliaceae |
| | ASV667 | 5.61 | 2.21E-10 | Eukaryota | Ciliophora | Prostomatea | Prorodontida | Colepidae |
| | ASV1076 | 5.51 | 8.06E-15 | Eukaryota | Cercozoa | Protozoa classis incertae sedis | Protozoa ordo incertae sedis | Protozoa familia incertae sedis |
| | ASV2202 | 5.49 | 4.70E-17 | Eukaryota | Myzozoa | Dinophyceae | Peridiniales | Diplopsaliaceae |
| | ASV167 | 5.41 | 2.64E-12 | Eukaryota | Ciliophora | Spirotrichea | Urostylida | Holostichidae |
| | ASV1725 | 5.26 | 4.09E-10 | Eukaryota | Ciliophora | Prostomatea | Prorodontida | Colepidae |
| | ASV1005 | 5.25 | 6.34E-09 | Eukaryota | Ciliophora | Prostomatea | Prorodontida | Colepidae |
| | ASV638 | 5.20 | 5.87E-13 | Eukaryota | Cryptophyta | Cryptophyceae | Cryptomonadales | Cryptomonadaceae |
| | ASV2268 | 5.17 | 4.59E-19 | Eukaryota | Cryptophyta | Cryptophyceae | Cryptomonadales | Cryptomonadaceae |
| | ASV2153 | 5.16 | 4.44E-08 | Eukaryota | Bigyra | Bikosea | Bicosoecida | Bicosoecidae |
| | ASV1797 | 5.15 | 3.34E-26 | Eukaryota | Myzozoa | Dinophyceae | Prorocentrales | Prorocentraceae |
| | ASV842 | 5.13 | 6.21E-11 | Eukaryota | Myzozoa | Dinophyceae | Suessiales | Suessiaceae |

| | | | | | | | | |
|---|---|---|---|---|---|---|---|---|
| | ASV2002 | 5.12 | 3.00E-23 | Eukaryota | Myzozoa | Dinophyceae | Prorocentrales | Prorocentraceae |
| | ASV2827 | 5.05 | 2.55E-15 | Eukaryota | Myzozoa | Dinophyceae | Peridiniales | Diplopsaliaceae |
| | ASV2487 | 5.04 | 5.15E-07 | Eukaryota | Bigyra | Bikosea | Bicosoecida | Bicosoecidae |
| | ASV2591 | 5.04 | 1.32E-08 | Eukaryota | Ciliophora | Prostomatea | Prorodontida | Colepidae |
| | ASV748 | 4.99 | 4.03E-06 | Eukaryota | Ciliophora | Litostomatea | Haptorida | Spathidiidae |
| | ASV2650 | 4.98 | 1.63E-12 | Eukaryota | Cryptophyta | Cryptophyceae | Cryptomonadales | Cryptomonadaceae |
| | ASV1209 | 4.97 | 2.01E-08 | Eukaryota | Ciliophora | Spirotrichea | Oligotrichia | Strombidiidae |
| | ASV1077 | 4.96 | 2.12E-08 | Eukaryota | Ciliophora | Spirotrichea | Oligotrichia | Strombidiidae |
| | ASV2138 | 4.92 | 3.91E-06 | Eukaryota | Ciliophora | Litostomatea | Haptorida | Spathidiidae |
| | ASV2565 | 4.87 | 1.45E-06 | Eukaryota | Ciliophora | Litostomatea | Haptorida | Spathidiidae |
| | ASV1495 | 4.84 | 1.56E-05 | Eukaryota | Ciliophora | Spirotrichea | Tintinnida | Tintinnidae |
| | ASV1391 | 4.83 | 7.97E-28 | Eukaryota | Myzozoa | Dinophyceae | Prorocentrales | Prorocentraceae |
| | ASV1531 | 4.82 | 0.001188 | Eukaryota | Ochrophyta | Chrysophyceae | Synurales | Mallomonadaceae |
| | ASV171 | 4.81 | 9.98E-11 | Eukaryota | Myzozoa | Dinophyceae | Suessiales | Suessiaceae |
| | ASV790 | 4.81 | 3.35E-08 | Eukaryota | Bigyra | Bikosea | Bicosoecida | Bicosoecidae |
| | ASV1231 | 4.78 | 7.53E-09 | Eukaryota | Ciliophora | Spirotrichea | Oligotrichia | Strombidiidae |
| | ASV2359 | 4.77 | 2.40E-15 | Eukaryota | Myzozoa | Dinophyceae | Peridiniales | Diplopsaliaceae |
| | ASV198 | 4.71 | 5.61E-09 | Eukaryota | Chlorophyta | Chlorophyceae | Sphaeropleales | Sphaeropleaceae |
| | ASV2200 | 4.69 | 6.96E-09 | Eukaryota | Myzozoa | Dinophyceae | Suessiales | Suessiaceae |
| | ASV41 | 4.67 | 1.51E-07 | Eukaryota | Bigyra | Bikosea | Bicosoecida | Bicosoecidae |
| | ASV1706 | 4.66 | 2.81E-11 | Eukaryota | Cryptophyta | Cryptophyceae | Cryptomonadales | Cryptomonadaceae |
| | ASV4 | 4.65 | 8.16E-07 | Eukaryota | Myzozoa | Dinophyceae | Gonyaulacales | Ceratiaceae |
| | ASV554 | 4.63 | 2.07E-08 | Eukaryota | Myzozoa | Dinophyceae | Suessiales | Suessiaceae |
| | ASV641 | 4.61 | 5.48E-05 | Eukaryota | Ciliophora | Spirotrichea | Tintinnida | Tintinnidae |
| | ASV2176 | 4.60 | 2.23E-08 | Eukaryota | Bacillariophyta | Bacillariophyceae | Fragilariales | Fragilariaceae |
| | ASV1469 | 4.59 | 4.20E-09 | Eukaryota | Cryptophyta | Cryptophyceae | Cryptomonadales | Cryptomonadaceae |
| | ASV2593 | 4.54 | 3.64E-09 | Eukaryota | Chlorophyta | Chlorophyceae | Sphaeropleales | Sphaeropleaceae |
| | ASV2208 | 4.53 | 6.34E-09 | Eukaryota | Chlorophyta | Chlorophyceae | Sphaeropleales | Sphaeropleaceae |
| | ASV2474 | 4.50 | 1.44E-16 | Eukaryota | Myzozoa | Dinophyceae | Prorocentrales | Prorocentraceae |
| | ASV1307 | 4.49 | 1.62E-07 | Eukaryota | Bacillariophyta | Bacillariophyceae | Fragilariales | Fragilariaceae |

| | | | | | | | | |
|---|---|---|---|---|---|---|---|---|
| | ASV2105 | 4.49 | 4.71E-06 | Eukaryota | Ciliophora | Litostomatea | Haptorida | Spathidiidae |
| | ASV1251 | 4.48 | 2.23E-11 | Eukaryota | Myzozoa | Dinophyceae | Suessiales | Suessiaceae |
| | ASV2095 | 4.45 | 2.14E-05 | Eukaryota | Ciliophora | Spirotrichea | Tintinnida | Tintinnidae |
| | ASV2671 | 4.45 | 5.87E-11 | Eukaryota | Cryptophyta | Cryptophyceae | Cryptomonadales | Cryptomonadaceae |
| | ASV1278 | 4.44 | 2.11E-06 | Eukaryota | Myzozoa | Dinophyceae | Gonyaulacales | Ceratiaceae |
| | ASV2545 | 4.41 | 2.51E-11 | Eukaryota | Ochrophyta | Chrysophyceae | Synurales | Mallomonadaceae |
| | ASV160 | 4.41 | 3.09E-10 | Eukaryota | Cercozoa | Imbricatea | Thaumatomonadida | Thaumatomastigidae |
| | ASV2026 | 4.37 | 2.58E-10 | Eukaryota | Cryptophyta | Cryptophyceae | Cryptomonadales | Cryptomonadaceae |
| | ASV462 | 4.35 | 0.002397 | Eukaryota | Ochrophyta | Chrysophyceae | Synurales | Mallomonadaceae |
| | ASV2800 | 4.31 | 7.22E-06 | Eukaryota | Myzozoa | Dinophyceae | Gonyaulacales | Ceratiaceae |
| | ASV2578 | 4.29 | 2.75E-11 | Eukaryota | Myzozoa | Dinophyceae | Suessiales | Suessiaceae |
| | ASV2662 | 4.28 | 1.50E-07 | Eukaryota | Ciliophora | Spirotrichea | Oligotrichia | Strombidiidae |
| | ASV787 | 4.27 | 5.19E-12 | Eukaryota | Cryptophyta | Cryptophyceae | Cryptomonadales | Cryptomonadaceae |
| | ASV439 | 4.27 | 3.87E-09 | Eukaryota | Ciliophora | Prostomatea | Prorodontida | Colepidae |
| | ASV651 | 4.25 | 1.29E-10 | Eukaryota | Myzozoa | Dinophyceae | Suessiales | Suessiaceae |
| | ASV89 | 4.22 | 5.97E-06 | Eukaryota | Myzozoa | Dinophyceae | Gonyaulacales | Ceratiaceae |
| | ASV1966 | 4.17 | 0.000124 | Eukaryota | Ciliophora | Spirotrichea | Tintinnida | Tintinnidae |
| | ASV2409 | 4.16 | 0.003994 | Eukaryota | Ochrophyta | Chrysophyceae | Synurales | Mallomonadaceae |
| | ASV702 | 4.15 | 4.91E-10 | Eukaryota | Myzozoa | Dinophyceae | Suessiales | Suessiaceae |
| | ASV2255 | 4.08 | 4.98E-08 | Eukaryota | Chlorophyta | Chlorophyceae | Sphaeropleales | Sphaeropleaceae |
| | ASV1850 | 4.06 | 9.99E-11 | Eukaryota | Ochrophyta | Chrysophyceae | Synurales | Mallomonadaceae |
| | ASV986 | 4.03 | 1.02E-12 | Eukaryota | Ciliophora | Heterotrichea | Heterotrichida | Spirostomidae |
| | ASV2149 | 4.03 | 1.40E-10 | Eukaryota | Myzozoa | Dinophyceae | Suessiales | Suessiaceae |
| | ASV624 | 4.03 | 2.17E-13 | Eukaryota | Ciliophora | Oligohymenophorea | Pleuronematida | Conchophthiridae |
| | ASV2718 | 4.00 | 2.55E-08 | Eukaryota | Myzozoa | Dinophyceae | Suessiales | Suessiaceae |
| | ASV2758 | 3.99 | 2.81E-12 | Eukaryota | Cercozoa | Thecofilosea | Cryomonadida | Protaspidae |
| | ASV141 | 3.99 | 1.21E-09 | Eukaryota | Ciliophora | Prostomatea | Prorodontida | Colepidae |
| | ASV2344 | 3.98 | 1.84E-07 | Eukaryota | Cercozoa | Thecofilosea | Cryomonadida | Protaspidae |
| | ASV523 | 3.95 | 2.58E-10 | Eukaryota | Myzozoa | Dinophyceae | Suessiales | Suessiaceae |
| | ASV1948 | 3.90 | 4.29E-08 | Eukaryota | Chlorophyta | Chlorophyceae | Chlamydomonadales | Chlamydomonadaceae |

| | | | | | | | | |
|---|---|---|---|---|---|---|---|---|
| | ASV848 | 3.87 | 7.22E-07 | Eukaryota | Chlorophyta | Chlorophyceae | Chlamydomonadales | Chlamydomonadaceae |
| | ASV2783 | 3.86 | 1.31E-11 | Eukaryota | Ciliophora | Heterotrichea | Heterotrichida | Spirostomidae |
| | ASV480 | 3.85 | 1.50E-09 | Eukaryota | Cercozoa | Imbricatea | Thaumatomonadida | Thaumatomastigidae |
| | ASV2684 | 3.84 | 7.84E-09 | Eukaryota | Myzozoa | Dinophyceae | Suessiales | Suessiaceae |
| | ASV2245 | 3.83 | 7.72E-11 | Eukaryota | Ciliophora | Oligohymenophorea | Hymenostomatida | Ichthyophthiriidae |
| | ASV1491 | 3.83 | 6.12E-08 | Eukaryota | Myzozoa | Dinophyceae | Peridiniales | Peridiniaceae |
| | ASV1902 | 3.82 | 1.18E-11 | Eukaryota | Ciliophora | Heterotrichea | Heterotrichida | Spirostomidae |
| | ASV2254 | 3.82 | 9.06E-08 | Eukaryota | Cercozoa | Thecofilosea | Cryomonadida | Protaspidae |
| | ASV355 | 3.79 | 1.69E-07 | Eukaryota | Myzozoa | Dinophyceae | Peridiniales | Peridiniaceae |
| | ASV691 | 3.79 | 5.32E-12 | Eukaryota | Ciliophora | Heterotrichea | Heterotrichida | Spirostomidae |
| | ASV186 | 3.79 | 0.005986 | Eukaryota | Ochrophyta | Chrysophyceae | Synurales | Mallomonadaceae |
| | ASV1243 | 3.78 | 1.62E-09 | Eukaryota | Chlorophyta | Chlorophyceae | Chlamydomonadales | Volvocaceae |
| | ASV2839 | 3.76 | 8.34E-10 | Eukaryota | Bacillariophyta | Bacillariophyceae | Fragilariales | Fragilariaceae |
| | ASV1098 | 3.76 | 1.36E-07 | Eukaryota | Myzozoa | Dinophyceae | Suessiales | Suessiaceae |
| | ASV2003 | 3.74 | 3.17E-08 | Eukaryota | Myzozoa | Dinophyceae | Peridiniales | Peridiniaceae |
| | ASV1778 | 3.72 | 2.27E-07 | Eukaryota | Bacillariophyta | Bacillariophyceae | Fragilariales | Fragilariaceae |
| | ASV1716 | 3.70 | 1.52E-11 | Eukaryota | Myzozoa | Dinophyceae | Suessiales | Suessiaceae |
| | ASV2753 | 3.70 | 1.29E-10 | Eukaryota | Cercozoa | Imbricatea | Thaumatomonadida | Thaumatomastigidae |
| | ASV751 | 3.67 | 5.58E-08 | Eukaryota | Bacillariophyta | Bacillariophyceae | Fragilariales | Fragilariaceae |
| | ASV763 | 3.67 | 5.17E-10 | Eukaryota | Myzozoa | Dinophyceae | Suessiales | Suessiaceae |
| | ASV80 | 3.67 | 3.54E-10 | Eukaryota | Chlorophyta | Chlorophyceae | Sphaeropleales | Sphaeropleaceae |
| | ASV1367 | 3.66 | 8.39E-08 | Eukaryota | Myzozoa | Dinophyceae | Peridiniales | Peridiniaceae |
| | ASV2175 | 3.62 | 0.000277 | Eukaryota | Bacillariophyta | Mediophyceae | Stephanodiscales | Stephanodiscaceae |
| | ASV2549 | 3.61 | 5.61E-09 | Fungi | Blastocladiomycota | Blastocladiomycetes | Blastocladiales | Catenariaceae |
| | ASV1816 | 3.61 | 4.59E-10 | Eukaryota | Chlorophyta | Chlorophyceae | Sphaeropleales | Sphaeropleaceae |
| | ASV630 | 3.61 | 5.35E-13 | Eukaryota | Ciliophora | Oligohymenophorea | Astomatida | Haptophryidae |
| | ASV1718 | 3.57 | 3.72E-13 | Eukaryota | Oomycota | Peronosporea | Saprolegniales | Saprolegniaceae |
| | ASV1142 | 3.56 | 7.91E-13 | Eukaryota | Ciliophora | Oligohymenophorea | Pleuronematida | Conchophthiridae |
| | ASV1908 | 3.56 | 0.000383 | Eukaryota | Bacillariophyta | Mediophyceae | Stephanodiscales | Stephanodiscaceae |
| | ASV693 | 3.55 | 4.58E-15 | Eukaryota | Chlorophyta | Chlorophyceae | Chlamydomonadales | Chlamydomonadaceae |

| | | | | | | | | |
|---|---|---|---|---|---|---|---|---|
| | ASV1727 | 3.54 | 5.46E-08 | Eukaryota | Bacillariophyta | Bacillariophyceae | Fragilariales | Fragilariaceae |
| | ASV964 | 3.53 | 9.17E-08 | Eukaryota | Myzozoa | Dinophyceae | Suessiales | Suessiaceae |
| | ASV1537 | 3.53 | 6.00E-09 | Eukaryota | Chlorophyta | Chlorophyceae | Chlamydomonadales | Volvocaceae |
| | ASV657 | 3.53 | 5.08E-13 | Eukaryota | Ciliophora | Oligohymenophorea | Pleuronematida | Conchophthiridae |
| | ASV1804 | 3.52 | 8.54E-11 | Eukaryota | Ochrophyta | Chrysophyceae | Synurales | Mallomonadaceae |
| | ASV2760 | 3.51 | 8.91E-11 | Eukaryota | Myzozoa | Dinophyceae | Suessiales | Suessiaceae |
| | ASV2366 | 3.51 | 6.13E-09 | Eukaryota | Ciliophora | Spirotrichea | Tintinnida | Tintinnidae |
| | ASV340 | 3.51 | 8.18E-08 | Eukaryota | Ciliophora | Prostomatea | Prorodontida | Colepidae |
| | ASV438 | 3.50 | 1.37E-11 | Eukaryota | Chlorophyta | Chlorophyceae | Sphaeropleales | Neochloridaceae |
| | ASV2090 | 3.49 | 3.35E-12 | Eukaryota | Myzozoa | Dinophyceae | Gonyaulacales | Goniodomataceae |
| | ASV2618 | 3.47 | 3.48E-10 | Eukaryota | Ciliophora | Oligohymenophorea | Hymenostomatida | Ichthyophthiriidae |
| | ASV1263 | 3.45 | 6.18E-13 | Eukaryota | Ciliophora | Oligohymenophorea | Astomatida | Anoplophryidae |
| | ASV391 | 3.44 | 5.90E-13 | Eukaryota | Myzozoa | Dinophyceae | Suessiales | Suessiaceae |
| | ASV97 | 3.44 | 9.80E-09 | Eukaryota | Chlorophyta | Chlorophyceae | Sphaeropleales | Sphaeropleaceae |
| | ASV775 | 3.43 | 3.61E-12 | Eukaryota | Ciliophora | Oligohymenophorea | Pleuronematida | Conchophthiridae |
| | ASV482 | 3.43 | 1.38E-07 | Eukaryota | Myzozoa | Dinophyceae | Peridiniales | Peridiniaceae |
| | ASV1904 | 3.42 | 3.36E-12 | Eukaryota | Ciliophora | Oligohymenophorea | Astomatida | Haptophryidae |
| | ASV532 | 3.42 | 3.42E-06 | Eukaryota | Cercozoa | Thecofilosea | Cryomonadida | Protaspidae |
| | ASV1728 | 3.42 | 2.28E-11 | Eukaryota | Myzozoa | Dinophyceae | Suessiales | Suessiaceae |
| | ASV656 | 3.41 | 7.37E-09 | Eukaryota | Cercozoa | Protozoa classis incertae sedis | Protozoa ordo incertae sedis | Protozoa familia incertae sedis |
| | ASV431 | 3.41 | 8.03E-10 | Eukaryota | Ciliophora | Spirotrichea | Tintinnida | Tintinnidae |
| | ASV2336 | 3.41 | 4.19E-05 | Eukaryota | Bacillariophyta | Mediophyceae | Stephanodiscales | Stephanodiscaceae |
| | ASV1250 | 3.40 | 3.17E-08 | Fungi | Blastocladiomycota | Blastocladiomycetes | Blastocladiales | Catenariaceae |
| | ASV436 | 3.39 | 1.32E-10 | Eukaryota | Cercozoa | Thecofilosea | Cryomonadida | Protaspidae |
| | ASV2527 | 3.39 | 3.08E-07 | Eukaryota | Ciliophora | Prostomatea | Prorodontida | Colepidae |
| | ASV1325 | 3.38 | 3.40E-06 | Eukaryota | Myzozoa | Dinophyceae | Suessiales | Suessiaceae |
| | ASV770 | 3.37 | 7.48E-07 | Eukaryota | Myzozoa | Dinophyceae | Peridiniales | Peridiniaceae |
| | ASV442 | 3.35 | 6.43E-10 | Eukaryota | Ochrophyta | Chrysophyceae | Synurales | Mallomonadaceae |
| | ASV655 | 3.35 | 1.30E-05 | Eukaryota | Oomycota | Peronosporea | Saprolegniales | Saprolegniaceae |
| | ASV1410 | 3.34 | 2.58E-10 | Eukaryota | Chlorophyta | Chlorophyceae | Sphaeropleales | Sphaeropleaceae |

| | | | | | | | | |
|---|---|---|---|---|---|---|---|---|
| | ASV2849 | 3.33 | 1.84E-08 | Eukaryota | Ciliophora | Spirotrichea | Tintinnida | Tintinnidae |
| | ASV1269 | 3.33 | 1.66E-06 | Eukaryota | Cercozoa | Thecofilosea | Cryomonadida | Protaspidae |
| | ASV2691 | 3.31 | 5.27E-12 | Eukaryota | Oomycota | Peronosporea | Saprolegniales | Saprolegniaceae |
| | ASV84 | 3.31 | 2.07E-08 | Eukaryota | Chlorophyta | Chlorophyceae | Chlamydomonadales | Volvocaceae |
| | ASV1483 | 3.30 | 6.08E-06 | Eukaryota | Ciliophora | Prostomatea | Prorodontida | Colepidae |
| | ASV1242 | 3.29 | 5.82E-10 | Eukaryota | Chlorophyta | Chlorophyceae | Sphaeropleales | Sphaeropleaceae |
| | ASV707 | 3.29 | 6.12E-08 | Eukaryota | Ciliophora | Prostomatea | Prorodontida | Colepidae |
| | ASV875 | 3.29 | 2.86E-07 | Eukaryota | Myzozoa | Dinophyceae | Peridiniales | Pfiesteriaceae |
| | ASV2860 | 3.28 | 2.74E-09 | Eukaryota | Chlorophyta | Chlorophyceae | Chlamydomonadales | Volvocaceae |
| | ASV48 | 3.28 | 5.78E-08 | Eukaryota | Cercozoa | Thecofilosea | Cryomonadida | Protaspidae |
| | ASV2232 | 3.27 | 7.67E-10 | Eukaryota | Cercozoa | Protozoa classis incertae sedis | Protozoa ordo incertae sedis | Protozoa familia incertae sedis |
| | ASV766 | 3.26 | 0.000636 | Eukaryota | Chlorophyta | Chlorophyceae | Chlamydomonadales | Chlamydomonadaceae |
| | ASV2034 | 3.25 | 6.64E-07 | Eukaryota | Myzozoa | Dinophyceae | Peridiniales | Peridiniaceae |
| | ASV2168 | 3.24 | 2.85E-11 | Eukaryota | Ciliophora | Oligohymenophorea | Astomatida | Haptophryidae |
| | ASV1557 | 3.23 | 7.49E-05 | Eukaryota | Chlorophyta | Chlorophyceae | Chlamydomonadales | Chlamydomonadaceae |
| | ASV2372 | 3.22 | 0.000755 | Eukaryota | Chlorophyta | Chlorophyceae | Chlamydomonadales | Chlamydomonadaceae |
| | ASV1522 | 3.22 | 4.63E-06 | Eukaryota | Ciliophora | Prostomatea | Prorodontida | Colepidae |
| | ASV559 | 3.20 | 1.62E-09 | Eukaryota | Chlorophyta | Chlorophyceae | Sphaeropleales | Neochloridaceae |
| | ASV121 | 3.20 | 5.05E-05 | Eukaryota | Myzozoa | Dinophyceae | Suessiales | Suessiaceae |
| | ASV2576 | 3.17 | 4.27E-10 | Eukaryota | Myzozoa | Dinophyceae | Suessiales | Suessiaceae |
| | ASV2287 | 3.17 | 2.62E-05 | Eukaryota | Ciliophora | Spirotrichea | Urostylida | Holostichidae |
| | ASV2414 | 3.16 | 4.66E-11 | Eukaryota | Oomycota | Peronosporea | Saprolegniales | Saprolegniaceae |
| | ASV1389 | 3.14 | 0.000149 | Eukaryota | Ciliophora | Spirotrichea | Urostylida | Holostichidae |
| | ASV2838 | 3.14 | 2.81E-12 | Fungi | Chytridiomycota | Chytridiomycetes | Chytridiales | Chytriomycetaceae |
| | ASV1170 | 3.13 | 7.03E-12 | Eukaryota | Chlorophyta | Trebouxiophyceae | Chlorellales | Oocystaceae |
| | ASV341 | 3.11 | 0.000862 | Eukaryota | Chlorophyta | Chlorophyceae | Chlamydomonadales | Chlamydomonadaceae |
| | ASV35 | 3.11 | 5.98E-07 | Eukaryota | Myzozoa | Dinophyceae | Peridiniales | Pfiesteriaceae |
| | ASV506 | 3.09 | 7.69E-08 | Eukaryota | Bigyra | Bikosea | Bicosoecida | Bicosoecidae |
| | ASV896 | 3.09 | 1.08E-14 | Eukaryota | Ochrophyta | Chrysophyceae | Chromulinales | Dinobryaceae |
| | ASV545 | 3.08 | 0.000415 | Eukaryota | Bacillariophyta | Mediophyceae | Stephanodiscales | Stephanodiscaceae |

| | | | | | | | | |
|---|---|---|---|---|---|---|---|---|
| | ASV222 | 3.07 | 6.22E-11 | Eukaryota | Myzozoa | Dinophyceae | Peridiniales | Peridiniaceae |
| | ASV235 | 3.06 | 2.05E-08 | Eukaryota | Chlorophyta | Chlorophyceae | Chlamydomonadales | Volvocaceae |
| | ASV1881 | 3.06 | 0.000177 | Eukaryota | Ciliophora | Spirotrichea | Urostylida | Holostichidae |
| | ASV883 | 3.04 | 3.62E-09 | Eukaryota | Cercozoa | Protozoa classis incertae sedis | Protozoa ordo incertae sedis | Protozoa familia incertae sedis |
| | ASV1047 | 3.04 | 2.16E-06 | Eukaryota | Myzozoa | Dinophyceae | Peridiniales | Pfiesteriaceae |
| | ASV1712 | 3.02 | 3.36E-05 | Eukaryota | Ciliophora | Prostomatea | Prorodontida | Colepidae |
| | ASV1932 | 3.02 | 5.33E-10 | Eukaryota | Chlorophyta | Chlorophyceae | Sphaeropleales | Neochloridaceae |
| | ASV190 | 3.01 | 3.46E-10 | Eukaryota | Myzozoa | Dinophyceae | Gonyaulacales | Goniodomataceae |
| | ASV663 | 2.99 | 5.27E-12 | Fungi | Chytridiomycota | Chytridiomycetes | Chytridiales | Chytriomycetaceae |
| | ASV1892 | 2.99 | 3.13E-12 | Eukaryota | Chlorophyta | Trebouxiophyceae | Chlorellales | Oocystaceae |
| | ASV990 | 2.98 | 3.27E-13 | Eukaryota | Chlorophyta | Chlorophyceae | Chlamydomonadales | Chlamydomonadaceae |
| | ASV2289 | 2.97 | 3.58E-07 | Eukaryota | Cercozoa | Thecofilosea | Cryomonadida | Protaspidae |
| | ASV2314 | 2.96 | 8.41E-06 | Eukaryota | Myzozoa | Dinophyceae | Peridiniales | Peridiniaceae |
| | ASV2695 | 2.96 | 3.47E-05 | Eukaryota | Ciliophora | Prostomatea | Prorodontida | Colepidae |
| | ASV2686 | 2.96 | 1.77E-09 | Eukaryota | Myzozoa | Dinophyceae | Peridiniales | Peridiniaceae |
| | ASV2521 | 2.95 | 0.000138 | Eukaryota | Ciliophora | Spirotrichea | Urostylida | Holostichidae |
| | ASV1095 | 2.94 | 5.11E-05 | Eukaryota | Ochrophyta | Chrysophyceae | Chrysosphaerales | Chrysosphaeraceae |
| | ASV2807 | 2.94 | 1.22E-11 | Eukaryota | Chlorophyta | Trebouxiophyceae | Chlorellales | Oocystaceae |
| | ASV118 | 2.92 | 4.93E-08 | Eukaryota | Protozoa incertae sedis | Protozoa classis incertae sedis | Protozoa ordo incertae sedis | Alphamonaceae |
| | ASV1588 | 2.91 | 4.47E-10 | Eukaryota | Myzozoa | Dinophyceae | Suessiales | Suessiaceae |
| | ASV2567 | 2.90 | 0.000661 | Eukaryota | Myzozoa | Dinophyceae | Gymnodiniales | Gymnodiniaceae |
| | ASV1787 | 2.90 | 0.000113 | Eukaryota | Oomycota | Peronosporea | Saprolegniales | Saprolegniaceae |
| | ASV1637 | 2.89 | 6.08E-07 | Fungi | Blastocladiomycota | Blastocladiomycetes | Blastocladiales | Catenariaceae |
| | ASV1886 | 2.88 | 6.98E-05 | Eukaryota | Chlorophyta | Chlorophyceae | Chlamydomonadales | Chlamydomonadaceae |
| | ASV595 | 2.88 | 4.60E-08 | Fungi | Blastocladiomycota | Blastocladiomycetes | Blastocladiales | Catenariaceae |
| | ASV858 | 2.87 | 4.50E-07 | Eukaryota | Bigyra | Labyrinthulea | Thraustochytrida | Thraustochytriaceae |
| | ASV2210 | 2.87 | 1.35E-10 | Eukaryota | Ochrophyta | Chrysophyceae | Paraphysomonadales | Chrysosphaerellaceae |
| | ASV1656 | 2.87 | 4.93E-09 | Eukaryota | Myzozoa | Dinophyceae | Peridiniales | Peridiniaceae |
| | ASV2076 | 2.86 | 0.000898 | Eukaryota | Myzozoa | Dinophyceae | Gymnodiniales | Gymnodiniaceae |
| | ASV2191 | 2.86 | 1.74E-05 | Eukaryota | Chlorophyta | Chlorophyceae | Chlamydomonadales | Chlamydomonadaceae |

| | | | | | | | | |
|---|---|---|---|---|---|---|---|---|
| | ASV623 | 2.85 | 1.46E-09 | Eukaryota | Ciliophora | Oligohymenophorea | Hymenostomatida | Ichthyophthiriidae |
| | ASV1246 | 2.84 | 9.70E-08 | Eukaryota | Protozoa incertae sedis | Protozoa classis incertae sedis | Protozoa ordo incertae sedis | Alphamonaceae |
| | ASV206 | 2.84 | 8.56E-10 | Eukaryota | Myzozoa | Dinophyceae | Gonyaulacales | Goniodomataceae |
| | ASV920 | 2.82 | 2.47E-07 | Eukaryota | Bigyra | Bikosea | Bicosoecida | Bicosoecidae |
| | ASV1835 | 2.82 | 1.08E-07 | Fungi | Blastocladiomycota | Blastocladiomycetes | Blastocladiales | Catenariaceae |
| | ASV968 | 2.82 | 2.71E-09 | Eukaryota | Myzozoa | Dinophyceae | Suessiales | Suessiaceae |
| | ASV476 | 2.81 | 7.84E-09 | Eukaryota | Myzozoa | Dinophyceae | Peridiniales | Peridiniaceae |
| | ASV676 | 2.80 | 0.000452 | Eukaryota | Myzozoa | Dinophyceae | Suessiales | Suessiaceae |
| | ASV761 | 2.80 | 0.000852 | Eukaryota | Myzozoa | Dinophyceae | Gymnodiniales | Gymnodiniaceae |
| | ASV2136 | 2.77 | 1.32E-05 | Fungi | Chytridiomycota | Chytridiomycetes | Rhizophydiales | Kappamycetaceae |
| | ASV2024 | 2.77 | 3.70E-10 | Eukaryota | Myzozoa | Dinophyceae | Peridiniales | Peridiniaceae |
| | ASV622 | 2.77 | 4.93E-09 | Eukaryota | Myzozoa | Dinophyceae | Peridiniales | Peridiniaceae |
| | ASV338 | 2.75 | 8.73E-09 | Eukaryota | Ciliophora | Oligohymenophorea | Astomatida | Haptophryidae |
| | ASV1333 | 2.75 | 4.82E-13 | Fungi | Chytridiomycota | Chytridiomycetes | Rhizophydiales | Uebelmesseromycetaceae |
| | ASV2238 | 2.74 | 1.94E-09 | Eukaryota | Chlorophyta | Chlorophyceae | Sphaeropleales | Sphaeropleaceae |
| | ASV2717 | 2.73 | 1.24E-10 | Eukaryota | Myzozoa | Dinophyceae | Suessiales | Suessiaceae |
| | ASV632 | 2.72 | 7.78E-11 | Fungi | Chytridiomycota | Chytridiomycetes | Chytridiales | Chytriomycetaceae |
| | ASV768 | 2.70 | 0.000178 | Eukaryota | Myzozoa | Dinophyceae | Suessiales | Suessiaceae |
| | ASV861 | 2.70 | 4.42E-05 | Fungi | Chytridiomycota | Chytridiomycetes | Rhizophydiales | Kappamycetaceae |
| | ASV437 | 2.69 | 1.77E-09 | Eukaryota | Myzozoa | Dinophyceae | Peridiniales | Podolampaceae |
| | ASV1252 | 2.69 | 8.56E-10 | Eukaryota | Myzozoa | Dinophyceae | Peridiniales | Peridiniaceae |
| | ASV2592 | 2.69 | 0.00244 | Eukaryota | Chlorophyta | Chlorophyceae | Chlamydomonadales | Chlamydomonadaceae |
| | ASV1693 | 2.69 | 9.63E-09 | Eukaryota | Myzozoa | Dinophyceae | Peridiniales | Peridiniaceae |
| | ASV2334 | 2.68 | 1.45E-08 | Eukaryota | Myzozoa | Dinophyceae | Peridiniales | Peridiniaceae |
| | ASV2458 | 2.68 | 0.002035 | Eukaryota | Ciliophora | Spirotrichea | Tintinnida | Codonellidae |
| | ASV629 | 2.68 | 2.88E-07 | Eukaryota | Ochrophyta | Chrysophyceae | Paraphysomonadales | Chrysosphaerellaceae |
| | ASV330 | 2.68 | 1.10E-09 | Eukaryota | Cercozoa | Imbricatea | Thaumatomonadida | Thaumatomastigidae |
| | ASV152 | 2.67 | 0.000287 | Eukaryota | Bigyra | incertae sedis | incertae sedis | Pseudophyllomitidae |
| | ASV985 | 2.66 | 0.000234 | Eukaryota | Chlorophyta | Chlorophyceae | Chlamydomonadales | Chlamydomonadaceae |
| | ASV2714 | 2.66 | 8.28E-07 | Eukaryota | Chlorophyta | Chlorophyceae | Chlamydomonadales | Chlamydomonadaceae |

| | | | | | | | | |
|---|---|---|---|---|---|---|---|---|
| | ASV47 | 2.66 | 4.93E-09 | Eukaryota | Ciliophora | Prostomatea | Prorodontida | Colepidae |
| | ASV1788 | 2.65 | 0.000767 | Eukaryota | Myzozoa | Dinophyceae | Gymnodiniales | Gymnodiniaceae |
| | ASV1201 | 2.65 | 2.69E-08 | Eukaryota | Myzozoa | Dinophyceae | Peridiniales | Peridiniaceae |
| | ASV2505 | 2.65 | 1.66E-10 | Eukaryota | Chlorophyta | Trebouxiophyceae | Chlorellales | Eremosphaeraceae |
| | ASV1022 | 2.64 | 1.64E-08 | Eukaryota | Myzozoa | Dinophyceae | Peridiniales | Podolampaceae |
| | ASV125 | 2.64 | 0.000113 | Eukaryota | Myzozoa | Dinophyceae | Peridiniales | Podolampaceae |
| | ASV2064 | 2.63 | 0.000805 | Eukaryota | Myzozoa | Dinophyceae | Suessiales | Suessiaceae |
| | ASV2482 | 2.62 | 1.00E-05 | Eukaryota | Ciliophora | Litostomatea | Haptorida | Spathidiidae |
| | ASV1123 | 2.60 | 0.000342 | Eukaryota | Chlorophyta | Chlorophyceae | Chlamydomonadales | Chlamydomonadaceae |
| | ASV38 | 2.60 | 0.000552 | Eukaryota | Chlorophyta | Chlorophyceae | Chlamydomonadales | Chlamydomonadaceae |
| | ASV2183 | 2.58 | 4.84E-06 | Eukaryota | Myzozoa | Dinophyceae | Peridiniales | Pfiesteriaceae |
| | ASV2357 | 2.56 | 9.26E-09 | Eukaryota | Bacillariophyta | Bacillariophyceae | Fragilariales | Fragilariaceae |
| | ASV2788 | 2.56 | 4.78E-06 | Eukaryota | Bacillariophyta | Bacillariophyceae | Fragilariales | Fragilariaceae |
| | ASV1412 | 2.55 | 1.40E-10 | Eukaryota | Ochrophyta | Chrysophyceae | Chromulinales | Dinobryaceae |
| | ASV1183 | 2.53 | 9.72E-06 | Eukaryota | Ciliophora | Litostomatea | Haptorida | Spathidiidae |
| | ASV1224 | 2.53 | 7.20E-08 | Eukaryota | Cryptophyta | Cryptophyceae | Kathablepharidacea | Katablepharidaceae |
| | ASV1920 | 2.53 | 9.99E-11 | Eukaryota | Choanozoa | Ichthyosporea | Ichthyophonida | Ichthyophonidae |
| | ASV557 | 2.52 | 0.000138 | Eukaryota | Oomycota | Peronosporea | Saprolegniales | Saprolegniaceae |
| | ASV2664 | 2.51 | 4.38E-06 | Fungi | Blastocladiomycota | Blastocladiomycetes | Blastocladiales | Catenariaceae |
| | ASV1694 | 2.51 | 2.07E-08 | Eukaryota | Ciliophora | Oligohymenophorea | Hymenostomatida | Ichthyophthiriidae |
| | ASV56 | 2.50 | 0.002561 | Eukaryota | Cryptophyta | Cryptophyceae | Cryptomonadales | Cryptomonadaceae |
| | ASV2782 | 2.50 | 1.04E-05 | Eukaryota | Ciliophora | Litostomatea | Haptorida | Spathidiidae |
| | ASV747 | 2.50 | 3.44E-08 | Eukaryota | Chlorophyta | Chlorophyceae | Chlamydomonadales | Chlamydomonadaceae |
| | ASV535 | 2.49 | 1.30E-10 | Eukaryota | Chlorophyta | Chlorodendrophyceae | Chlorodendrales | Chlorodendraceae |
| | ASV1318 | 2.49 | 3.81E-10 | Eukaryota | Choanozoa | Choanoflagellata | Craspedida | Codonosigaceae |
| | ASV729 | 2.48 | 1.49E-06 | Fungi | Chytridiomycota | Chytridiomycetes | Rhizophydiales | Uebelmesseromycetaceae |
| | ASV1400 | 2.48 | 8.67E-07 | Eukaryota | Myzozoa | Dinophyceae | Suessiales | Suessiaceae |
| | ASV1195 | 2.47 | 7.22E-07 | Fungi | Chytridiomycota | Chytridiomycetes | Rhizophydiales | Uebelmesseromycetaceae |
| | ASV839 | 2.46 | 5.09E-08 | Eukaryota | Myzozoa | Dinophyceae | Peridiniales | Peridiniaceae |
| | ASV2733 | 2.45 | 8.82E-05 | Eukaryota | Charophyta | Zygnemophyceae | Desmidiales | Closteriaceae |

| | | | | | | | | |
|---|---|---|---|---|---|---|---|---|
| | ASV1435 | 2.44 | 3.07E-08 | Eukaryota | Cryptophyta | Cryptophyceae | Kathablepharidacea | Katablepharidaceae |
| | ASV369 | 2.44 | 3.65E-05 | Fungi | Chytridiomycota | Chytridiomycetes | Rhizophydiales | Kappamycetaceae |
| | ASV1084 | 2.43 | 2.86E-06 | Eukaryota | Ciliophora | Spirotrichea | Sporadotrichida | Oxytrichidae |
| | ASV2876 | 2.43 | 4.20E-06 | Eukaryota | Cercozoa | Thecofilosea | Cryomonadida | Protaspidae |
| | ASV334 | 2.43 | 0.003715 | Eukaryota | Oomycota | Peronosporea | Saprolegniales | Saprolegniaceae |
| | ASV1073 | 2.42 | 9.46E-05 | Eukaryota | Cercozoa | Thecofilosea | Cryomonadida | Protaspidae |
| | ASV1310 | 2.42 | 4.92E-10 | Eukaryota | Myzozoa | Dinophyceae | Peridiniales | Podolampaceae |
| | ASV867 | 2.42 | 0.000194 | Eukaryota | Oomycota | Peronosporea | Saprolegniales | Saprolegniaceae |
| | ASV1597 | 2.41 | 1.69E-08 | Eukaryota | Bacillariophyta | Bacillariophyceae | Fragilariales | Fragilariaceae |
| | ASV140 | 2.39 | 0.004993 | Eukaryota | Ciliophora | Spirotrichea | Tintinnida | Codonellidae |
| | ASV878 | 2.38 | 4.60E-08 | Eukaryota | Myzozoa | Dinophyceae | Peridiniales | Peridiniaceae |
| | ASV631 | 2.37 | 8.18E-10 | Fungi | Chytridiomycota | Chytridiomycetes | Chytridiales | Chytriomycetaceae |
| | ASV484 | 2.37 | 3.17E-08 | Eukaryota | Myzozoa | Dinophyceae | Peridiniales | Peridiniaceae |
| | ASV838 | 2.35 | 1.35E-05 | Eukaryota | Ciliophora | Spirotrichea | Sporadotrichida | Oxytrichidae |
| | ASV1112 | 2.33 | 2.41E-05 | Eukaryota | Chlorophyta | Chlorophyceae | Chlamydomonadales | Chlamydomonadaceae |
| | ASV1427 | 2.33 | 4.64E-11 | Fungi | Ascomycota | Leotiomycetes | Chaetomellales | Chaetomellaceae |
| | ASV760 | 2.33 | 1.82E-07 | Fungi | Blastocladiomycota | Blastocladiomycetes | Blastocladiales | Catenariaceae |
| | ASV1838 | 2.32 | 0.002623 | Eukaryota | Ciliophora | Spirotrichea | Tintinnida | Tintinnidae |
| | ASV1501 | 2.32 | 4.03E-06 | Eukaryota | Bacillariophyta | Bacillariophyceae | Fragilariales | Fragilariaceae |
| | ASV1514 | 2.31 | 0.000149 | Eukaryota | Ciliophora | Prostomatea | Prorodontida | Colepidae |
| | ASV1293 | 2.31 | 1.31E-06 | Eukaryota | Ciliophora | Litostomatea | Haptorida | Spathidiidae |
| | ASV745 | 2.31 | 3.64E-06 | Eukaryota | Myzozoa | Dinophyceae | Suessiales | Suessiaceae |
| | ASV78 | 2.30 | 3.10E-06 | Eukaryota | Ochrophyta | Chrysophyceae | Synurales | Mallomonadaceae |
| | ASV1237 | 2.29 | 2.24E-09 | Eukaryota | Chlorophyta | Chlorophyceae | Sphaeropleales | Neochloridaceae |
| | ASV1790 | 2.29 | 1.19E-07 | Eukaryota | Oomycota | Peronosporea | Saprolegniales | Saprolegniaceae |
| | ASV1109 | 2.29 | 3.28E-06 | Fungi | Blastocladiomycota | Blastocladiomycetes | Blastocladiales | Catenariaceae |
| | ASV1841 | 2.29 | 0.000224 | Eukaryota | Myzozoa | Dinophyceae | Peridiniales | Podolampaceae |
| | ASV2317 | 2.28 | 2.23E-08 | Eukaryota | Ciliophora | Odontostomatea | Odontostomatida | Epalxellidae |
| | ASV1586 | 2.27 | 0.000319 | Eukaryota | Ochrophyta | Chrysophyceae | Synurales | Mallomonadaceae |
| | ASV2348 | 2.27 | 0.00097 | Eukaryota | Myzozoa | Dinophyceae | Peridiniales | Podolampaceae |

| | | | | | | | | |
|---|---|---|---|---|---|---|---|---|
| | ASV2689 | 2.25 | 8.39E-08 | Eukaryota | Ciliophora | Prostomatea | Prorodontida | Colepidae |
| | ASV1625 | 2.25 | 0.001365 | Eukaryota | Ciliophora | Prostomatea | Prorodontida | Colepidae |
| | ASV423 | 2.25 | 3.47E-05 | Fungi | Chytridiomycota | Chytridiomycetes | Rhizophydiales | Kappamycetaceae |
| | ASV2330 | 2.25 | 0.00565 | Eukaryota | Bacillariophyta | Mediophyceae | Stephanodiscales | Stephanodiscaceae |
| | ASV859 | 2.24 | 5.25E-07 | Eukaryota | Chlorophyta | Chlorophyceae | Chlamydomonadales | Chlamydomonadaceae |
| | ASV2550 | 2.24 | 4.93E-09 | Eukaryota | Chlorophyta | Trebouxiophyceae | Chlorellales | Eremosphaeraceae |
| | ASV519 | 2.23 | 3.17E-08 | Fungi | Chytridiomycota | Chytridiomycetes | Chytridiales | Chytriomycetaceae |
| | ASV816 | 2.22 | 1.64E-08 | Eukaryota | Chlorophyta | Chlorophyceae | Chlamydomonadales | Chlamydomonadaceae |
| | ASV1105 | 2.22 | 0.000358 | Eukaryota | Ciliophora | Prostomatea | Prorodontida | Colepidae |
| | ASV1319 | 2.21 | 0.000994 | Eukaryota | Ciliophora | Prostomatea | Prorodontida | Colepidae |
| | ASV1539 | 2.21 | 0.002564 | Eukaryota | Ciliophora | Spirotrichea | Urostylida | Holostichidae |
| | ASV1632 | 2.21 | 0.002639 | Eukaryota | Myzozoa | Dinophyceae | Peridiniales | Podolampaceae |
| | ASV607 | 2.21 | 0.000118 | Eukaryota | Cercozoa | Thecofilosea | Cryomonadida | Protaspidae |
| | ASV1223 | 2.21 | 6.59E-09 | Eukaryota | Myzozoa | Dinophyceae | Peridiniales | Peridiniaceae |
| | ASV1009 | 2.20 | 4.08E-06 | Eukaryota | Chlorophyta | Chlorophyceae | Chlamydomonadales | Chlamydomonadaceae |
| | ASV155 | 2.19 | 0.000636 | Eukaryota | Ciliophora | Prostomatea | Prorodontida | Colepidae |
| | ASV2605 | 2.19 | 4.72E-06 | Eukaryota | Chlorophyta | Chlorophyceae | Chlamydomonadales | Chlamydomonadaceae |
| | ASV704 | 2.19 | 7.18E-05 | Eukaryota | Ciliophora | Litostomatea | Haptorida | Spathidiidae |
| | ASV359 | 2.18 | 7.78E-07 | Eukaryota | Bigyra | Bikosea | Bicosoecida | Bicosoecidae |
| | ASV2515 | 2.18 | 7.10E-10 | Eukaryota | Chlorophyta | Trebouxiophyceae | Chlorellales | Eremosphaeraceae |
| | ASV368 | 2.17 | 0.002991 | Eukaryota | Chlorophyta | Chlorophyceae | Chlamydomonadales | Chlamydomonadaceae |
| | ASV266 | 2.16 | 3.12E-06 | Eukaryota | Protozoa incertae sedis | Protozoa classis incertae sedis | Protozoa ordo incertae sedis | Alphamonaceae |
| | ASV1913 | 2.16 | 2.36E-07 | Eukaryota | Myzozoa | Dinophyceae | Peridiniales | Podolampaceae |
| | ASV1710 | 2.16 | 2.90E-07 | Eukaryota | Charophyta | Zygnemophyceae | Desmidiales | Desmidiaceae |
| | ASV2692 | 2.16 | 0.000118 | Eukaryota | Charophyta | Zygnemophyceae | Desmidiales | Closteriaceae |
| | ASV2422 | 2.16 | 1.26E-08 | Eukaryota | Cryptophyta | Cryptophyceae | Cryptomonadales | Cryptomonadaceae |
| | ASV1890 | 2.16 | 1.05E-08 | Eukaryota | Cryptophyta | Cryptophyceae | Kathablepharidacea | Katablepharidaceae |
| | ASV1032 | 2.15 | 8.61E-08 | Eukaryota | Oomycota | Peronosporea | Saprolegniales | Saprolegniaceae |
| | ASV777 | 2.15 | 4.70E-10 | Eukaryota | Bigyra | Bikosea | Bicosoecida | Bicosoecidae |
| | ASV2447 | 2.15 | 3.91E-06 | Eukaryota | Ochrophyta | Chrysophyceae | Paraphysomonadales | Chrysosphaerellaceae |

| | | | | | | | | |
|---|---|---|---|---|---|---|---|---|
| | ASV1285 | 2.14 | 0.000108 | Eukaryota | Cercozoa | Protozoa classis incertae sedis | Protozoa ordo incertae sedis | Protozoa familia incertae sedis |
| | ASV2862 | 2.14 | 1.15E-06 | Eukaryota | Cercozoa | Imbricatea | Spongomonadida | Spongomonadidae |
| | ASV2863 | 2.14 | 2.42E-06 | Fungi | Chytridiomycota | Chytridiomycetes | Rhizophydiales | Uebelmesseromycetaceae |
| | ASV2534 | 2.13 | 0.002964 | Eukaryota | Ciliophora | Spirotrichea | Urostylida | Holostichidae |
| | ASV2117 | 2.13 | 0.005047 | Eukaryota | Ciliophora | Spirotrichea | Urostylida | Holostichidae |
| | ASV180 | 2.13 | 2.75E-06 | Eukaryota | Cercozoa | Thecofilosea | Cryomonadida | Protaspidae |
| | ASV2052 | 2.13 | 2.24E-08 | Eukaryota | Chlorophyta | Chlorophyceae | Sphaeropleales | Scenedesmaceae |
| | ASV1425 | 2.12 | 0.004993 | Eukaryota | Chlorophyta | Chlorophyceae | Chlamydomonadales | Chlamydomonadaceae |
| | ASV2460 | 2.12 | 6.49E-07 | Eukaryota | Myzozoa | Dinophyceae | Gymnodiniales | Gymnodiniaceae |
| | ASV611 | 2.11 | 2.79E-07 | Eukaryota | Oomycota | Peronosporea | Saprolegniales | Saprolegniaceae |
| | ASV402 | 2.09 | 0.004718 | Eukaryota | Ciliophora | Spirotrichea | Urostylida | Holostichidae |
| | ASV2857 | 2.09 | 7.13E-08 | Eukaryota | Chlorophyta | Chlorophyceae | Chlamydomonadales | Chlamydomonadaceae |
| | ASV1802 | 2.08 | 0.000265 | Eukaryota | Myzozoa | Dinophyceae | Suessiales | Suessiaceae |
| | ASV2391 | 2.07 | 0.000178 | Eukaryota | Ochrophyta | Chrysophyceae | Chrysosphaerales | Chrysosphaeraceae |
| | ASV1221 | 2.05 | 1.34E-06 | Eukaryota | Ciliophora | Spirotrichea | Sporadotrichida | Oxytrichidae |
| | ASV2185 | 2.05 | 0.002415 | Eukaryota | Cryptophyta | Cryptophyceae | Kathablepharidacea | Katablepharidaceae |
| | ASV2808 | 2.05 | 0.000958 | Eukaryota | Myzozoa | Dinophyceae | Prorocentrales | Prorocentraceae |
| | ASV92 | 2.04 | 0.006192 | Eukaryota | Ochrophyta | Chrysophyceae | Chrysosphaerales | Chrysosphaeraceae |
| | ASV980 | 2.04 | 0.000108 | Eukaryota | Ciliophora | Spirotrichea | Sporadotrichida | Oxytrichidae |
| | ASV2280 | 2.03 | 0.008355 | Eukaryota | Bacillariophyta | Mediophyceae | Stephanodiscales | Stephanodiscaceae |
| | ASV2546 | 2.03 | 6.43E-05 | Eukaryota | Myzozoa | Dinophyceae | Prorocentrales | Prorocentraceae |
| | ASV1715 | 2.02 | 5.33E-05 | Eukaryota | Bacillariophyta | Bacillariophyceae | Fragilariales | Fragilariaceae |
| | ASV1093 | 2.02 | 0.000719 | Eukaryota | Ochrophyta | Chrysophyceae | Ochromonadales | Ochromonadaceae |
| | ASV1029 | 2.02 | 0.000952 | Eukaryota | Ciliophora | Spirotrichea | Choreotrichida | Strombidinopsidae |
| | ASV2116 | 2.01 | 3.99E-06 | Eukaryota | Ochrophyta | Chrysophyceae | Synurales | Mallomonadaceae |
| | ASV1030 | 2.00 | 0.000919 | Eukaryota | Ochrophyta | Chrysophyceae | Synurales | Mallomonadaceae |
| 18S-PP | ASV1076 | 8.71 | 2.35E-12 | Eukaryota | Cercozoa | Protozoa classis incertae sedis | Protozoa ordo incertae sedis | Protozoa familia incertae sedis |
| | ASV711 | 8.37 | 6.69E-34 | Eukaryota | Myzozoa | Dinophyceae | Peridiniales | Protoperidiniaceae |
| | ASV2002 | 8.34 | 5.49E-32 | Eukaryota | Myzozoa | Dinophyceae | Prorocentrales | Prorocentraceae |
| | ASV1217 | 8.23 | 6.04E-30 | Eukaryota | Myzozoa | Dinophyceae | Peridiniales | Protoperidiniaceae |

| | | | | | | | | |
|---|---|---|---|---|---|---|---|---|
| | ASV2474 | 8.19 | 1.00E-27 | Eukaryota | Myzozoa | Dinophyceae | Prorocentrales | Prorocentraceae |
| | ASV2232 | 8.05 | 2.92E-14 | Eukaryota | Cercozoa | Protozoa classis incertae sedis | Protozoa ordo incertae sedis | Protozoa familia incertae sedis |
| | ASV2302 | 8.01 | 7.57E-32 | Eukaryota | Myzozoa | Dinophyceae | Peridiniales | Protoperidiniaceae |
| | ASV1797 | 7.98 | 1.07E-31 | Eukaryota | Myzozoa | Dinophyceae | Prorocentrales | Prorocentraceae |
| | ASV656 | 7.91 | 6.05E-13 | Eukaryota | Cercozoa | Protozoa classis incertae sedis | Protozoa ordo incertae sedis | Protozoa familia incertae sedis |
| | ASV883 | 7.61 | 2.90E-14 | Eukaryota | Cercozoa | Protozoa classis incertae sedis | Protozoa ordo incertae sedis | Protozoa familia incertae sedis |
| | ASV1391 | 7.58 | 5.32E-27 | Eukaryota | Myzozoa | Dinophyceae | Prorocentrales | Prorocentraceae |
| | ASV184 | 7.41 | 4.24E-25 | Eukaryota | Myzozoa | Dinophyceae | Peridiniales | Protoperidiniaceae |
| | ASV389 | 7.12 | 3.30E-28 | Eukaryota | Myzozoa | Dinophyceae | Peridiniales | Diplopsaliaceae |
| | ASV2202 | 7.04 | 1.80E-26 | Eukaryota | Myzozoa | Dinophyceae | Peridiniales | Diplopsaliaceae |
| | ASV2827 | 6.72 | 2.88E-26 | Eukaryota | Myzozoa | Dinophyceae | Peridiniales | Diplopsaliaceae |
| | ASV2726 | 6.62 | 2.92E-14 | Eukaryota | Bigyra | incertae sedis | incertae sedis | Pseudophyllomitidae |
| | ASV1022 | 6.57 | 2.85E-14 | Eukaryota | Myzozoa | Dinophyceae | Peridiniales | Podolampaceae |
| | ASV1615 | 6.49 | 2.36E-13 | Eukaryota | Ciliophora | Spirotrichea | Urostylida | Holostichidae |
| | ASV437 | 6.46 | 1.16E-13 | Eukaryota | Myzozoa | Dinophyceae | Peridiniales | Podolampaceae |
| | ASV2359 | 6.45 | 3.84E-26 | Eukaryota | Myzozoa | Dinophyceae | Peridiniales | Diplopsaliaceae |
| | ASV2090 | 6.38 | 4.79E-14 | Eukaryota | Myzozoa | Dinophyceae | Gonyaulacales | Goniodomataceae |
| | ASV638 | 6.28 | 9.57E-10 | Eukaryota | Cryptophyta | Cryptophyceae | Cryptomonadales | Cryptomonadaceae |
| | ASV787 | 6.25 | 5.35E-11 | Eukaryota | Cryptophyta | Cryptophyceae | Cryptomonadales | Cryptomonadaceae |
| | ASV1862 | 6.24 | 2.46E-15 | Eukaryota | Bigyra | incertae sedis | incertae sedis | Pseudophyllomitidae |
| | ASV458 | 6.23 | 1.17E-15 | Eukaryota | Chlorophyta | Chlorophyceae | Sphaeropleales | Radiococcaceae |
| | ASV2587 | 6.23 | 1.13E-12 | Eukaryota | Bigyra | incertae sedis | incertae sedis | Pseudophyllomitidae |
| | ASV2557 | 6.20 | 2.94E-16 | Eukaryota | Chlorophyta | Chlorophyceae | Sphaeropleales | Radiococcaceae |
| | ASV1010 | 6.20 | 2.86E-12 | Eukaryota | Ciliophora | Spirotrichea | Urostylida | Holostichidae |
| | ASV2650 | 6.15 | 2.81E-10 | Eukaryota | Cryptophyta | Cryptophyceae | Cryptomonadales | Cryptomonadaceae |
| | ASV1310 | 6.13 | 2.90E-14 | Eukaryota | Myzozoa | Dinophyceae | Peridiniales | Podolampaceae |
| | ASV190 | 6.13 | 4.27E-13 | Eukaryota | Myzozoa | Dinophyceae | Gonyaulacales | Goniodomataceae |
| | ASV1913 | 6.10 | 8.17E-14 | Eukaryota | Myzozoa | Dinophyceae | Peridiniales | Podolampaceae |
| | ASV554 | 6.05 | 4.26E-15 | Eukaryota | Myzozoa | Dinophyceae | Suessiales | Suessiaceae |
| | ASV2200 | 6.04 | 1.92E-16 | Eukaryota | Myzozoa | Dinophyceae | Suessiales | Suessiaceae |

| | | | | | | | | |
|---|---|---|---|---|---|---|---|---|
| | ASV597 | 6.04 | 2.94E-16 | Eukaryota | Chlorophyta | Chlorophyceae | Chlamydomonadales | Volvocaceae |
| | ASV2877 | 6.03 | 2.89E-13 | Eukaryota | Bigyra | incertae sedis | incertae sedis | Pseudophyllomitidae |
| | ASV206 | 5.96 | 6.58E-13 | Eukaryota | Myzozoa | Dinophyceae | Gonyaulacales | Goniodomataceae |
| | ASV167 | 5.91 | 1.79E-12 | Eukaryota | Ciliophora | Spirotrichea | Urostylida | Holostichidae |
| | ASV2268 | 5.91 | 2.42E-10 | Eukaryota | Cryptophyta | Cryptophyceae | Cryptomonadales | Cryptomonadaceae |
| | ASV511 | 5.85 | 1.76E-13 | Eukaryota | Myzozoa | Dinophyceae | Gonyaulacales | Goniodomataceae |
| | ASV1157 | 5.82 | 2.79E-11 | Eukaryota | Ciliophora | Spirotrichea | Urostylida | Holostichidae |
| | ASV1153 | 5.80 | 3.82E-11 | Eukaryota | Chlorophyta | Chlorophyceae | Sphaeropleales | Radiococcaceae |
| | ASV2802 | 5.80 | 5.87E-12 | Eukaryota | Chlorophyta | Chlorophyceae | Sphaeropleales | Radiococcaceae |
| | ASV235 | 5.79 | 2.59E-16 | Eukaryota | Chlorophyta | Chlorophyceae | Chlamydomonadales | Volvocaceae |
| | ASV345 | 5.78 | 3.75E-12 | Eukaryota | Chlorophyta | Chlorophyceae | Sphaeropleales | Radiococcaceae |
| | ASV843 | 5.77 | 1.26E-14 | Eukaryota | Chlorophyta | Chlorophyceae | Chlamydomonadales | Volvocaceae |
| | ASV988 | 5.71 | 6.20E-13 | Eukaryota | Chlorophyta | Chlorophyceae | Sphaeropleales | Radiococcaceae |
| | ASV431 | 5.69 | 9.52E-11 | Eukaryota | Ciliophora | Spirotrichea | Tintinnida | Tintinnidae |
| | ASV2849 | 5.63 | 3.15E-09 | Eukaryota | Ciliophora | Spirotrichea | Tintinnida | Tintinnidae |
| | ASV2366 | 5.61 | 3.42E-10 | Eukaryota | Ciliophora | Spirotrichea | Tintinnida | Tintinnidae |
| | ASV1158 | 5.60 | 1.70E-10 | Eukaryota | Cercozoa | Sarcomonadea | Glissomonadida | Viridiraptoridae |
| | ASV1292 | 5.58 | 2.77E-13 | Eukaryota | Chlorophyta | Chlorophyceae | Sphaeropleales | Radiococcaceae |
| | ASV1740 | 5.55 | 3.09E-12 | Eukaryota | Chlorophyta | Chlorophyceae | Sphaeropleales | Radiococcaceae |
| | ASV2092 | 5.54 | 5.94E-12 | Eukaryota | Chlorophyta | Chlorophyceae | Sphaeropleales | Radiococcaceae |
| | ASV555 | 5.53 | 3.63E-12 | Eukaryota | Chlorophyta | Chlorophyceae | Sphaeropleales | Radiococcaceae |
| | ASV706 | 5.52 | 2.28E-14 | Eukaryota | Chlorophyta | Chlorophyceae | Sphaeropleales | Radiococcaceae |
| | ASV560 | 5.49 | 1.54E-13 | Eukaryota | Chlorophyta | Chlorophyceae | Sphaeropleales | Radiococcaceae |
| | ASV842 | 5.44 | 1.07E-12 | Eukaryota | Myzozoa | Dinophyceae | Suessiales | Suessiaceae |
| | ASV1469 | 5.37 | 2.81E-09 | Eukaryota | Cryptophyta | Cryptophyceae | Cryptomonadales | Cryptomonadaceae |
| | ASV217 | 5.34 | 2.18E-11 | Eukaryota | Chlorophyta | Chlorophyceae | Sphaeropleales | Radiococcaceae |
| | ASV2850 | 5.32 | 1.05E-14 | Eukaryota | Chlorophyta | Chlorophyceae | Chlamydomonadales | Volvocaceae |
| | ASV1878 | 5.32 | 4.64E-09 | Eukaryota | Cercozoa | Sarcomonadea | Cercomonadida | Cercomonadidae |
| | ASV139 | 5.30 | 6.40E-10 | Eukaryota | Chlorophyta | Chlorophyceae | Sphaeropleales | Radiococcaceae |
| | ASV427 | 5.27 | 1.72E-11 | Eukaryota | Bigyra | incertae sedis | incertae sedis | Pseudophyllomitidae |

| | | | | | | | | |
|---|---|---|---|---|---|---|---|---|
| | ASV641 | 5.24 | 8.08E-06 | Eukaryota | Ciliophora | Spirotrichea | Tintinnida | Tintinnidae |
| | ASV1041 | 5.23 | 5.92E-12 | Eukaryota | Chlorophyta | Chlorophyceae | Sphaeropleales | Radiococcaceae |
| | ASV1177 | 5.21 | 1.45E-12 | Eukaryota | Chlorophyta | Chlorophyceae | Sphaeropleales | Radiococcaceae |
| | ASV1495 | 5.15 | 1.44E-05 | Eukaryota | Ciliophora | Spirotrichea | Tintinnida | Tintinnidae |
| | ASV1077 | 5.12 | 4.28E-09 | Eukaryota | Ciliophora | Oligotrichea | Oligotrichida | Strombidiidae |
| | ASV1209 | 5.11 | 2.01E-09 | Eukaryota | Ciliophora | Oligotrichea | Oligotrichida | Strombidiidae |
| | ASV1706 | 5.01 | 1.72E-08 | Eukaryota | Cryptophyta | Cryptophyceae | Cryptomonadales | Cryptomonadaceae |
| | ASV328 | 5.01 | 3.07E-09 | Eukaryota | Bigyra | Bikosea | Bicosoecida | Siluaniidae |
| | ASV2118 | 5.00 | 2.94E-16 | Fungi | Chytridiomycota | Chytridiomycetes | Rhizophydiales | Uebelmesseromycetaceae |
| | ASV1246 | 4.97 | 3.48E-11 | Eukaryota | Protozoa incertae sedis | Protozoa classis incertae sedis | Protozoa ordo incertae sedis | Alphamonaceae |
| | ASV1966 | 4.97 | 1.52E-05 | Eukaryota | Ciliophora | Spirotrichea | Tintinnida | Tintinnidae |
| | ASV118 | 4.92 | 1.82E-12 | Eukaryota | Protozoa incertae sedis | Protozoa classis incertae sedis | Protozoa ordo incertae sedis | Alphamonaceae |
| | ASV679 | 4.92 | 3.59E-15 | Fungi | Chytridiomycota | Chytridiomycetes | Rhizophydiales | Uebelmesseromycetaceae |
| | ASV1164 | 4.92 | 1.03E-08 | Eukaryota | Cercozoa | Sarcomonadea | Cercomonadida | Cercomonadidae |
| | ASV2026 | 4.90 | 9.45E-10 | Eukaryota | Cryptophyta | Cryptophyceae | Cryptomonadales | Cryptomonadaceae |
| | ASV1733 | 4.88 | 8.54E-14 | Eukaryota | Myzozoa | Dinophyceae | Peridiniales | Podolampaceae |
| | ASV2153 | 4.83 | 6.40E-07 | Eukaryota | Bigyra | Bikosea | Bicosoecida | Bicosoecidae |
| | ASV1522 | 4.83 | 7.94E-07 | Eukaryota | Ciliophora | Prostomatea | Prorodontida | Colepidae |
| | ASV1616 | 4.80 | 1.95E-16 | Eukaryota | Myzozoa | Dinophyceae | Peridiniales | Podolampaceae |
| | ASV171 | 4.75 | 2.18E-11 | Eukaryota | Myzozoa | Dinophyceae | Suessiales | Suessiaceae |
| | ASV158 | 4.74 | 3.13E-12 | Eukaryota | Myzozoa | Dinophyceae | Peridiniales | Podolampaceae |
| | ASV1231 | 4.73 | 8.55E-09 | Eukaryota | Ciliophora | Oligotrichea | Oligotrichida | Strombidiidae |
| | ASV2487 | 4.72 | 3.02E-06 | Eukaryota | Bigyra | Bikosea | Bicosoecida | Bicosoecidae |
| | ASV57 | 4.69 | 2.74E-12 | Eukaryota | Bigyra | incertae sedis | incertae sedis | Pseudophyllomitidae |
| | ASV2785 | 4.69 | 6.06E-09 | Eukaryota | Bigyra | Bikosea | Bicosoecida | Siluaniidae |
| | ASV2838 | 4.68 | 2.02E-13 | Fungi | Chytridiomycota | Chytridiomycetes | Chytridiales | Chytriomycetaceae |
| | ASV663 | 4.65 | 5.48E-13 | Fungi | Chytridiomycota | Chytridiomycetes | Chytridiales | Chytriomycetaceae |
| | ASV1483 | 4.63 | 3.02E-06 | Eukaryota | Ciliophora | Prostomatea | Prorodontida | Colepidae |
| | ASV1333 | 4.61 | 2.98E-14 | Fungi | Chytridiomycota | Chytridiomycetes | Rhizophydiales | Uebelmesseromycetaceae |
| | ASV2662 | 4.58 | 1.25E-08 | Eukaryota | Ciliophora | Oligotrichea | Oligotrichida | Strombidiidae |

| | | | | | | | | |
|---|---|---|---|---|---|---|---|---|
| | ASV632 | 4.55 | 1.21E-13 | Fungi | Chytridiomycota | Chytridiomycetes | Chytridiales | Chytriomycetaceae |
| | ASV702 | 4.55 | 1.84E-13 | Eukaryota | Myzozoa | Dinophyceae | Suessiales | Suessiaceae |
| | ASV2095 | 4.52 | 6.98E-05 | Eukaryota | Ciliophora | Spirotrichea | Tintinnida | Tintinnidae |
| | ASV1911 | 4.49 | 4.28E-09 | Fungi | Ascomycota | Leotiomycetes | Chaetomellales | Chaetomellaceae |
| | ASV1785 | 4.46 | 2.35E-12 | Eukaryota | Protozoa incertae sedis | Protozoa classis incertae sedis | Protozoa ordo incertae sedis | Alphamonaceae |
| | ASV2625 | 4.45 | 3.37E-11 | Eukaryota | incertae sedis | incertae sedis | Aquavolonida | incertae sedis |
| | ASV905 | 4.42 | 1.57E-13 | Fungi | Chytridiomycota | Chytridiomycetes | Chytridiales | Chytriomycetaceae |
| | ASV1835 | 4.40 | 3.68E-06 | Fungi | Blastocladiomycota | Blastocladiomycetes | Blastocladiales | Catenariaceae |
| | ASV2246 | 4.38 | 1.16E-07 | Eukaryota | Choanozoa | Ichthyosporea | Ichthyophonida | incertae sedis |
| | ASV396 | 4.34 | 7.43E-08 | Eukaryota | Choanozoa | Ichthyosporea | Ichthyophonida | incertae sedis |
| | ASV266 | 4.34 | 1.11E-10 | Eukaryota | Protozoa incertae sedis | Protozoa classis incertae sedis | Protozoa ordo incertae sedis | Alphamonaceae |
| | ASV41 | 4.32 | 1.72E-06 | Eukaryota | Bigyra | Bikosea | Bicosoecida | Bicosoecidae |
| | ASV9 | 4.31 | 2.60E-16 | Eukaryota | Myzozoa | Dinophyceae | Gymnodiniales | Polykrikaceae |
| | ASV2671 | 4.30 | 4.67E-08 | Eukaryota | Cryptophyta | Cryptophyceae | Cryptomonadales | Cryptomonadaceae |
| | ASV1527 | 4.30 | 2.77E-13 | Fungi | Chytridiomycota | Chytridiomycetes | Rhizophydiales | Uebelmesseromycetaceae |
| | ASV2428 | 4.29 | 2.54E-08 | Fungi | Ascomycota | Leotiomycetes | Chaetomellales | Chaetomellaceae |
| | ASV595 | 4.29 | 9.94E-06 | Fungi | Blastocladiomycota | Blastocladiomycetes | Blastocladiales | Catenariaceae |
| | ASV651 | 4.26 | 2.87E-11 | Eukaryota | Myzozoa | Dinophyceae | Suessiales | Suessiaceae |
| | ASV755 | 4.26 | 3.70E-09 | Eukaryota | Cryptophyta | Cryptophyceae | Cyathomonadacea | Goniomonadaceae |
| | ASV2812 | 4.24 | 5.47E-09 | Eukaryota | Cryptophyta | Cryptophyceae | Cyathomonadacea | Goniomonadaceae |
| | ASV1251 | 4.24 | 4.36E-12 | Eukaryota | Myzozoa | Dinophyceae | Suessiales | Suessiaceae |
| | ASV1003 | 4.23 | 8.90E-15 | Eukaryota | Myzozoa | Dinophyceae | Gymnodiniales | Polykrikaceae |
| | ASV790 | 4.23 | 1.62E-06 | Eukaryota | Bigyra | Bikosea | Bicosoecida | Bicosoecidae |
| | ASV2141 | 4.22 | 2.59E-09 | Fungi | Ascomycota | Leotiomycetes | Chaetomellales | Chaetomellaceae |
| | ASV198 | 4.22 | 1.56E-09 | Eukaryota | Chlorophyta | Chlorophyceae | Sphaeropleales | Sphaeropleaceae |
| | ASV1109 | 4.19 | 5.59E-06 | Fungi | Blastocladiomycota | Blastocladiomycetes | Blastocladiales | Catenariaceae |
| | ASV2160 | 4.16 | 9.22E-08 | Eukaryota | Choanozoa | Ichthyosporea | Ichthyophonida | incertae sedis |
| | ASV2528 | 4.15 | 1.34E-10 | Eukaryota | Bigyra | Bikosea | Bicosoecida | Siluaniidae |
| | ASV1712 | 4.12 | 4.09E-05 | Eukaryota | Ciliophora | Prostomatea | Prorodontida | Colepidae |
| | ASV2578 | 4.12 | 2.22E-12 | Eukaryota | Myzozoa | Dinophyceae | Suessiales | Suessiaceae |

| | | | | | | | | |
|---|---|---|---|---|---|---|---|---|
| | ASV2149 | 4.07 | 1.38E-12 | Eukaryota | Myzozoa | Dinophyceae | Suessiales | Suessiaceae |
| | ASV760 | 4.04 | 4.36E-06 | Fungi | Blastocladiomycota | Blastocladiomycetes | Blastocladiales | Catenariaceae |
| | ASV944 | 4.04 | 2.50E-10 | Eukaryota | incertae sedis | incertae sedis | Aquavolonida | incertae sedis |
| | ASV1401 | 4.03 | 9.65E-08 | Eukaryota | Cercozoa | Sarcomonadea | Glissomonadida | Viridiraptoridae |
| | ASV887 | 4.02 | 9.65E-15 | Eukaryota | Myzozoa | Dinophyceae | Gymnodiniales | Polykrikaceae |
| | ASV2684 | 4.01 | 4.99E-11 | Eukaryota | Myzozoa | Dinophyceae | Suessiales | Suessiaceae |
| | ASV2593 | 3.98 | 5.04E-09 | Eukaryota | Chlorophyta | Chlorophyceae | Sphaeropleales | Sphaeropleaceae |
| | ASV1330 | 3.96 | 5.30E-11 | Eukaryota | Myzozoa | Dinophyceae | Peridiniales | Podolampaceae |
| | ASV2208 | 3.92 | 4.51E-09 | Eukaryota | Chlorophyta | Chlorophyceae | Sphaeropleales | Sphaeropleaceae |
| | ASV3 | 3.89 | 5.28E-05 | Eukaryota | Myzozoa | Dinophyceae | Prorocentrales | Prorocentraceae |
| | ASV40 | 3.87 | 4.49E-12 | Eukaryota | Myzozoa | Dinophyceae | Prorocentrales | Prorocentraceae |
| | ASV2719 | 3.85 | 1.43E-08 | Eukaryota | Cercozoa | Sarcomonadea | Cercomonadida | Cercomonadidae |
| | ASV1283 | 3.83 | 4.36E-12 | Eukaryota | Ciliophora | Prostomatea | Prorodontida | Colepidae |
| | ASV2695 | 3.77 | 9.49E-05 | Eukaryota | Ciliophora | Prostomatea | Prorodontida | Colepidae |
| | ASV1637 | 3.77 | 1.23E-06 | Fungi | Blastocladiomycota | Blastocladiomycetes | Blastocladiales | Catenariaceae |
| | ASV2152 | 3.74 | 1.60E-07 | Eukaryota | Cercozoa | Sarcomonadea | Cercomonadida | Cercomonadidae |
| | ASV2549 | 3.73 | 9.84E-06 | Fungi | Blastocladiomycota | Blastocladiomycetes | Blastocladiales | Catenariaceae |
| | ASV1399 | 3.73 | 6.37E-07 | Eukaryota | Cercozoa | Sarcomonadea | Cercomonadida | Cercomonadidae |
| | ASV463 | 3.71 | 4.90E-07 | Fungi | Ascomycota | Leotiomycetes | Chaetomellales | Chaetomellaceae |
| | ASV1569 | 3.70 | 2.13E-14 | Eukaryota | Myzozoa | Dinophyceae | Gymnodiniales | Polykrikaceae |
| | ASV763 | 3.69 | 2.81E-10 | Eukaryota | Myzozoa | Dinophyceae | Suessiales | Suessiaceae |
| | ASV2255 | 3.68 | 9.66E-09 | Eukaryota | Chlorophyta | Chlorophyceae | Sphaeropleales | Sphaeropleaceae |
| | ASV1344 | 3.67 | 4.43E-05 | Eukaryota | Myzozoa | Dinophyceae | Prorocentrales | Prorocentraceae |
| | ASV1094 | 3.64 | 6.06E-09 | Eukaryota | Cercozoa | Sarcomonadea | Glissomonadida | Viridiraptoridae |
| | ASV1280 | 3.63 | 1.46E-07 | Eukaryota | Myzozoa | Dinophyceae | Amphidiniales | Amphidiniaceae |
| | ASV1095 | 3.63 | 9.26E-06 | Eukaryota | Ochrophyta | Chrysophyceae | Chrysosphaerales | Chrysosphaeraceae |
| | ASV1496 | 3.60 | 7.59E-06 | Eukaryota | Bigyra | Bikosea | Bicosoecida | Siluaniidae |
| | ASV1571 | 3.59 | 9.34E-10 | Eukaryota | Ciliophora | Prostomatea | Prorodontida | Colepidae |
| | ASV2384 | 3.57 | 1.34E-10 | Eukaryota | Cercozoa | Protozoa classis incertae sedis | Protozoa ordo incertae sedis | Protozoa familia incertae sedis |
| | ASV800 | 3.57 | 3.78E-09 | Eukaryota | Myzozoa | Dinophyceae | Peridiniales | Podolampaceae |

| | | | | | | | | |
|---|---|---|---|---|---|---|---|---|
| | ASV1697 | 3.57 | 1.82E-09 | Eukaryota | Myzozoa | Dinophyceae | Peridiniales | Podolampaceae |
| | ASV2765 | 3.57 | 1.25E-07 | Fungi | Chytridiomycota | Chytridiomycetes | Rhizophydiales | Uebelmesseromycetaceae |
| | ASV340 | 3.56 | 8.88E-05 | Eukaryota | Ciliophora | Prostomatea | Prorodontida | Colepidae |
| | ASV2567 | 3.54 | 1.53E-05 | Eukaryota | Myzozoa | Dinophyceae | Gymnodiniales | Gymnodiniaceae |
| | ASV202 | 3.53 | 0.000231 | Eukaryota | Myzozoa | Dinophyceae | Prorocentrales | Prorocentraceae |
| | ASV523 | 3.48 | 3.94E-09 | Eukaryota | Myzozoa | Dinophyceae | Suessiales | Suessiaceae |
| | ASV2250 | 3.47 | 1.94E-05 | Eukaryota | Cercozoa | Imbricatea | Spongomonadida | Spongomonadidae |
| | ASV1627 | 3.47 | 8.55E-09 | Eukaryota | incertae sedis | incertae sedis | Aquavolonida | incertae sedis |
| | ASV1159 | 3.47 | 5.30E-11 | Eukaryota | Myzozoa | Dinophyceae | Prorocentrales | Prorocentraceae |
| | ASV1585 | 3.47 | 5.00E-09 | Eukaryota | Myzozoa | Dinophyceae | Peridiniales | Podolampaceae |
| | ASV2076 | 3.43 | 0.000107 | Eukaryota | Myzozoa | Dinophyceae | Gymnodiniales | Gymnodiniaceae |
| | ASV1969 | 3.43 | 4.42E-08 | Eukaryota | Myzozoa | Dinophyceae | Peridiniales | Podolampaceae |
| | ASV1725 | 3.42 | 3.29E-05 | Eukaryota | Ciliophora | Prostomatea | Prorodontida | Colepidae |
| | ASV667 | 3.41 | 0.000345 | Eukaryota | Ciliophora | Prostomatea | Prorodontida | Colepidae |
| | ASV576 | 3.41 | 1.60E-07 | Eukaryota | Cercozoa | Sarcomonadea | Cercomonadida | Cercomonadidae |
| | ASV1873 | 3.40 | 1.64E-06 | Fungi | Chytridiomycota | Chytridiomycetes | Rhizophydiales | Uebelmesseromycetaceae |
| | ASV2716 | 3.40 | 4.19E-07 | Eukaryota | Ciliophora | Prostomatea | Prorodontida | Colepidae |
| | ASV2261 | 3.40 | 3.10E-06 | Eukaryota | Ciliophora | Prostomatea | Prorodontida | Colepidae |
| | ASV1196 | 3.39 | 0.000311 | Eukaryota | Myzozoa | Dinophyceae | Peridiniales | Peridiniaceae |
| | ASV563 | 3.37 | 0.000508 | Eukaryota | Myzozoa | Dinophyceae | Prorocentrales | Prorocentraceae |
| | ASV151 | 3.37 | 7.00E-06 | Eukaryota | Choanozoa | Choanoflagellata | Craspedida | Codonosigaceae |
| | ASV2507 | 3.35 | 1.50E-06 | Eukaryota | Ciliophora | Prostomatea | Prorodontida | Colepidae |
| | ASV2485 | 3.34 | 3.42E-06 | Eukaryota | Cercozoa | Sarcomonadea | Cercomonadida | Cercomonadidae |
| | ASV1250 | 3.33 | 8.57E-05 | Fungi | Blastocladiomycota | Blastocladiomycetes | Blastocladiales | Catenariaceae |
| | ASV526 | 3.32 | 2.78E-07 | Eukaryota | Myzozoa | Dinophyceae | Amphidiniales | Amphidiniaceae |
| | ASV1427 | 3.30 | 2.99E-07 | Fungi | Ascomycota | Leotiomycetes | Chaetomellales | Chaetomellaceae |
| | ASV451 | 3.28 | 1.97E-05 | Eukaryota | Choanozoa | Ichthyosporea | Ichthyophonida | incertae sedis |
| | ASV2591 | 3.28 | 0.000134 | Eukaryota | Ciliophora | Prostomatea | Prorodontida | Colepidae |
| | ASV2664 | 3.27 | 1.65E-05 | Fungi | Blastocladiomycota | Blastocladiomycetes | Blastocladiales | Catenariaceae |
| | ASV948 | 3.26 | 2.79E-10 | Eukaryota | Ochrophyta | Eustigmatophyceae | Goniochloridales | Goniochloridaceae |

| | | | | | | | | |
|---|---|---|---|---|---|---|---|---|
| | ASV92 | 3.24 | 3.83E-05 | Eukaryota | Ochrophyta | Chrysophyceae | Chrysosphaerales | Chrysosphaeraceae |
| | ASV750 | 3.22 | 0.000116 | Eukaryota | Bigyra | Bikosea | Bicosoecida | Siluaniidae |
| | ASV1319 | 3.22 | 9.43E-05 | Eukaryota | Ciliophora | Prostomatea | Prorodontida | Colepidae |
| | ASV1549 | 3.21 | 0.001091 | Eukaryota | Myzozoa | Dinophyceae | Peridiniales | Peridiniaceae |
| | ASV1541 | 3.21 | 6.29E-08 | Eukaryota | Myzozoa | Dinophyceae | Amphidiniales | Amphidiniaceae |
| | ASV1783 | 3.19 | 5.62E-05 | Eukaryota | Choanozoa | Choanoflagellata | Craspedida | Codonosigaceae |
| | ASV1 | 3.19 | 4.54E-11 | Eukaryota | Choanozoa | Choanoflagellata | Craspedida | Codonosigaceae |
| | ASV141 | 3.18 | 0.000945 | Eukaryota | Ciliophora | Prostomatea | Prorodontida | Colepidae |
| | ASV1563 | 3.18 | 5.58E-10 | Eukaryota | Ciliophora | Oligohymenophorea | Astomatida | Haptophryidae |
| | ASV2179 | 3.16 | 3.13E-06 | Fungi | Chytridiomycota | Chytridiomycetes | Rhizophydiales | Uebelmesseromycetaceae |
| | ASV1803 | 3.16 | 2.71E-05 | Eukaryota | Ciliophora | Prostomatea | Prorodontida | Colepidae |
| | ASV1061 | 3.15 | 4.87E-07 | Eukaryota | Ochrophyta | Chrysophyceae | Ochromonadales | Ochromonadaceae |
| | ASV2394 | 3.15 | 0.000207 | Eukaryota | Ciliophora | Prostomatea | Prorodontida | Colepidae |
| | ASV1788 | 3.14 | 0.000104 | Eukaryota | Myzozoa | Dinophyceae | Gymnodiniales | Gymnodiniaceae |
| | ASV761 | 3.13 | 0.000188 | Eukaryota | Myzozoa | Dinophyceae | Gymnodiniales | Gymnodiniaceae |
| | ASV2294 | 3.12 | 0.000188 | Eukaryota | Myzozoa | Dinophyceae | Peridiniales | Podolampaceae |
| | ASV2033 | 3.10 | 1.97E-05 | Eukaryota | Myzozoa | Dinophyceae | Peridiniales | Podolampaceae |
| | ASV1904 | 3.09 | 3.30E-10 | Eukaryota | Ciliophora | Oligohymenophorea | Astomatida | Haptophryidae |
| | ASV2233 | 3.05 | 0.000143 | Eukaryota | Myzozoa | Dinophyceae | Peridiniales | Podolampaceae |
| | ASV85 | 3.05 | 0.002877 | Eukaryota | Myzozoa | Dinophyceae | Peridiniales | Peridiniaceae |
| | ASV1531 | 3.05 | 9.49E-05 | Eukaryota | Ochrophyta | Chrysophyceae | Synurales | Mallomonadaceae |
| | ASV1917 | 3.04 | 2.05E-06 | Eukaryota | Myzozoa | Dinophyceae | Peridiniales | Podolampaceae |
| | ASV792 | 3.03 | 0.006972 | Eukaryota | Cercozoa | Sarcomonadea | Cercomonadida | Cercomonadidae |
| | ASV2521 | 3.02 | 0.001749 | Eukaryota | Ciliophora | Spirotrichea | Urostylida | Holostichidae |
| | ASV1881 | 3.02 | 0.003301 | Eukaryota | Ciliophora | Spirotrichea | Urostylida | Holostichidae |
| | ASV2511 | 3.01 | 2.23E-10 | Eukaryota | Cercozoa | Thecofilosea | Ebriida | Ebriidae |
| | ASV462 | 3.01 | 0.000122 | Eukaryota | Ochrophyta | Chrysophyceae | Synurales | Mallomonadaceae |
| | ASV2424 | 3.01 | 0.000103 | Eukaryota | Myzozoa | Dinophyceae | Peridiniales | Podolampaceae |
| | ASV469 | 3.00 | 0.00121 | Eukaryota | Bigyra | Bikosea | Bicosoecida | Siluaniidae |
| | ASV1005 | 3.00 | 0.000976 | Eukaryota | Ciliophora | Prostomatea | Prorodontida | Colepidae |

| | | | | | | | | |
|---|---|---|---|---|---|---|---|---|
| | ASV1894 | 3.00 | 1.94E-07 | Fungi | Ascomycota | Dothideomycetes | Venturiales | Sympoventuriaceae |
| | ASV2374 | 3.00 | 7.59E-06 | Eukaryota | Cercozoa | Imbricatea | Spongomonadida | Spongomonadidae |
| | ASV64 | 2.96 | 8.09E-05 | Eukaryota | Choanozoa | Choanoflagellata | Craspedida | Codonosigaceae |
| | ASV561 | 2.94 | 2.88E-07 | Eukaryota | Choanozoa | Choanoflagellata | Craspedida | Codonosigaceae |
| | ASV2752 | 2.94 | 6.20E-07 | Eukaryota | Ochrophyta | Chrysophyceae | Paraphysomonadales | Paraphysomonadaceae |
| | ASV155 | 2.94 | 0.000657 | Eukaryota | Ciliophora | Prostomatea | Prorodontida | Colepidae |
| | ASV2723 | 2.93 | 4.47E-06 | Eukaryota | Ciliophora | Spirotrichea | Sporadotrichida | Oxytrichidae |
| | ASV2456 | 2.91 | 6.88E-05 | Eukaryota | Bigyra | Bikosea | Bicosoecida | Siluaniidae |
| | ASV1716 | 2.90 | 2.32E-09 | Eukaryota | Myzozoa | Dinophyceae | Suessiales | Suessiaceae |
| | ASV898 | 2.89 | 4.89E-06 | Eukaryota | Ciliophora | Spirotrichea | Sporadotrichida | Oxytrichidae |
| | ASV2624 | 2.88 | 0.000466 | Eukaryota | Bigyra | Bikosea | Bicosoecida | Siluaniidae |
| | ASV1626 | 2.87 | 1.14E-06 | Eukaryota | Choanozoa | Choanoflagellata | Craspedida | Codonosigaceae |
| | ASV488 | 2.86 | 7.41E-07 | Eukaryota | Bigyra | Bikosea | Bicosoecida | Siluaniidae |
| | ASV359 | 2.85 | 3.22E-08 | Eukaryota | Bigyra | Bikosea | Bicosoecida | Bicosoecidae |
| | ASV562 | 2.83 | 1.36E-05 | Fungi | Chytridiomycota | Chytridiomycetes | Rhizophydiales | Uebelmesseromycetaceae |
| | ASV2576 | 2.83 | 1.87E-10 | Eukaryota | Myzozoa | Dinophyceae | Suessiales | Suessiaceae |
| | ASV1872 | 2.82 | 6.38E-06 | Eukaryota | incertae sedis | Developea | incertae sedis | Developayellaceae |
| | ASV2287 | 2.82 | 0.002613 | Eukaryota | Ciliophora | Spirotrichea | Urostylida | Holostichidae |
| | ASV257 | 2.81 | 0.00091 | Eukaryota | Ciliophora | Litostomatea | Haptorida | Dimacrocaryonidae |
| | ASV76 | 2.80 | 3.67E-09 | Eukaryota | Myzozoa | Dinophyceae | Prorocentrales | Prorocentraceae |
| | ASV608 | 2.79 | 0.000147 | Eukaryota | Ciliophora | Prostomatea | Prorodontida | Colepidae |
| | ASV391 | 2.78 | 1.90E-10 | Eukaryota | Myzozoa | Dinophyceae | Suessiales | Suessiaceae |
| | ASV2220 | 2.77 | 0.001003 | Eukaryota | Ciliophora | Litostomatea | Haptorida | Dimacrocaryonidae |
| | ASV382 | 2.77 | 5.42E-06 | Eukaryota | Bigyra | Bikosea | Bicosoecida | Siluaniidae |
| | ASV951 | 2.77 | 0.001358 | Eukaryota | Chlorophyta | Chlorophyceae | Chlamydomonadales | Chlamydomonadaceae |
| | ASV439 | 2.76 | 1.84E-06 | Eukaryota | Ciliophora | Prostomatea | Prorodontida | Colepidae |
| | ASV2048 | 2.76 | 0.00024 | Eukaryota | Ochrophyta | Dictyochophyceae | Pedinellales | Actinomonadaceae |
| | ASV2834 | 2.76 | 0.00868 | Eukaryota | Myzozoa | Dinophyceae | Peridiniales | Peridiniaceae |
| | ASV968 | 2.75 | 2.91E-11 | Eukaryota | Myzozoa | Dinophyceae | Suessiales | Suessiaceae |
| | ASV690 | 2.75 | 5.46E-07 | Eukaryota | Bigyra | Bikosea | Bicosoecida | Siluaniidae |

| | | | | | | | | |
|---|---|---|---|---|---|---|---|---|
| | ASV2523 | 2.75 | 5.16E-05 | Eukaryota | Ochrophyta | Chrysophyceae | Ochromonadales | Ochromonadaceae |
| | ASV411 | 2.75 | 3.27E-06 | Eukaryota | Choanozoa | Choanoflagellata | Craspedida | Codonosigaceae |
| | ASV1389 | 2.74 | 0.008066 | Eukaryota | Ciliophora | Spirotrichea | Urostylida | Holostichidae |
| | ASV2413 | 2.74 | 1.78E-07 | Eukaryota | Chlorophyta | Trebouxiophyceae | Chlorellales | Chlorellaceae |
| | ASV2197 | 2.72 | 0.000134 | Eukaryota | Cryptophyta | Cryptophyceae | Cyathomonadacea | Goniomonadaceae |
| | ASV1940 | 2.70 | 0.000127 | Eukaryota | Ciliophora | Prostomatea | Prorodontida | Colepidae |
| | ASV1043 | 2.69 | 9.80E-08 | Eukaryota | Ochrophyta | Chrysophyceae | Chromulinales | Chrysoamoebidaceae |
| | ASV189 | 2.69 | 9.96E-06 | Eukaryota | Bigyra | Bikosea | Bicosoecida | Siluaniidae |
| | ASV1221 | 2.66 | 1.29E-05 | Eukaryota | Ciliophora | Spirotrichea | Sporadotrichida | Oxytrichidae |
| | ASV2038 | 2.66 | 0.001311 | Eukaryota | Bigyra | Bikosea | Bicosoecida | Siluaniidae |
| | ASV2854 | 2.65 | 2.73E-08 | Eukaryota | Bigyra | Bikosea | Bicosoecida | Siluaniidae |
| | ASV298 | 2.65 | 0.000135 | Eukaryota | incertae sedis | incertae sedis | Aquavolonida | incertae sedis |
| | ASV1181 | 2.64 | 1.40E-06 | Eukaryota | Ochrophyta | Dictyochophyceae | Pedinellales | Actinomonadaceae |
| | ASV196 | 2.64 | 0.000646 | Eukaryota | Cryptophyta | Cryptophyceae | Cryptomonadales | Cryptomonadaceae |
| | ASV1587 | 2.62 | 9.17E-07 | Eukaryota | Bigyra | Labyrinthulea | Thraustochytrida | Thraustochytriaceae |
| | ASV1062 | 2.62 | 3.04E-06 | Eukaryota | incertae sedis | Developea | incertae sedis | Developayellaceae |
| | ASV1242 | 2.62 | 1.11E-09 | Eukaryota | Chlorophyta | Chlorophyceae | Sphaeropleales | Sphaeropleaceae |
| | ASV351 | 2.61 | 8.38E-07 | Eukaryota | Bigyra | Bikosea | Bicosoecida | Siluaniidae |
| | ASV17 | 2.60 | 0.000249 | Eukaryota | Bigyra | Bikosea | Bicosoecida | Siluaniidae |
| | ASV1118 | 2.59 | 2.01E-06 | Eukaryota | Cryptophyta | Cryptophyceae | Cyathomonadacea | Goniomonadaceae |
| | ASV112 | 2.59 | 0.001008 | Eukaryota | Cryptophyta | Cryptophyceae | Cryptomonadales | Cryptomonadaceae |
| | ASV1525 | 2.59 | 0.000633 | Eukaryota | Cryptophyta | Cryptophyceae | Cryptomonadales | Cryptomonadaceae |
| | ASV736 | 2.59 | 4.08E-05 | Eukaryota | Oomycota | Hyphochytrea | Pirsoniales | Pirsoniaceae |
| | ASV662 | 2.57 | 6.73E-08 | Eukaryota | incertae sedis | incertae sedis | Aquavolonida | incertae sedis |
| | ASV291 | 2.57 | 5.35E-06 | Eukaryota | Ciliophora | Spirotrichea | Sporadotrichida | Oxytrichidae |
| | ASV1730 | 2.55 | 0.000887 | Eukaryota | Myzozoa | Dinophyceae | Gymnodiniales | Gymnodiniaceae |
| | ASV2526 | 2.54 | 0.007645 | Eukaryota | Bigyra | Bikosea | Bicosoecida | Siluaniidae |
| | ASV1028 | 2.53 | 0.000646 | Eukaryota | Chlorophyta | Chlorophyceae | Chlamydomonadales | Chlamydomonadaceae |
| | ASV820 | 2.53 | 0.000939 | Eukaryota | Cryptophyta | Cryptophyceae | Cyathomonadacea | Goniomonadaceae |
| | ASV626 | 2.53 | 1.71E-07 | Eukaryota | Ochrophyta | Dictyochophyceae | Pedinellales | Actinomonadaceae |

| | | | | | | | | |
|---|---|---|---|---|---|---|---|---|
| | ASV953 | 2.53 | 2.39E-07 | Eukaryota | Cercozoa | Sarcomonadea | Cercomonadida | Cercomonadidae |
| | ASV2041 | 2.52 | 0.002063 | Eukaryota | Cryptophyta | Cryptophyceae | Cryptomonadales | Cryptomonadaceae |
| | ASV1840 | 2.52 | 1.09E-09 | Eukaryota | Ciliophora | Litostomatea | Haptorida | Spathidiidae |
| | ASV1789 | 2.51 | 1.40E-06 | Eukaryota | Chlorophyta | Mamiellophyceae | Dolichomastigales | Crustomastigaceae |
| | ASV1313 | 2.50 | 1.23E-05 | Eukaryota | Choanozoa | Choanoflagellata | Craspedida | Codonosigaceae |
| | ASV2562 | 2.50 | 2.15E-05 | Eukaryota | Bigyra | Bikosea | Bicosoecida | Siluaniidae |
| | ASV2640 | 2.49 | 3.75E-06 | Eukaryota | Ochrophyta | Dictyochophyceae | Pedinellales | Actinomonadaceae |
| | ASV113 | 2.49 | 0.001575 | Eukaryota | Chlorophyta | Chlorophyceae | Chlamydomonadales | Characiochloridaceae |
| | ASV7 | 2.49 | 0.00809 | Eukaryota | Cercozoa | Sarcomonadea | Cercomonadida | Cercomonadidae |
| | ASV1343 | 2.48 | 0.003157 | Eukaryota | Choanozoa | Choanoflagellata | Craspedida | Codonosigaceae |
| | ASV1779 | 2.48 | 0.001049 | Eukaryota | Cryptophyta | Cryptophyceae | Cyathomonadacea | Goniomonadaceae |
| | ASV1852 | 2.48 | 0.000113 | Eukaryota | Cryptophyta | Cryptophyceae | Cyathomonadacea | Goniomonadaceae |
| | ASV402 | 2.46 | 0.002367 | Eukaryota | Ciliophora | Spirotrichea | Urostylida | Holostichidae |
| | ASV56 | 2.46 | 0.000204 | Eukaryota | Cryptophyta | Cryptophyceae | Cryptomonadales | Cryptomonadaceae |
| | ASV1574 | 2.45 | 0.002348 | Eukaryota | Myzozoa | Dinophyceae | Gymnodiniales | Gymnodiniaceae |
| | ASV2370 | 2.45 | 0.00045 | Eukaryota | Cryptophyta | Cryptophyceae | Cyathomonadacea | Goniomonadaceae |
| | ASV540 | 2.44 | 0.002281 | Eukaryota | Chlorophyta | Chlorophyceae | Chlamydomonadales | Characiochloridaceae |
| | ASV14 | 2.44 | 0.005602 | Eukaryota | Chlorophyta | Chlorophyceae | Chlamydomonadales | Chlamydomonadaceae |
| | ASV2764 | 2.44 | 7.48E-09 | Eukaryota | Bigyra | Bikosea | Bicosoecida | Siluaniidae |
| | ASV2577 | 2.42 | 0.002978 | Eukaryota | Ciliophora | Litostomatea | Haptorida | Dimacrocaryonidae |
| | ASV2266 | 2.42 | 0.001736 | Eukaryota | Cryptophyta | Cryptophyceae | Cryptomonadales | Cryptomonadaceae |
| | ASV2348 | 2.41 | 0.00121 | Eukaryota | Myzozoa | Dinophyceae | Peridiniales | Podolampaceae |
| | ASV1018 | 2.41 | 5.12E-05 | Eukaryota | Chlorophyta | Mamiellophyceae | Dolichomastigales | Crustomastigaceae |
| | ASV1654 | 2.40 | 0.003157 | Eukaryota | Ciliophora | Prostomatea | Prorodontida | Colepidae |
| | ASV696 | 2.40 | 8.35E-07 | Eukaryota | Cryptophyta | Cryptophyceae | Cryptomonadales | Cryptomonadaceae |
| | ASV394 | 2.39 | 7.98E-05 | Eukaryota | Myzozoa | Dinophyceae | Prorocentrales | Prorocentraceae |
| | ASV2338 | 2.39 | 7.42E-07 | Eukaryota | Choanozoa | Choanoflagellata | Craspedida | Codonosigaceae |
| | ASV1025 | 2.39 | 0.001784 | Eukaryota | Ciliophora | Litostomatea | Haptorida | Dimacrocaryonidae |
| | ASV1632 | 2.39 | 0.002198 | Eukaryota | Myzozoa | Dinophyceae | Peridiniales | Podolampaceae |
| | ASV325 | 2.38 | 0.001484 | Eukaryota | Ciliophora | Prostomatea | Prorodontida | Colepidae |

| | | | | | | | | |
|---|---|---|---|---|---|---|---|---|
| | ASV1093 | 2.38 | 0.006253 | Eukaryota | Ochrophyta | Chrysophyceae | Ochromonadales | Ochromonadaceae |
| | ASV872 | 2.37 | 2.79E-07 | Eukaryota | Cercozoa | Sarcomonadea | Cercomonadida | Cercomonadidae |
| | ASV2215 | 2.35 | 0.007829 | Eukaryota | Ochrophyta | Chrysophyceae | Chromulinales | Chrysoamoebidaceae |
| | ASV635 | 2.35 | 0.000935 | Eukaryota | Myzozoa | Dinophyceae | Gymnodiniales | Gymnodiniaceae |
| | ASV2545 | 2.34 | 9.98E-05 | Eukaryota | Ochrophyta | Chrysophyceae | Synurales | Mallomonadaceae |
| | ASV1787 | 2.34 | 0.00122 | Eukaryota | Oomycota | Peronosporea | Saprolegniales | Saprolegniaceae |
| | ASV89 | 2.32 | 8.09E-05 | Eukaryota | Myzozoa | Dinophyceae | Gonyaulacales | Ceratiaceae |
| | ASV1601 | 2.32 | 0.009703 | Eukaryota | Ciliophora | Spirotrichea | Choreotrichida | Strombidinopsidae |
| | ASV1278 | 2.31 | 0.00061 | Eukaryota | Myzozoa | Dinophyceae | Gonyaulacales | Ceratiaceae |
| | ASV1534 | 2.30 | 2.92E-06 | Eukaryota | Bigyra | Bikosea | Bicosoecida | Siluaniidae |
| | ASV729 | 2.29 | 4.99E-05 | Fungi | Chytridiomycota | Chytridiomycetes | Rhizophydiales | Uebelmesseromycetaceae |
| | ASV1864 | 2.28 | 0.00226 | Eukaryota | Bigyra | Bikosea | Bicosoecida | Siluaniidae |
| | ASV1883 | 2.28 | 0.000127 | Eukaryota | Ochrophyta | Dictyochophyceae | Pedinellales | Actinomonadaceae |
| | ASV1539 | 2.27 | 0.00571 | Eukaryota | Ciliophora | Spirotrichea | Urostylida | Holostichidae |
| | ASV2025 | 2.27 | 0.006504 | Eukaryota | Chlorophyta | Chlorophyceae | Chlamydomonadales | Chlamydomonadaceae |
| | ASV1919 | 2.27 | 0.007652 | Eukaryota | Ochrophyta | Chrysophyceae | Chromulinales | Chrysoamoebidaceae |
| | ASV442 | 2.26 | 7.63E-05 | Eukaryota | Ochrophyta | Chrysophyceae | Synurales | Mallomonadaceae |
| | ASV655 | 2.26 | 0.000866 | Eukaryota | Oomycota | Peronosporea | Saprolegniales | Saprolegniaceae |
| | ASV1834 | 2.26 | 0.002613 | Eukaryota | Cryptophyta | Cryptophyceae | Cyathomonadacea | Goniomonadaceae |
| | ASV1739 | 2.25 | 0.000466 | Fungi | Cryptomycota | Cryptomycota classis incertae sedis | Cryptomycota ordo incertae sedis | Cryptomycota familia incertae sedis |
| | ASV1841 | 2.24 | 0.001341 | Eukaryota | Myzozoa | Dinophyceae | Peridiniales | Podolampaceae |
| | ASV1286 | 2.24 | 7.23E-05 | Eukaryota | Bigyra | Bikosea | Bicosoecida | Siluaniidae |
| | ASV1946 | 2.23 | 0.001966 | Eukaryota | Cryptophyta | Cryptophyceae | Cyathomonadacea | Goniomonadaceae |
| | ASV179 | 2.23 | 0.006298 | Eukaryota | Charophyta | Zygnemophyceae | Desmidiales | Closteriaceae |
| | ASV920 | 2.22 | 2.05E-06 | Eukaryota | Bigyra | Bikosea | Bicosoecida | Bicosoecidae |
| | ASV2216 | 2.21 | 9.12E-06 | Eukaryota | Bigyra | Bikosea | Bicosoecida | Bicosoecidae |
| | ASV1463 | 2.20 | 1.97E-05 | Eukaryota | Cryptophyta | Cryptophyceae | Cyathomonadacea | Goniomonadaceae |
| | ASV2835 | 2.19 | 0.000103 | Eukaryota | Ochrophyta | Chrysophyceae | Paraphysomonadales | Paraphysomonadaceae |
| | ASV1829 | 2.18 | 2.52E-05 | Eukaryota | Myzozoa | Dinophyceae | Amphidiniales | Amphidiniaceae |

| | | | | | | | | |
|---|---|---|---|---|---|---|---|---|
| | ASV2552 | 2.18 | 0.005469 | Eukaryota | Chlorophyta | Chlorophyceae | Chlamydomonadales | Characiochloridaceae |
| | ASV2733 | 2.18 | 0.003866 | Eukaryota | Charophyta | Zygnemophyceae | Desmidiales | Closteriaceae |
| | ASV1812 | 2.18 | 0.000124 | Eukaryota | Cryptophyta | Cryptophyceae | Cyathomonadacea | Goniomonadaceae |
| | ASV1123 | 2.17 | 0.000516 | Eukaryota | Chlorophyta | Chlorophyceae | Chlamydomonadales | Chlamydomonadaceae |
| | ASV1850 | 2.17 | 0.000134 | Eukaryota | Ochrophyta | Chrysophyceae | Synurales | Mallomonadaceae |
| | ASV365 | 2.17 | 0.001103 | Fungi | Cryptomycota | Cryptomycota classis incertae sedis | Cryptomycota ordo incertae sedis | Cryptomycota familia incertae sedis |
| | ASV2534 | 2.16 | 0.002229 | Eukaryota | Ciliophora | Spirotrichea | Urostylida | Holostichidae |
| | ASV1439 | 2.16 | 0.000127 | Eukaryota | Ochrophyta | Chrysophyceae | Ochromonadales | Ochromonadaceae |
| | ASV260 | 2.15 | 3.72E-05 | Eukaryota | Ochrophyta | Chrysophyceae | Ochromonadales | Ochromonadaceae |
| | ASV2323 | 2.14 | 5.61E-06 | Eukaryota | incertae sedis | incertae sedis | Aquavolonida | incertae sedis |
| | ASV2188 | 2.14 | 0.001313 | Eukaryota | Bigyra | Bikosea | Bicosoecida | Siluaniidae |
| | ASV424 | 2.13 | 1.25E-05 | Eukaryota | Myzozoa | Dinophyceae | Prorocentrales | Prorocentraceae |
| | ASV4 | 2.13 | 0.006395 | Eukaryota | Myzozoa | Dinophyceae | Gonyaulacales | Ceratiaceae |
| | ASV146 | 2.12 | 7.59E-06 | Fungi | Chytridiomycota | Chytridiomycetes | Chytridiales | Chytriomycetaceae |
| | ASV1550 | 2.12 | 0.005049 | Eukaryota | Ciliophora | Prostomatea | Prorodontida | Colepidae |
| | ASV1083 | 2.11 | 0.003249 | Eukaryota | Chlorophyta | Chlorophyceae | Chlamydomonadales | Characiochloridaceae |
| | ASV2470 | 2.11 | 5.03E-05 | Eukaryota | Myzozoa | Dinophyceae | Thoracosphaerales | Thoracosphaeraceae |
| | ASV2391 | 2.11 | 0.000602 | Eukaryota | Ochrophyta | Chrysophyceae | Chrysosphaerales | Chrysosphaeraceae |
| | ASV1358 | 2.11 | 0.002256 | Eukaryota | Chlorophyta | Trebouxiophyceae | Chlorellales | Chlorellaceae |
| | ASV1766 | 2.10 | 1.52E-05 | Eukaryota | Ochrophyta | Chrysophyceae | Ochromonadales | Ochromonadaceae |
| | ASV186 | 2.09 | 0.000127 | Eukaryota | Ochrophyta | Chrysophyceae | Synurales | Mallomonadaceae |
| | ASV1186 | 2.07 | 0.000111 | Eukaryota | Chlorophyta | Mamiellophyceae | Dolichomastigales | Crustomastigaceae |
| | ASV61 | 2.07 | 0.000923 | Eukaryota | Ochrophyta | Chrysophyceae | Chromulinales | Paraphysomonadaceae |
| | ASV1266 | 2.07 | 0.00067 | Eukaryota | Ochrophyta | Dictyochophyceae | Pedinellales | Actinomonadaceae |
| | ASV140 | 2.06 | 0.000267 | Eukaryota | Ciliophora | Spirotrichea | Tintinnida | Codonellidae |
| | ASV271 | 2.04 | 0.00554 | Eukaryota | Ochrophyta | Chrysophyceae | Chromulinales | Chrysoamoebidaceae |
| | ASV936 | 2.03 | 0.000862 | Eukaryota | Bigyra | Bikosea | Bicosoecida | Siluaniidae |
| | ASV37 | 2.03 | 0.000665 | Eukaryota | Ochrophyta | Chrysophyceae | Ochromonadales | Ochromonadaceae |
| | ASV881 | 2.03 | 0.002367 | Eukaryota | Ciliophora | Prostomatea | Prorodontida | Colepidae |

| | | | | | | | | |
|---|---|---|---|---|---|---|---|---|
| | ASV764 | 2.02 | 0.005683 | Eukaryota | Ochrophyta | Chrysophyceae | Chromulinales | Chrysoamoebidaceae |
| | ASV1741 | 2.02 | 0.000286 | Eukaryota | Ochrophyta | Chrysophyceae | Ochromonadales | Ochromonadaceae |
| | ASV2689 | 2.02 | 0.000164 | Eukaryota | Ciliophora | Prostomatea | Prorodontida | Colepidae |
| | ASV333 | 2.01 | 0.001735 | Eukaryota | Ciliophora | Prostomatea | Prorodontida | Colepidae |
| | ASV1216 | 2.01 | 1.14E-06 | Eukaryota | Cercozoa | Sarcomonadea | Cercomonadida | Cercomonadidae |
| | ASV25 | 2.00 | 0.002687 | Eukaryota | Ochrophyta | Chrysophyceae | Ochromonadales | Ochromonadaceae |
| | ASV1804 | 2.00 | 0.0001 | Eukaryota | Ochrophyta | Chrysophyceae | Synurales | Mallomonadaceae |
| 16S-PP | ASV538 | 6.90 | 1.20E-15 | Bacteria | Bacteroidota | Bacteroidia | Chitinophagales | bacI |
| | ASV862 | 6.64 | 8.63E-20 | Bacteria | Proteobacteria | Gammaproteobacteria | Gammaproteobacteria_Incertae_Sedis | Unknown_Family |
| | ASV1127 | 6.61 | 6.08E-20 | Bacteria | Proteobacteria | Gammaproteobacteria | Gammaproteobacteria_Incertae_Sedis | Unknown_Family |
| | ASV1308 | 6.40 | 2.69E-17 | Bacteria | Bacteroidota | Bacteroidia | Chitinophagales | bacI |
| | ASV456 | 6.35 | 9.59E-15 | Bacteria | Bacteroidota | Bacteroidia | Chitinophagales | Chitinophagaceae |
| | ASV362 | 6.19 | 1.32E-15 | Bacteria | Bacteroidota | Bacteroidia | Flavobacteriales | bacV |
| | ASV682 | 6.15 | 9.43E-13 | Bacteria | Proteobacteria | Alphaproteobacteria | Acetobacterales | alfVIII |
| | ASV1380 | 6.10 | 9.67E-12 | Bacteria | Proteobacteria | Gammaproteobacteria | Burkholderiales | betI |
| | ASV1449 | 5.92 | 3.42E-20 | Bacteria | Proteobacteria | Gammaproteobacteria | Gammaproteobacteria_Incertae_Sedis | Unknown_Family |
| | ASV1457 | 5.91 | 2.39E-18 | Bacteria | Actinobacteriota | Actinobacteria | Frankiales | acI |
| | ASV493 | 5.82 | 9.59E-15 | Bacteria | Bacteroidota | Bacteroidia | Chitinophagales | bacI |
| | ASV1481 | 5.79 | 1.52E-14 | Bacteria | Proteobacteria | Gammaproteobacteria | Burkholderiales | Rhodocyclaceae |
| | ASV13 | 5.75 | 1.35E-12 | Bacteria | Actinobacteriota | Acidimicrobiia | Microtrichales | acIV |
| | ASV34 | 5.70 | 6.44E-10 | Bacteria | Bacteroidota | Kapabacteria | Kapabacteriales | Kapabacteriales |
| | ASV839 | 5.62 | 9.35E-11 | Bacteria | Acidobacteriota | Acidobacteriae | Acidobacteriae | Acidobacteriae |
| | ASV938 | 5.30 | 1.03E-09 | Bacteria | Bacteroidota | Bacteroidia | Chitinophagales | uncultured |
| | ASV1094 | 5.30 | 2.50E-18 | Bacteria | Actinobacteriota | Acidimicrobiia | Microtrichales | acIV |
| | ASV1077 | 5.26 | 2.45E-15 | Bacteria | Proteobacteria | Gammaproteobacteria | Burkholderiales | Rhodocyclaceae |
| | ASV481 | 5.24 | 2.41E-10 | Bacteria | Bacteroidota | Bacteroidia | Flavobacteriales | Crocinitomicaceae |
| | ASV1337 | 5.21 | 3.35E-16 | Bacteria | Bacteroidota | Bacteroidia | Chitinophagales | Chitinophagaceae |
| | ASV1352 | 5.15 | 1.42E-17 | Bacteria | Bacteroidota | Bacteroidia | Flavobacteriales | bacV |
| | ASV1355 | 5.05 | 1.22E-16 | Bacteria | Proteobacteria | Gammaproteobacteria | Burkholderiales | Rhodocyclaceae |
| | ASV1459 | 5.05 | 1.81E-15 | Bacteria | Proteobacteria | Gammaproteobacteria | Burkholderiales | Rhodocyclaceae |

| | | | | | | | | |
|---|---|---|---|---|---|---|---|---|
| | ASV1139 | 4.98 | 1.62E-10 | Bacteria | Bacteroidota | Bacteroidia | Flavobacteriales | Cryomorphaceae |
| | ASV1219 | 4.93 | 2.45E-12 | Bacteria | Bacteroidota | Bacteroidia | Flavobacteriales | Crocinitomicaceae |
| | ASV1366 | 4.68 | 1.51E-14 | Bacteria | Proteobacteria | Gammaproteobacteria | Burkholderiales | Rhodocyclaceae |
| | ASV407 | 4.64 | 3.42E-20 | Bacteria | Proteobacteria | Alphaproteobacteria | Rhizobiales | alfI |
| | ASV1245 | 4.63 | 3.72E-12 | Bacteria | Proteobacteria | Gammaproteobacteria | Burkholderiales | Rhodocyclaceae |
| | ASV1388 | 4.62 | 4.25E-14 | Bacteria | Bacteroidota | Bacteroidia | Sphingobacteriales | NS11-12_marine_group |
| | ASV517 | 4.59 | 7.18E-10 | Bacteria | SAR324_clade(Marine_group_B) | SAR324_clade(Marine_group_B) | SAR324_clade(Marine_group_B) | SAR324_clade(Marine_group_B) |
| | ASV507 | 4.59 | 1.43E-09 | Bacteria | Bacteroidota | Bacteroidia | Chitinophagales | Chitinophagaceae |
| | ASV3 | 4.58 | 1.04E-13 | Bacteria | Actinobacteriota | Actinobacteria | Micrococcales | Luna1 |
| | ASV1383 | 4.53 | 2.30E-15 | Bacteria | Bacteroidota | Bacteroidia | Sphingobacteriales | env.OPS_17 |
| | ASV1159 | 4.53 | 5.50E-17 | Bacteria | Verrucomicrobiota | Verrucomicrobiae | Chthoniobacterales | Chthoniobacteraceae |
| | ASV304 | 4.50 | 3.75E-08 | Bacteria | Bacteroidota | Bacteroidia | Chitinophagales | bacI |
| | ASV1375 | 4.47 | 1.84E-10 | Bacteria | Bacteroidota | Bacteroidia | Cytophagales | bacIII |
| | ASV1119 | 4.46 | 2.02E-16 | Bacteria | Proteobacteria | Alphaproteobacteria | Rhodospirillales | Magnetospirillaceae |
| | ASV1086 | 4.44 | 1.52E-11 | Bacteria | Actinobacteriota | Thermoleophilia | Gaiellales | uncultured |
| | ASV1098 | 4.37 | 3.12E-19 | Bacteria | Proteobacteria | Gammaproteobacteria | Burkholderiales | Comamonadaceae |
| | ASV1146 | 4.35 | 2.92E-14 | Bacteria | Proteobacteria | Gammaproteobacteria | Burkholderiales | Comamonadaceae |
| | ASV96 | 4.28 | 7.85E-08 | Bacteria | Cyanobacteria | Cyanobacteriia | Synechococcales | Cyanobiaceae |
| | ASV691 | 4.28 | 9.34E-10 | Bacteria | Proteobacteria | Gammaproteobacteria | Burkholderiales | betI |
| | ASV305 | 4.25 | 1.17E-10 | Bacteria | Proteobacteria | Gammaproteobacteria | Legionellales | Legionellaceae |
| | ASV1396 | 4.24 | 1.11E-07 | Bacteria | Bacteroidota | Bacteroidia | Flavobacteriales | bacII |
| | ASV1341 | 4.20 | 4.87E-11 | Bacteria | Proteobacteria | Gammaproteobacteria | Burkholderiales | Rhodocyclaceae |
| | ASV1058 | 4.19 | 8.34E-12 | Bacteria | Actinobacteriota | Acidimicrobiia | Microtrichales | Ilumatobacteraceae |
| | ASV354 | 4.16 | 1.10E-11 | Bacteria | Bacteroidota | Bacteroidia | Flavobacteriales | bacV |
| | ASV1057 | 4.14 | 3.90E-11 | Bacteria | Bdellovibrionota | Bdellovibrionia | Bdellovibrionales | Bdellovibrionaceae |
| | ASV1467 | 4.13 | 2.39E-12 | Bacteria | Proteobacteria | Gammaproteobacteria | Burkholderiales | betI |
| | ASV299 | 4.13 | 8.22E-12 | Bacteria | Verrucomicrobiota | Verrucomicrobiae | Verrucomicrobiales | Verrucomicrobiaceae |
| | ASV140 | 4.11 | 6.87E-10 | Bacteria | Proteobacteria | Alphaproteobacteria | Caulobacterales | Caulobacteraceae |
| | ASV1464 | 4.10 | 2.33E-13 | Bacteria | Bacteroidota | Bacteroidia | Flavobacteriales | Cryomorphaceae |
| | ASV1050 | 4.10 | 1.51E-13 | Bacteria | Proteobacteria | Gammaproteobacteria | Burkholderiales | Comamonadaceae |

| | | | | | | | | |
|---|---|---|---|---|---|---|---|---|
| | ASV1095 | 4.05 | 9.43E-13 | Bacteria | Bacteroidota | Bacteroidia | Sphingobacteriales | Lentimicrobiaceae |
| | ASV1068 | 4.00 | 7.88E-13 | Bacteria | Proteobacteria | Alphaproteobacteria | Rhodobacterales | Rhodobacteraceae |
| | ASV1097 | 3.99 | 2.85E-08 | Bacteria | Proteobacteria | Alphaproteobacteria | Rickettsiales | Rickettsiaceae |
| | ASV954 | 3.99 | 4.43E-06 | Bacteria | Proteobacteria | Alphaproteobacteria | Acetobacterales | alfVIII |
| | ASV1310 | 3.96 | 1.01E-13 | Bacteria | Proteobacteria | Gammaproteobacteria | Burkholderiales | Gallionellaceae |
| | ASV1436 | 3.93 | 9.15E-10 | Bacteria | Bacteroidota | Bacteroidia | Sphingobacteriales | NS11-12_marine_group |
| | ASV1088 | 3.91 | 2.50E-12 | Bacteria | Proteobacteria | Gammaproteobacteria | Burkholderiales | Rhodocyclaceae |
| | ASV1279 | 3.91 | 5.37E-12 | Bacteria | Bacteroidota | Bacteroidia | Sphingobacteriales | NS11-12_marine_group |
| | ASV1090 | 3.90 | 1.56E-10 | Bacteria | Acidobacteriota | Vicinamibacteria | Vicinamibacterales | uncultured |
| | ASV1346 | 3.87 | 6.75E-10 | Bacteria | Proteobacteria | Gammaproteobacteria | Burkholderiales | betI |
| | ASV556 | 3.80 | 3.39E-05 | Bacteria | Bacteroidota | Bacteroidia | Sphingobacteriales | bacVI |
| | ASV1386 | 3.78 | 3.00E-10 | Bacteria | Proteobacteria | Gammaproteobacteria | Methylococcales | Methylomonadaceae |
| | ASV1053 | 3.77 | 9.12E-05 | Bacteria | Cyanobacteria | Cyanobacteriia | Synechococcales | Cyanobiaceae |
| | ASV1414 | 3.77 | 6.02E-10 | Bacteria | Bacteroidota | Bacteroidia | Sphingobacteriales | NS11-12_marine_group |
| | ASV1357 | 3.76 | 1.65E-09 | Bacteria | Bacteroidota | Bacteroidia | Sphingobacteriales | Lentimicrobiaceae |
| | ASV967 | 3.75 | 9.44E-14 | Bacteria | Proteobacteria | Gammaproteobacteria | Oceanospirillales | Pseudohongiellaceae |
| | ASV1261 | 3.74 | 5.99E-12 | Bacteria | Proteobacteria | Gammaproteobacteria | Burkholderiales | Rhodocyclaceae |
| | ASV1432 | 3.72 | 3.82E-05 | Bacteria | Bacteroidota | Bacteroidia | Sphingobacteriales | Sphingobacteriaceae |
| | ASV1141 | 3.71 | 3.07E-14 | Bacteria | Proteobacteria | Gammaproteobacteria | Burkholderiales | Rhodocyclaceae |
| | ASV1226 | 3.67 | 5.57E-13 | Bacteria | Proteobacteria | Alphaproteobacteria | Rhodobacterales | Rhodobacteraceae |
| | ASV1148 | 3.66 | 7.19E-11 | Bacteria | Bacteroidota | Bacteroidia | Flavobacteriales | Crocinitomicaceae |
| | ASV364 | 3.66 | 1.49E-06 | Bacteria | Proteobacteria | Gammaproteobacteria | Burkholderiales | betI |
| | ASV1399 | 3.65 | 2.23E-10 | Bacteria | Bacteroidota | Bacteroidia | Sphingobacteriales | Lentimicrobiaceae |
| | ASV1290 | 3.64 | 2.33E-13 | Bacteria | Bacteroidota | Kryptonia | Kryptoniales | BSV26 |
| | ASV1133 | 3.64 | 6.87E-10 | Bacteria | Actinobacteriota | Acidimicrobiia | Microtrichales | acIV |
| | ASV681 | 3.63 | 4.54E-12 | Bacteria | Bacteroidota | Bacteroidia | Sphingobacteriales | NS11-12_marine_group |
| | ASV1495 | 3.56 | 4.46E-14 | Bacteria | Bacteroidota | Bacteroidia | Flavobacteriales | Crocinitomicaceae |
| | ASV1209 | 3.55 | 6.87E-10 | Bacteria | Proteobacteria | Gammaproteobacteria | Burkholderiales | Rhodocyclaceae |
| | ASV746 | 3.53 | 7.77E-07 | Bacteria | Bacteroidota | Bacteroidia | Chitinophagales | bacI |
| | ASV334 | 3.47 | 3.25E-08 | Bacteria | Verrucomicrobiota | Verrucomicrobiae | Verrucomicrobiales | Verrucomicrobiaceae |

| | | | | | | | | |
|---|---|---|---|---|---|---|---|---|
| | ASV429 | 3.46 | 6.80E-07 | Bacteria | Bacteroidota | Bacteroidia | Sphingobacteriales | NS11-12_marine_group |
| | ASV1025 | 3.42 | 1.94E-09 | Bacteria | Verrucomicrobiota | Verrucomicrobiae | Chthoniobacterales | Chthoniobacteraceae |
| | ASV1123 | 3.40 | 9.67E-12 | Bacteria | Bacteroidota | Bacteroidia | Bacteroidales | Prolixibacteraceae |
| | ASV1066 | 3.40 | 2.85E-11 | Bacteria | Bacteroidota | Bacteroidia | Chitinophagales | bacI |
| | ASV1453 | 3.39 | 2.76E-12 | Bacteria | Proteobacteria | Gammaproteobacteria | Coxiellales | Coxiellaceae |
| | ASV1215 | 3.38 | 1.88E-12 | Bacteria | Proteobacteria | Alphaproteobacteria | uncultured | uncultured |
| | ASV1444 | 3.38 | 6.02E-08 | Bacteria | Proteobacteria | Gammaproteobacteria | Burkholderiales | betI |
| | ASV1158 | 3.36 | 2.80E-12 | Bacteria | Planctomycetota | Phycisphaerae | Phycisphaerales | Phycisphaeraceae |
| | ASV1265 | 3.32 | 8.04E-08 | Bacteria | Cyanobacteria | Cyanobacteriia | Synechococcales | Cyanobiaceae |
| | ASV1362 | 3.31 | 3.45E-11 | Bacteria | Verrucomicrobiota | Verrucomicrobiae | Pedosphaerales | Pedosphaeraceae |
| | ASV794 | 3.30 | 0.001696 | Bacteria | Proteobacteria | Gammaproteobacteria | Burkholderiales | Gallionellaceae |
| | ASV1233 | 3.30 | 1.16E-11 | Bacteria | Bacteroidota | Bacteroidia | Sphingobacteriales | NS11-12_marine_group |
| | ASV1334 | 3.29 | 2.50E-13 | Bacteria | Proteobacteria | Gammaproteobacteria | Burkholderiales | Rhodocyclaceae |
| | ASV1177 | 3.28 | 3.00E-11 | Bacteria | Proteobacteria | Gammaproteobacteria | Burkholderiales | Comamonadaceae |
| | ASV632 | 3.28 | 2.99E-07 | Bacteria | Bacteroidota | Bacteroidia | Flavobacteriales | NS9_marine_group |
| | ASV1054 | 3.28 | 1.66E-10 | Bacteria | Bacteroidota | Bacteroidia | Flavobacteriales | bacII |
| | ASV1309 | 3.26 | 1.23E-11 | Bacteria | Proteobacteria | Gammaproteobacteria | Burkholderiales | Nitrosomonadaceae |
| | ASV1205 | 3.25 | 0.000292 | Bacteria | Proteobacteria | Gammaproteobacteria | Burkholderiales | Methylophilaceae |
| | ASV952 | 3.25 | 8.22E-12 | Bacteria | Bacteroidota | Bacteroidia | Sphingobacteriales | env.OPS_17 |
| | ASV1407 | 3.24 | 1.05E-13 | Bacteria | Bacteroidota | Bacteroidia | Flavobacteriales | NS9_marine_group |
| | ASV1317 | 3.23 | 9.94E-13 | Bacteria | Chloroflexi | Anaerolineae | Anaerolineales | Anaerolineaceae |
| | ASV1311 | 3.22 | 2.73E-10 | Bacteria | Proteobacteria | Gammaproteobacteria | Burkholderiales | Oxalobacteraceae |
| | ASV146 | 3.22 | 5.50E-05 | Bacteria | Proteobacteria | Alphaproteobacteria | Micropepsales | Micropepsaceae |
| | ASV1170 | 3.21 | 3.89E-09 | Bacteria | Bacteroidota | Kryptonia | Kryptoniales | BSV26 |
| | ASV424 | 3.21 | 7.90E-05 | Bacteria | Bacteroidota | Bacteroidia | Flavobacteriales | bacII |
| | ASV294 | 3.20 | 2.68E-08 | Bacteria | Planctomycetota | OM190 | OM190 | OM190 |
| | ASV679 | 3.19 | 7.74E-14 | Bacteria | Proteobacteria | Gammaproteobacteria | Burkholderiales | betI |
| | ASV763 | 3.16 | 7.35E-07 | Bacteria | Verrucomicrobiota | Verrucomicrobiae | Chthoniobacterales | Chthoniobacteraceae |
| | ASV1185 | 3.14 | 4.56E-15 | Bacteria | Bacteroidota | Kapabacteria | Kapabacteriales | Kapabacteriales |
| | ASV281 | 3.12 | 1.07E-06 | Bacteria | SAR324_clade(Marine_group_B) | SAR324_clade(Marine_group_B) | SAR324_clade(Marine_group_B) | SAR324_clade(Marine_group_B) |

| | | | | | | | | |
|---|---|---|---|---|---|---|---|---|
| | ASV1316 | 3.11 | 7.29E-08 | Bacteria | Gemmatimonadota | Gemmatimonadetes | Gemmatimonadales | Gemmatimonadaceae |
| | ASV1235 | 3.11 | 7.48E-12 | Bacteria | Proteobacteria | Gammaproteobacteria | Burkholderiales | Gallionellaceae |
| | ASV185 | 3.06 | 2.15E-05 | Bacteria | Bacteroidota | Bacteroidia | Flavobacteriales | NS9_marine_group |
| | ASV947 | 3.04 | 0.000271 | Bacteria | Bacteroidota | Bacteroidia | Flavobacteriales | bacII |
| | ASV1155 | 3.02 | 5.37E-09 | Bacteria | Actinobacteriota | Actinobacteria | Frankiales | acI |
| | ASV901 | 3.00 | 1.02E-05 | Bacteria | Planctomycetota | Phycisphaerae | Phycisphaerales | Phycisphaeraceae |
| | ASV506 | 2.99 | 5.30E-05 | Bacteria | Actinobacteriota | Actinobacteria | Micrococcales | Luna1 |
| | ASV1298 | 2.98 | 4.55E-12 | Bacteria | Gemmatimonadota | Gemmatimonadetes | Gemmatimonadales | Gemmatimonadaceae |
| | ASV1264 | 2.98 | 7.23E-09 | Bacteria | Proteobacteria | Gammaproteobacteria | Burkholderiales | Comamonadaceae |
| | ASV384 | 2.96 | 2.12E-05 | Bacteria | Proteobacteria | Gammaproteobacteria | Burkholderiales | betII |
| | ASV1450 | 2.95 | 2.14E-10 | Bacteria | Bacteroidota | Kapabacteria | Kapabacteriales | Kapabacteriales |
| | ASV201 | 2.95 | 0.000901 | Bacteria | Proteobacteria | Alphaproteobacteria | Sphingomonadales | Sphingomonadaceae |
| | ASV1418 | 2.95 | 1.28E-11 | Bacteria | Desulfobacterota | Desulfuromonadia | PB19 | PB19 |
| | ASV1281 | 2.94 | 1.68E-09 | Bacteria | Bdellovibrionota | Bdellovibrionia | Bdellovibrionales | Bdellovibrionaceae |
| | ASV1196 | 2.93 | 3.67E-08 | Bacteria | Planctomycetota | Phycisphaerae | Phycisphaerales | Phycisphaeraceae |
| | ASV1336 | 2.92 | 0.000166 | Bacteria | Verrucomicrobiota | Verrucomicrobiae | uncultured | uncultured |
| | ASV1502 | 2.91 | 1.44E-05 | Bacteria | Actinobacteriota | Thermoleophilia | Gaiellales | uncultured |
| | ASV1242 | 2.91 | 0.00068 | Bacteria | Proteobacteria | Gammaproteobacteria | Methylococcales | Methylomonadaceae |
| | ASV252 | 2.87 | 0.000271 | Bacteria | Bacteroidota | Bacteroidia | Flavobacteriales | Crocinitomicaceae |
| | ASV587 | 2.86 | 6.52E-06 | Bacteria | Chloroflexi | Chloroflexia | Chloroflexales | Roseiflexaceae |
| | ASV1475 | 2.86 | 1.56E-10 | Bacteria | Verrucomicrobiota | Verrucomicrobiae | Pedosphaerales | Pedosphaeraceae |
| | ASV1110 | 2.83 | 1.08E-06 | Bacteria | Actinobacteriota | Actinobacteria | Frankiales | acSTL |
| | ASV32 | 2.80 | 1.02E-05 | Bacteria | Verrucomicrobiota | Verrucomicrobiae | uncultured | uncultured |
| | ASV541 | 2.79 | 2.36E-11 | Bacteria | Proteobacteria | Gammaproteobacteria | Burkholderiales | Oxalobacteraceae |
| | ASV1203 | 2.78 | 2.07E-10 | Bacteria | Bdellovibrionota | Oligoflexia | 0319-6G20 | 0319-6G20 |
| | ASV1143 | 2.75 | 2.55E-07 | Bacteria | Proteobacteria | Gammaproteobacteria | Burkholderiales | Comamonadaceae |
| | ASV363 | 2.74 | 9.85E-07 | Bacteria | Bacteroidota | Bacteroidia | Sphingobacteriales | bacVI |
| | ASV1430 | 2.74 | 1.22E-07 | Bacteria | Bacteroidota | Bacteroidia | Chitinophagales | Chitinophagaceae |
| | ASV488 | 2.71 | 2.21E-10 | Bacteria | Desulfobacterota | Desulfuromonadia | PB19 | PB19 |
| | ASV787 | 2.71 | 3.11E-05 | Bacteria | Planctomycetota | Phycisphaerae | Phycisphaerales | Phycisphaeraceae |

| | | | | | | | | |
|---|---|---|---|---|---|---|---|---|
| | ASV1083 | 2.71 | 2.48E-08 | Bacteria | Proteobacteria | Gammaproteobacteria | Burkholderiales | Rhodocyclaceae |
| | ASV337 | 2.71 | 7.22E-06 | Bacteria | Verrucomicrobiota | Verrucomicrobiae | Chthoniobacterales | Chthoniobacteraceae |
| | ASV285 | 2.68 | 0.004819 | Bacteria | Bacteroidota | Bacteroidia | Sphingobacteriales | bacVI |
| | ASV1304 | 2.66 | 7.18E-10 | Bacteria | Firmicutes | Bacilli | Paenibacillales | Paenibacillaceae |
| | ASV1505 | 2.64 | 0.000143 | Bacteria | Proteobacteria | Gammaproteobacteria | Methylococcales | Methylomonadaceae |
| | ASV579 | 2.60 | 1.15E-07 | Bacteria | Proteobacteria | Gammaproteobacteria | Burkholderiales | Comamonadaceae |
| | ASV582 | 2.60 | 3.29E-05 | Bacteria | Bdellovibrionota | Bdellovibrionia | Bdellovibrionales | Bdellovibrionaceae |
| | ASV1255 | 2.58 | 1.31E-07 | Bacteria | Bacteroidota | Bacteroidia | Sphingobacteriales | NS11-12_marine_group |
| | ASV1319 | 2.57 | 6.83E-10 | Bacteria | Proteobacteria | Gammaproteobacteria | Methylococcales | Methylomonadaceae |
| | ASV1165 | 2.55 | 3.34E-10 | Bacteria | Bdellovibrionota | Oligoflexia | 053A03-B-DI-P58 | 053A03-B-DI-P58 |
| | ASV1354 | 2.54 | 0.000112 | Bacteria | Bacteroidota | Bacteroidia | Flavobacteriales | Crocinitomicaceae |
| | ASV177 | 2.53 | 4.07E-06 | Bacteria | Bacteroidota | Bacteroidia | Chitinophagales | 37-13 |
| | ASV1118 | 2.52 | 2.47E-06 | Bacteria | Proteobacteria | Gammaproteobacteria | Burkholderiales | Hydrogenophilaceae |
| | ASV1268 | 2.52 | 6.45E-08 | Bacteria | Bdellovibrionota | Oligoflexia | Oligoflexales | uncultured |
| | ASV157 | 2.49 | 6.84E-07 | Bacteria | Bacteroidota | Bacteroidia | Bacteroidales | Bacteroidaceae |
| | ASV1491 | 2.47 | 2.62E-05 | Bacteria | Actinobacteriota | Actinobacteria | Micrococcales | Luna1 |
| | ASV628 | 2.47 | 0.000101 | Bacteria | Proteobacteria | Gammaproteobacteria | Xanthomonadales | Xanthomonadaceae |
| | ASV176 | 2.45 | 3.07E-05 | Bacteria | Bacteroidota | Bacteroidia | Flavobacteriales | Crocinitomicaceae |
| | ASV226 | 2.45 | 0.002424 | Bacteria | Bacteroidota | Bacteroidia | Chitinophagales | bacI |
| | ASV376 | 2.44 | 0.000473 | Bacteria | Planctomycetota | Planctomycetes | Gemmatales | Gemmataceae |
| | ASV1223 | 2.44 | 0.003613 | Bacteria | Proteobacteria | Gammaproteobacteria | Burkholderiales | Comamonadaceae |
| | ASV1062 | 2.41 | 5.16E-08 | Bacteria | Verrucomicrobiota | Verrucomicrobiae | Pedosphaerales | Pedosphaeraceae |
| | ASV1342 | 2.41 | 2.52E-07 | Bacteria | Proteobacteria | Gammaproteobacteria | Burkholderiales | Comamonadaceae |
| | ASV1404 | 2.40 | 1.02E-09 | Bacteria | Verrucomicrobiota | Verrucomicrobiae | Opitutales | Opitutaceae |
| | ASV1333 | 2.39 | 1.22E-07 | Bacteria | Proteobacteria | Alphaproteobacteria | Rhizobiales | Beijerinckiaceae |
| | ASV1438 | 2.38 | 1.11E-07 | Bacteria | Dependentiae | Babeliae | Babeliales | Babeliales |
| | ASV890 | 2.38 | 2.27E-05 | Bacteria | Proteobacteria | Gammaproteobacteria | Coxiellales | Coxiellaceae |
| | ASV754 | 2.38 | 3.06E-05 | Bacteria | Proteobacteria | Alphaproteobacteria | Rhodospirillales | Rhodospirillaceae |
| | ASV1349 | 2.37 | 0.00142 | Bacteria | Bacteroidota | Bacteroidia | Flavobacteriales | bacV |
| | ASV1435 | 2.37 | 9.41E-05 | Bacteria | Bacteroidota | Bacteroidia | Chitinophagales | Saprospiraceae |

| | | | | | | | | |
|---|---|---|---|---|---|---|---|---|
| | ASV1116 | 2.35 | 9.91E-05 | Bacteria | Cyanobacteria | Cyanobacteriia | Synechococcales | Cyanobiaceae |
| | ASV1225 | 2.34 | 1.12E-07 | Bacteria | Proteobacteria | Gammaproteobacteria | Burkholderiales | Burkholderiaceae |
| | ASV194 | 2.34 | 0.000103 | Bacteria | Proteobacteria | Gammaproteobacteria | Burkholderiales | betII |
| | ASV1122 | 2.32 | 0.000537 | Bacteria | Proteobacteria | Gammaproteobacteria | Burkholderiales | betI |
| | ASV381 | 2.32 | 1.10E-09 | Bacteria | Actinobacteriota | Actinobacteria | Frankiales | acI |
| | ASV1393 | 2.30 | 0.002772 | Bacteria | Bacteroidota | Bacteroidia | Flavobacteriales | Crocinitomicaceae |
| | ASV1348 | 2.30 | 4.12E-09 | Bacteria | Proteobacteria | Gammaproteobacteria | Burkholderiales | Rhodocyclaceae |
| | ASV528 | 2.28 | 0.001195 | Bacteria | Proteobacteria | Gammaproteobacteria | Burkholderiales | betI |
| | ASV1113 | 2.28 | 4.34E-09 | Bacteria | Proteobacteria | Gammaproteobacteria | Burkholderiales | betII |
| | ASV747 | 2.24 | 2.96E-05 | Bacteria | Verrucomicrobiota | Verrucomicrobiae | Verrucomicrobiales | Rubritaleaceae |
| | ASV1076 | 2.21 | 2.30E-07 | Bacteria | Bacteroidota | Bacteroidia | Sphingobacteriales | Lentimicrobiaceae |
| | ASV393 | 2.21 | 1.15E-07 | Bacteria | Verrucomicrobiota | Verrucomicrobiae | Verrucomicrobiales | Rubritaleaceae |
| | ASV1313 | 2.19 | 1.33E-05 | Bacteria | Actinobacteriota | Acidimicrobiia | Microtrichales | acIV |
| | ASV1403 | 2.18 | 1.06E-07 | Bacteria | Proteobacteria | Gammaproteobacteria | Burkholderiales | Rhodocyclaceae |
| | ASV730 | 2.17 | 1.79E-09 | Bacteria | Proteobacteria | Alphaproteobacteria | Paracaedibacterales | Paracaedibacteraceae |
| | ASV1423 | 2.15 | 1.55E-06 | Bacteria | Bacteroidota | Bacteroidia | Flavobacteriales | Crocinitomicaceae |
| | ASV78 | 2.14 | 1.96E-05 | Bacteria | Bdellovibrionota | Bdellovibrionia | Bdellovibrionales | Bdellovibrionaceae |
| | ASV1405 | 2.14 | 3.79E-06 | Bacteria | Bdellovibrionota | Oligoflexia | 0319-6G20 | 0319-6G20 |
| | ASV455 | 2.13 | 0.000166 | Bacteria | Proteobacteria | Gammaproteobacteria | Burkholderiales | Comamonadaceae |
| | ASV1294 | 2.13 | 1.25E-09 | Bacteria | Proteobacteria | Alphaproteobacteria | Rhodospirillales | Magnetospirillaceae |
| | ASV1411 | 2.13 | 0.000124 | Bacteria | Proteobacteria | Gammaproteobacteria | Burkholderiales | betI |
| | ASV1434 | 2.12 | 1.74E-06 | Bacteria | Verrucomicrobiota | Verrucomicrobiae | Pedosphaerales | Pedosphaeraceae |
| | ASV1186 | 2.11 | 1.57E-06 | Bacteria | Bacteroidota | Bacteroidia | Chitinophagales | bacI |
| | ASV1206 | 2.10 | 0.000137 | Bacteria | Verrucomicrobiota | Verrucomicrobiae | Chthoniobacterales | Chthoniobacteraceae |
| | ASV983 | 2.10 | 7.33E-05 | Bacteria | Actinobacteriota | Acidimicrobiia | Microtrichales | Ilumatobacteraceae |
| | ASV1377 | 2.09 | 1.03E-08 | Bacteria | Bacteroidota | Bacteroidia | Sphingobacteriales | bacVI |
| | ASV1472 | 2.08 | 5.80E-06 | Bacteria | Proteobacteria | Gammaproteobacteria | Burkholderiales | Rhodocyclaceae |
| | ASV1232 | 2.08 | 7.21E-07 | Bacteria | Bacteroidota | Bacteroidia | Flavobacteriales | bacV |
| | ASV479 | 2.07 | 0.000206 | Bacteria | Bacteroidota | Bacteroidia | Sphingobacteriales | bacVI |
| | ASV1192 | 2.07 | 6.02E-06 | Bacteria | Bacteroidota | Bacteroidia | Flavobacteriales | bacII |

| | | | | | | | | |
|---|---|---|---|---|---|---|---|---|
| | ASV641 | 2.06 | 6.76E-05 | Bacteria | Bdellovibrionota | Bdellovibrionia | Bdellovibrionales | Bdellovibrionaceae |
| | ASV1300 | 2.05 | 1.32E-05 | Bacteria | Proteobacteria | Gammaproteobacteria | Burkholderiales | Neisseriaceae |
| | ASV1297 | 2.05 | 5.68E-08 | Bacteria | Margulisbacteria | Margulisbacteria | Margulisbacteria | Margulisbacteria |
| | ASV1138 | 2.04 | 1.24E-07 | Bacteria | Bdellovibrionota | Oligoflexia | 0319-6G20 | 0319-6G20 |
| | ASV1443 | 2.03 | 2.15E-06 | Bacteria | Proteobacteria | Gammaproteobacteria | Burkholderiales | Comamonadaceae |
| | ASV454 | 2.03 | 0.000606 | Bacteria | Actinobacteriota | Actinobacteria | Micrococcales | Luna1 |
| | ASV244 | 2.03 | 8.95E-05 | Bacteria | Planctomycetota | Planctomycetes | Gemmatales | Gemmataceae |
| | ASV1238 | 2.03 | 6.52E-08 | Bacteria | Firmicutes | Bacilli | Lactobacillales | Streptococcaceae |
| | ASV1193 | 2.03 | 7.60E-06 | Bacteria | Proteobacteria | Gammaproteobacteria | Burkholderiales | Comamonadaceae |
| | ASV1476 | 2.01 | 4.15E-05 | Bacteria | Bacteroidota | Bacteroidia | Flavobacteriales | bacV |
| | ASV1500 | 2.00 | 1.48E-06 | Bacteria | Bacteroidota | Bacteroidia | Chitinophagales | Saprospiraceae |

**Table S5: Number of affected ASVs per family and their proportion in the compared to the total number of ASVs in the family in ULC.** Families are ordered by the proportion of ASVs affected and categorised as follows: eukaryotic nanoplankton (18S-NP), eukaryotic picoplankton (18S-PP) and prokaryotic picoplankton (16S-PP).

| Microbial plankton | Family | Total ASVs | Affected ASVs | Proportion | Kingdom | Phylum | Class | Order |
|---|---|---|---|---|---|---|---|---|
| 18S-NP | Amphisiellidae | 4 | 4 | 100.0 | Eukaryota | Ciliophora | Spirotrichea | Stichotrichida |
| | Noelaerhabdaceae | 4 | 4 | 100.0 | Eukaryota | Haptophyta | Coccolithophyceae | Isochrysidales |
| | Chytridiaceae | 5 | 4 | 80.0 | Fungi | Chytridiomycota | Chytridiomycetes | Chytridiales |
| | Holophryidae | 5 | 4 | 80.0 | Eukaryota | Ciliophora | Prostomatea | Prorodontida |
| | Peronosporaceae | 5 | 4 | 80.0 | Eukaryota | Oomycota | Peronosporea | Peronosporales |
| | Stephanodiscaceae | 46 | 35 | 76.1 | Eukaryota | Bacillariophyta | Mediophyceae | Stephanodiscales |
| | Didiniidae | 6 | 4 | 66.7 | Eukaryota | Ciliophora | Litostomatea | Haptorida |
| | Geminigeraceae | 12 | 8 | 66.7 | Eukaryota | Cryptophyta | Cryptophyceae | Pyrenomonadales |
| | Tetrahymenidae | 3 | 2 | 66.7 | Eukaryota | Ciliophora | Oligohymenophorea | Hymenostomatida |
| | Gymnodiniaceae | 44 | 24 | 54.5 | Eukaryota | Myzozoa | Dinophyceae | Gymnodiniales |
| | Chlorodendraceae | 8 | 4 | 50.0 | Eukaryota | Chlorophyta | Chlorodendrophyceae | Chlorodendrales |
| | Massisteriidae | 2 | 1 | 50.0 | Eukaryota | Cercozoa | Granofilosea | Leucodictyida |
| | Thalassiosiraceae | 17 | 7 | 41.2 | Eukaryota | Bacillariophyta | Mediophyceae | Thalassiosirales |
| | Protozoa familia incertae sedis | 32 | 12 | 37.5 | Eukaryota | Cercozoa | Protozoa classis incertae sedis | Protozoa ordo incertae sedis |
| | Telonemia familia ineditae | 16 | 6 | 37.5 | Eukaryota | Telonemia | Telonemia classis ineditae | Telonemida |
| | Amphidiniaceae | 11 | 4 | 36.4 | Eukaryota | Myzozoa | Dinophyceae | Amphidiniales |
| | Pfiesteriaceae | 22 | 8 | 36.4 | Eukaryota | Myzozoa | Dinophyceae | Peridiniales |
| | Mallomonadaceae | 37 | 13 | 35.1 | Eukaryota | Ochrophyta | Chrysophyceae | Synurales |
| | Colepidae | 66 | 23 | 34.8 | Eukaryota | Ciliophora | Prostomatea | Prorodontida |
| | Protaspidae | 59 | 20 | 33.9 | Eukaryota | Cercozoa | Thecofilosea | Cryomonadida |
| | Cercomonadidae | 3 | 1 | 33.3 | Eukaryota | Cercozoa | Sarcomonadea | Cercomonadida |
| | Strobilidiidae | 18 | 6 | 33.3 | Eukaryota | Ciliophora | Spirotrichea | Choreotrichida |
| | Tabellariaceae | 9 | 3 | 33.3 | Eukaryota | Bacillariophyta | Bacillariophyceae | Rhabdonematales |
| | Katablepharidaceae | 17 | 5 | 29.4 | Eukaryota | Cryptophyta | Cryptophyceae | Kathablepharidacea |

| | | | | | | | | |
|---|---|---|---|---|---|---|---|---|
| | Chrysochromulinaceae | 11 | 3 | 27.3 | Eukaryota | Haptophyta | Coccolithophyceae | Prymnesiales |
| | Spathidiidae | 54 | 13 | 24.1 | Eukaryota | Ciliophora | Litostomatea | Haptorida |
| | Thoracosphaeraceae | 9 | 2 | 22.2 | Eukaryota | Myzozoa | Dinophyceae | Thoracosphaerales |
| | Paraphysomonadaceae | 54 | 11 | 20.4 | Eukaryota | Ochrophyta | Chrysophyceae | Chromulinales |
| | Histiobalantiidae | 5 | 1 | 20.0 | Eukaryota | Ciliophora | Oligohymenophorea | Pleuronematida |
| | Tintinnidae | 21 | 4 | 19.0 | Eukaryota | Ciliophora | Spirotrichea | Tintinnida |
| | Strombidiidae | 28 | 4 | 14.3 | Eukaryota | Ciliophora | Spirotrichea | Oligotrichia |
| | Suessiaceae | 35 | 4 | 11.4 | Eukaryota | Myzozoa | Dinophyceae | Suessiales |
| | Chlamydomonadaceae | 36 | 4 | 11.1 | Eukaryota | Chlorophyta | Chlorophyceae | Chlamydomonadales |
| | Bicosoecidae | 19 | 2 | 10.5 | Eukaryota | Bigyra | Bikosea | Bicosoecida |
| | Pseudophyllomitidae | 10 | 1 | 10.0 | Eukaryota | Bigyra | incertae sedis | incertae sedis |
| 18S-PP | Amphisiellidae | 4 | 4 | 100.0 | Eukaryota | Ciliophora | Spirotrichea | Stichotrichida |
| | Ancistridae | 3 | 3 | 100.0 | Eukaryota | Ciliophora | Oligohymenophorea | Thigmotrichida |
| | Chlorellales incertae sedis | 4 | 4 | 100.0 | Eukaryota | Chlorophyta | Trebouxiophyceae | Chlorellales |
| | Mychonastaceae | 3 | 3 | 100.0 | Eukaryota | Chlorophyta | Chlorophyceae | Sphaeropleales |
| | Peronosporaceae | 2 | 2 | 100.0 | Eukaryota | Oomycota | Peronosporea | Peronosporales |
| | Stephanodiscaceae | 21 | 18 | 85.7 | Eukaryota | Bacillariophyta | Mediophyceae | Stephanodiscales |
| | Geminigeraceae | 28 | 21 | 75.0 | Eukaryota | Cryptophyta | Cryptophyceae | Pyrenomonadales |
| | Noelaerhabdaceae | 4 | 3 | 75.0 | Eukaryota | Haptophyta | Coccolithophyceae | Isochrysidales |
| | Pfiesteriaceae | 12 | 8 | 66.7 | Eukaryota | Myzozoa | Dinophyceae | Peridiniales |
| | Thalassiosiraceae | 3 | 2 | 66.7 | Eukaryota | Bacillariophyta | Mediophyceae | Thalassiosirales |
| | Dolichomastigaceae | 7 | 4 | 57.1 | Eukaryota | Chlorophyta | Mamiellophyceae | Dolichomastigales |
| | Gymnodiniaceae | 54 | 28 | 51.9 | Eukaryota | Myzozoa | Dinophyceae | Gymnodiniales |
| | Acanthoecidae | 2 | 1 | 50.0 | Eukaryota | Choanozoa | Choanoflagellata | Acanthoecida |
| | Chlorellaceae | 8 | 4 | 50.0 | Eukaryota | Chlorophyta | Trebouxiophyceae | Chlorellales |
| | Chytridiaceae | 8 | 4 | 50.0 | Fungi | Chytridiomycota | Chytridiomycetes | Chytridiales |
| | Telonemia familia ineditae | 19 | 9 | 47.4 | Eukaryota | Telonemia | Telonemia classis ineditae | Telonemida |
| | Protozoa familia incertae sedis | 34 | 13 | 38.2 | Eukaryota | Cercozoa | Protozoa classis incertae sedis | Protozoa ordo incertae sedis |

| | | | | | | | | |
|---|---|---|---|---|---|---|---|---|
| | Filasterea familia incertae sedis | 12 | 4 | 33.3 | Eukaryota | Choanozoa | Filasterea | Filasterea ordo incertae sedis |
| | Pirsoniaceae | 6 | 2 | 33.3 | Eukaryota | Oomycota | Hyphochytrea | Pirsoniales |
| | Triparmaceae | 6 | 2 | 33.3 | Eukaryota | Ochrophyta | Bolidophyceae | Parmales |
| | Chrysochromulinaceae | 27 | 8 | 29.6 | Eukaryota | Haptophyta | Coccolithophyceae | Prymnesiales |
| | Erysiphaceae | 4 | 1 | 25.0 | Fungi | Ascomycota | Leotiomycetes | Erysiphales |
| | Colepidae | 62 | 15 | 24.2 | Eukaryota | Ciliophora | Prostomatea | Prorodontida |
| | Pseudophyllomitidae | 21 | 5 | 23.8 | Eukaryota | Bigyra | incertae sedis | incertae sedis |
| | Uebelmesseromycetaceae | 17 | 4 | 23.5 | Fungi | Chytridiomycota | Chytridiomycetes | Rhizophydiales |
| | Protaspidae | 30 | 7 | 23.3 | Eukaryota | Cercozoa | Thecofilosea | Cryomonadida |
| | Amphidiniaceae | 35 | 8 | 22.9 | Eukaryota | Myzozoa | Dinophyceae | Amphidiniales |
| | Katablepharidaceae | 35 | 8 | 22.9 | Eukaryota | Cryptophyta | Cryptophyceae | Kathablepharidacea |
| | Mallomonadaceae | 40 | 8 | 20.0 | Eukaryota | Ochrophyta | Chrysophyceae | Synurales |
| | Suessiaceae | 16 | 3 | 18.8 | Eukaryota | Myzozoa | Dinophyceae | Suessiales |
| | Codonosigaceae | 48 | 8 | 16.7 | Eukaryota | Choanozoa | Choanoflagellata | Craspedida |
| | Actinomonadaceae | 46 | 7 | 15.2 | Eukaryota | Ochrophyta | Dictyochophyceae | Pedinellales |
| | Chrysolepidomonadaceae | 27 | 4 | 14.8 | Eukaryota | Ochrophyta | Chrysophyceae | Chromulinales |
| | Strombidiidae | 28 | 4 | 14.3 | Eukaryota | Ciliophora | Oligotrichea | Oligotrichida |
| | Paraphysomonadaceae | 129 | 18 | 14.0 | Eukaryota | Ochrophyta | Chrysophyceae | Chromulinales |
| | Crustomastigaceae | 12 | 1 | 8.3 | Eukaryota | Chlorophyta | Mamiellophyceae | Dolichomastigales |
| | Ochromonadaceae | 50 | 3 | 6.0 | Eukaryota | Ochrophyta | Chrysophyceae | Ochromonadales |
| | Prorocentraceae | 69 | 4 | 5.8 | Eukaryota | Myzozoa | Dinophyceae | Prorocentrales |
| 16S-PP | AncK6 | 1 | 1 | 100.0 | Bacteria | AncK6 | AncK6 | AncK6 |
| | bacteriap25 | 1 | 1 | 100.0 | Bacteria | Myxococcota | bacteriap25 | bacteriap25 |
| | BD2-11_terrestrial_group | 1 | 1 | 100.0 | Bacteria | Gemmatimonadota | BD2-11_terrestrial_group | BD2-11_terrestrial_group |
| | Clade_I | 1 | 1 | 100.0 | Bacteria | Proteobacteria | Alphaproteobacteria | SAR11_clade |
| | EF100-94H03 | 1 | 1 | 100.0 | Bacteria | Proteobacteria | Alphaproteobacteria | Puniceispirillales |
| | Gaiellaceae | 1 | 1 | 100.0 | Bacteria | Actinobacteriota | Thermoleophilia | Gaiellales |
| | KD4-96 | 1 | 1 | 100.0 | Bacteria | Chloroflexi | KD4-96 | KD4-96 |

| | | | | | | | | |
|---|---|---|---|---|---|---|---|---|
| | KI89A_clade | 1 | 1 | 100.0 | Bacteria | Proteobacteria | Gammaproteobacteria | KI89A_clade |
| | Nitrosopumilaceae | 1 | 1 | 100.0 | Archaea | Crenarchaeota | Nitrososphaeria | Nitrosopumilales |
| | P2-11E | 1 | 1 | 100.0 | Bacteria | Chloroflexi | P2-11E | P2-11E |
| | Bryobacteraceae | 4 | 3 | 75.0 | Bacteria | Acidobacteriota | Acidobacteriae | Bryobacterales |
| | JG30-KF-CM66 | 8 | 6 | 75.0 | Bacteria | Chloroflexi | JG30-KF-CM66 | JG30-KF-CM66 |
| | Reyranellaceae | 4 | 3 | 75.0 | Bacteria | Proteobacteria | Alphaproteobacteria | Reyranellales |
| | Vicinamibacteraceae | 20 | 15 | 75.0 | Bacteria | Acidobacteriota | Vicinamibacteria | Vicinamibacterales |
| | Anaerolineaceae | 3 | 2 | 66.7 | Bacteria | Chloroflexi | Anaerolineae | Anaerolineales |
| | KD3-93 | 3 | 2 | 66.7 | Bacteria | Bacteroidota | Bacteroidia | Sphingobacteriales |
| | Nitrospiraceae | 3 | 2 | 66.7 | Bacteria | Nitrospirota | Nitrospiria | Nitrospirales |
| | CCM19a | 2 | 1 | 50.0 | Bacteria | Proteobacteria | Gammaproteobacteria | CCM19a |
| | Clade_II | 2 | 1 | 50.0 | Bacteria | Proteobacteria | Alphaproteobacteria | SAR11_clade |
| | Ferrovibrionales | 2 | 1 | 50.0 | Bacteria | Proteobacteria | Alphaproteobacteria | Ferrovibrionales |
| | Iamiaceae | 2 | 1 | 50.0 | Bacteria | Actinobacteriota | Acidimicrobiia | Microtrichales |
| | IMCC26256 | 2 | 1 | 50.0 | Bacteria | Actinobacteriota | Acidimicrobiia | IMCC26256 |
| | Rhodospirillaceae | 8 | 4 | 50.0 | Bacteria | Proteobacteria | Alphaproteobacteria | Rhodospirillales |
| | Solirubrobacteraceae | 2 | 1 | 50.0 | Bacteria | Actinobacteriota | Thermoleophilia | Solirubrobacterales |
| | UBA12409 | 2 | 1 | 50.0 | Bacteria | Dependentiae | Babeliae | Babeliales |
| | Acetobacteraceae | 12 | 5 | 41.7 | Bacteria | Proteobacteria | Alphaproteobacteria | Acetobacterales |
| | Xanthobacteraceae | 11 | 4 | 36.4 | Bacteria | Proteobacteria | Alphaproteobacteria | Rhizobiales |
| | betIV | 6 | 2 | 33.3 | Bacteria | Proteobacteria | Gammaproteobacteria | Burkholderiales |
| | BSV26 | 3 | 1 | 33.3 | Bacteria | Bacteroidota | Kryptonia | Kryptoniales |
| | KF-JG30-B3 | 3 | 1 | 33.3 | Bacteria | Proteobacteria | Alphaproteobacteria | Rhizobiales |
| | Micropepsaceae | 3 | 1 | 33.3 | Bacteria | Proteobacteria | Alphaproteobacteria | Micropepsales |
| | Saccharimonadales | 3 | 1 | 33.3 | Bacteria | Patescibacteria | Saccharimonadia | Saccharimonadales |
| | TRA3-20 | 15 | 5 | 33.3 | Bacteria | Proteobacteria | Gammaproteobacteria | Burkholderiales |
| | Phycisphaeraceae | 22 | 7 | 31.8 | Bacteria | Planctomycetota | Phycisphaerae | Phycisphaerales |
| | Pedosphaeraceae | 19 | 6 | 31.6 | Bacteria | Verrucomicrobiota | Verrucomicrobiae | Pedosphaerales |
| | Gemmatimonadaceae | 10 | 3 | 30.0 | Bacteria | Gemmatimonadota | Gemmatimonadetes | Gemmatimonadales |
| | Nitrosomonadaceae | 14 | 4 | 28.6 | Bacteria | Proteobacteria | Gammaproteobacteria | Burkholderiales |

| | | | | | | | | |
|---|---|---|---|---|---|---|---|---|
| | uncultured | 52 | 14 | 26.9 | Bacteria | unknown | unknown | unknown |
| | Acidobacteriae | 4 | 1 | 25.0 | Bacteria | Acidobacteriota | Acidobacteriae | Acidobacteriae |
| | Hyphomicrobiaceae | 8 | 2 | 25.0 | Bacteria | Proteobacteria | Alphaproteobacteria | Rhizobiales |
| | Ilumatobacteraceae | 8 | 2 | 25.0 | Bacteria | Actinobacteriota | Acidimicrobiia | Microtrichales |
| | Pseudohongiellaceae | 9 | 2 | 22.2 | Bacteria | Proteobacteria | Gammaproteobacteria | Oceanospirillales |
| | Blfdi19 | 10 | 2 | 20.0 | Bacteria | Myxococcota | Polyangia | Blfdi19 |
| | Opitutaceae | 10 | 2 | 20.0 | Bacteria | Verrucomicrobiota | Verrucomicrobiae | Opitutales |
| | Rhodobacteraceae | 10 | 2 | 20.0 | Bacteria | Proteobacteria | Alphaproteobacteria | Rhodobacterales |
| | NS11-12_marine_group | 29 | 5 | 17.2 | Bacteria | Bacteroidota | Bacteroidia | Sphingobacteriales |
| | 0319-6G20 | 6 | 1 | 16.7 | Bacteria | Bdellovibrionota | Oligoflexia | 0319-6G20 |
| | betIII | 6 | 1 | 16.7 | Bacteria | Proteobacteria | Gammaproteobacteria | Burkholderiales |
| | Margulisbacteria | 6 | 1 | 16.7 | Bacteria | Margulisbacteria | Margulisbacteria | Margulisbacteria |
| | OM190 | 6 | 1 | 16.7 | Bacteria | Planctomycetota | OM190 | OM190 |
| | Sporichthyaceae | 6 | 1 | 16.7 | Bacteria | Actinobacteriota | Actinobacteria | Frankiales |
| | Unknown_Family | 6 | 1 | 16.7 | Bacteria | Proteobacteria | Gammaproteobacteria | Gammaproteobacteria_Incertae_Sedis |
| | Methylomonadaceae | 13 | 2 | 15.4 | Bacteria | Proteobacteria | Gammaproteobacteria | Methylococcales |
| | alfI | 7 | 1 | 14.3 | Bacteria | Proteobacteria | Alphaproteobacteria | Rhizobiales |
| | Gemmataceae | 7 | 1 | 14.3 | Bacteria | Planctomycetota | Planctomycetes | Gemmatales |
| | Kapabacteriales | 7 | 1 | 14.3 | Bacteria | Bacteroidota | Kapabacteria | Kapabacteriales |
| | Sphingobacteriaceae | 7 | 1 | 14.3 | Bacteria | Bacteroidota | Bacteroidia | Sphingobacteriales |
| | Chitinophagaceae | 27 | 3 | 11.1 | Bacteria | Bacteroidota | Bacteroidia | Chitinophagales |
| | Cryomorphaceae | 9 | 1 | 11.1 | Bacteria | Bacteroidota | Bacteroidia | Flavobacteriales |
| | Oxalobacteraceae | 9 | 1 | 11.1 | Bacteria | Proteobacteria | Gammaproteobacteria | Burkholderiales |
| | alfII | 10 | 1 | 10.0 | Bacteria | Proteobacteria | Alphaproteobacteria | Caulobacterales |
| | PeM15 | 10 | 1 | 10.0 | Bacteria | Actinobacteriota | Actinobacteria | PeM15 |
| | SAR324_clade(Marine_group_B) | 11 | 1 | 9.1 | Bacteria | SAR324_clade(Marine_group_B) | SAR324_clade(Marine_group_B) | SAR324_clade(Marine_group_B) |
| | env.OPS_17 | 12 | 1 | 8.3 | Bacteria | Bacteroidota | Bacteroidia | Sphingobacteriales |
| | NS9_marine_group | 12 | 1 | 8.3 | Bacteria | Bacteroidota | Bacteroidia | Flavobacteriales |
| | Beijerinckiaceae | 13 | 1 | 7.7 | Bacteria | Proteobacteria | Alphaproteobacteria | Rhizobiales |
| | Legionellaceae | 29 | 2 | 6.9 | Bacteria | Proteobacteria | Gammaproteobacteria | Legionellales |

| | | | | | | | | |
|---|---|---|---|---|---|---|---|---|
| | Paracaedibacteraceae | 15 | 1 | 6.7 | Bacteria | Proteobacteria | Alphaproteobacteria | Paracaedibacterales |
| | Bdellovibrionaceae | 31 | 2 | 6.5 | Bacteria | Bdellovibrionota | Bdellovibrionia | Bdellovibrionales |
| | Chthoniobacteraceae | 16 | 1 | 6.3 | Bacteria | Verrucomicrobiota | Verrucomicrobiae | Chthoniobacterales |
| | Rickettsiaceae | 17 | 1 | 5.9 | Bacteria | Proteobacteria | Alphaproteobacteria | Rickettsiales |
| | Sphingomonadaceae | 18 | 1 | 5.6 | Bacteria | Proteobacteria | Alphaproteobacteria | Sphingomonadales |
| | Crocinitomicaceae | 33 | 1 | 3.0 | Bacteria | Bacteroidota | Bacteroidia | Flavobacteriales |
| | betI | 77 | 1 | 1.3 | Bacteria | Proteobacteria | Gammaproteobacteria | Burkholderiales |
| | Comamonadaceae | 84 | 1 | 1.2 | Bacteria | Proteobacteria | Gammaproteobacteria | Burkholderiales |

**Table S6: Number of affected ASVs per family and their proportion in the compared to the total number of ASVs in the family in LLC.** Families are ordered by the proportion of ASVs affected and categorised as follows: eukaryotic nanoplankton (18S-NP), eukaryotic picoplankton (18S-PP) and prokaryotic picoplankton (16S-PP).

| Microbial plankton | Family | Total ASVs | Affected ASVs | Proportion | Kingdom | Phylum | Class | Order |
|---|---|---|---|---|---|---|---|---|
| 18S-NP | Anoplophryidae | 1 | 1 | 100.0 | Eukaryota | Ciliophora | Oligohymenophorea | Astomatida |
| | Ceratiaceae | 4 | 4 | 100.0 | Eukaryota | Myzozoa | Dinophyceae | Gonyaulacales |
| | Conchophthiridae | 4 | 4 | 100.0 | Eukaryota | Ciliophora | Oligohymenophorea | Pleuronematida |
| | Desmidiaceae | 1 | 1 | 100.0 | Eukaryota | Charophyta | Zygnemophyceae | Desmidiales |
| | Dimacrocaryonidae | 4 | 4 | 100.0 | Eukaryota | Ciliophora | Litostomatea | Haptorida |
| | Diplopsaliaceae | 4 | 4 | 100.0 | Eukaryota | Myzozoa | Dinophyceae | Peridiniales |
| | Epalxellidae | 1 | 1 | 100.0 | Eukaryota | Ciliophora | Odontostomatea | Odontostomatida |
| | Eremosphaeraceae | 3 | 3 | 100.0 | Eukaryota | Chlorophyta | Trebouxiophyceae | Chlorellales |
| | Goniodomataceae | 3 | 3 | 100.0 | Eukaryota | Myzozoa | Dinophyceae | Gonyaulacales |
| | Haptophryidae | 4 | 4 | 100.0 | Eukaryota | Ciliophora | Oligohymenophorea | Astomatida |
| | Holostichidae | 12 | 12 | 100.0 | Eukaryota | Ciliophora | Spirotrichea | Urostylida |
| | Ichthyophonidae | 1 | 1 | 100.0 | Eukaryota | Choanozoa | Ichthyosporea | Ichthyophonida |
| | Ichthyophthiriidae | 4 | 4 | 100.0 | Eukaryota | Ciliophora | Oligohymenophorea | Hymenostomatida |
| | Neochloridaceae | 4 | 4 | 100.0 | Eukaryota | Chlorophyta | Chlorophyceae | Sphaeropleales |
| | Oocystaceae | 3 | 3 | 100.0 | Eukaryota | Chlorophyta | Trebouxiophyceae | Chlorellales |
| | Protoperidiniaceae | 4 | 4 | 100.0 | Eukaryota | Myzozoa | Dinophyceae | Peridiniales |
| | Spirostomidae | 4 | 4 | 100.0 | Eukaryota | Ciliophora | Heterotrichea | Heterotrichida |
| | Thaumatomastigidae | 4 | 4 | 100.0 | Eukaryota | Cercozoa | Imbricatea | Thaumatomonadida |
| | Volvocaceae | 5 | 5 | 100.0 | Eukaryota | Chlorophyta | Chlorophyceae | Chlamydomonadales |
| | Peridiniaceae | 23 | 22 | 95.7 | Eukaryota | Myzozoa | Dinophyceae | Peridiniales |
| | Suessiaceae | 40 | 31 | 77.5 | Eukaryota | Myzozoa | Dinophyceae | Suessiales |
| | Sphaeropleaceae | 14 | 10 | 71.4 | Eukaryota | Chlorophyta | Chlorophyceae | Sphaeropleales |
| | Oxytrichidae | 6 | 4 | 66.7 | Eukaryota | Ciliophora | Spirotrichea | Sporadotrichida |
| | Saprolegniaceae | 17 | 11 | 64.7 | Eukaryota | Oomycota | Peronosporea | Saprolegniales |

| | | | | | | | | |
|---|---|---|---|---|---|---|---|---|
| | Chytriomycetaceae | 8 | 5 | 62.5 | Fungi | Chytridiomycota | Chytridiomycetes | Chytridiales |
| | Catenariaceae | 14 | 8 | 57.1 | Fungi | Blastocladiomycota | Blastocladiomycetes | Blastocladiales |
| | Chaetomellaceae | 2 | 1 | 50.0 | Fungi | Ascomycota | Leotiomycetes | Chaetomellales |
| | Codonellidae | 4 | 2 | 50.0 | Eukaryota | Ciliophora | Spirotrichea | Tintinnida |
| | Thraustochytriaceae | 2 | 1 | 50.0 | Eukaryota | Bigyra | Labyrinthulea | Thraustochytrida |
| | Uebelmesseromycetaceae | 8 | 4 | 50.0 | Fungi | Chytridiomycota | Chytridiomycetes | Rhizophydiales |
| | Cryptomonadaceae | 22 | 10 | 45.5 | Eukaryota | Cryptophyta | Cryptophyceae | Cryptomonadales |
| | Kappamycetaceae | 9 | 4 | 44.4 | Fungi | Chytridiomycota | Chytridiomycetes | Rhizophydiales |
| | Podolampaceae | 18 | 8 | 44.4 | Eukaryota | Myzozoa | Dinophyceae | Peridiniales |
| | Tintinnidae | 18 | 8 | 44.4 | Eukaryota | Ciliophora | Spirotrichea | Tintinnida |
| | Fragilariaceae | 25 | 11 | 44.0 | Eukaryota | Bacillariophyta | Bacillariophyceae | Fragilariales |
| | Closteriaceae | 5 | 2 | 40.0 | Eukaryota | Charophyta | Zygnemophyceae | Desmidiales |
| | Chlamydomonadaceae | 64 | 24 | 37.5 | Eukaryota | Chlorophyta | Chlorophyceae | Chlamydomonadales |
| | Bicosoecidae | 23 | 8 | 34.8 | Eukaryota | Bigyra | Bikosea | Bicosoecida |
| | Alphamonaceae | 10 | 3 | 30.0 | Eukaryota | Protozoa incertae sedis | Protozoa classis incertae sedis | Protozoa ordo incertae sedis |
| | Colepidae | 78 | 20 | 25.6 | Eukaryota | Ciliophora | Prostomatea | Prorodontida |
| | Dinobryaceae | 8 | 2 | 25.0 | Eukaryota | Ochrophyta | Chrysophyceae | Chromulinales |
| | Pfiesteriaceae | 16 | 4 | 25.0 | Eukaryota | Myzozoa | Dinophyceae | Peridiniales |
| | Prorocentraceae | 24 | 6 | 25.0 | Eukaryota | Myzozoa | Dinophyceae | Prorocentrales |
| | Mallomonadaceae | 51 | 12 | 23.5 | Eukaryota | Ochrophyta | Chrysophyceae | Synurales |
| | Chrysosphaerellaceae | 14 | 3 | 21.4 | Eukaryota | Ochrophyta | Chrysophyceae | Paraphysomonadales |
| | Scenedesmaceae | 5 | 1 | 20.0 | Eukaryota | Chlorophyta | Chlorophyceae | Sphaeropleales |
| | Protozoa familia incertae sedis | 27 | 5 | 18.5 | Eukaryota | Cercozoa | Protozoa classis incertae sedis | Protozoa ordo incertae sedis |
| | Spathidiidae | 53 | 9 | 17.0 | Eukaryota | Ciliophora | Litostomatea | Haptorida |
| | Chrysosphaeraceae | 18 | 3 | 16.7 | Eukaryota | Ochrophyta | Chrysophyceae | Chrysosphaerales |
| | Spongomonadidae | 6 | 1 | 16.7 | Eukaryota | Cercozoa | Imbricatea | Spongomonadida |
| | Protaspidae | 74 | 12 | 16.2 | Eukaryota | Cercozoa | Thecofilosea | Cryomonadida |
| | Strombidiidae | 25 | 4 | 16.0 | Eukaryota | Ciliophora | Spirotrichea | Oligotrichia |
| | Katablepharidaceae | 26 | 4 | 15.4 | Eukaryota | Cryptophyta | Cryptophyceae | Kathablepharidacea |
| | Stephanodiscaceae | 44 | 6 | 13.6 | Eukaryota | Bacillariophyta | Mediophyceae | Stephanodiscales |

| | | | | | | | | |
|---|---|---|---|---|---|---|---|---|
| | Gymnodiniaceae | 39 | 5 | 12.8 | Eukaryota | Myzozoa | Dinophyceae | Gymnodiniales |
| | Strombidinopsidae | 8 | 1 | 12.5 | Eukaryota | Ciliophora | Spirotrichea | Choreotrichida |
| | Chlorodendraceae | 10 | 1 | 10.0 | Eukaryota | Chlorophyta | Chlorodendrophyceae | Chlorodendrales |
| | Pseudophyllomitidae | 13 | 1 | 7.7 | Eukaryota | Bigyra | incertae sedis | incertae sedis |
| | Codonosigaceae | 18 | 1 | 5.6 | Eukaryota | Choanozoa | Choanoflagellata | Craspedida |
| | Ochromonadaceae | 20 | 1 | 5.0 | Eukaryota | Ochrophyta | Chrysophyceae | Ochromonadales |
| 18S-PP | Chaetomellaceae | 5 | 5 | 100.0 | Fungi | Ascomycota | Leotiomycetes | Chaetomellales |
| | Characiochloridaceae | 4 | 4 | 100.0 | Eukaryota | Chlorophyta | Chlorophyceae | Chlamydomonadales |
| | Dimacrocaryonidae | 4 | 4 | 100.0 | Eukaryota | Ciliophora | Litostomatea | Haptorida |
| | Diplopsaliaceae | 4 | 4 | 100.0 | Eukaryota | Myzozoa | Dinophyceae | Peridiniales |
| | Ebriidae | 1 | 1 | 100.0 | Eukaryota | Cercozoa | Thecofilosea | Ebriida |
| | Goniochloridaceae | 1 | 1 | 100.0 | Eukaryota | Ochrophyta | Eustigmatophyceae | Goniochloridales |
| | Goniodomataceae | 4 | 4 | 100.0 | Eukaryota | Myzozoa | Dinophyceae | Gonyaulacales |
| | Haptophryidae | 2 | 2 | 100.0 | Eukaryota | Ciliophora | Oligohymenophorea | Astomatida |
| | Oxytrichidae | 4 | 4 | 100.0 | Eukaryota | Ciliophora | Spirotrichea | Sporadotrichida |
| | Protoperidiniaceae | 4 | 4 | 100.0 | Eukaryota | Myzozoa | Dinophyceae | Peridiniales |
| | Radiococcaceae | 16 | 16 | 100.0 | Eukaryota | Chlorophyta | Chlorophyceae | Sphaeropleales |
| | Uebelmesseromycetaceae | 9 | 9 | 100.0 | Fungi | Chytridiomycota | Chytridiomycetes | Rhizophydiales |
| | Volvocaceae | 4 | 4 | 100.0 | Eukaryota | Chlorophyta | Chlorophyceae | Chlamydomonadales |
| | Holostichidae | 12 | 11 | 91.7 | Eukaryota | Ciliophora | Spirotrichea | Urostylida |
| | Ceratiaceae | 4 | 3 | 75.0 | Eukaryota | Myzozoa | Dinophyceae | Gonyaulacales |
| | Podolampaceae | 27 | 20 | 74.1 | Eukaryota | Myzozoa | Dinophyceae | Peridiniales |
| | Cryptomonadaceae | 21 | 15 | 71.4 | Eukaryota | Cryptophyta | Cryptophyceae | Cryptomonadales |
| | Sphaeropleaceae | 7 | 5 | 71.4 | Eukaryota | Chlorophyta | Chlorophyceae | Sphaeropleales |
| | Suessiaceae | 24 | 16 | 66.7 | Eukaryota | Myzozoa | Dinophyceae | Suessiales |
| | Catenariaceae | 15 | 8 | 53.3 | Fungi | Blastocladiomycota | Blastocladiomycetes | Blastocladiales |
| | incertae sedis | 19 | 10 | 52.6 | Eukaryota | incertae sedis | incertae sedis | Aquavolonida |
| | Chytriomycetaceae | 10 | 5 | 50.0 | Fungi | Chytridiomycota | Chytridiomycetes | Chytridiales |

| | | | | | | | | |
|---|---|---|---|---|---|---|---|---|
| | Closteriaceae | 4 | 2 | 50.0 | Eukaryota | Charophyta | Zygnemophyceae | Desmidiales |
| | Codonellidae | 2 | 1 | 50.0 | Eukaryota | Ciliophora | Spirotrichea | Tintinnida |
| | Developayellaceae | 4 | 2 | 50.0 | Eukaryota | incertae sedis | Developea | incertae sedis |
| | Saprolegniaceae | 4 | 2 | 50.0 | Eukaryota | Oomycota | Peronosporea | Saprolegniales |
| | Thraustochytriaceae | 2 | 1 | 50.0 | Eukaryota | Bigyra | Labyrinthulea | Thraustochytrida |
| | Tintinnidae | 15 | 7 | 46.7 | Eukaryota | Ciliophora | Spirotrichea | Tintinnida |
| | Bicosoecidae | 16 | 7 | 43.8 | Eukaryota | Bigyra | Bikosea | Bicosoecida |
| | Cercomonadidae | 30 | 12 | 40.0 | Eukaryota | Cercozoa | Sarcomonadea | Cercomonadida |
| | Alphamonaceae | 11 | 4 | 36.4 | Eukaryota | Protozoa incertae sedis | Protozoa classis incertae sedis | Protozoa ordo incertae sedis |
| | Goniomonadaceae | 33 | 12 | 36.4 | Eukaryota | Cryptophyta | Cryptophyceae | Cyathomonadacea |
| | Peridiniaceae | 11 | 4 | 36.4 | Eukaryota | Myzozoa | Dinophyceae | Peridiniales |
| | Gymnodiniaceae | 21 | 7 | 33.3 | Eukaryota | Myzozoa | Dinophyceae | Gymnodiniales |
| | Pirsoniaceae | 3 | 1 | 33.3 | Eukaryota | Oomycota | Hyphochytrea | Pirsoniales |
| | Polykrikaceae | 12 | 4 | 33.3 | Eukaryota | Myzozoa | Dinophyceae | Gymnodiniales |
| | Viridiraptoridae | 9 | 3 | 33.3 | Eukaryota | Cercozoa | Sarcomonadea | Glissomonadida |
| | Colepidae | 92 | 28 | 30.4 | Eukaryota | Ciliophora | Prostomatea | Prorodontida |
| | Cryptomycota familia incertae sedis | 7 | 2 | 28.6 | Fungi | Cryptomycota | Cryptomycota classis incertae sedis | Cryptomycota ordo incertae sedis |
| | Chrysosphaeraceae | 11 | 3 | 27.3 | Eukaryota | Ochrophyta | Chrysophyceae | Chrysosphaerales |
| | Prorocentraceae | 49 | 13 | 26.5 | Eukaryota | Myzozoa | Dinophyceae | Prorocentrales |
| | Pseudophyllomitidae | 25 | 6 | 24.0 | Eukaryota | Bigyra | incertae sedis | incertae sedis |
| | Thoracosphaeraceae | 5 | 1 | 20.0 | Eukaryota | Myzozoa | Dinophyceae | Thoracosphaerales |
| | Strombidiidae | 21 | 4 | 19.0 | Eukaryota | Ciliophora | Oligotrichea | Oligotrichida |
| | Protozoa familia incertae sedis | 27 | 5 | 18.5 | Eukaryota | Cercozoa | Protozoa classis incertae sedis | Protozoa ordo incertae sedis |
| | Amphidiniaceae | 22 | 4 | 18.2 | Eukaryota | Myzozoa | Dinophyceae | Amphidiniales |
| | Siluaniidae | 147 | 24 | 16.3 | Eukaryota | Bigyra | Bikosea | Bicosoecida |
| | Chlamydomonadaceae | 32 | 5 | 15.6 | Eukaryota | Chlorophyta | Chlorophyceae | Chlamydomonadales |
| | Chrysoamoebidaceae | 32 | 5 | 15.6 | Eukaryota | Ochrophyta | Chrysophyceae | Chromulinales |

| | | | | | | | | |
|---|---|---|---|---|---|---|---|---|
| | Spongomonadidae | 13 | 2 | 15.4 | Eukaryota | Cercozoa | Imbricatea | Spongomonadida |
| | Ochromonadaceae | 60 | 9 | 15.0 | Eukaryota | Ochrophyta | Chrysophyceae | Ochromonadales |
| | Codonosigaceae | 72 | 10 | 13.9 | Eukaryota | Choanozoa | Choanoflagellata | Craspedida |
| | Strombidinopsidae | 8 | 1 | 12.5 | Eukaryota | Ciliophora | Spirotrichea | Choreotrichida |
| | Crustomastigaceae | 25 | 3 | 12.0 | Eukaryota | Chlorophyta | Mamiellophyceae | Dolichomastigales |
| | Chlorellaceae | 19 | 2 | 10.5 | Eukaryota | Chlorophyta | Trebouxiophyceae | Chlorellales |
| | Mallomonadaceae | 71 | 7 | 9.9 | Eukaryota | Ochrophyta | Chrysophyceae | Synurales |
| | Sympoventuriaceae | 11 | 1 | 9.1 | Fungi | Ascomycota | Dothideomycetes | Venturiales |
| | Actinomonadaceae | 72 | 6 | 8.3 | Eukaryota | Ochrophyta | Dictyochophyceae | Pedinellales |
| | Spathidiidae | 16 | 1 | 6.3 | Eukaryota | Ciliophora | Litostomatea | Haptorida |
| | Paraphysomonadaceae | 105 | 3 | 2.9 | Eukaryota | Ochrophyta | Chrysophyceae | Paraphysomonadales |
| | Paraphysomonadaceae | 105 | 3 | 2.9 | Eukaryota | Ochrophyta | Chrysophyceae | Chromulinales |
| 16S-PP | Coxiellaceae | 2 | 2 | 100.0 | Bacteria | Proteobacteria | Gammaproteobacteria | Coxiellales |
| | BSV26 | 3 | 2 | 66.7 | Bacteria | Bacteroidota | Kryptonia | Kryptoniales |
| | Lentimicrobiaceae | 7 | 4 | 57.1 | Bacteria | Bacteroidota | Bacteroidia | Sphingobacteriales |
| | Rhodocyclaceae | 30 | 16 | 53.3 | Bacteria | Proteobacteria | Gammaproteobacteria | Burkholderiales |
| | 0319-6G20 | 6 | 3 | 50.0 | Bacteria | Bdellovibrionota | Oligoflexia | 0319-6G20 |
| | 053A03-B-DI-P58 | 2 | 1 | 50.0 | Bacteria | Bdellovibrionota | Oligoflexia | 053A03-B-DI-P58 |
| | 37-13 | 2 | 1 | 50.0 | Bacteria | Bacteroidota | Bacteroidia | Chitinophagales |
| | Paenibacillaceae | 2 | 1 | 50.0 | Bacteria | Firmicutes | Bacilli | Paenibacillales |
| | Prolixibacteraceae | 2 | 1 | 50.0 | Bacteria | Bacteroidota | Bacteroidia | Bacteroidales |
| | Streptococcaceae | 2 | 1 | 50.0 | Bacteria | Firmicutes | Bacilli | Lactobacillales |
| | Unknown_Family | 6 | 3 | 50.0 | Bacteria | Proteobacteria | Gammaproteobacteria | Gammaproteobacteria_Incertae_Sedis |
| | Gallionellaceae | 7 | 3 | 42.9 | Bacteria | Proteobacteria | Gammaproteobacteria | Burkholderiales |
| | Kapabacteriales | 7 | 3 | 42.9 | Bacteria | Bacteroidota | Kapabacteria | Kapabacteriales |
| | alfVIII | 5 | 2 | 40.0 | Bacteria | Proteobacteria | Alphaproteobacteria | Acetobacterales |
| | Magnetospirillaceae | 5 | 2 | 40.0 | Bacteria | Proteobacteria | Alphaproteobacteria | Rhodospirillales |
| | PB19 | 5 | 2 | 40.0 | Bacteria | Desulfobacterota | Desulfuromonadia | PB19 |
| | bacVI | 13 | 5 | 38.5 | Bacteria | Bacteroidota | Bacteroidia | Sphingobacteriales |

| | | | | | | | | |
|---|---|---|---|---|---|---|---|---|
| | Anaerolineaceae | 3 | 1 | 33.3 | Bacteria | Chloroflexi | Anaerolineae | Anaerolineales |
| | Hydrogenophilaceae | 3 | 1 | 33.3 | Bacteria | Proteobacteria | Gammaproteobacteria | Burkholderiales |
| | Micropepsaceae | 3 | 1 | 33.3 | Bacteria | Proteobacteria | Alphaproteobacteria | Micropepsales |
| | Chthoniobacteraceae | 16 | 5 | 31.3 | Bacteria | Verrucomicrobiota | Verrucomicrobiae | Chthoniobacterales |
| | Methylomonadaceae | 13 | 4 | 30.8 | Bacteria | Proteobacteria | Gammaproteobacteria | Methylococcales |
| | Gemmataceae | 7 | 2 | 28.6 | Bacteria | Planctomycetota | Planctomycetes | Gemmatales |
| | NS11-12_marine_group | 29 | 8 | 27.6 | Bacteria | Bacteroidota | Bacteroidia | Sphingobacteriales |
| | Crocinitomicaceae | 33 | 9 | 27.3 | Bacteria | Bacteroidota | Bacteroidia | Flavobacteriales |
| | Acidobacteriae | 4 | 1 | 25.0 | Bacteria | Acidobacteriota | Acidobacteriae | Acidobacteriae |
| | acSTL | 4 | 1 | 25.0 | Bacteria | Actinobacteriota | Actinobacteria | Frankiales |
| | Bacteroidaceae | 4 | 1 | 25.0 | Bacteria | Bacteroidota | Bacteroidia | Bacteroidales |
| | Ilumatobacteraceae | 8 | 2 | 25.0 | Bacteria | Actinobacteriota | Acidimicrobiia | Microtrichales |
| | NS9_marine_group | 12 | 3 | 25.0 | Bacteria | Bacteroidota | Bacteroidia | Flavobacteriales |
| | Roseiflexaceae | 4 | 1 | 25.0 | Bacteria | Chloroflexi | Chloroflexia | Chloroflexales |
| | Luna1 | 17 | 4 | 23.5 | Bacteria | Actinobacteriota | Actinobacteria | Micrococcales |
| | bacI | 35 | 8 | 22.9 | Bacteria | Bacteroidota | Bacteroidia | Chitinophagales |
| | Cryomorphaceae | 9 | 2 | 22.2 | Bacteria | Bacteroidota | Bacteroidia | Flavobacteriales |
| | Oxalobacteraceae | 9 | 2 | 22.2 | Bacteria | Proteobacteria | Gammaproteobacteria | Burkholderiales |
| | betII | 14 | 3 | 21.4 | Bacteria | Proteobacteria | Gammaproteobacteria | Burkholderiales |
| | Pedosphaeraceae | 19 | 4 | 21.1 | Bacteria | Verrucomicrobiota | Verrucomicrobiae | Pedosphaerales |
| | Babeliales | 5 | 1 | 20.0 | Bacteria | Dependentiae | Babeliae | Babeliales |
| | Gemmatimonadaceae | 10 | 2 | 20.0 | Bacteria | Gemmatimonadota | Gemmatimonadetes | Gemmatimonadales |
| | Rhodobacteraceae | 10 | 2 | 20.0 | Bacteria | Proteobacteria | Alphaproteobacteria | Rhodobacterales |
| | Saprospiraceae | 10 | 2 | 20.0 | Bacteria | Bacteroidota | Bacteroidia | Chitinophagales |
| | Xanthomonadaceae | 5 | 1 | 20.0 | Bacteria | Proteobacteria | Gammaproteobacteria | Xanthomonadales |
| | Phycisphaeraceae | 22 | 4 | 18.2 | Bacteria | Planctomycetota | Phycisphaerae | Phycisphaerales |
| | SAR324_clade(Marine_group_B) | 11 | 2 | 18.2 | Bacteria | SAR324_clade(Marine_group_B) | SAR324_clade(Marine_group_B) | SAR324_clade(Marine_group_B) |
| | env.OPS_17 | 12 | 2 | 16.7 | Bacteria | Bacteroidota | Bacteroidia | Sphingobacteriales |
| | Margulisbacteria | 6 | 1 | 16.7 | Bacteria | Margulisbacteria | Margulisbacteria | Margulisbacteria |
| | Methylophilaceae | 6 | 1 | 16.7 | Bacteria | Proteobacteria | Gammaproteobacteria | Burkholderiales |

| | | | | | | | | |
|---|---|---|---|---|---|---|---|---|
| | OM190 | 6 | 1 | 16.7 | Bacteria | Planctomycetota | OM190 | OM190 |
| | Bdellovibrionaceae | 31 | 5 | 16.1 | Bacteria | Bdellovibrionota | Bdellovibrionia | Bdellovibrionales |
| | acIV | 25 | 4 | 16.0 | Bacteria | Actinobacteriota | Acidimicrobiia | Microtrichales |
| | bacV | 39 | 6 | 15.4 | Bacteria | Bacteroidota | Bacteroidia | Flavobacteriales |
| | Cyanobiaceae | 26 | 4 | 15.4 | Bacteria | Cyanobacteria | Cyanobacteriia | Synechococcales |
| | uncultured | 52 | 8 | 15.4 | Bacteria | unknown | unknown | unknown |
| | Chitinophagaceae | 27 | 4 | 14.8 | Bacteria | Bacteroidota | Bacteroidia | Chitinophagales |
| | alfI | 7 | 1 | 14.3 | Bacteria | Proteobacteria | Alphaproteobacteria | Rhizobiales |
| | bacIII | 7 | 1 | 14.3 | Bacteria | Bacteroidota | Bacteroidia | Cytophagales |
| | Caulobacteraceae | 7 | 1 | 14.3 | Bacteria | Proteobacteria | Alphaproteobacteria | Caulobacterales |
| | Comamonadaceae | 84 | 12 | 14.3 | Bacteria | Proteobacteria | Gammaproteobacteria | Burkholderiales |
| | Sphingobacteriaceae | 7 | 1 | 14.3 | Bacteria | Bacteroidota | Bacteroidia | Sphingobacteriales |
| | Verrucomicrobiaceae | 15 | 2 | 13.3 | Bacteria | Verrucomicrobiota | Verrucomicrobiae | Verrucomicrobiales |
| | betI | 77 | 10 | 13.0 | Bacteria | Proteobacteria | Gammaproteobacteria | Burkholderiales |
| | Rhodospirillaceae | 8 | 1 | 12.5 | Bacteria | Proteobacteria | Alphaproteobacteria | Rhodospirillales |
| | bacII | 42 | 5 | 11.9 | Bacteria | Bacteroidota | Bacteroidia | Flavobacteriales |
| | Burkholderiaceae | 9 | 1 | 11.1 | Bacteria | Proteobacteria | Gammaproteobacteria | Burkholderiales |
| | Pseudohongiellaceae | 9 | 1 | 11.1 | Bacteria | Proteobacteria | Gammaproteobacteria | Oceanospirillales |
| | Neisseriaceae | 10 | 1 | 10.0 | Bacteria | Proteobacteria | Gammaproteobacteria | Burkholderiales |
| | Opitutaceae | 10 | 1 | 10.0 | Bacteria | Verrucomicrobiota | Verrucomicrobiae | Opitutales |
| | Rubritaleaceae | 20 | 2 | 10.0 | Bacteria | Verrucomicrobiota | Verrucomicrobiae | Verrucomicrobiales |
| | acI | 36 | 3 | 8.3 | Bacteria | Actinobacteriota | Actinobacteria | Frankiales |
| | Beijerinckiaceae | 13 | 1 | 7.7 | Bacteria | Proteobacteria | Alphaproteobacteria | Rhizobiales |
| | Nitrosomonadaceae | 14 | 1 | 7.1 | Bacteria | Proteobacteria | Gammaproteobacteria | Burkholderiales |
| | Paracaedibacteraceae | 15 | 1 | 6.7 | Bacteria | Proteobacteria | Alphaproteobacteria | Paracaedibacterales |
| | Rickettsiaceae | 17 | 1 | 5.9 | Bacteria | Proteobacteria | Alphaproteobacteria | Rickettsiales |
| | Sphingomonadaceae | 18 | 1 | 5.6 | Bacteria | Proteobacteria | Alphaproteobacteria | Sphingomonadales |
| | Legionellaceae | 29 | 1 | 3.4 | Bacteria | Proteobacteria | Gammaproteobacteria | Legionellales |
